# The Temporal Statistics of Musical Rhythm across Western Genres: An Amplitude Modulation Phase Hierarchy Model

**DOI:** 10.1101/2020.08.18.255117

**Authors:** Tatsuya Daikoku, Usha Goswami

## Abstract

Statistical learning by the human brain plays a core role in the development of cognitive systems like language and music. Both music and speech have structured inherent rhythms, however the acoustic sources of these rhythms are debated. Theoretically, rhythm structures in both systems may be related to a novel set of acoustic statistics embedded in the amplitude envelope, statistics originally revealed by modelling children’s nursery rhymes. Here we apply similar modelling to explore whether the amplitude modulation (AM) timescales underlying rhythm in music match those in child-directed speech (CDS). Utilising AM-driven phase hierarchy modelling previously applied to infant-directed speech (IDS), adult-directed speech (ADS) and CDS, we test whether the physical stimulus characteristics that yield speech rhythm in IDS and CDS describe rhythm in music. Two models were applied. One utilized a low-dimensional representation of the auditory signal adjusted for known mechanisms of the human cochlear, and the second utilized probabilistic amplitude demodulation, estimating the modulator (envelope) and carriers using Bayesian inference. Both models revealed a similar hierarchically-nested temporal modulation structure across Western musical genres and instruments. Core bands of AM and spectral patterning matched prior analyses of IDS and CDS, and music showed strong phase dependence between slower bands of AMs, again matching IDS and CDS. This phase dependence is critical to the perception of rhythm. Control analyses modelling other natural sounds (wind, rain, storms, rivers) did not show similar temporal modulation structures and phase dependencies. We conclude that acoustic rhythm in language and music has a shared statistical basis.

---

The potential parallels between language and music have long fascinated researchers in cognitive science. In this paper, we examine whether a statistical learning approach previously applied to understand the development of phonology as a cognitive system in language-learning infants and children may enable theoretical advances in understanding the acoustic basis of rhythm in music. Although language acquisition by human infants was once thought to require specialized neural architecture, studies of infant statistical learning have revealed that basic acoustic processing mechanisms are sufficient for infants to learn phonology (speech sound structure at different linguistic levels such as words, syllables, rhymes and phonemes; e.g. Saffran, 2001). Most statistical learning studies have focused on syllable and word learning, however infant language learning has been argued to begin with speech rhythm (Mehler et al., 1988). Children who exhibit difficulties with phonological learning also exhibit rhythm processing difficulties (Goswami, 2015, for review). Modelling of the speech signal aimed at understanding the potential sensory basis of these phonological and rhythmic difficulties has revealed a novel set of acoustic statistics that underpin speech rhythm in infant- and child-directed speech (IDS and CDS). These novel statistics were discovered by applying an amplitude modulation (AM) phase hierarchy modelling approach based on the neural speech encoding literature to children’s nursery rhymes and to baby talk (Leong & Goswami, 2015; Leong et al., 2017).

The theoretical framework underpinning this novel modelling approach was Temporal Sampling theory (TS theory, Goswami, 2011). TS theory was initially developed to provide a systematic sensory/neural/cognitive framework for explaining childhood language disorders, with a view to supporting musical remediation of such disorders (Bhide et al., 2013). TS theory is based on the perception of the amplitude envelope of speech, and proposes that accurate sensory/neural processing of the envelope is one foundation of language acquisition (Goswami, 2019a). The amplitude envelope of any sound is the slower changes in AM (intensity or signal energy) that unfold over time. By TS theory, phonological learning in infancy depends in part on automatic sensory learning of the statistical structure of the amplitude envelope. TS theory has led to the modelling of both IDS and rhythmic CDS (English nursery rhymes) in terms of patterns of AM in the amplitude envelope (Spectral-Amplitude Modulation Phase Hierarchy or S-AMPH modelling, see Leong, 2012; Leong & Goswami, 2015; Leong et al., 2014, 2017). This modelling revealed that an AM phase hierarchy between three broad bands of AMs (at rates of ∼2 Hz, 5 Hz and 20 Hz) described rhythm patterns in both IDS and CDS, with the phase relations between peaks and troughs in AM in the two slower bands being particularly important for perceiving metrical patterning (Leong et al., 2014, 2017). For example, in experimental work with adults, perceptual experience of a trochaic rhythm could be changed to an iambic rhythm by phase-shifting the AM peaks in the slowest (∼2 Hz) AM band by 180’, so that they now aligned with different AM peaks in the adjacent (∼5 Hz) AM band compared to their alignment prior to the phase shift (Leong et al., 2014). Theoretically, it is plausible that the physical stimulus characteristics that describe CDS and IDS may also describe the hierarchical rhythmic relationships that characterize modern music.

Rhythmic structure is a fundamental feature of both language and music. Rhythm involves sequences of events (such as syllables, notes, or drum beats) and these events have systematic patterns of timing, accent, and grouping (Patel, 2008). IDS, also called baby talk or Parentese, has been described as sing-song speech, and has particular prosodic or quasi-musical characteristics that have been suggested to explain both natural selection for human language from an anthropological perspective (Falk, 2004), and to facilitate infant learning of the phonological structure of human languages (Nazzi et al., 1998). The potential role of the physical structure of the amplitude envelope in children’s phonological learning can be investigated by separating the AM characteristics of speech from the frequency modulation (FM) characteristics. This is achieved by acoustic engineering methods for decomposing the amplitude envelope (called demodulation; Turner & Sahani, 2011). The AM patterns are associated with fluctuations in loudness or sound intensity, a primary acoustic correlate of perceived rhythm based on onset timing, beat, accent, and grouping. In contrast, the FM patterns can be interpreted as fluctuations in pitch and noise (Turner, 2010). Particularly relevant to an application to music, the demodulation approach to modelling IDS and CDS revealed (a) that the modulation peak in IDS is ∼2 Hz, in direct contrast to the modulation peak found for adult-directed speech (ADS, also modelled, ∼5 Hz) (Leong et al., 2017), and (b) that rhythmic patterning in IDS and CDS is represented in the phase relations of slower AM bands in the amplitude envelope, particularly those corresponding temporally to delta and theta bands in electroencephalography (EEG, Leong & Goswami, 2015). Prior analyses of the temporal modulation spectrum in Western musical genres have also revealed a peak at ∼2 Hz, matching the peak in IDS (Ding et al., 2017). When the S-AMPH modelling approach was applied to adult-directed speech (ADS), a different set of acoustic statistics was revealed (Leong et al., 2017; Araujo et al., 2018). The adult modelling showed that the amplitude phase relations in ADS do not foreground rhythm, but rather increase the salience of acoustic information related to phonemes and syllables (Araujo et al., 2018). This suggests that the different modulation statistics characterizing IDS versus ADS represent structural acoustic differences that are important for early language learning, which is known to utilise rhythm. The different statistical structure of ADS may reflect the acquisition of literacy, which is known to re-map phonology to reflect sound categories like phonemes that are highlighted by spelling systems (Ziegler & Goswami, 2005).

Music and language are ubiquitous in human societies (Mehr et al., 2020), but literacy is a relatively recent cultural acquisition. Accordingly, given that the temporal modulation structure of speech varies depending on whether an infant or an adult is being addressed, as well as with the literacy level of the speaker, we can ask which kind of human speech should be used as a basis for extracting the physical stimulus characteristics that are to be compared with music. It is notable that prior music/speech analyses across languages have emphasized *differences* regarding the temporal modulation structure of music and speech, but have depended on analyses of speech produced by highly literate adult speakers (Ding et al., 2017). Comparisons based on IDS and CDS may reveal similarities rather than differences. The strong AM phase relations of IDS in essence underpin its prosodic structure, which is thought to have played a key role in the pre- linguistic foundations of the protolanguage(s) evolved by early hominins (Falk, 2004). This prosodic structure appears to have arisen from hominid soothing routines, devised to settle infants placed on the ground during foraging behavior: “affectively positive, rhythmic melodies” (Falk, 2004). Intriguingly, current analyses of the lullabies sung by mothers to their infants across cultures reveals a beat rate of ∼2 Hz (120 beats per minute, bpm, see Trehub & Trainor, 1998), matching the modulation peak found in studies of IDS (Leong et al., 2017) as well as music (Ding et al., 2017). The strong phase synchronization of delta- and theta-rate AMs in IDS reflects stronger rhythmic synchronization and acoustic temporal regularity, contributing to the “sing-song” nature of Motherese remarked by Falk (2004). Human song, and possibly also human music across genres, may thus exhibit similar physical stimulus characteristics to IDS.

In principle, therefore, it can be argued that statistical comparisons between music and the rhythmic speech directed to infants and young children may be theoretically more appropriate than current comparisons between music and ADS (Ding et al., 2017).

Of course, music shares many commonalities with speech aside from rhythm at both an acoustic and cognitive level (Hayes, 1995; Tierney & Kraus, 2013; Schön et al., 2010; Peretz et al., 2015). However, the human peripheral auditory system responds to “broadband” sounds like speech or music in the same manner, by filtering the sounds into narrowband signals with a wide range of center frequencies (Moore, 2012). Like pre-verbal infants listening to spoken language, the music listener experiences a coherent perceptual signal with accentuation and rhythms that are captured by the overall amplitude envelope (the broadband sound). The music listener needs to identify discrete units to gain meaning, like musical notes and phrasing, analogous to infants needing to identify discrete units like syllables, words and syntactic phrases from the prosodic rhythm structure of IDS. This segmentation task is aided by the filtering that occurs in the human cochlea, and each of the resultant narrowband signals can be modelled as a rapidly oscillating carrier signal with a relatively slowly varying amplitude envelope (the cochlear filterbank model of human hearing, Moore, 2019; Glasberg & Moore, 1990; Elliott & Theunissen, 2009). In terms of parsing the signal into discrete units like musical notes and phrasing, the prior linguistic analyses have revealed that the systematic patterns of AM nested in the amplitude envelope of both IDS and CDS support the identification of discrete units like syllables. For example, when one particular AM cycle is assumed to match a particular speech unit, application of the S-AMPH to English nursery rhyme corpora identifies 72% of stressed syllables correctly, 82% of syllables correctly, and 78% of onset-rime units correctly (Leong & Goswami, 2015). If the nursery rhymes are chanted to a regular 2 Hz beat (a temporal rate also dominant in music, 120 bpm), then the model identifies over 90% of each type of linguistic unit correctly. Accordingly, decomposition of the amplitude envelope of different musical genres may identify similar hierarchical AM structures in spectral (pitch) bandings matching IDS and CDS. For music, such a phase hierarchy may provide a perceptual basis for perceiving rhythm patterns, musical notes and musical phrasing. Whether music would exhibit similar salient bands of AMs, similar spectral banding and similar phase dependencies between AM bands as CDS and IDS is explored here.

This theoretical question is addressed using two contrasting mathematical approaches to demodulation of the amplitude envelope of music. The same genres of Western music studied by Ding et al. (2017) were modelled using two different algorithms, the S-AMPH (Leong & Goswami, 2015), and PAD (Probabilistic Amplitude Demodulation, Turner & Sahani, 2011; Turner, 2010). Both models parse the amplitude envelope of the music signals into an hierarchy of AM bands, but the principles underpinning their operation are different. The S-AMPH simulates the frequency decomposition known to be carried out by the cochlea (Moore, 2012; Zeng et al., 2005; Dau et al., 1997b), thereby aiming to decompose the amplitude envelope of music in the same way as the human ear. PAD infers the modulators and carriers in the envelope based purely on Bayesian inference, thereby carrying out amplitude demodulation on a neutral statistical basis that makes no adjustments for the human hearing system. PAD is thus a “brain-neutral” approach. In our prior work with English nursery rhymes (Leong et al., 2014), both S-AMPH and PAD modelling showed that adult perception of linguistic rhythm patterns such as trochaic and iambic meters depended on the temporal alignment of modulation peaks in the delta- and theta-rate bands of AM. Accordingly, here we predict that musical meter may also depend primarily on the temporal alignment of modulation peaks in the delta- and theta-rate bands of AM, across musical genres. To test the specificity of this prediction regarding speech and music, we also modelled other quasi-rhythmic natural sounds such as wind and rain, using both the S-AMPH and PAD approaches.

Our prediction that the perception of musical meter may depend on the temporal alignment of different AM bands across musical genres also relates to linguistic theory (Liberman & Prince, 1977; Selkirk, 1984; Selkirk, 1980). Linguistically, hierarchical structures like the phonological hierarchy of prosodic, syllabic, rhyme and phoneme levels nested within speech rhythm are classically represented as a tree that captures the *relative prominence* of units (Hayes, 1995; Selkirk, 1980). Such tree representations may also provide a good model regarding the core principles of metrical structure in music (Lerdahl et al., 1983). In the tree representation, a “parent” node (element) at one tier of the hierarchy encompasses one or more “daughter” nodes at a lower level of the hierarchy. The adjacent connection between the parent and daughter nodes are indicated as “branches” in the tree. To give an example from CDS, a parent node such as the trisyllabic word “pussycat” in the nursery rhyme “Pussycat pussycat where have you been,” which is also the prosodic foot, would have 3 daughter nodes at the next hierarchical level, comprising the three syllables. By the S-AMPH model, the level of the prosodic foot would be captured by the cycles of AM at the delta-band (∼2 Hz) rate, while the individual syllables would be captured by the cycles of AM at the theta-band (∼5 Hz) rate. As noted, when modelled with the S-AMPH, English nursery rhymes with different metrical structures like “Jack and Jill went up the hill” (trochaic rhythm), “As I was going to St Ives” (iambic rhythm) and “Pussycat pussycat where have you been” (dactyl rhythm) all showed the same acoustic hierarchical AM structure. Which metrical structure was perceived by the listener depended on the temporal alignment of AM peaks in the delta- and theta-rate AM bands (Leong et al., 2014).

It should be noted that the terms “delta-rate” and “theta-rate” AM bands were adopted to describe the results of the speech demodulation analyses because TS theory was based in part on the neural oscillatory bands that track human speech in adult cortex (Luo & Poeppel, 2007; Ahissar et al., 2001; Giraud & Poeppel, 2012; Henry & Obleser, 2012; Overath et al., 2015; Ding et al., 2016; Park et al., 2015). These AM bands equate temporally to electrophysiological rhythms found across the brain at the oscillatory rates of delta, theta and beta-low gamma. In adult work, neural (“speech-brain”) alignment has been shown to contribute to parsing of the speech signal into phonological units such as syllables and words (Ding et al., 2016). It is known that human speech perception relies in part on neural tracking of the temporal modulation patterns in speech at different timescales simultaneously. These temporal modulation patterns are then bound into a single percept, “multi-time resolution processing” (Luo & Poeppel, 2007; Ahissar et al., 2001; Giraud & Poeppel, 2012; Poeppel, 2003). This neural tracking (also described as phase alignment, temporal alignment or entrainment) relies on oscillatory cortical activity. For both music and speech, oscillatory activity is known to align with selected rhythmic features of the input such as syllables or musical beats (Obleser & Kayser, 2019; Gross et al., 2013; Di Liberto et al., 2015; Baltzell et al., 2019; Fujioka et al., 2015). Indeed, if musical rhythms and language rhythms are designed to be identical in a particular stimulus set (achieved by creating matching hierarchical rhythmic structures in words in German sentences and Waltz-like 3-count musical pieces), then EEG recordings show similar phase-locked neural responses to both the speech and musical signals in listening adults (Harding et al., 2019). For language, delta, theta, and beta/gamma oscillators in auditory cortex contribute to the perception of prosodic, syllabic, and phonetic information respectively (Luo & Poeppel, 2007; Park et al., 2015; Fontolan et al., 2014). For music, possible correspondences between different oscillators and musical units has yet to be investigated.

Of core relevance to the demodulation modelling approach adopted here, there is also growing biological evidence that these adjacent-band neural oscillators are not independent of, but interdependent on, each other (Gross et al., 2013; Lakatos et al., 2005). For example, the phase of delta oscillators modulates the phase of theta oscillators, and theta phase modulates beta/gamma power (Gross et al., 2013). Accordingly, there are mechanistic phase dependencies in the neural system, which mirror the acoustic phase dependencies between AM bands revealed by the S-AMPH modelling of speech. Thus, musical rhythm may depend on the same acoustic phase dependencies. To date, despite a number of studies of music encompassing brain-based analyses (Doelling & Poeppel, 2015; Norman-Haignere et al., 2015; Nozaradan et al., 2011; Baltzell et al., 2019), no studies have examined the temporal correlates of musical rhythm from an hierarchical AM modelling perspective. As outlined above, it is biologically plausible to suggest that rhythm perception in music and language may depend on neural entrainment to the AM hierarchies nested in the amplitude envelope of music versus speech respectively. Most of the slow energy modulations within the speech amplitude envelope reflect intensity patterns associated with syllable production (Greenberg, 2006). However, within the overall speech envelope there are many amplitude envelopes of the different constituent (spectral) frequencies changing at different temporal rates, which can be modelled by decomposing the overall envelope. The resulting “temporal modulation spectrum” of speech has a relatively straightforward neurophysiological interpretation which could also apply to musical signals. The cochlea decomposes acoustic signals into narrow frequency bands (Moore, 2012), which are then low-pass filtered at different stages of the auditory pathway (Dau et al., 1997a), thereby extracting the temporal modulation envelope of the narrowband signals (Yang et al., 1992). Oscillatory cortical networks track the temporal modulation envelope < ∼40 Hz (Joris et al., 2004; Shamma, 2001), with different oscillators (delta: <4 Hz, theta: 4–8 Hz, alpha: 8–12 Hz, beta: 12–30 Hz, gamma: 30–80 Hz) phase-synchronizing with different AM patterns at matching rates. Amplitude “rise times” (time to modulation peak) are used as automatic triggers for this phase re-setting, so that amplitude envelopes at different frequencies in the signal become temporally aligned with neural oscillators at these frequencies (Doelling et al., 2014; Luo & Poeppel, 2007; Ahissar et al., 2001). These populations of auditory neurons thereby encode the modulation spectra of the temporal envelope at different neural signaling rates (Luo & Poeppel, 2007; Ahissar et al., 2001; Giraud & Poeppel, 2012; Henry & Obleser, 2012; Overath et al., 2015; Ding et al., 2016; Di Liberto et al., 2015; Barton et al., 2012; Santoro et al., 2014).

Regarding musical signals, it has already been shown that neural phase locking to periodic rhythms present in musical tempi is selectively enhanced compared to frequencies unrelated to the beat and meter (Nozaradan et al., 2011). However, to date the amplitude envelope of different musical inputs has not been decomposed in order to discover whether beat and meter are systematically related to adjacent bands of AMs that are physically connected by mutual phase dependencies. Aside from their mechanistic role in phase-resetting, amplitude rise times are important for the perception of rhythm because they determine the acoustic experience of “P-centers.” P-centers are the perceptual moment of occurrence (“perceptual center”) of each musical beat or syllable for the listener (Morton et al., 1976; Hoequist, 1983). Amplitude rise times are typically called attack times in the musical literature (Scott, 1993; Gordon, 1987). Perceiving the beat structure in both music and speech is known to be developmentally inter-related (Huss et al., 2011). Children’s prosodic perception and their musical rhythm perception are both related to individual differences in amplitude rise time perception, and children’s performance in a musical beat perception task uniquely predicts their performance on tests of phonological awareness, both concurrently and longitudinally (Huss et al., 2011; Goswami et al., 2013). As noted, amplitude rise times also provide sensory landmarks that *automatically* trigger brain rhythms and speech rhythms into temporal alignment, acting as acoustic landmarks that phase-reset ongoing neural oscillatory activity (Doelling et al., 2014).

Accordingly, exploring the physical characteristics of musical rhythm across Western musical genres and instruments by utilizing an AM phase hierarchy modelling approach is supported theoretically by neural, acoustic and developmental data. We hypothesized here that the two chosen models (filtering [S-AMPH] vs probabilistic [PAD]) would demonstrate the same core acoustic principles underlying the structure of musical rhythm. Given the biological evidence that each neural oscillator modulates the adjacent-band oscillator during speech perception (Gross et al., 2013; Lakatos et al., 2005), and our prior acoustic modelling data with IDS, CDS and ADS, we also hypothesized that the adjacent tiers in the temporal hierarchies of music would be highly dependent on each other compared with non-adjacent tiers, particularly for delta-theta AM coupling. By hypothesis, phase locking to different bands of AM present in the amplitude envelope of each genre may enable parsing of the signal to yield the perceptual experience of musical components such as minim, crotchet, and quaver (half, quarter, and eighth notes). The acoustic structure of the amplitude envelope should also contribute systematically to the perceptual experience of beat, tempo, and musical phrasing.

Note that our modelling approach is conceptually distinct from models that identify the tactus or beat markers in singing (Coath et al., 2010), models of pulse perception based on neural resonance (Large et al., 2019), and oscillatory models of auditory attention based on dynamic attending (Large & Jones, 1999) via its focus on decomposition of the amplitude envelope of the acoustic input. To test whether these predicted commonalities across speech and music would be unique to these two culturally-acquired systems, we also used both the PAD and S-AMPH models to examine mutual dependency in natural sounds with quasi-rhythmic content such as rain, wind, fire, storms and rivers. These natural sounds have also been present since early hominid times, but their statistical structure has not been constrained by the human brain.

## Materials and Methods

The music samples for modelling consisted of the music corpora used in the study by Ding et al. (Ding et al., 2017), with the addition of 23 children’s songs, in order to characterize more general properties of modulation spectra across musical genres. The final samples consisted of over 39 h of recordings (sampling rate = 44.1 kHz) of Western music (Western-classical music, Jazz, Rock, and Children’s song) and musical instruments (single-voice: Violin, Viola, Cello, and Bass; multi-voice: Piano and Guitar). In addition, a range of natural sounds like wind and rain were extracted from sound files available on the internet (https://mixkit.co; https://www.zapsplat.com). The full list is provided in S1 Appendix. The acoustic signals were normalized based on z-score (mean = 0, SD = 1). The spectro-temporal modulation of the signals was analyzed using two different algorithms for deriving the dominant amplitude modulation (AM) patterns: Spectral Amplitude Modulation Phase Hierarchy (S-AMPH; Leong et al., 2014), and Probability Amplitude Demodulation based on Bayesian inference (PAD; Large et al., 2019). The S-AMPH model is a low-dimensional representation of the auditory signal, using equivalent rectangular bandwidth (ERB_N_) filterbank, which simulates the frequency decomposition by the cochlea function in a normal human (Moore et al., 2012; Dau et al., 1997). This model can generate a hierarchical representation of the core spectral (acoustic frequency spanning 100–7,250 Hz) and temporal (oscillatory rate spanning 0.9–40 Hz) modulation hierarchies in the amplitude envelopes of speech and music. The number and the edge of bands are determined by principal component analysis (PCA) dimensionality reduction of original high-dimensional spectral and temporal envelope representations (Figure 1). This modulation filterbank can generate a cascade of amplitude modulators at different oscillatory rates, producing the AM hierarchy. On the other hand, the filterbank may partially introduce artificial modulations into the stimuli because the bandpass filters can introduce modulations near the center-frequency of the filter through “ringing.” Therefore, we also used a second AM-hierarchy extraction method (i.e., PAD) as a control. The PAD model does not implement the Hilbert transform, filtering, and PCA, but infers the modulators and a carrier based on Bayesian inference, identifying the envelope which best matches the data and the a priori assumptions. PAD can be run recursively using different demodulation parameters each time, which generates a cascade of amplitude modulators at different oscillatory rates (i.e., delta, theta, alpha, beta, and gamma), forming an AM hierarchy (Turner, 2010) (Figure 2).

**Figure 1.**
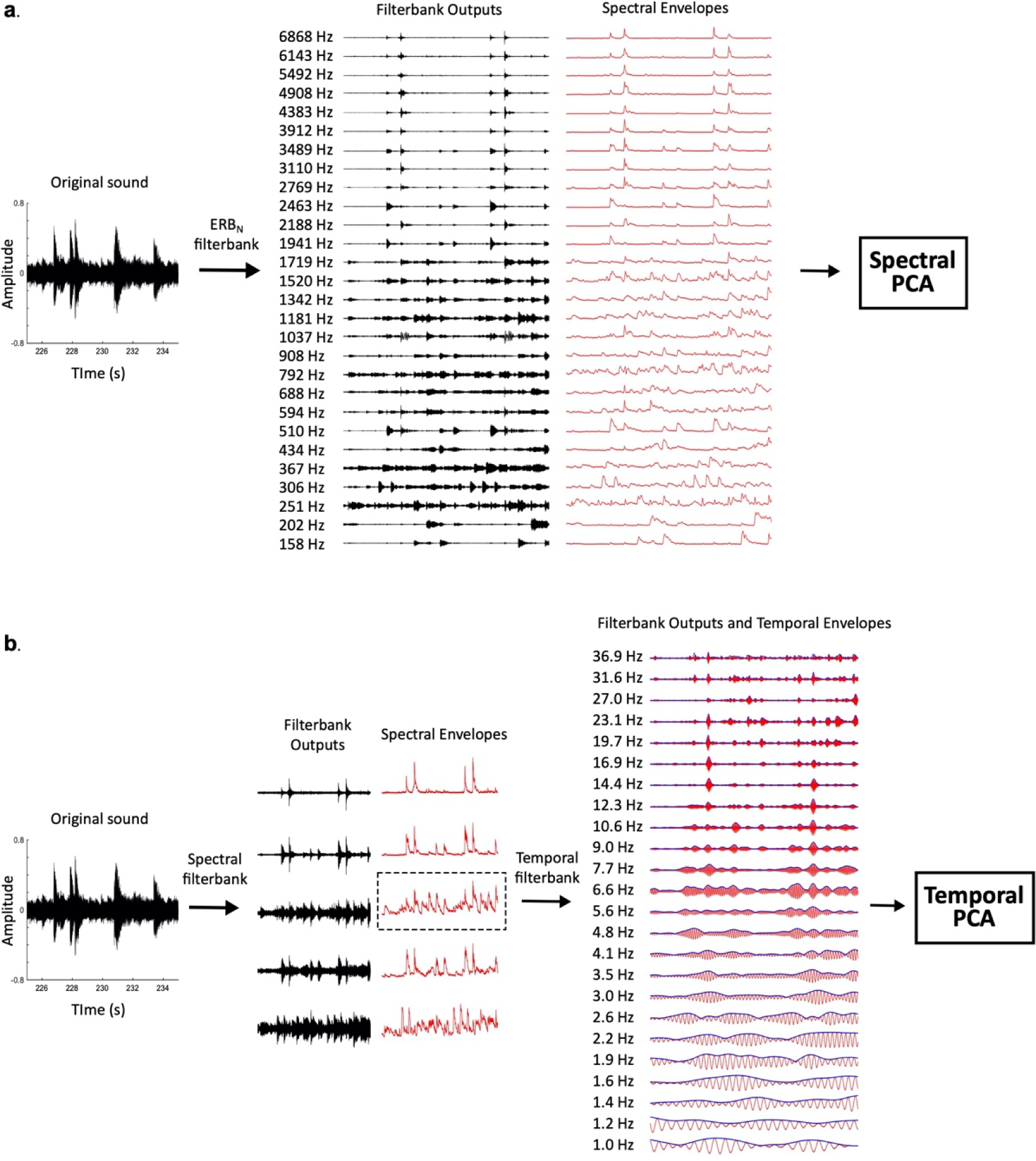
Signal Processing Steps in S-AMPH Model. *Note*. (**a**) The original sound is a part of a sample used in this study (Beethoven Piano Sonata No.14, Op27 No 2 “Moonlight”). Original sound is passed through an ERB_N_-spaced filterbank, yielding a set of high-dimensional spectral channel outputs. The envelope is extracted from each spectral channel output using the Hilbert transform, and these envelopes are entered into the spectral PCA to identify patterns of covariation across spectral channels. (**b**) The original sound is passed through a low-dimensional spectral filterbank, yielding a small set of core spectral band outputs. The parameters of the low-dimensional spectral filterbank were determined in the Spectral PCA procedure (**a**). The envelopes are extracted from each spectral band output using the Hilbert transform. Each envelope is further passed through a high-dimensional modulation filterbank, yielding a set of high-dimensional modulation rate envelopes. This rate-filtering is performed for each spectral band envelope, but for simplicity, only the modulation rate envelopes from a single spectral band are shown in this figure. Finally, the power profiles of the modulation rate envelopes (bold blue line) are entered into a temporal PCA to identify patterns of covariation across modulation rates.

**Figure 2.**
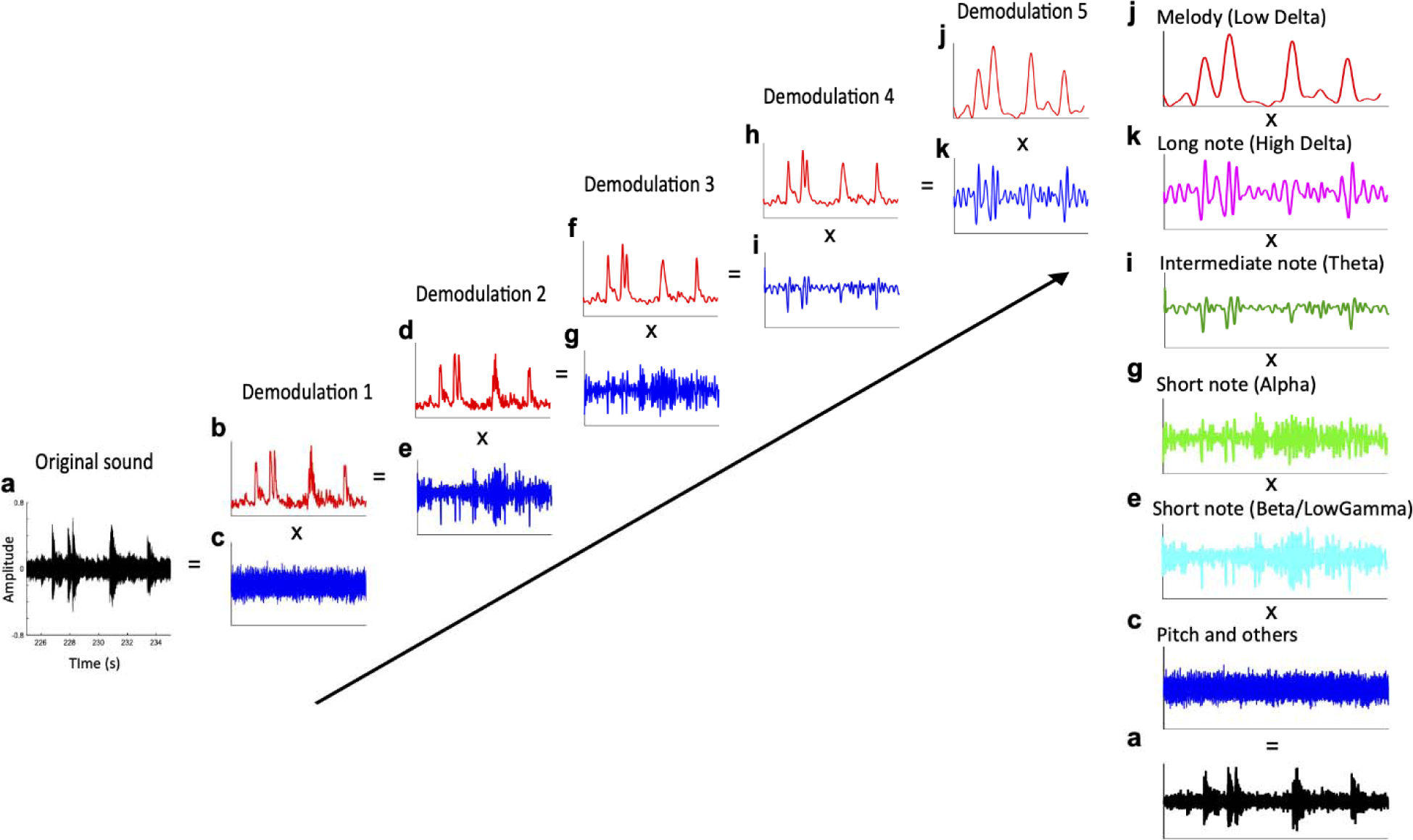
Signal Processing Steps in PAD Model. *Note*. Example of an amplitude modulation (AM) hierarchy derived by recursive application of PAD. In the first demodulation round (left column), the data (**a**) are demodulated using PAD set to a fast time scale. This yields a relatively quickly-varying envelope (**b**) and a carrier (**c**). In the second demodulation round (middle column), the demodulation process is re-applied to the extracted envelope b, using a slower time scale than before. This yields a slower daughter envelope (**d**) and a faster daughter envelope (**e**). Daughter envelopes d and e form the two tiers of the resulting AM hierarchy (right column). Mathematically, these two tiers (**d** and **e**) can be multiplied back with the very first carrier (c, bottom left) to yield the original signal (**a**).

### Spectral Amplitude Modulation Phase Hierarchy (S-AMPH) Model

#### Signal Processing: Spectral and Temporal Modulations

The methodologies were based on a previous study by Leong and Goswami (Leong & Goswami, 2015). To establish the patterns of spectral modulation, the raw acoustic signal was passed through a 28 log-spaced ERB_N_ filterbank spanning 100–7250 Hz, which simulates the frequency decomposition by the cochlea in a normal human (Moore, 2012; Dau et al., 1997b). For further technical details of the filterbank design, see Stone and Moore (Stone & Moore, 2003). The parameters of the ERB_N_ filterbanks and the frequency response characteristics are provided in S2 Appendix. Then, the Hilbert envelope was obtained for each of the 28 filtered-signals. Using the 28 Hilbert envelopes, the core spectral patterning was defined by PCA. This can identify the appropriate number and spacing of non-redundant spectral bands, by detecting co-modulation in the high-dimensional ERB_N_ representation. To establish the patterns of temporal modulation, the raw acoustic signal was filtered into the number of spectral bands that were identified in the spectral PCA analysis. Then, the Hilbert envelope was extracted from each of the spectral bands. Further, the Hilbert envelope of each of the spectral bands were passed through a 24 log-spaced ERB_N_ filterbank spanning 0.9–40 Hz. Using the 24 Hilbert envelopes in each of the spectral bands, the core AM hierarchy was defined by PCA. This can clarify co-activation patterns across modulation rate channels

To determine the number and the edge of the core spectral (acoustic frequency spanning 100–7,250 Hz) and temporal (oscillatory rate spanning 0.9–40 Hz) modulation bands, PCA was applied separately for spectral and temporal dimensionality reductions. PCA has previously been used for dimensionality reduction in speech studies (e.g., Klein et al., 1970; Pols et al., 1973). The present study focused on the absolute value of component loadings rather than the component scores. The loadings indicate the underlying patterns of correlation between high-dimensional channels. That is, PCA loading was adopted to identify patterns of covariation between the high-dimensional channels of spectral (28 channels) and temporal (24 channels) modulations, and to determine groups (or clusters) of channels that belonged to the same core modulation bands.

#### PCA to Finding the Core Modulation Hierarchy in the High-dimensional ERB_N_ Representation

In spectral PCA, the 28 spectral channels were taken as separate variables, yielding a total of 28 principal components. Only the top 5 principal components (PC) were considered for the further analysis, because these already cumulatively accounted for over 58% (on average) of the total variance in the original sound signal. In temporal PCA, the 24 channels in each of the spectral bands were entered as separate variables. Only the top 3 were considered for further analysis, because these cumulatively accounted for over 55% of the total variance in the original sound signal. Each of PC loading value was averaged across all samples in each genres (Western-classical music, Jazz, Rock, and Children’s song) and musical instruments (single-voice: Violin, Viola, Cello, and Bass; multi-voice: Piano and Guitar). The absolute value of the PC loading were used to avoid mutual cancellation by averaging an opposite valence across samples (Leong et al., 2014). Then, peaks in the grand average PC loading patterns were taken to identify the core modulation hierarchy. Troughs were also identified because they reflect boundaries of edges between co-modulated clusters of channels. To ensure that there would be an adequate spacing between the resulting inferred modulation bands, a minimum peak-to-peak distance of 2 and 5 channels was set for the spectral and temporal PCAs, respectively. After detecting all the peaks and troughs, the core spectral and temporal modulation bands were determined based on the criteria that at least 2 of the 5 PCs and 1 of the 3 PCs showed a peak for spectral and temporal bands, respectively. On the other hand, the boundary edges between modulation bands were determined based on the most consistent locations of “*flanking*” troughs for each group of PC peaks that indicated the presence of a band. The detailed methodologies and examples are shown in Leong and Goswami (Leong & Goswami, 2015).

#### Probability Amplitude Demodulation (PAD) Model Based on Bayesian Inference

Amplitude demodulation is the process by which a signal (y_t_) is decomposed into a slowly-varying modulator (m_t_) and quickly-varying carrier (c_t_):

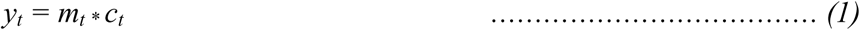

Probabilistic amplitude demodulation (PAD) (Turner & Sahani, 2011) implements the amplitude demodulation as a problem of learning and inference. Learning corresponds to the estimation of the parameters that describe these distributional constraints such as the expected time-scale of variation of the modulator. Inference corresponds to the estimation of the modulator and carrier from the signals based on the learned or manually defined parametric distributional constraints. This information is encoded probabilistically in the likelihood: *P(y_1:T_|c_1:T_, m_1:T_,θ)*, prior distribution over the carrier: *p(*c*_1:T_|θ)*, and prior distribution over the modulators: *p(m_1:T_|θ)*. Here, the notation x_1:T_ represents all the samples of the signal x, running from 1 to a maximum value T. Each of these distributions depends on a set of parameters θ, which controls factors such as the typical time-scale of variation of the modulator or the frequency content of the carrier. For more detail, the parametrized joint probability of the signal, carrier and modulator is:

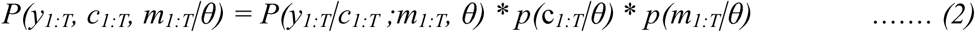

Bayes’ theorem is applied for inference, forming the posterior distribution over the modulators and carriers, given the signal:

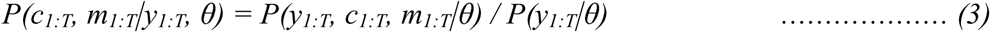

The full solution to PAD is a distribution over possible pairs of modulator and carrier.

The most probable pair of modulator and carrier given the signal is returned:

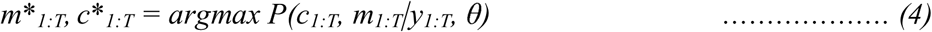

That is, compared with S-AMPH, PAD does not implement the Hilbert transform, filtering, and PCA, but estimates the most appropriate modulator (envelope) and carrier based on Bayesian inference that identifies the envelope which best matches the data and the a priori assumptions (Turner & Sahani, 2011; Turner, 2010). The solution takes the form of a probability distribution which describes how probable a particular setting of the modulator and carrier is, given the observed signal. Thus, PAD summarizes the posterior distribution by returning the specific envelope and carrier that have the highest posterior probability and therefore represent the best match to the data. As noted, PAD can be run recursively using different demodulation parameters each time, thereby generating a cascade of amplitude modulators at different oscillatory rates (Turner, 2010). The positive slow envelope is modelled by applying an exponential nonlinear function to a stationary Gaussian process. This produces a positive-valued envelope whose mean is constant over time. The degree of correlation between points in the envelope can be constrained by the timescale parameters of variation of the modulator (envelope), which may either be entered manually or learned from the data. In the present study, we manually entered the PAD parameters to produce the modulators at each of five tiers of oscillatory band (i.e., delta: -4 Hz, theta: 4-8 Hz, alpha: 8-12 Hz, beta: 12-30 Hz, and gamma: 30-50 Hz). Note that manual entry of these parameters does not predetermine the results, rather it enables exploration of whether there is a prominent peak frequency observed in each oscillatory rate band regardless of any tempo variations (such as speeding up or slowing down) that may depend on the performer or the particular music. Accordingly, it determines the frequencies that comprise the core temporal modulation structure of each musical genre. The carrier is interpreted as components including noise and pitches whose frequencies are much higher than the core modulation bands in phrase, prosodic, syllabic, phonological components. In each of music samples, the modulators (envelopes) of the five oscillatory bands were converted into the frequency domains by the Fast Fourier Transform (FFT). Spectral analysis of the modulator reflects how fast sound intensity fluctuates over time. High modulation frequency corresponds to fast modulations and vice versa (see Figure 2). The modulation spectra were averaged across all samples in each genre (Western-classical music, Jazz, Rock, and Children’s song) and musical instruments (single-voice: Violin, Viola, Cello, and Bass; multi-voice: Piano and Guitar).

### Mutual Information Between Different Modulation Bands

We also examined whether one tier of the temporal hierarchy of music may be mutually dependent on the timing of another tier by conducting mutual information analyses. Mutual information is a measure of the mutual dependence between the two variables. The mutual information can also be expressed as

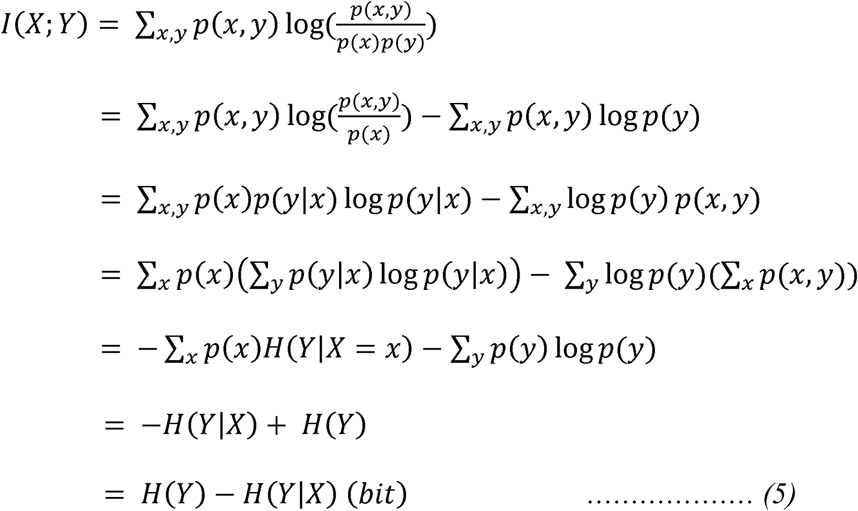

where p(x,y) is the joint probability function of X and Y, p(x) and p(y) are the marginal probability distribution functions of the X and Y respectively, H(X) and H(Y) are the marginal entropies, H(X|Y) and H(Y|X) are the conditional entropies, and (X,Y) is the joint entropy of X and Y (Daikoku, 2018).

This analysis could answer a question on how a certain oscillatory rhythm X (i.e., delta, theta, alpha and beta) is dependent on another oscillatory rhythm Y. Previous evidence suggests that adjacent-band oscillators are not independent of, but interdependent on each other (Gross et al., 2013; Lakatos et al., 2005). Given the evidence, we hypothesize that the adjacent tiers that can connect via so-called “*branches*” in the AM hierarchy are mutually dependent on each other compared with non-adjacent tiers. If so, the results may support a hierarchical “*tree-based*” structure of musical rhythm, highlighting the applicability of an AM hierarchy to music as well as speech.

To explore this, we adopted the phase angle “θ” of the core temporal modulation envelopes corresponding to delta, theta, alpha and beta/gamma waves, which was detected in each of S-AMPH and PAD approaches. That is, in S-AMPH model, the 5 spectral envelopes (see Figure 1b) were passed through a second series of band-pass filters to isolate the 4 different AM bands based on the results of temporal PCA (channel edge frequencies: 0.9, 2.5, 7, 17 and 30 Hz). The phase angles were then calculated using each of the 4x5 temporal modulation envelopes. In the PAD model, the phase angles were calculated using the four core modulators (envelopes) that have been detected in the last analyses, which correspond to delta, theta, alpha, and beta/gamma bands, respectively (for example, k, I, g, e in Figure 2). Then, using the phase angle values, the mutual information between different temporal modulation bands was measured.

#### Phase Synchronization Analyses

Based on the findings of mutual information, we further investigated possible multi-timescale phase synchronization between bands by computing the integer ratios between “*adjacent*” AM hierarchies (i.e., the number of parent vs. daughter elements in an AM hierarchy). This analysis addresses how many daughter elements a parent element encompasses in general in a particular musical genre. We adopted the core temporal modulation envelopes corresponding to delta, theta, alpha and beta/gamma waves, which was detected in each of S-AMPH and PAD approaches. That is, in S-AMPH model, the five spectral envelopes (see Figure 1b) was passed through a second series of band-pass filters to isolate the four different AM bands based on the results of temporal PCA (channel edge frequencies: 0.9, 2.5, 7, 17 and 30 Hz). In the end, the total numbers were 4x5 temporal modulation envelopes in the S-AMPH model. In contrast, in PAD model, we made use of the four core modulators (envelopes) corresponding to delta, theta, alpha, and beta/gamma bands, respectively (for example, k, I, g, e in Figure 2).

The Phase Synchronization Index (PSI) was computed between adjacent AM bands in the S-AMPH representation for each of the five spectral bands and in the PAD representation (i.e., delta vs. theta, theta vs. alpha, alpha vs. beta, beta vs. gamma phase synchronizations). The n:m PSI was originally conceptualized to quantify phase synchronization between two oscillators of different frequencies (e.g., muscle activity; Tass et al., 1998), and was subsequently adapted for neural analyses of oscillatory phase-locking (Schack & Weiss, 2005). For example, if the integer ratio is 1:2, then the parent element encompasses 2 daughter elements for the rhythm. The PSI was computed as:

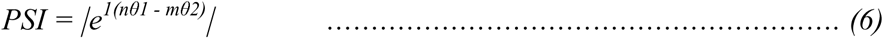

n and m are integers describing the frequency relationship between lower and higher AM bands, respectively. An n : m ratio for each PSI was defined as n & m < 10, and 1 < n/m < 3. The values θ1 and θ2 refer to the instantaneous phase of the two AMs at each point in time. Therefore, (nθ1–mθ2) is the generalized phase difference between the two AMs, which was computed by taking the circular distance (modulus 2π) between the two instantaneous phase angles. The angled brackets denote averaging of this phase difference over all time-points. The PSI is the absolute value of this average, and can take values between 0 and 1 (i.e., no to perfect synchronizations) (Leong et al., 2017). A sound with a PSI of 1 is perceived as being perfectly rhythmically regular (a repeating pattern of strong and weak beats), whereas a sound with a PSI of 0 is perceived as being random in rhythm.

To see if the detected findings properly represent systematic characteristics of natural musical rhythm, we further conducted simulation analyses. We generated synthesized sounds that consisted of four temporal modulation envelopes (i.e., modulator) and one spectral frequency (carrier). That is, 2 Hz, 4 Hz, 8 Hz and 16 Hz sine waves were summarized to synthesize one compound tone waveform. The compound tone waveform was, then, multiplied by a 200 Hz sine waves. The synthesized waveform was assumed as a sound that includes temporal information of delta, theta, alpha and gamma rhythms, and spectral information of a pitch around to natural human voices. It is important to note that all of the temporal envelopes comprised of simple sine waves with frequencies of a power of 2. Hence, we can hypothesize that 1:2 integer ratios should clearly and consistently be appeared compared with other integer ratios. If the PSIs of natural music show different findings from those of artificial sounds, then the results may indicate that natural musical rhythm has covert and systematic integer ratios in an AM hierarchy.

## Results

### Amplitude Modulation Properties of Western Music from S-AMPH

#### Spectral PCA

S3 Appendix shows the grand average as well as the loading patterns and cumulative contribution ratios for each music genre and instrument. The first to fifth principal component (PC1 to PC5) accounted for, on average, 33%, 8%, 6%, 5%, and 5% of the total variances, respectively. One peak (∼3000 Hz) was identified from the loading pattern of PC1. Thus, PC1 was assumed to reflect the global correlation between spectral channels. The peak of ∼300 Hz and the “*flanking*” trough of ∼350 Hz were identical between PC3 and PC4, providing corroborating evidence for a lowest spectral band at this spectral location with a potential boundary between the first and second spectral bands at ∼350 Hz (troughs indicate potential boundaries between modulation rate bands). Further peaks and troughs were identified providing evidence for four further spectral bands, please see S3 Appendix and Methods. Based on the a priori criteria (see Methods), the spectral PCA thus provided evidence for the presence of 5 core spectral bands in the spectral modulation data (300, 500, 1000, 2500 and 5500 Hz), with at least 2 out of 5 PCs showing peaks in each of these 5 spectral regions. Furthermore, we consistently observed 4 boundaries between these 5 spectral bands (350, 700, 1750 and 3900 Hz). Table a in the S3 Appendix provides a summary of these 5 spectral bands and their boundaries. It is noteworthy that these 5 spectral bands, which were consistent across musical instruments and the human voice, are proportionately-scaled with respect to the logarithmic frequency sensitivity of human hearing. As predicted, these results are similar to the spectral bands previously revealed by modelling IDS and CDS (Leong et al., 2017; Leong et al., 2014; Leong & Goswami, 2015). It can also be noted that the loading patterns for the 5 PCA components showed roughly similar characteristics across the genres. There was some individual variation at each spectral modulation band, see Appendix S3.

### Temporal PCA

S3 Appendix also shows the temporal loading patterns and cumulative contribution ratios for each music genre and each instrument, while Figure 3 shows the grand average loading patterns (absolute value) for the first three principal components arising from the temporal PCA of each of the 5 spectral bands determined in the spectral PCA (Table a in Appendix S3). The colors in Figure 3 represent the 5 spectral bands, while the types of lines (i.e., bold, dashed and dotted) represent PCs 1,2 and 3, respectively. As may be clearly observed, the loading showed consistent patterns between the 3 PCA loading patterns.

The first to third principal component (PC1 to PC3) accounted for, on average, 49%, 11% and 6% of the total variances, respectively (for more detail, see Table b in S3 Appendix). PC1 showed a moderate peak at acoustic frequencies of 7-9 Hz in all of the 5 spectral bands. As observed in the spectral PCA, PC1 in the temporal PCA might reflect the global correlation between temporal channels. As no troughs were detected in PC1 (indicating no potential boundaries), our analysis focused on PC2 and PC3. The loading patterns of PC2 resulted in 2 strong peaks at acoustic frequencies of 1-2 Hz (evidence for a delta-rate band of AMs) and 20-30 Hz (evidence for a beta-gamma rate band of AMs), and 1 strong flanking trough at acoustic frequencies of ∼7 Hz. These findings were consistent between the 5 spectral bands, suggesting the potential existence of at least 2 core temporal bands. Compared with PC1 and PC2, PC3 loading patterns varied across spectral bands. As detected in PC2, all the spectral bands showed a peak in loading at ∼30 Hz, and spectral band 2 similarly showed a peak at ∼1 Hz. PC3 also showed an additional mid-rate peak at ∼5 Hz (theta-rate band) and ∼9 Hz (alpha-rate band). The flanking troughs for these peaks occurred 2-3 Hz and 15-18 Hz. Based on the a priori criteria (Methods), the temporal PCA thus provided evidence for the presence of 4 core bands with 3 boundaries across the different musical genres and instruments (see Table 1). Perceptually, cycles in these AM bands may yield the experience of crotchets, quavers, demiquavers and onsets, as shown in Table 1.

These AM bands in music matched those previously found in IDS, but the AM bands in the nature sounds did not (PC3 in Figure 4). In particular, the strong peaks in the delta and theta bands, along with the strong flanking trough between these bands, are clearly visible for music and speech compared with the nature sounds. As predicted, therefore, the results of the temporal PCA match prior studies of CDS and IDS (Leong et al., 2017; Leong et al., 2014; Leong & Goswami, 2015).

**Figure 3.**
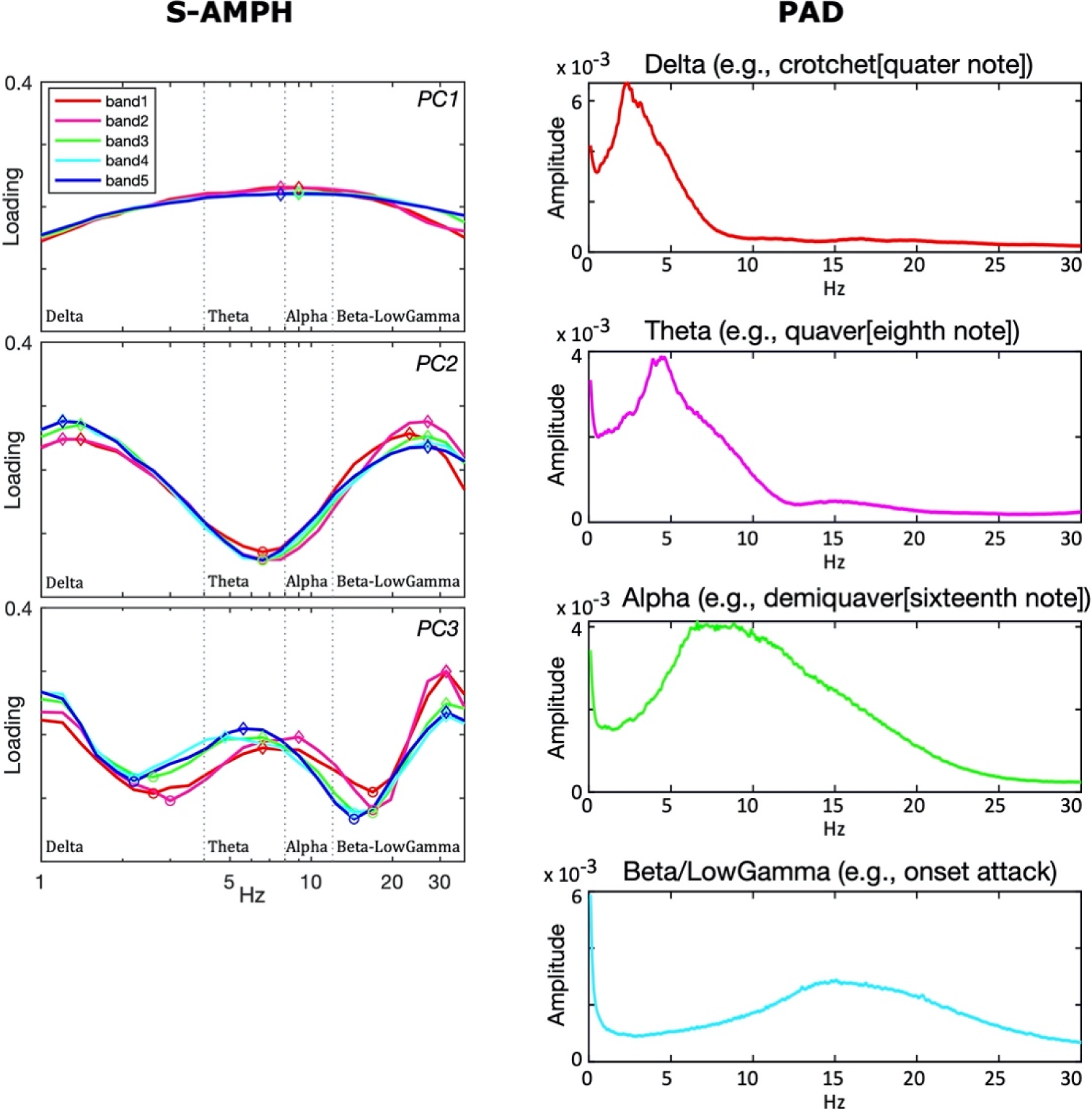
Core Temporal Modulation Rates. *Note*. Grand average absolute value of temporal PCA component loading patterns in the S-AMPH (a) model and modulation spectra of FFT in the PAD model (b). Both models showed an amplitude modulations (AM) hierarchy that consisted of delta-, theta-, alpha- and beta-rate AM bands.

**Figure 4.**
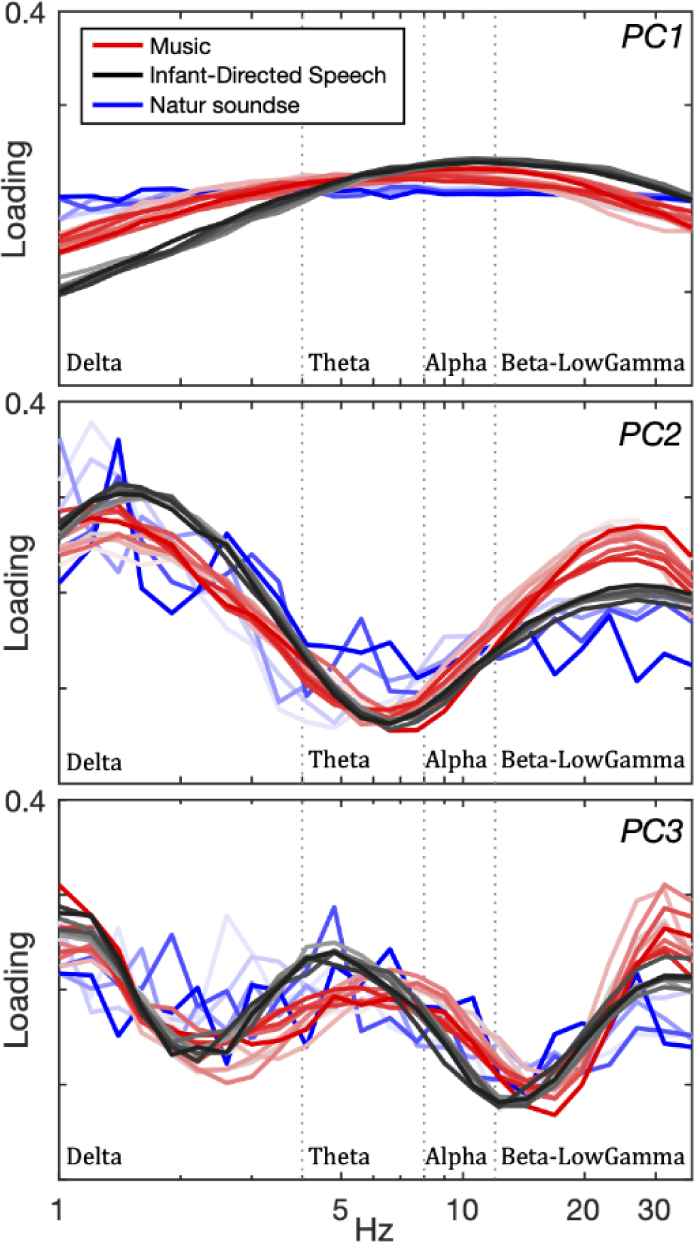
Core Temporal Modulation Rates of Music, Speech, and Nature Sounds. *Note*. Grand average absolute value of temporal PCA component loading patterns in the S-AMPH. Red, grey, and blue color scales represent music, speech, and nature sounds, respectively. Individual lines represent different speakers, musical genres and nature sounds.

**Table 1.** Summary of the 4 Temporal Bands Identified from PCA by Both S-AMPH and PAD.

| Temporal bands | S-AMPH | PAD | Neural Oscillatory Rates | Note value |
| --- | --- | --- | --- | --- |
| Band 1 | 0.9-2.5 Hz | -4 Hz | Delta | Crotchet (quarter note) |
| Band 2 | 2.5-7 Hz | 4-8 Hz | Theta | Quaver (eighth note) |
| Band 3 | 7-17 Hz | 8-12 Hz | Alpha | Demiquaver (sixteenth note) |
| Band 4 | 17-40 Hz | 12-30 Hz | Beta/LowGamma | Onset attack |

### Amplitude Modulation Properties of Music from PAD

PAD ignores the logarithmic frequency sensitivity of human hearing, implementing amplitude demodulation by estimating the most appropriate modulator (envelope) and carrier based on Bayesian inference (for more detail, see Methods). Accordingly, PAD provides a good test of the assumption that there is a systematic hierarchy of temporal modulations underpinning both Western music and (English) IDS. Further, PAD is exempt from the possibility that the filterbank used in the S-AMPH may have partially introduced artificial modulations into the stimuli through “ringing”. We had hypothesized that modelling with PAD should reveal the same core principles as the S-AMPH modelling regarding hierarchical AM patterns.

S3 Appendix shows the grand average for the modulation spectra of FFT as well as the loading patterns and cumulative contribution ratios for each music genre and instrument along with individual variation. Overall, low spectral frequencies (∼5Hz) showed higher consistency across musical genres and instruments while mid-to-high spectral frequencies showed some individual variation, as also found for S-AMPH. Figure 3 represents the grand average modulation spectra of FFT for each oscillatory band. Four peak frequencies were detected at ∼2.4 Hz, ∼4.8 Hz, ∼9 Hz and 16 Hz. Accordingly, PAD yielded similar findings to the S-AMPH model, detecting peak frequencies in AM bands corresponding temporally to neural delta, theta, alpha and beta/gamma neural oscillatory bands (see Table 1). As predicted, the modelling suggests the same core principles of AM structure across musical genres as the S-AMPH model. Accordingly, the AM structure of music shown in Table 1 is found irrespective of the modelling approach adopted.

### Mutual Information in Both Models

To examine whether mutual dependencies between AM bands in the temporal modulation structure of different musical genres was more similar to the dependencies identified in IDS rather than in ADS, a mutual information analysis method was employed. As noted earlier, prior modelling of IDS and CDS has revealed a significantly higher phase dependency between delta- and theta-rate AM bands compared to ADS. ADS by contrast shows a significantly higher phase dependency between theta- and beta/low gamma rate AM bands compared to IDS. S4 Appendix shows the MI for each music genre and instrument, revealing high consistency between Western music genres and instruments in both S-AMPH and PAD models. Therefore, to probe potentially ubiquitous mutual dependencies across Western genres and instruments, we focused on the grand average (Figure 5). The results showed that adjacent tiers of the AM hierarchy were mutually dependent on each other compared with nonadjacent tiers. Further, mutual dependence between delta- and theta-rate AM bands was the strongest of all mutual dependence in both models. This matched the results of the prior speech-based modelling with IDS and CDS rather than ADS (Leong et al., 2017; Leong et al., 2014; Leong & Goswami, 2015).

As an additional test, we also examined the MI between AM tiers of natural quasi-rhythmic non-musical inputs experienced by humans (utilizing sounds such as rain, wind, fire, storms and rivers; see S1 Appendix), using both the S-AMPH and PAD models. The results were notably different to music. Inspection of the figures in S4 Appendix shows that compared with music, the mutual dependence between delta- and theta-rate AM bands of natural sounds was similar to non-adjacent tiers (S4 Appendix). The current modelling suggests that for music, delta-theta phase alignment of AM bands underpins metrical structure, at least for Western musical genres. Accordingly, metrical structure, a feature shared by both music and speech, depends on the same core delta-theta AM phase relations in both domains.

**Figure 5.**
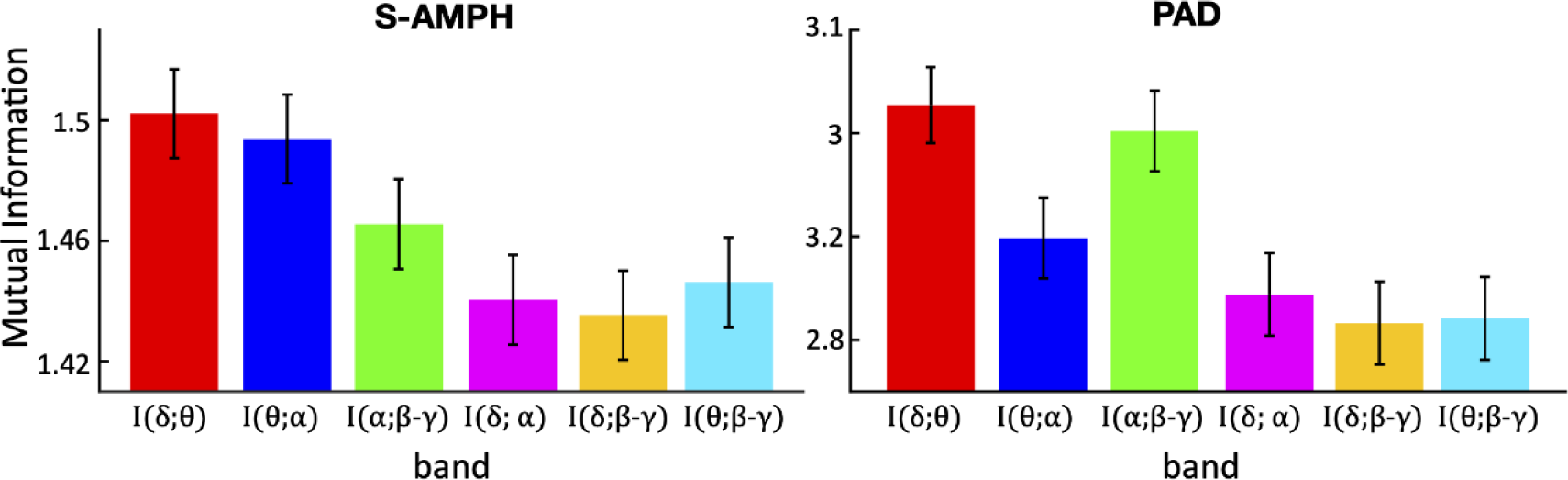
Mutual Information Between Different Tiers in an Amplitude Modulation Hierarchy. *Note*. Both S-AMPH (a) and PAD models (b) showed stronger dependency between each adjacent tier of an amplitude modulations (AM) hierarchy. Further, mutual dependence between delta- and theta-rate AM bands was the strongest of all mutual dependence in both models.

### Multi-Timescale Phase Synchronization in Both Models

The demonstration of mutual dependency does not by itself capture metrical structure, as each AM cycle at a particular timescale may encompass one or more AM cycles at a faster timescale. To identify how many daughter elements a parent element could encompass in general, we next investigated the integer ratios between “*adjacent*” AM bands. For example, if the integer ratio is 1:2, then the parent element encompasses 2 daughter elements for the rhythm. An example from speech would be a tongue twister like “Peter Piper picked a peck of pickled peppers,” which follows a 1:2 ratio (two syllables in each prosodic foot). To assess the integer ratios for each pair of mutually dependent AM bands in our selected musical genres, we used PSI indices. S5 Appendix shows the PSI for each music genre and instrument, revealing high consistency between music genres in both S-AMPH and PAD models. Further analysis focused on the grand average (shown in Figure 6). The PSI of the S-AMPH model suggested that the PSI of 1:2 integer ratios is the highest in all of the adjacent oscillatory bands. The PSIs of 1:3 and 2:3 integer ratios were also higher than the other integer ratios, suggesting that the simpler integer ratios (i.e., m/n) were likely to synchronize in phase with each other. For spoken languages, the m/n ratio between two adjacent AM bands tends to vary with linguistic factors such as how many phonemes typically comprise a syllable (e.g. 2 phonemes per syllable for a language with a consonant-vowel syllable structure like Spanish, hence a theta-beta/low gamma PSI of 1:2, but 3 phonemes per syllable for a language with largely consonant-vowel-consonant syllable structures like English, hence a theta-beta/low gamma PSI of 1:3). For music, the dominance of PSIs 1:3 and 2:3 across genres and instruments suggests more tightly controlled rhythmic dependencies than for speech. The PSIs of the PAD model were similar to the S-AMPH, but PAD was more sensitive to the simple integer ratios. In PAD, the PSIs of not only the 1:2 integer ratios, but also those of the 2:3, 3:4 and 4:5 integer ratios were notably higher than the other integer ratios. This may have arisen because the filterbank used in the S-AMPH model may partially introduce some artificial modulations into the stimuli through “ringing.” However, the ERB_N_ filterbank in the S-AMPH model is the filtering process that reflects the frequency decomposition by cochlear function in the normal human ear. Hence, the different findings between S-AMPH and PAD models regarding phase synchronization may imply that there are differences between auditory signals as perceived by the human brain and the purely physical and statistical structure of music.

**Figure 6.**
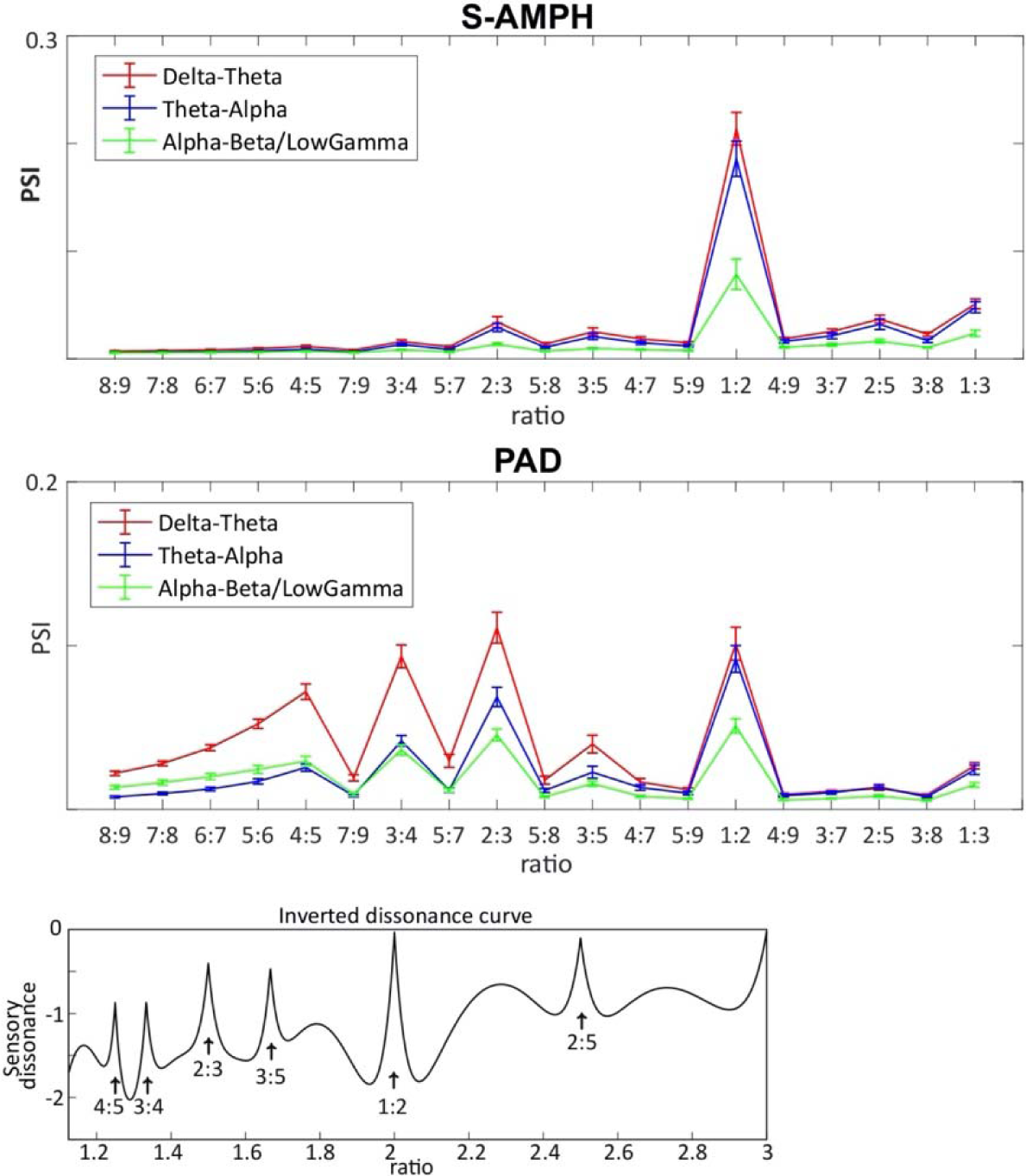
Phase Synchronization Index Between Different Tiers in an Amplitude Modulation Hierarchy. *Note*. Both S-AMPH (a) and PAD models (b) showed that the simpler integer ratios (i.e., m/n) synchronize their phase with each other. The inverted dissonance curve **(c)** was obtained by including the first five upper partials of tones with a 440 Hz (i.e., pitch standard, A4) fundamental frequency in calculating the total dissonance of intervals (Plomp & Levelt, 1965). It is of note that the peaks of PSI (a) correspond to those of the dissonance curve.

Nevertheless, as shown in Figure 6, the PSI between delta- and theta-rate AM bands was consistently the largest PSI in both the S-AMPH and PAD models. Again, this finding is consistent with our prior findings for IDS and rhythmic CDS (Leong et al., 2017; Leong & Goswami, 2015). As a further check, we also examined the PSI of sounds found in nature. The human hearing system has been receiving these quasi-rhythmic sounds at least as long as it has been receiving language and music, but unlike language and music, these sounds have not been produced by humans and shaped by human physiology and culture. Accordingly, it would not be expected that the temporal modulation structure of these natural sounds would be shared with IDS and CDS. The results showed that compared with music, the PSI between delta- and theta-rate AM bands was not consistently the largest PSI (S5 Appendix). This shows that the strong phase dependence between slower bands of AMs revealed for music and for IDS/CDS is not an artifact of the modelling approaches employed, but a core physical feature of their rhythmic structure. This shared structure is demonstrated in Figure 7, which depicts music, IDS, nature sounds (averaged) and a man-made rhythmic sound (a machine) from an envelope demodulation perspective. Comparison of the temporal structures of these sounds for the low-frequency modulation rates (0 – 5 Hz) shows that only music and speech show strong delta- and theta-AM band patterning. The nested structure of AM patterning across the higher modulation bands (12-40Hz) is also clearly visible for each quasi-rhythmic sound for all the natural sounds. This patterning is clearly absent for the man-made rhythmic sound of a machine.

**Figure 7.**
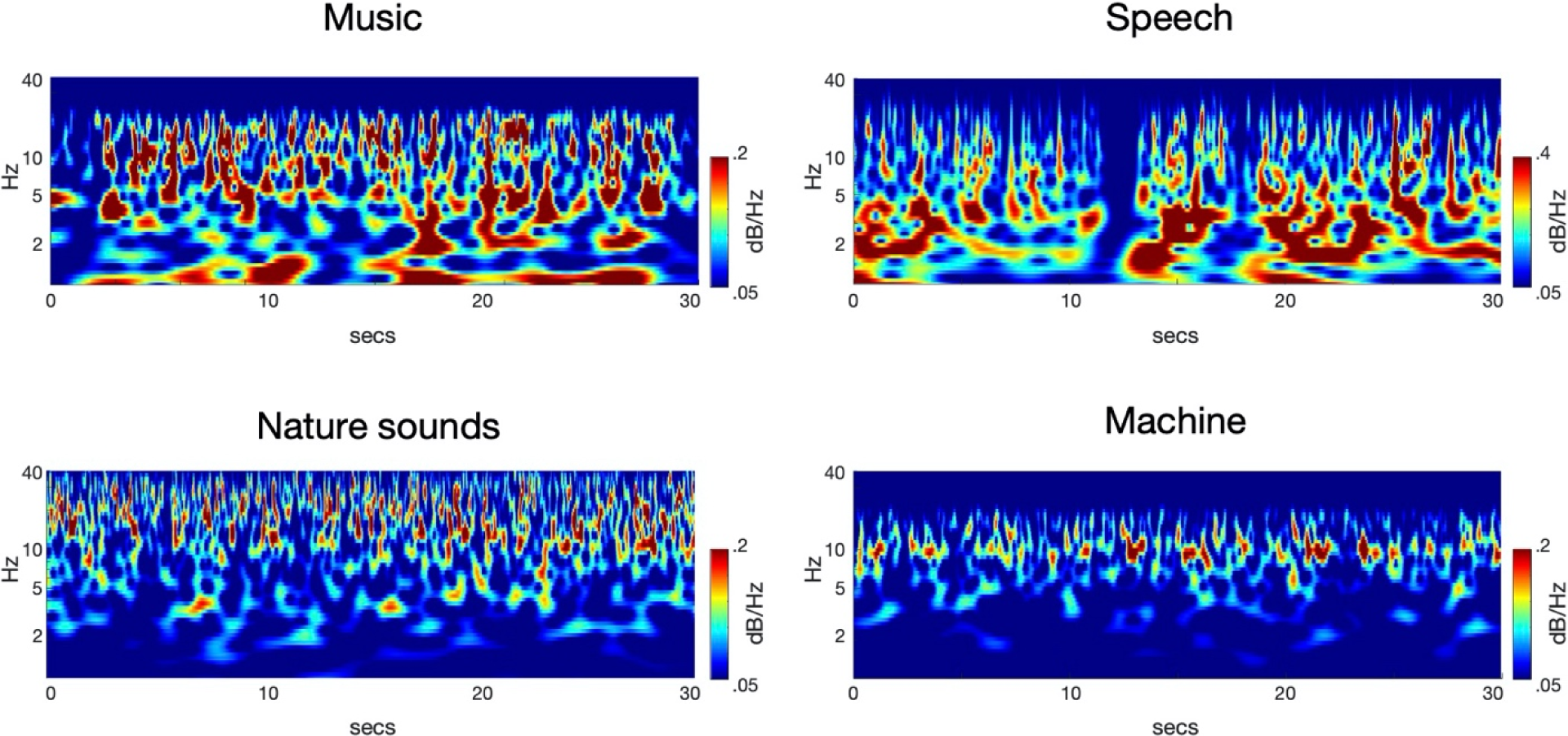
Scalograms Depicting the Amplitude Modulation (AM) Envelopes Derived by Recursive Application of PAD (i.e., Carrier) *Note*. Continuous Wavelet Transform (CWT) was run on each AM envelope from randomly chosen 30-s excerpts of music, speech, nature sounds, and machine sounds. The x-axis denotes time (30 s) and the y-axis denotes modulation rate (0.1-40Hz). The maximal amplitude is normalized to 0 dB. It should be noted the low frequency structure (<5 Hz) visible in music and IDS is absent for the nature and machine sounds.

Accordingly, the strong rhythmic character and acoustic temporal regularity of both infant- and child-directed speech and Western music appears to be influenced by AMs in the delta band (a 2 Hz modulation peak, in music reflecting a 120 bpm rate) and by delta-theta AM phase alignment. Our modelling data for temporal frequency (i.e., “*rhythm*”) also map nicely to the Plomp and Levelt (Plomp & Levelt, 1965) modeling of the dissonance curve for spectral frequency (i.e., “*pitch*”) (shown in Figure 6, Bottom). This may imply that these physical properties of fast spectral frequencies are also involved in very slow temporal modulation envelopes below 40 Hz.

### Simulation Analyses

Finally, to investigate whether the detected (dissonance curve-like) characteristics really represented systematic features of natural musical rhythm, we conducted simulation analyses with synthesized rhythmic but non-musical sounds. The final synthesized waveform comprised a sound that included clear rhythmic information of delta (2Hz), theta (4Hz), alpha (8Hz) and gamma (16Hz) rhythms, and spectral information of a pitch around that of natural human voices (200 Hz) (for the figure, see S5 Appendix). The percept is similar to a harsh rhythmic whisper. As all of the temporal envelopes were comprised of simple sine waves with frequencies of a power of 2, PSI analyses of these artificial sounds should clearly and consistently reveal 1:2 integer ratios compared with other integer ratios. This was the case. Thus, the simulation analyses revealed that the PSIs for natural Western musical genres were different from those for artificial rhythmic sounds. This suggests that natural musical rhythm has covert and systematic integer ratios (i.e., 2:3, 3:4 and 4:5 as well as 1:2) within the AM hierarchy, at least when considering Western musical genres.

## Discussion

Here we tested the prediction that the physical stimulus characteristics (acoustic statistics) that describe IDS and CDS from a demodulation perspective would also describe the hierarchical rhythmic relationships that characterize music. Decomposition of the amplitude envelope of IDS and CDS has previously revealed that (a) the modulation peak in IDS is ∼2 Hz (Leong et al., 2017), (b) that perceived rhythmic patterning depends on the three core AM bands in the amplitude envelope that are found systematically across the spectral range of speech (Leong & Goswami, 2015), and (c) that varying metrical patterns such as trochaic and iambic meters can be identified by the phase relations between two of these bands of AMs (delta- and theta-rate AMs) (Leong et al., 2014). The phase alignment (rhythmic synchronicity) of these relatively slow AM rates represents a unique statistical clue to rhythmic patterning (Goswami, 2019b). Accordingly, we predicted that the physical stimulus characteristics of the amplitude envelope of different musical genres and of music produced by different instruments would yield similar acoustic statistics that described the underlying rhythmic structures.

Our demodulation perspective indeed revealed an hierarchy of temporal modulations that systematically described the acoustic properties of musical rhythm for a range of Western musical genres and instruments. The modelling indicated highly similar acoustic statistical properties to IDS and CDS: a 2Hz modulation peak, particularly strong phase alignment between delta- and theta-rate AM bands across genres, and a distinct set of preferred PSIs that indicated multi-timescale synchronization across different AM bands. As the brain begins learning language using IDS, and consolidates this learning via the rhythmic routines of the nursery (CDS), the present findings are consistent with the theoretical view that perceiving rhythm in both music and language may (at least early in development, prior to acquiring expertise) rely on statistical learning of the same physical stimulus characteristics. Although not tested directly here, it is likely that similar neural oscillatory entrainment mechanisms are used for encoding this hierarchical AM structure in both domains (Doelling & Poeppel, 2015; Norman-Haignere et al., 2015; Nozaradan et al., 2011; Baltzell et al., 2019).

The multi-timescale synchronization found here was systematic across Western musical genres and instruments, suggesting that this AM hierarchy contributes to building perceived rhythmic structures. The nested AM hierarchies in music may yield nested musical units (crotchets, quavers, demiquavers and onsets), just as nested AM hierarchies in CDS yield linguistic units like syllables and rhymes (Leong & Goswami, 2015). The current modelling shows that acoustically-emergent musical units can in principle be parsed reliably from the temporal modulation spectra of the different musical genres examined, and that these units are reflected in each of delta-, theta-, alpha- and beta/gamma-rate bands of AM (Figure 8). To the best of our knowledge, our study is the first to reveal a shared hierarchical AM structure related to musical rhythm.

**Figure 8.**
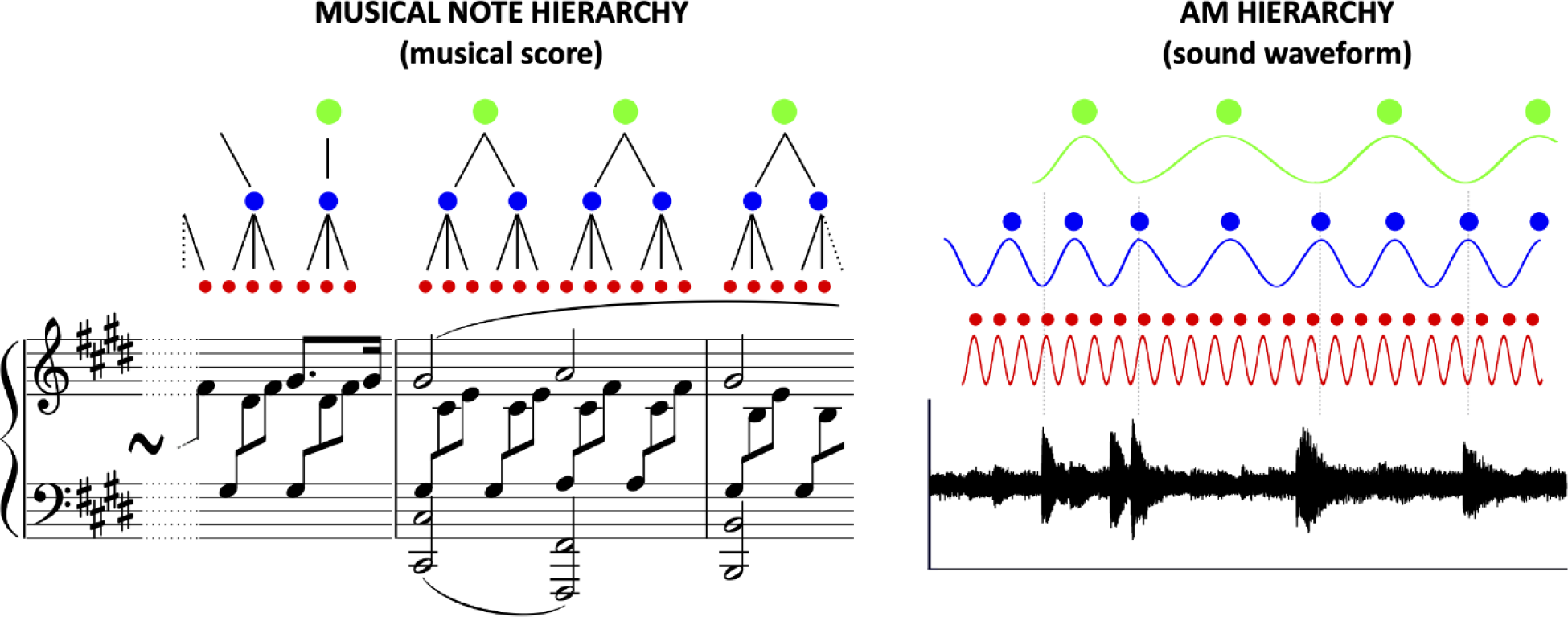
Hierarchical Structure of Rhythm in Music. *Note*. Left and right are the representation by musical score and the corresponding sound waveform of a part of the 33 Variations on a waltz by Anton Diabelli, Op. 120 (commonly known as the Diabelli Variations) by <u>Ludwig van Beethoven</u>. The musical rhythm could be hierarchically organized based either on note values (left) or amplitude modulations (AM, right). As shown by the black lines in the musical note hierarchy (left) and the dotted vertical lines in the AM hierarchy (right), the adjacent tiers of the hierarchy (i.e., green & blue and blue & red) are mutually dependent on each other compared with non-adjacent hierarchical relations (i.e., green-red).

The modelling further revealed strong mutual dependence (using MI estimates) between adjacent bands in the AM hierarchy across musical genres (Western classical, jazz, rock, children’s songs) and musical instruments (piano, guitar, violin, viola, cello, bass, single-voice, multi-voice). In particular, the mutual dependence between delta- and theta-rate bands of AM was the strongest dependence identified by both models. Stronger mutual dependence between delta- and theta-rate AM bands characterizes IDS and CDS, but could not be detected in non-music nature sounds that are quasi-rhythmic (e.g., fire, wind, rain, storms and rivers). By contrast, in ADS there is stronger mutual phase dependence between theta- and beta/low gamma-rate AM bands (Leong et al., 2017); Araujo et al., 2018). The current modelling thus suggests that for music, delta-theta phase alignment of AM bands may underpin metrical rhythmic patterns, at least for Western musical genres.

Convergent results from the phase synchronization analyses further showed that multi-timescale synchronization between delta- and theta-rate AM bands was always higher than the other PSIs regardless of the integer ratios. This was not replicated for nature sounds. The phase alignment of delta- and theta-rate bands of AM has been suggested to be a key acoustic statistic for the language-learning brain (Flanagan & Goswami, 2018; Goswami, 2019b). For example, individual differences in sensitivity to delta- and theta- AM rates and their associated rise times are known to be implicated in disorders of language development (Goswami, 2019a; Goswami, 2019b). In experimental studies, the phase alignment between the slower AM bands (delta and theta) in speech has been demonstrated to play a key role in the perception of metrical patterning (for example, judging whether tone-vocoded nursery rhymes were trochaic or iambic) (Leong et al., 2014). Perceptual data from adults showed that a strong or stressed syllable was perceived when delta and theta modulation peaks were in alignment. The placement of stressed syllables governs metrical patterning in speech (e.g., trochaic, iambic and dactyl meters). The present findings concerning mutual dependence and phase synchronization indicate that music may share these properties: phase alignment between delta- and theta-rate AM bands may contribute to establishing musical metrical structure as well.

Accordingly, prior claims that the rhythmic properties of music and language are distinct (Ding et al., 2017), with the modulation spectrum for music peaking at 2 Hz and the modulation spectrum for speech peaking at 5 Hz, appear to arise from the exclusive reliance of the speech modelling on ADS. By contrast, our modelling approach shows better matching with temporal data from studies of IDS, where the modulation spectrum also peaks at 2 Hz, as well as a similar set of phase relations (the latter were not explored by Ding et al., 2017). We would predict that the statistical regularities in temporal modulations may be the same for other forms of music, and for IDS and CDS in other languages, this remains to be explored. The demonstration that temporal modulation bands play a key role in rhythm hierarchies in music as well as in speech may also suggest that the same evolutionary adaptations underpin both music and language.

Another interesting result from the phase synchronization analyses was the appearance of systematic integer ratios within the AM hierarchy. While the 1:2 integer ratio was strongest for both models, the PSIs for 1:3 and 2:3 were also higher than the other integer ratios explored, for both models. For the PAD modelling approach, which does not make any adjustments for the cochlea, the 2:3, 3:4 and 4:5 integer ratios were also prominent. This statistical patterning may be related to the different metrical structures and integer ratios that characterize music from different cultures (Mehr et al., 2020; McPherson et al., 2020), and even the songs of different species (Roeske et al., 2020). For example, even prior to the acquisition of culture-specific biases of musical rhythm, young infants (5-month-olds) are influenced by ratio complexity (Hannon et al., 2011). Our modelling further suggests that the AM bands in music are related by integer ratios in a similar way to the integer ratios relating notes of different fundamental frequencies that create harmonicity (see the similarity between the PSIs for the two models shown in Figure 7 and the dissonance curve measured by Plomp & Levelt, 1965). Converging prior modelling of speech has shown that the probability distribution of amplitude–frequency combinations in human speech sounds relates statistically to the harmonicity patterns that comprise musical universals (Schwartz et al., 2003). Our modelling appears to suggest that the simple integer ratios (i.e., 1:2, 1:3, and 2:3) in the AM hierarchy comprise a fundamental set of statistics for musical rhythm perception. This fits well with prior data from Jacoby and McDermott (2017), who demonstrated that certain integer ratios are prominent across music from both Western and non-Western cultures. The modelling suggests that AM phase hierarchies may play as strong a role as harmonicity regarding universal mechanisms of human hearing that are important for both music and language.

The modelling presented here is also relevant to the remediation of childhood language disorders. The possible utility of musical interventions for children with disorders of language learning such as developmental language disorder (DLD) and developmental dyslexia has long been recognized (Ladányi et al., 2020; Cumming et al., 2015; Kodály, 1974; Jacques-Dalcroze, 1980; Elliott & Theunissen, 2009). Such interventions are likely to be most beneficial when the temporal hierarchy of the music corresponds to the temporal hierarchy underpinning speech rhythm (Goswami, 2019a; Goswami, 2019b). Careful consideration of the statistical rhythm structures characterizing speech in different languages may thus lead to better remedial outcomes. Similar interventions could be beneficial for second language learners. A caveat is that here we modelled musical genres that could be designated WEIRD corpora (originating from Westernized, educated, industrialized, rich and democratic societies). Accordingly, further studies are necessary to understand how music interventions can contribute to improving speech processing in other languages.

In conclusion, the present study revealed that the acoustic statistics that describe rhythm in Western musical genres from an amplitude envelope decomposition perspective match those that describe IDS and CDS. The physical stimulus characteristics that describe ADS are different, suggesting that studies aiming to discover commonalities between music and speech should not rely exclusively on adult speech corpora. The modelling demonstrates a core acoustic hierarchy of AMs that yield musical rhythm across the amplitude envelopes of different Western musical genres and instruments, with mutual dependencies between AM bands playing a key role in organizing rhythmic units in the musical hierarchy for each genre. Accordingly, biological mechanisms that exploit AM hierarchies may underpin the perception and development of both language and music. In terms of evolution, the novel acoustic statistics revealed here could also explain cross-cultural regularities in musical systems (McPherson et al., 2020); this remains to be tested.

## Supporting information

S1 Appendix. Music Corpora (DOCX).

S2 Appendix. Parameters of the ERBN and modulation filterbanks, and the frequency response characteristics.

S3 Appendix. Individual Variation of PCA Loadings in S-AMPH model and those of FFT in the PAD model.

S4 Appendix. Individual Variation of Mutual Information.

S5 Appendix. Individual Variation of PSI in Each Integer Ratio.

