## Supplementary material for "The Temporal Statistics of Musical Rhythm across Western Genres: An Amplitude Modulation Phase Hierarchy Model": S1 Appendix (from another version of this preprint: version 2, 2021-04-01)

**Music Catalog**

|  | Single instrument | Ensemble |
| --- | --- | --- |
| Classical | ○ | ○ |
| Jazz |  | ○ |
| Rock |  | ○ |
| Children’s song (English) |  | ○ |

***1) Single instrument materials***

| Instrument | Representative  Composers | Duration (minutes) | # of artists |
| --- | --- | --- | --- |
| Piano | Beethoven, Mozart | 230 | 4 |
| Cello | Bach,  Ysaÿe | 139 | 2 |
| Bass | Bach | 60 | 1 |
| Viola | Bach, Hindemith | 210 | 3 |
| Violin | Bach,  Ysaÿe | 204 | 2 |
| Guitar | Bach,  Sanz | 216 | 2 |

***2) Ensemble recordings***

| Genre | Composers or  performers | Duration  (minutes) | # pieces |
| --- | --- | --- | --- |
| Symphonic | Bach, Mozart, Beethoven | 815 | 19 |
| Jazz | Miles Davis, Dave Brubeck | 148 | 20 |
| Rock | The Beatles, U2 | 224 | 50 |
| Children’s song (English) | various |  | 23 |

**Piano**

**[1] Mozart Piano Sonatas K. 310, K. 331, K. 533/494**

Perahia, Murra, pianist.
Sony Classical, SK48233 (1995)

*Mozart WA, Piano Sonata in A minor, K 330 (mistaken) -> K310
Mozart WA, Piano Sonata in A major, K 331
Mozart WA, Piano Sonata in F major, K 533/494*

**[2] Beethoven: Sonata, Op.27,No.2 / Franck: Prelude, choral et fugue / Brahms: Paganini Variations**

Kissin, Evgeny, pianist.
RCA, BMG 09026-68910-2 (1998)

*Beethoven, L. Sonata Op. 27 No. 2 'Moonlight' in C Sharp
Franck, C. Prelude, Choral et Fugue
Brahms, J. Paganini Variations*

### [3] Beethoven: Piano Sonatas, Op.81a & 106 Brendel, Alfred, pianist.

### Phillips Import, Philips 446 093*-2* (1996)

*Beethoven, L. Sonata for piano No. 29 in B flat major ("Hammerklavier") Op. 106*

*Beethoven, L.* [*Sonata for piano No. 26 in E flat major ("Les Adieux") Op. 81a*](http://sg1.allmusic.com/cg/smp.dll?link=p9ieifhgh8bxo475ms6lf4g&r=96.mp3)

##### [4] Mozart: Klaviersonaten

[Pogorelich](http://www.allmusic.com/artist/ivo-pogorelich-mn0002200619), Ivo, pianist.

Dg Imports, DG4377632 (1995)

*Mozart WA, Fantasia for Piano in D minor K 397
Mozart WA, Piano Sonata in G major K 283
Mozart WA, Piano Sonata in A major K 331*

### Cello

#### [5] Unaccompanied cello suites. Johann Sebastian Bach.

Ma, Yo-Yo, cellist.

Columbia, M2K 37867 (1983)

*Bach JS. No. 1, S. BWV 1007, G major
Bach JS. No. 4, S. BWV 1010, E-flat major
Bach JS. No. 5, S. BWV 1011, C minor
Bach JS. No. 2, S. BWV 1008, D minor
Bach JS. No. 3, S. BWV 1009, C major
Bach JS. No. 6, S. BWV 1012, D major*

**[6] Cello Sonatas**
Epperson, [Gordon](http://bobcat.library.nyu.edu/primo_library/libweb/action/search.do?vl%28freeText0%29=Gordon+Epperson&vl%28212921975UI0%29=creator&vl%28378633853UI1%29=all_items&vl%281UIStartWith0%29=exact&fn=search&tab=all&mode=Basic&vid=NYU&scp.scps=scope%3a%28NS%29%2cscope%3a%28CU%29%2cscope%3a%28%22BHS%22%29%2cscope%3a%28NYU%29%2cscope%3a%28%22NYSID%22%29%2cscope%3a%28%22NYHS%22%29%2cscope%3a%28GEN%29%2cscope%3a%28%22NYUAD%22%29), cellist.

Centaur, CRC 2228 (1995)

*Eugène Ysaÿe . Sonate pour violoncelle seul, op. 28
George Crumb . Sonata for solo violoncello
Zoltán Kodály. Sonate pour violoncelle seul, op. 8*

**Bass**

#### [7] [Unaccompanied cello suites: performed on double bass](https://getit.library.nyu.edu/resolve?&ctx_ver=Z39.88-2004&ctx_enc=info:ofi/enc:UTF-8&ctx_tim=2014-03-04T15%3A47%3A28IST&url_ver=Z39.88-2004&url_ctx_fmt=infofi/fmt:kev:mtx:ctx&rfr_id=info:sid/primo.exlibrisgroup.com:primo-nyu_aleph001715054&rft_val_fmt=info:ofi/fmt:kev:mtx:audio&rft.genre=audio&rft.jtitle=&rft.btitle=Unaccompanied%20cello%20suites%20%5bsound%20recording%5d%20:%20performed%20on%20double%20bass&rft.aulast=Bach&rft.aufirst=Johann%20Sebastian%2C&rft.auinit=&rft.auinit1=&rft.auinitm=&rft.ausuffix=&rft.au=Bach%2C%20Johann%20Sebastian%2C%201685-1750&rft.aucorp=&rft.volume=&rft.issue=&rft.part=&rft.quarter=&rft.ssn=&rft.spage=&rft.epage=&rft.pages=&rft.artnum=&rft.pub=Sony%20Classical&rft.place=New%20York%2C%20NY&rft.issn=&rft.eissn=&rft.isbn=&rft.sici=&rft.coden=&rft_id=info:doi/&rft.object_id=&rft.primo=nyu_aleph001715054&rft.eisbn=&rft_dat=%3Cnyu_aleph%3E001715054%3C/nyu_aleph%3E&rft_id=info:oai/)

Meyer, Edgar, bassist.

Sony Classical, SK 89183 (2000)

*Bach, JS. Suites, violoncello, BWV 1008
Bach, JS. Suites, violoncello, BWV 1007
Bach, JS. Suites, violoncello, BWV 1011*

**Viola**

#### [8] Bach, JS. [Six cello suites, on viola](https://getit.library.nyu.edu/resolve?&ctx_ver=Z39.88-2004&ctx_enc=info:ofi/enc:UTF-8&ctx_tim=2014-03-05T13%3A03%3A42IST&url_ver=Z39.88-2004&url_ctx_fmt=infofi/fmt:kev:mtx:ctx&rfr_id=info:sid/primo.exlibrisgroup.com:primo-nyu_aleph003475787&rft_val_fmt=info:ofi/fmt:kev:mtx:audio&rft.genre=audio&rft.jtitle=&rft.btitle=Six%20cello%20suites%2C%20on%20viola%20%5bsound%20recording%5d%20=%20Six%20suites%20pour%20violoncelle%2C%20%C3%A0%20l%27alto&rft.aulast=Bach&rft.aufirst=Johann%20Sebastian%2C&rft.auinit=&rft.auinit1=&rft.auinitm=&rft.ausuffix=&rft.au=Bach%2C%20Johann%20Sebastian%2C%201685-1750&rft.aucorp=&rft.volume=&rft.issue=&rft.part=&rft.quarter=&rft.ssn=&rft.spage=&rft.epage=&rft.pages=&rft.artnum=&rft.pub=Analekta&rft.place=Montr%C3%A9al&rft.issn=&rft.eissn=&rft.isbn=&rft.sici=&rft.coden=&rft_id=info:doi/&rft.object_id=&rft.primo=nyu_aleph003475787&rft.eisbn=&rft_dat=%3Cnyu_aleph%3E003475787%3C/nyu_aleph%3E&rft_id=info:oai/)

Callus, [Helen, violist.](http://bobcat.library.nyu.edu/primo_library/libweb/action/search.do?vl%28freeText0%29=Helen+Callus&vl%28212921975UI0%29=creator&vl%28378633853UI1%29=all_items&vl%281UIStartWith0%29=exact&fn=search&tab=all&mode=Basic&vid=NYU&scp.scps=scope%3a%28NS%29%2cscope%3a%28CU%29%2cscope%3a%28%22BHS%22%29%2cscope%3a%28NYU%29%2cscope%3a%28%22NYSID%22%29%2cscope%3a%28%22NYHS%22%29%2cscope%3a%28GEN%29%2cscope%3a%28%22NYUAD%22%29)

Analekta, AN 2 9968-9 (2011)

*Bach, JS. Suite no. 1 in G major BWV 1007
Bach, JS. Suite no. 2 in D minor, BWV 1008
Bach, JS. Suite no. 3 in C major BWV 1009
Bach, JS. Suite no. 4 in E flat major BWV 1010
Bach, JS. Suite no. 5 in C minor BWV 1011
Bach, JS. Suite no. 6 in D BWV 1012 (transposed in G major)*

**[9] Hindemith, Viola Sonatas**

Levin, [Robert D.](http://bobcat.library.nyu.edu/primo_library/libweb/action/search.do?vl%28freeText0%29=+Robert+D+Levin&vl%28212921975UI0%29=creator&vl%28378633853UI1%29=all_items&vl%281UIStartWith0%29=exact&vl%28freeText0%29=Kim+Kashkashian&vl%28212921975UI0%29=creator&vl%28378633853UI1%29=all_items&vl%281UIStartWith0%29=exact&fn=search&tab=all&mode=Basic&vid=NYU&scp.scps=scope%3a%28NS%29%2cscope%3a%28CU%29%2cscope%3a%28%22BHS%22%29%2cscope%3a%28NYU%29%2cscope%3a%28%22NYSID%22%29%2cscope%3a%28%22NYHS%22%29%2cscope%3a%28GEN%29%2cscope%3a%28%22NYUAD%22%29) [Sonatas for viola alone; Sonatas for viola and piano](https://getit.library.nyu.edu/resolve?&ctx_ver=Z39.88-2004&ctx_enc=info:ofi/enc:UTF-8&ctx_tim=2014-03-05T13%3A08%3A41IST&url_ver=Z39.88-2004&url_ctx_fmt=infofi/fmt:kev:mtx:ctx&rfr_id=info:sid/primo.exlibrisgroup.com:primo-nyu_aleph001720022&rft_val_fmt=info:ofi/fmt:kev:mtx:audio&rft.genre=audio&rft.jtitle=&rft.btitle=Sonatas%20for%20viola%20alone%20%5bsound%20recording%5d%20%3B%20Sonatas%20for%20viola%20and%20piano&rft.aulast=Hindemith&rft.aufirst=Paul%2C&rft.auinit=&rft.auinit1=&rft.auinitm=&rft.ausuffix=&rft.au=Hindemith%2C%20Paul%2C%201895-1963&rft.aucorp=&rft.volume=&rft.issue=&rft.part=&rft.quarter=&rft.ssn=&rft.spage=&rft.epage=&rft.pages=&rft.artnum=&rft.pub=ECM&rft.place=M%C3%BCnchen&rft.issn=&rft.eissn=&rft.isbn=&rft.sici=&rft.coden=&rft_id=info:doi/&rft.object_id=&rft.primo=nyu_aleph001720022&rft.eisbn=&rft_dat=%3Cnyu_aleph%3E001720022%3C/nyu_aleph%3E&rft_id=info:oai/)
Kashkashian, [Kim, violist.](http://bobcat.library.nyu.edu/primo_library/libweb/action/search.do?vl%28freeText0%29=Kim+Kashkashian&vl%28212921975UI0%29=creator&vl%28378633853UI1%29=all_items&vl%281UIStartWith0%29=exact&fn=search&tab=all&mode=Basic&vid=NYU&scp.scps=scope%3a%28NS%29%2cscope%3a%28CU%29%2cscope%3a%28%22BHS%22%29%2cscope%3a%28NYU%29%2cscope%3a%28%22NYSID%22%29%2cscope%3a%28%22NYHS%22%29%2cscope%3a%28GEN%29%2cscope%3a%28%22NYUAD%22%29)

ECM New Series, ECM 1330-32 (1988)

#### [*Paul Hindemith*](http://bobcat.library.nyu.edu/primo_library/libweb/action/search.do?vl%28freeText0%29=Paul++Hindemith++1895-1963.&vl%28212921975UI0%29=creator&vl%28378633853UI1%29=all_items&vl%281UIStartWith0%29=exact&fn=search&tab=all&mode=Basic&vid=NYU&scp.scps=scope%3a%28NS%29%2cscope%3a%28CU%29%2cscope%3a%28%22BHS%22%29%2cscope%3a%28NYU%29%2cscope%3a%28%22NYSID%22%29%2cscope%3a%28%22NYHS%22%29%2cscope%3a%28GEN%29%2cscope%3a%28%22NYUAD%22%29)*. Sonatas for viola alone. op. 31/4* [*Paul Hindemith*](http://bobcat.library.nyu.edu/primo_library/libweb/action/search.do?vl%28freeText0%29=Paul++Hindemith++1895-1963.&vl%28212921975UI0%29=creator&vl%28378633853UI1%29=all_items&vl%281UIStartWith0%29=exact&fn=search&tab=all&mode=Basic&vid=NYU&scp.scps=scope%3a%28NS%29%2cscope%3a%28CU%29%2cscope%3a%28%22BHS%22%29%2cscope%3a%28NYU%29%2cscope%3a%28%22NYSID%22%29%2cscope%3a%28%22NYHS%22%29%2cscope%3a%28GEN%29%2cscope%3a%28%22NYUAD%22%29)*. Sonatas for viola alone op. 25/1* [*Paul Hindemith*](http://bobcat.library.nyu.edu/primo_library/libweb/action/search.do?vl%28freeText0%29=Paul++Hindemith++1895-1963.&vl%28212921975UI0%29=creator&vl%28378633853UI1%29=all_items&vl%281UIStartWith0%29=exact&fn=search&tab=all&mode=Basic&vid=NYU&scp.scps=scope%3a%28NS%29%2cscope%3a%28CU%29%2cscope%3a%28%22BHS%22%29%2cscope%3a%28NYU%29%2cscope%3a%28%22NYSID%22%29%2cscope%3a%28%22NYHS%22%29%2cscope%3a%28GEN%29%2cscope%3a%28%22NYUAD%22%29)*. Sonatas for viola alone op. 11/5*

**[10] Watras,** [**Melia**](http://bobcat.library.nyu.edu/primo_library/libweb/action/search.do?vl%28freeText0%29=Melia++Watras&vl%28212921975UI0%29=creator&vl%28378633853UI1%29=all_items&vl%281UIStartWith0%29=exact&fn=search&tab=all&mode=Basic&vid=NYU&scp.scps=scope%3a%28NS%29%2cscope%3a%28CU%29%2cscope%3a%28%22BHS%22%29%2cscope%3a%28NYU%29%2cscope%3a%28%22NYSID%22%29%2cscope%3a%28%22NYHS%22%29%2cscope%3a%28GEN%29%2cscope%3a%28%22NYUAD%22%29) **. Viola solo**
Fleur de Son Classics, FDS 57962 (2004)

*Arad, Atar. Sonata
Bach, Johann Sebastian. Chromatische Fantasie und Fuge. Fantasia.
Corigliano, John. Fancy on a Bach air
Waggoner, Andrew. Collines parmi étoiles
Stravinsky, Igor. Elegy.
Prestini, Paola. Sympathique.
Penderecki, Krzysztof. Cadenza*

**Violin**

**[11]** [**Ysaÿe**](http://bobcat.library.nyu.edu/primo_library/libweb/action/search.do?vl%28freeText0%29=Eug%c3%a8ne++Ysa%c3%bfe++1858-1931.&vl%28212921975UI0%29=creator&vl%28378633853UI1%29=all_items&vl%281UIStartWith0%29=exact&fn=search&tab=all&mode=Basic&vid=NYU&scp.scps=scope%3a%28NS%29%2cscope%3a%28CU%29%2cscope%3a%28%22BHS%22%29%2cscope%3a%28NYU%29%2cscope%3a%28%22NYSID%22%29%2cscope%3a%28%22NYHS%22%29%2cscope%3a%28GEN%29%2cscope%3a%28%22NYUAD%22%29) **Eugène. Six sonatas for solo violin, op. 27**

#### Murray, [Tai, violinst.](http://bobcat.library.nyu.edu/primo_library/libweb/action/search.do?vl%28freeText0%29=Tai+Murray&vl%28212921975UI0%29=creator&vl%28378633853UI1%29=all_items&vl%281UIStartWith0%29=exact&fn=search&tab=all&mode=Basic&vid=NYU&scp.scps=scope%3a%28NS%29%2cscope%3a%28CU%29%2cscope%3a%28%22BHS%22%29%2cscope%3a%28NYU%29%2cscope%3a%28%22NYSID%22%29%2cscope%3a%28%22NYHS%22%29%2cscope%3a%28GEN%29%2cscope%3a%28%22NYUAD%22%29)

Harmonia Mundi USA, HMU 907569 (2012)

[*Ysaÿe*](http://bobcat.library.nyu.edu/primo_library/libweb/action/search.do?vl%28freeText0%29=Eug%c3%a8ne++Ysa%c3%bfe++1858-1931.&vl%28212921975UI0%29=creator&vl%28378633853UI1%29=all_items&vl%281UIStartWith0%29=exact&fn=search&tab=all&mode=Basic&vid=NYU&scp.scps=scope%3a%28NS%29%2cscope%3a%28CU%29%2cscope%3a%28%22BHS%22%29%2cscope%3a%28NYU%29%2cscope%3a%28%22NYSID%22%29%2cscope%3a%28%22NYHS%22%29%2cscope%3a%28GEN%29%2cscope%3a%28%22NYUAD%22%29) *Eugène. Sonata no. 1*
[*Ysaÿe*](http://bobcat.library.nyu.edu/primo_library/libweb/action/search.do?vl%28freeText0%29=Eug%c3%a8ne++Ysa%c3%bfe++1858-1931.&vl%28212921975UI0%29=creator&vl%28378633853UI1%29=all_items&vl%281UIStartWith0%29=exact&fn=search&tab=all&mode=Basic&vid=NYU&scp.scps=scope%3a%28NS%29%2cscope%3a%28CU%29%2cscope%3a%28%22BHS%22%29%2cscope%3a%28NYU%29%2cscope%3a%28%22NYSID%22%29%2cscope%3a%28%22NYHS%22%29%2cscope%3a%28GEN%29%2cscope%3a%28%22NYUAD%22%29) *Eugène. Sonata no. 2*
[*Ysaÿe*](http://bobcat.library.nyu.edu/primo_library/libweb/action/search.do?vl%28freeText0%29=Eug%c3%a8ne++Ysa%c3%bfe++1858-1931.&vl%28212921975UI0%29=creator&vl%28378633853UI1%29=all_items&vl%281UIStartWith0%29=exact&fn=search&tab=all&mode=Basic&vid=NYU&scp.scps=scope%3a%28NS%29%2cscope%3a%28CU%29%2cscope%3a%28%22BHS%22%29%2cscope%3a%28NYU%29%2cscope%3a%28%22NYSID%22%29%2cscope%3a%28%22NYHS%22%29%2cscope%3a%28GEN%29%2cscope%3a%28%22NYUAD%22%29) *Eugène. Sonata no. 3*
[*Ysaÿe*](http://bobcat.library.nyu.edu/primo_library/libweb/action/search.do?vl%28freeText0%29=Eug%c3%a8ne++Ysa%c3%bfe++1858-1931.&vl%28212921975UI0%29=creator&vl%28378633853UI1%29=all_items&vl%281UIStartWith0%29=exact&fn=search&tab=all&mode=Basic&vid=NYU&scp.scps=scope%3a%28NS%29%2cscope%3a%28CU%29%2cscope%3a%28%22BHS%22%29%2cscope%3a%28NYU%29%2cscope%3a%28%22NYSID%22%29%2cscope%3a%28%22NYHS%22%29%2cscope%3a%28GEN%29%2cscope%3a%28%22NYUAD%22%29) *Eugène. Sonata no. 4*
[*Ysaÿe*](http://bobcat.library.nyu.edu/primo_library/libweb/action/search.do?vl%28freeText0%29=Eug%c3%a8ne++Ysa%c3%bfe++1858-1931.&vl%28212921975UI0%29=creator&vl%28378633853UI1%29=all_items&vl%281UIStartWith0%29=exact&fn=search&tab=all&mode=Basic&vid=NYU&scp.scps=scope%3a%28NS%29%2cscope%3a%28CU%29%2cscope%3a%28%22BHS%22%29%2cscope%3a%28NYU%29%2cscope%3a%28%22NYSID%22%29%2cscope%3a%28%22NYHS%22%29%2cscope%3a%28GEN%29%2cscope%3a%28%22NYUAD%22%29) *Eugène. Sonata no. 5*
[*Ysaÿe*](http://bobcat.library.nyu.edu/primo_library/libweb/action/search.do?vl%28freeText0%29=Eug%c3%a8ne++Ysa%c3%bfe++1858-1931.&vl%28212921975UI0%29=creator&vl%28378633853UI1%29=all_items&vl%281UIStartWith0%29=exact&fn=search&tab=all&mode=Basic&vid=NYU&scp.scps=scope%3a%28NS%29%2cscope%3a%28CU%29%2cscope%3a%28%22BHS%22%29%2cscope%3a%28NYU%29%2cscope%3a%28%22NYSID%22%29%2cscope%3a%28%22NYHS%22%29%2cscope%3a%28GEN%29%2cscope%3a%28%22NYUAD%22%29) *Eugène. Sonata no. 6*

#### [12] Bach, JS. [The complete sonatas and partitas for solo violin. Vol. 1](https://getit.library.nyu.edu/resolve?&ctx_ver=Z39.88-2004&ctx_enc=info:ofi/enc:UTF-8&ctx_tim=2014-03-05T13%3A24%3A43IST&url_ver=Z39.88-2004&url_ctx_fmt=infofi/fmt:kev:mtx:ctx&rfr_id=info:sid/primo.exlibrisgroup.com:primo-nyu_aleph000501198&rft_val_fmt=info:ofi/fmt:kev:mtx:audio&rft.genre=audio&rft.jtitle=&rft.btitle=The%20complete%20sonatas%20and%20partitas%20for%20solo%20violin.%20Vol.%201%20%5bsound%20recording%5d&rft.aulast=Bach&rft.aufirst=Johann%20Sebastian%2C&rft.auinit=&rft.auinit1=&rft.auinitm=&rft.ausuffix=&rft.au=Bach%2C%20Johann%20Sebastian%2C%201685-1750&rft.aucorp=&rft.volume=&rft.issue=&rft.part=&rft.quarter=&rft.ssn=&rft.spage=&rft.epage=&rft.pages=&rft.artnum=&rft.pub=Gaudeamus&rft.place=England%5d&rft.issn=&rft.eissn=&rft.isbn=&rft.sici=&rft.coden=&rft_id=info:doi/&rft.object_id=&rft.primo=nyu_aleph000501198&rft.eisbn=&rft_dat=%3Cnyu_aleph%3E000501198%3C/nyu_aleph%3E&rft_id=info:oai/)

Ross, Jacqueline, violinst.

Gaudeamus, GAU 358 (2007)

*Bach JS. Sonata no. 1 in g minor, BWV 1001
Bach JS. Partita no. 1 in B minor, BWV 1002
Bach JS. Sonata no. 2 in A minor, BWV 1003*

#### Bach, JS. [The complete sonatas and partitas for solo violin. Vol. 2](https://getit.library.nyu.edu/resolve?&ctx_ver=Z39.88-2004&ctx_enc=info:ofi/enc:UTF-8&ctx_tim=2014-03-05T13%3A24%3A43IST&url_ver=Z39.88-2004&url_ctx_fmt=infofi/fmt:kev:mtx:ctx&rfr_id=info:sid/primo.exlibrisgroup.com:primo-nyu_aleph000501198&rft_val_fmt=info:ofi/fmt:kev:mtx:audio&rft.genre=audio&rft.jtitle=&rft.btitle=The%20complete%20sonatas%20and%20partitas%20for%20solo%20violin.%20Vol.%201%20%5bsound%20recording%5d&rft.aulast=Bach&rft.aufirst=Johann%20Sebastian%2C&rft.auinit=&rft.auinit1=&rft.auinitm=&rft.ausuffix=&rft.au=Bach%2C%20Johann%20Sebastian%2C%201685-1750&rft.aucorp=&rft.volume=&rft.issue=&rft.part=&rft.quarter=&rft.ssn=&rft.spage=&rft.epage=&rft.pages=&rft.artnum=&rft.pub=Gaudeamus&rft.place=England%5d&rft.issn=&rft.eissn=&rft.isbn=&rft.sici=&rft.coden=&rft_id=info:doi/&rft.object_id=&rft.primo=nyu_aleph000501198&rft.eisbn=&rft_dat=%3Cnyu_aleph%3E000501198%3C/nyu_aleph%3E&rft_id=info:oai/).

Ross, Jacqueline.

Gaudeamus, GAU 359 (2007)

*Bach JS. Partita no. 2 in D minor, BWV 1004
Bach JS. Sonata no. 3 in C, BWV 1005
Bach JS. Partita no. 3 in E, BWV 1006.*

**Guitar**

**[13] Segovia,** [**Andrés**](http://bobcat.library.nyu.edu/primo_library/libweb/action/search.do?vl%28freeText0%29=Andr%c3%a9s++Segovia++1893-1987.&vl%28212921975UI0%29=creator&vl%28378633853UI1%29=all_items&vl%281UIStartWith0%29=exact&fn=search&tab=all&mode=Basic&vid=NYU&scp.scps=scope%3a%28NS%29%2cscope%3a%28CU%29%2cscope%3a%28%22BHS%22%29%2cscope%3a%28NYU%29%2cscope%3a%28%22NYSID%22%29%2cscope%3a%28%22NYHS%22%29%2cscope%3a%28GEN%29%2cscope%3a%28%22NYUAD%22%29)**. Art of Segovia**

Deutsche Grammophon, 289 471 697-2 (2002)
*Handel, George Frideric. Suite for harpsichord no. 4 in D minor, HWV 437.
 Sarabande
Bach JS. Suite for violoncello solo no. 1 in G major, BWV 1007. Prélude
Bach JS. Partita for violin solo no. 1 in B minor, BWV 1002. Tempo de bourrée
Bach JS. Suite for violoncello solo no. 3 in C major, BWV 1009. Courante
Bach JS. Partita for violin solo no. 3 E major, BWV 1006.Gavotte en rondeau Aria e corrente
Frescobaldi, Girolamo. Sonata in C minor, K. 11 (L. 352)
Scarlatti, Domenico. Nouvelles suites de pièces de clavecin.
Rameau, Jean-Philippe. Menuet in G major.
Manuel Ponce after Nicolò Paganini . Andantino variato
Chopin, Frédéric. 24 préludes, op. 28. No. 7 in A major
Mendelssohn, Felix. String quartet in E flat major, op. 12. Canzonetta
Franck, César. L'Organiste, FWV 41, Sept pièces en mi bémol majeur et mi bémol mineur. Quasi lento, Andantino poco allegretto
Mussorgsky, Modest. Pictures at an exhibition. Il vecchio castello
Grieg, Edvard. Lyric pieces IV, op. 47. Melodie
Debussy, Claude. Préludes, livre 1. La fille aux cheveux de lin
Scriabin, Alexander. 5 preludes, op. 16. No. 4 in E flat minor
Albéniz, Isaac. Suite española. Asturias. Leyenda-Preludio
Albéniz, Isaac. Piezas características. Zambra granadina
Segovia, Andrés. Estudio sin luz
Rodrigo, Joaquín. Fantasía para un gentilhombre for guitar and small orchestra.
 Danza de las hachas
Jordà, Enrique. Symphony of the Air*

**[14] Bream, Julian. Baroque Guitar**
RCA Victor Gold Seal, RCA 60494 (1991)

*Sanz, Gaspar. Pavanas
Sanz, Gaspar. Galliardas
Sanz, Gaspar. Passacalles
Sanz, Gaspar. Canarios
Guerau, Francisco. Villano
Guerau, Francisco. Canario
Bach, JS. Prelude in D minor, BWV 999
Bach, JS. Fugue in A minor, BWV 1000
Weiss, Sylvius Leopold. Passacaille
Weiss, Sylvius Leopold. Fantasie
Weiss, Sylvius Leopold. Tombeau sur la mort de M. Comte de Logy
de Visée, Robert. Suite in D minor
Frescobaldi, Girolamo. Aria con variazione detta la Frescobalda arr. Segovia
Scarlatti, Domenico. Sonata in E minor, K. 11
Scarlatti, Domenico. Sonata in E minor, K. 87, arr. Bream [K. 11], Segovia [K. 87]
Cimarosa, Domenico. Sonata in C# minor
Cimarosa, Domenico. Sonata in A, arr. Bream.*

**Symphonies**

### [15] The Beethoven Symphonies 1-9 Live From the Edinburgh Festival Mackeras, Sir Charles, conductor.

### Hyperion UK, CDS44301/5 (2007)

*Beethoven L. Symphony #1
Beethoven L. Symphony #2
Beethoven L. Symphony #3
Beethoven L. Symphony #4
Beethoven L. Symphony #5
Beethoven L. Symphony #6
Beethoven L. Symphony #7
Beethoven L. Symphony #8
Beethoven L. Symphony #9*

**[16] Jacobs,** [**René, conductor.**](http://www.allmusic.com/artist/renÃ©-jacobs-mn0001981440) [**Freiburger**](http://www.allmusic.com/artist/freiburger-barockorchester-mn0001519316) **Barockorchester.**

Mozart: Symphonies Nos. 39 & 40

Harmonia Mundi, HMC901959 (2010)

*Symphony 39 in E-flat major K 543
Symphony 40 in G minor K 550*

**[17] Brüggen,** [**Frans, conductor.**](http://www.allmusic.com/artist/frans-brÃ¼ggen-mn0000184236) [**Orchestra of the Eighteenth Century**](http://www.allmusic.com/artist/orchestra-of-the-eighteenth-century-mn0002207604)**.**

Mozart: Symphony No. 41; La Clemenza di Tito Overture

Philips Digital Classics, Phillips 420 241-1 (2010)

*Mozart WA, Symphony 41
Mozart WA, La Clemenza di Tito Overture*

**[18] Bach: Brandenburg Concertos**

Alessandrini, Rinaldo, conductor. Concerto Italiano.

Naïve, OP 30412 (2005)

*Bach JS. Brandenburg Concerto No. 1 in F major, BWV 1046*

*Bach JS. Brandenburg Concerto No. 2 in F major, BWV 1047*

*Bach JS. Brandenburg Concerto No. 3 in G major, BWV 1048*

*Bach JS. Brandenburg Concerto No. 4 in G major, BWV 1049*

*Bach JS. Brandenburg Concerto No. 5 in D major, BWV 1050*

*Bach JS. Brandenburg Concerto No. 6 in B flat major, BWV 1051*

**Jazz**

**[19] Davis, Miles. Kind of Blue (or. 1959)**
Columbia/Legacy, CN 90887 (2004)

*(1) So what
(2) Freddie Freeloader
(3) Blue in green
(4) All blues
(5) Flamenco sketches
(6) Flamenco sketches (alternate take)*

### [20] [Dave Brubeck Quartet.](http://bobcat.library.nyu.edu/primo_library/libweb/action/search.do?vl%28freeText0%29=Dave+Brubeck+Quartet.&vl%28212921975UI0%29=creator&vl%28378633853UI1%29=all_items&vl%281UIStartWith0%29=exact&fn=search&tab=all&mode=Basic&vid=NYU&scp.scps=scope%3a%28NS%29%2cscope%3a%28CU%29%2cscope%3a%28%22BHS%22%29%2cscope%3a%28NYU%29%2cscope%3a%28%22NYSID%22%29%2cscope%3a%28%22NYHS%22%29%2cscope%3a%28GEN%29%2cscope%3a%28%22NYUAD%22%29) Time Out (or. 1959) Columbia/Legacy, CN 88697 (2009)

### *(7) Blue rondo à la Turk (8) Strange meadow lark (9) Three to get ready (10) Kathy's waltz (11) Everybody's jumpin' (12) Pick up sticks (13) The Dave Brubeck Quartet live at Newport. St. Louis Blues (14) Waltz limp (15) Since love had its way (16) Koto song (17) Pennies from heaven (18) You go to my head (19) Blue rondo à la Turk (20) Take five*

**Rock**

**[21] The Beatles. The Beatles 1967-1970**

Apple Records, CDP 0777 7 97039 2 0 (1993)

*(1) Strawberry Fields Forever*

*(2) Penny Lane*

*(3) Sgt. Pepper's Lonely Hearts Club Band*

*(4) Here Comes the Sun*

*(5) Come Together*

*(6) Something*

*(7) Octopus's Garden*

*(8) Let It Be*

*(9) Across the Universe*

*(10) The Long and Winding Road*

**[22] The Beatles. The Beatles 1962-1966**

Apple Records, CDP 0777 7 97036 2 3 (1993)

(11) *Love Me Do*

*(12) Please Please Me*

*(13) From Me To You*

*(14) She Loves You*

*(15) I Want To Hold Your Hand*

*(16) All My Loving*

*(17) Can't Buy Me Love*

*(18) A Hard Day's Night*

*(19) And I Love Her*

*(20) Eight Days A Week*

*(21) I Feel Fine*

*(22) Ticket To Ride*

*(23) Yesterday*

*(24) Help!*

*(25) You've Got To Hide Your Love Away*

*(26) We Can Work It Out*

*(27) Day Tripper*

*(28) Drive My Car*

*(29) Norwegian Wood (This Bird Has Flown)*

*(30) Nowhere Man*

*(31) Michelle*

*(32) In My Life*

*(33) Girl*

*(34) Paperback Writer*

*(35) Eleanor Rigby*

*(36) Yellow Submarine*

**[23] U2. The Best of 1980-1990**

Island, ADV7963-2 (1998)

*(37) Pride (In The Name Of Love)*

*(38) New Year's Day*

*(39) With Or Without You*

*(40) I Still Haven't Found What I'm Looking For*

*(41) Sunday Bloody Sunday*

*(42) Bad*

*(43) Where The Streets Have No Name*

*(44) I Will Follow*

*(45) Unforgettable Fire*

*(46) Sweetest Thing*

*(47) Desire*

*(48) When Love Comes To Town*

*(49) Angel Of Harlem*

*(50) All I Want Is You*

**Children’s Song (English)**

**[24]** Twinkle Twinkle Little Star & Other Lullabies - Baby Lullaby Music and Childrens Songs for Bedtime Sleep

1. Twinkle Twinkle Little Star by Nursery Rhymes ABC

2. Lavender's Blue (Dilly Dilly) by Nursery Rhymes ABC

3. I Can Sing A Rainbow by Nursery Rhymes ABC

4. Hush a Bye Baby Lullaby by Nursery Rhymes ABC

5. Beyond the Sea (Finding Nemo) (Piano Lullaby Version) by Sleepyheadz

6. You Are My Sunshine by Nursery Rhymes ABC

7. A Whole New World (Aladdin) (Piano Lullaby Version) by Sleepyheadz

8. Time To Go To Sleep by Nursery Rhymes ABC

9. Stretch & Grow Lullaby by Nursery Rhymes ABC

10. Can You Feel the Love Tonight (The Lion King) (Piano Lullaby Version) by Sleepyheadz

11. I See The Moon by Nursery Rhymes ABC

12. Circle of Life (The Lion King) (Piano Lullaby Version) by Sleepyheadz

13. When You Wish Upon A Star (Pinocchio) (Piano Lullaby Version) by Sleepyheadz

14. Once Upon A Dream (Sleeping Beauty) (Piano Lullaby Version) by Sleepyheadz

15. Let It Go (Frozen) (Piano Lullaby Version) by Sleepyheadz

16. Someday My Prince Will Come (Snow White) (Piano Lullaby Version) by Sleepyheadz

17. How Far I'll Go (Moana) (Piano Lullaby Version) by Sleepyheadz

18. Something There (Beauty and the Beast) (Piano Lullaby Version)　by Sleepyheadz

19. Under the Sea (The Little Mermaid) (Piano Lullaby Version) by Sleepyheadz

20. Brahms Lullaby by Nursery Rhymes ABC

**[25]** Children's Birthday Party Music (Nursery Rhymes)

1. Humpty Dumpty by Nursery Rhymes

2. Little Bo Peep by Nursery Rhymes

3. The Farmer in the Dell by Nursery Rhymes

4. Jack & Jill by Nursery Rhymes

5. There Was a Crooked Man by Nursery Rhymes

6. Goosey Goosey Gander by Songs For Children

7. The Muffin Man by Songs For Children

8. Three Little Kittens by Children's Lullabyes

9. My Special Favorites : Old King Cole / If I Had a Donkey / Hickory Dickory Dock / Little Nut Tree / The Owl and the Pussycat by Nursery Rhymes

10. Lady Bird, Lady Bird by Children's Lullabyes

11. A-Tisket, A-Tasket by Nursery Rhymes

12. Old Mother Hubbard by Nursery Rhymes

13. Just Plain Silly : Jack and Jill / Three Blind Mice / Rub-A-Dub / Humpty Dumpty / What Are Little Boys Made of / I Saw a Ship a Sailing / Moses' Toeses / Peter Piper / The Cat and the Fiddle by Songs For Children

14. The Man in the Moon by Songs For Children

15. Playtime : Here We Go Round the Mulberry Bush / Pat a Cake / One, Two, Buckle My Shoe / Itsy Bitsy Spider / This Little Piggy by Children's Lullabyes

16. Skip to My Loo by Songs For Children

Nature sounds

<https://mixkit.co/free-sound-effects/nature/>

<https://www.zapsplat.com>

| Types | Duration (seconds) | # pieces |
| --- | --- | --- |
| Fire | 168 | 3 |
| Storm | 112 | 3 |
| Wind | 178 | 4 |
| Rain | 113 | 4 |
| River | 154 | 3 |
