## Supplementary material for "The Temporal Statistics of Musical Rhythm across Western Genres: An Amplitude Modulation Phase Hierarchy Model": S2 Appendix (from another version of this preprint: version 2, 2021-04-01)

**ERB filterbank (for 28 filter bank) and the frequency response characteristics (Fig. a).**

| Edge Number | Edge (Hz) |
| --- | --- |
| 1 (low-pass) | 100 |
| 2 | 137 |
| 3 | 179 |
| 4 | 225 |
| 5 | 277 |
| 6 | 334 |
| 7 | 398 |
| 8 | 470 |
| 9 | 549 |
| 10 | 638 |
| 11 | 736 |
| 12 | 846 |
| 13 | 969 |
| 14 | 1105 |
| 15 | 1257 |
| 16 | 1426 |
| 17 | 1614 |
| 18 | 1824 |
| 19 | 2057 |
| 20 | 2317 |
| 21 | 2607 |
| 22 | 2930 |
| 23 | 3289 |
| 24 | 3689 |
| 25 | 4135 |
| 26 | 4631 |
| 27 | 5184 |
| 28 | 5800 |
| 29 | 6486 |
| 30 | 7250 |

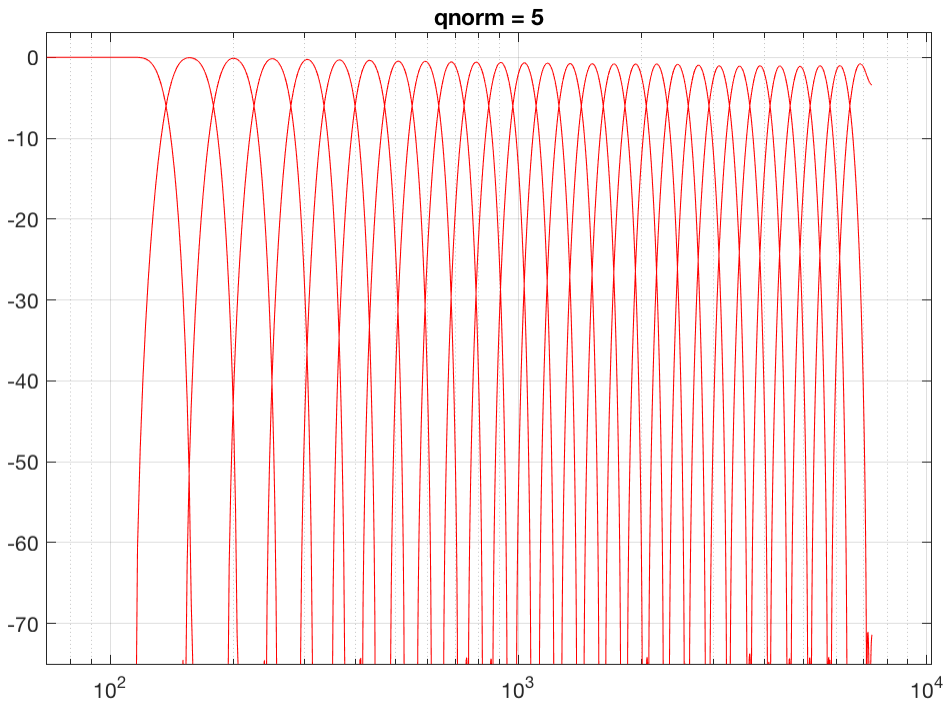

**Fig. a.** frequency response characteristics (28 filterbanks).

**Modulation filterbank (for 24 filter bank) and the frequency response characteristics (Fig. b and c).**

| Edge Number | Edge (Hz) |
| --- | --- |
| 1 (low-pass dummy channel,output discarded) | 0.79 |
| 2 | 0.93 |
| 3 | 1.09 |
| 4 | 1.27 |
| 5 | 1.49 |
| 6 | 1.74 |
| 7 | 2.03 |
| 8 | 2.38 |
| 9 | 2.78 |
| 10 | 3.25 |
| 11 | 3.80 |
| 12 | 4.45 |
| 13 | 5.20 |
| 14 | 6.08 |
| 15 | 7.11 |
| 16 | 8.32 |
| 17 | 9.72 |
| 18 | 11.38 |
| 19 | 13.30 |
| 20 | 15.56 |
| 21 | 18.20 |
| 22 | 21.28 |
| 23 | 24.89 |
| 24 | 29.11 |
| 25 | 34.04 |
| 26 | 39.81 |

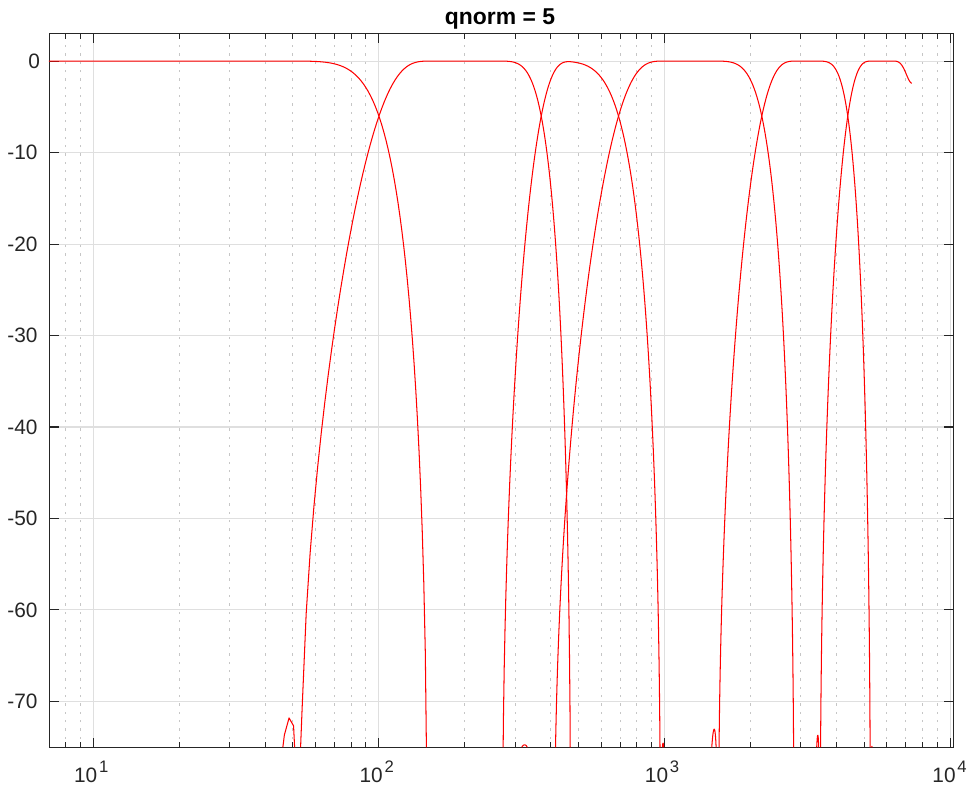

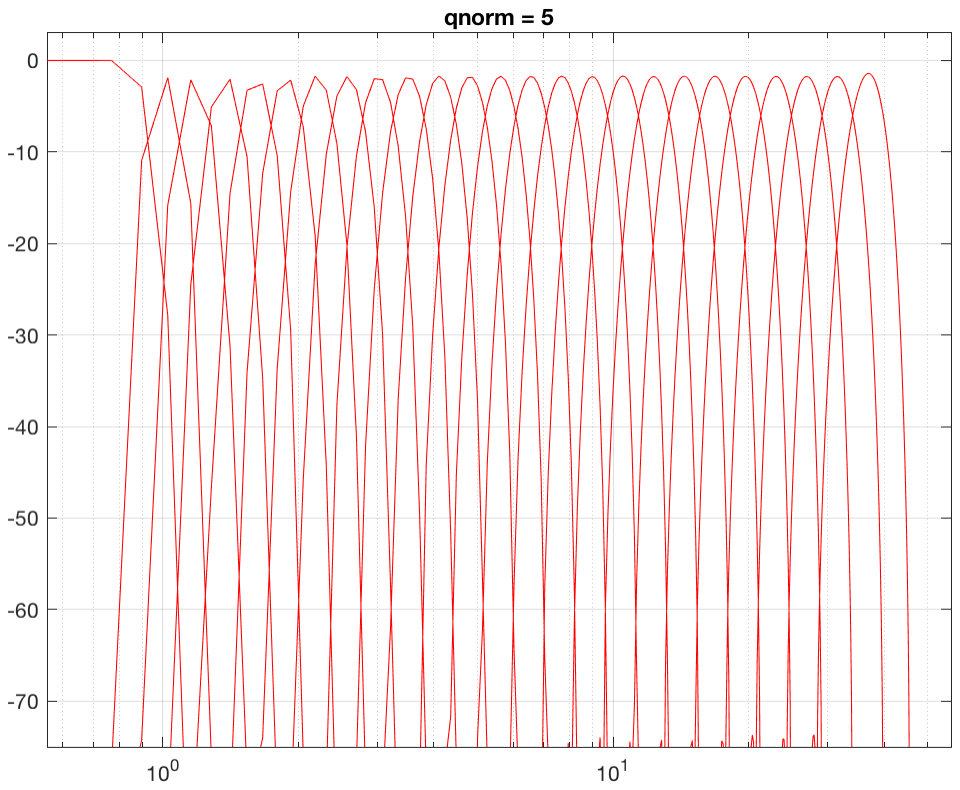
**Fig. b.** frequency response characteristics (5 filterbanks).

**Fig. a.** frequency response characteristics (24 filterbanks).
