## Supplementary material for "The Temporal Statistics of Musical Rhythm across Western Genres: An Amplitude Modulation Phase Hierarchy Model": S3 Appendix (from another version of this preprint: version 2, 2021-04-01)

**Individual Variation in Spectral PCA Component Loadings and the cumulative accounts**

Figure **a** shows the spectral PCA component loading patterns for each genre. In each subplot, the lines of different thickness indicate different PCA components. More important (lower numbered) components are shown in a thicker line. The loading patterns for the top 5 PCA components are similar across the 10 genres. It may be observed that the 10 genres produced consistent PCA loading patterns, particularly for the first 3 components. We, however, also find the individual variation at each spectral modulation band.

The present study considered the core spectral bands using the grand average PC loading patterns. In the grand average, a close peak of around 420-600 Hz and the identical flanking trough of ~700 Hz in PC3, PC4 and PC5, providing corroborating evidence for a second spectral band at this spectral location and a potential boundary between the second and third spectral bands at ~700 Hz. A close peak of 900-1100 Hz n PC3, PC4 and PC5, and a close flanking trough of 1750-2000 Hz in PC2, PC3 and PC5, provided corroborating evidence for a third spectral band at this spectral location and a potential boundary between the third and fourth spectral bands at ~1750 Hz. The peak of ~2500 Hz and the flanking trough of ~3900 Hz were identical between PC3 and PC5, providing corroborating evidence for a fourth spectral band at this spectral location and a potential boundary between the fourth and fifth spectral bands at ~3900 Hz. A peak of ~5500 Hz was identical between PC2 and PC3, providing corroborating evidence for a fifth spectral band at the spectral location. Further, this fifth peak was the highest peak observed in the spectral PCA loading patterns.

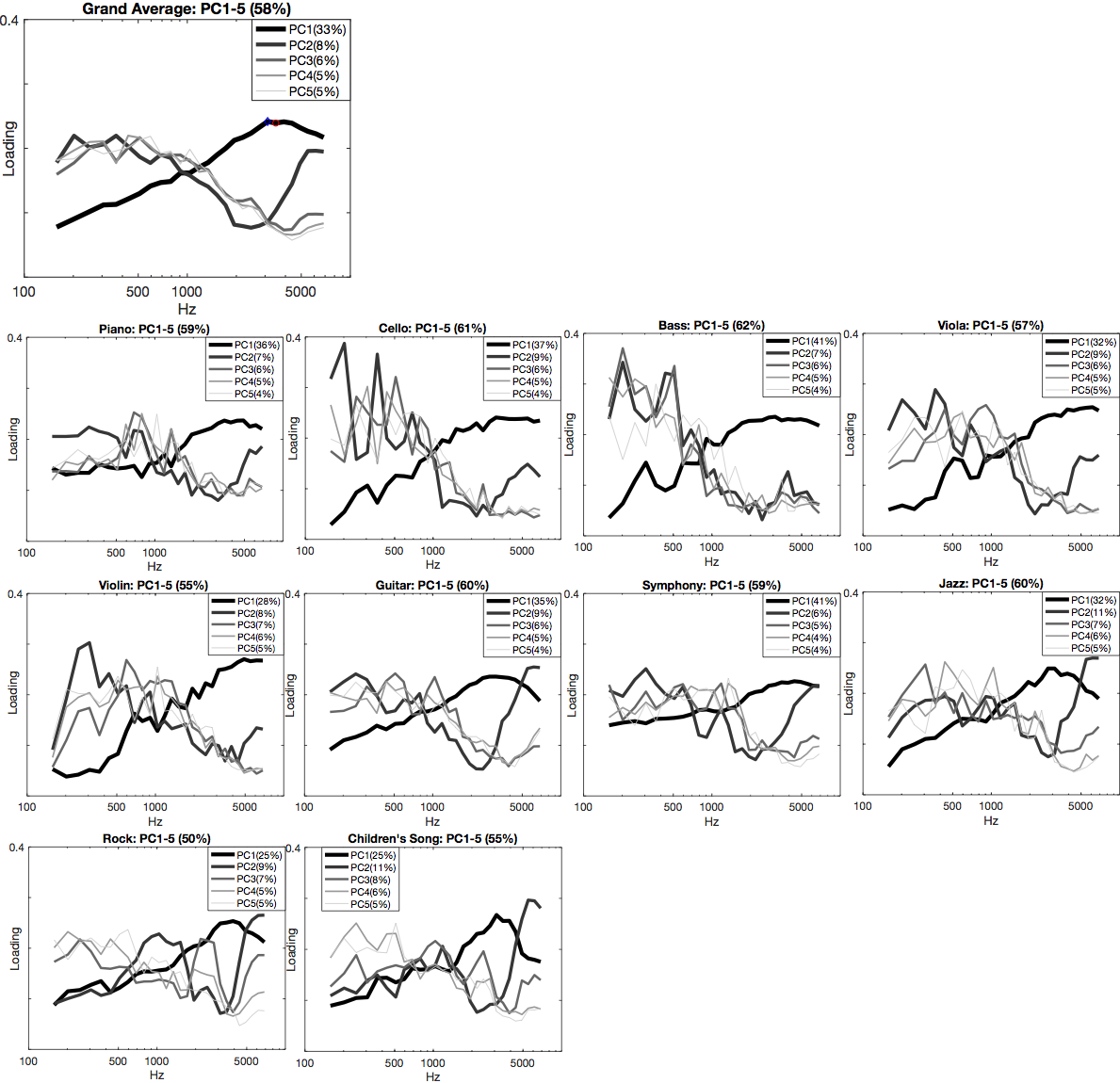

**Figure a**. Individual spectral PCA component loading patterns

**Table a. Summary of the 5 spectral bands and the 4 flanking boundaries indentified from spectral PCA.**

| **Spectral bands** | **Frequency range (Hz)** | **PC Peaks** |
| --- | --- | --- |
| Band 1 | 100-350 | PC2-PC5 |
| Band 2 | 350-700 | PC2-PC5 |
| Band 3 | 700-1750 | PC3-PC5 |
| Band 4 | 1750-3900 | PC1, PC3, PC5 |
| Band 5 | 3900-7250 | PC2, PC3 |

**Individual Variation in Temporal PCA Component Loadings**

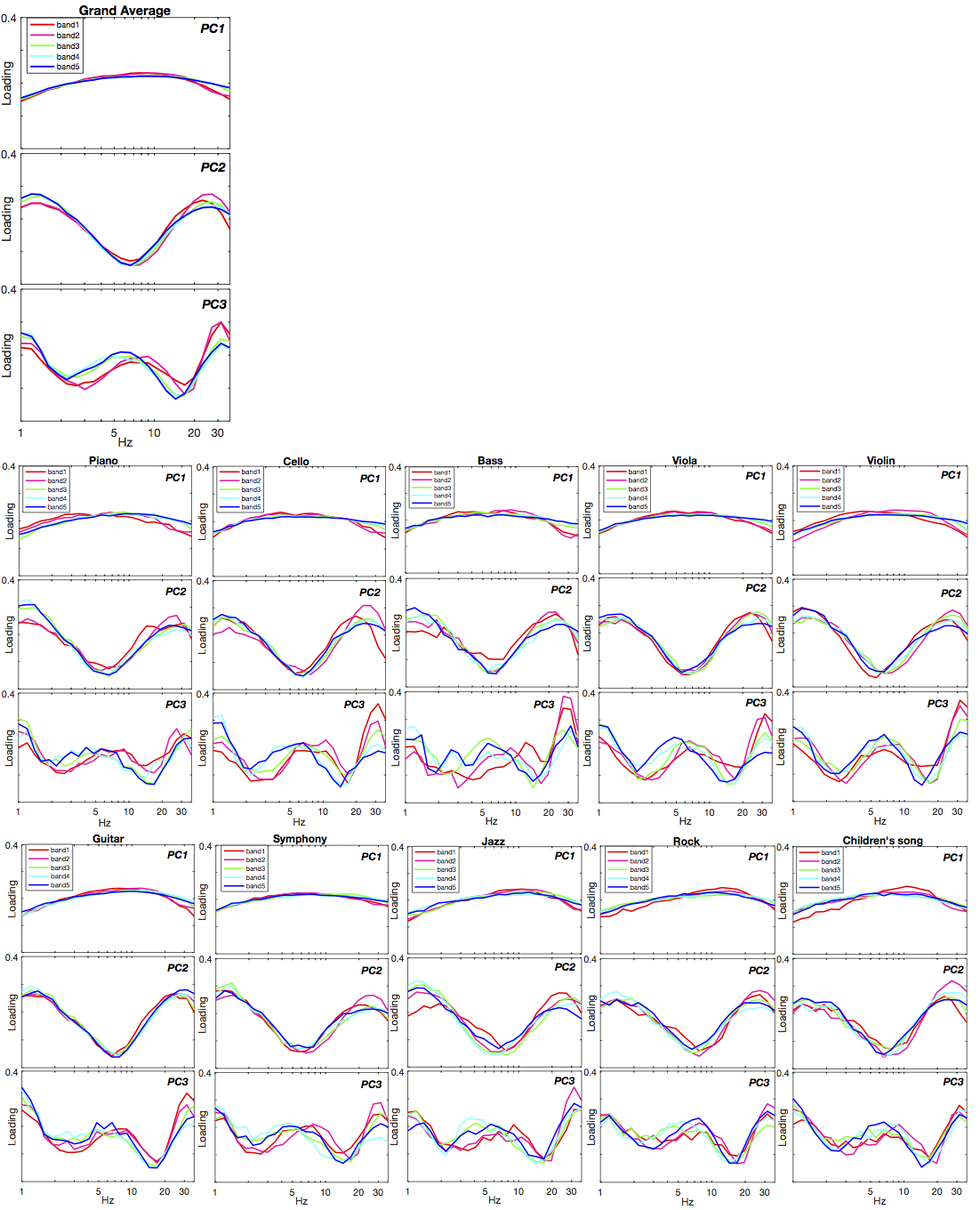
Figure **b** below shows the temporal PCA component loading patterns for each genre. In each subplot, the lines of different thickness indicate different PCA components. More important (lower numbered) components are shown in a thicker line, and the components from the 5 spectral bands are plotted in different colours. It may be observed that the 10 genres produced consistent PCA loading patterns. We, however, also find the individual variation at each spectral modulation band.

**Figure b.** Individual temporal PCA component loading patterns

**Table b. Cumulative accounts of temporal PCA of 5 bands indentified from spectral PCA.**

| **Genre** | **PC** | **band1** | **band2** | **band3** | **band4** | **band5** | **mean** |
| --- | --- | --- | --- | --- | --- | --- | --- |
| Grand average | PC1 | 0.43 | 0.44 | 0.48 | 0.52 | 0.56 | 0.486 |
| Piano | PC2 | 0.11 | 0.11 | 0.10 | 0.10 | 0.11 | 0.106 |
|  | PC3 | 0.07 | 0.07 | 0.06 | 0.05 | 0.05 | 0.06 |
|  | PC1 | 0.44 | 0.45 | 0.55 | 0.58 | 0.64 | 0.532 |
|  | PC2 | 0.12 | 0.11 | 0.11 | 0.11 | 0.12 | 0.114 |
|  | PC3 | 0.07 | 0.07 | 0.06 | 0.05 | 0.05 | 0.06 |
| Cello | PC1 | 0.40 | 0.42 | 0.46 | 0.53 | 0.58 | 0.478 |
|  | PC2 | 0.11 | 0.11 | 0.09 | 0.09 | 0.09 | 0.098 |
|  | PC3 | 0.07 | 0.07 | 0.06 | 0.05 | 0.05 | 0.06 |
| Bass | PC1 | 0.38 | 0.39 | 0.49 | 0.53 | 0.58 | 0.474 |
|  | PC2 | 0.11 | 0.12 | 0.11 | 0.09 | 0.09 | 0.104 |
|  | PC3 | 0.08 | 0.07 | 0.06 | 0.05 | 0.05 | 0.062 |
| Viola | PC1 | 0.48 | 0.47 | 0.49 | 0.53 | 0.59 | 0.512 |
|  | PC2 | 0.12 | 0.12 | 0.10 | 0.09 | 0.09 | 0.104 |
|  | PC3 | 0.06 | 0.06 | 0.05 | 0.05 | 0.05 | 0.054 |
| Violin | PC1 | 0.54 | 0.42 | 0.47 | 0.49 | 0.54 | 0.492 |
|  | PC2 | 0.13 | 0.12 | 0.10 | 0.10 | 0.10 | 0.11 |
|  | PC3 | 0.06 | 0.07 | 0.06 | 0.06 | 0.05 | 0.06 |
| Guitar | PC1 | 0.49 | 0.48 | 0.51 | 0.56 | 0.62 | 0.532 |
|  | PC2 | 0.12 | 0.11 | 0.11 | 0.11 | 0.13 | 0.116 |
|  | PC3 | 0.07 | 0.06 | 0.06 | 0.05 | 0.05 | 0.058 |
| Synphony | PC1 | 0.48 | 0.46 | 0.48 | 0.52 | 0.55 | 0.498 |
|  | PC2 | 0.10 | 0.10 | 0.09 | 0.09 | 0.10 | 0.096 |
|  | PC3 | 0.05 | 0.06 | 0.05 | 0.05 | 0.05 | 0.052 |
| Jazz | PC1 | 0.42 | 0.45 | 0.53 | 0.55 | 0.52 | 0.494 |
|  | PC2 | 0.12 | 0.11 | 0.12 | 0.11 | 0.12 | 0.116 |
|  | PC3 | 0.07 | 0.06 | 0.05 | 0.05 | 0.06 | 0.058 |
| Rock | PC1 | 0.32 | 0.38 | 0.37 | 0.38 | 0.44 | 0.378 |
|  | PC2 | 0.11 | 0.11 | 0.10 | 0.10 | 0.14 | 0.112 |
|  | PC3 | 0.07 | 0.06 | 0.06 | 0.06 | 0.07 | 0.064 |
| Children's song | PC1 | 0.37 | 0.49 | 0.47 | 0.51 | 0.53 | 0.474 |
|  | PC2 | 0.11 | 0.11 | 0.12 | 0.12 | 0.15 | 0.122 |
|  | PC3 | 0.08 | 0.06 | 0.07 | 0.06 | 0.06 | 0.066 |

**Table c. Summary of the 4 temporal bands and the 3 flanking boundaries indentified from temporal PCA.**

Using an ERB_N_ filterbank that simulated the frequency decomposition by the cochlea, the S-AMPH model generated an hierarchical representation of the core spectral (acoustic frequency spanning 100–7,250 Hz) and temporal (oscillatory rate spanning 0.9–40 Hz) modulation architectures in the envelope of a range of musical genres and instruments. In each genre and for each instrument, the AM hierarchies consisted of 5 spectral bands (100–350 Hz; 350–700 Hz; 700–1750 Hz; 1750–3900 Hz; 3900–7250 Hz), each containing 4 hierarchically-nested temporal bands (0.9–2.5 Hz, 2.5–7 Hz, 7–17 Hz, 17–40 Hz).

| **Temporal bands** | **Frequency range (Hz)** | **PC Peaks** |
| --- | --- | --- |
| Band 1 | 0.9-2.5 | PC2, PC3 in spectral band 1-5 |
| Band 2 | 2.5-7 | PC3 in spectral band 3-5 |
| Band 3 | 7-17 | PC3 in spectral band 1-2 |
| Band 4 | 17-40 | PC2, PC3 in spectral band 1-5 |

**Individual modulation spectra of FFT in the PAD model**

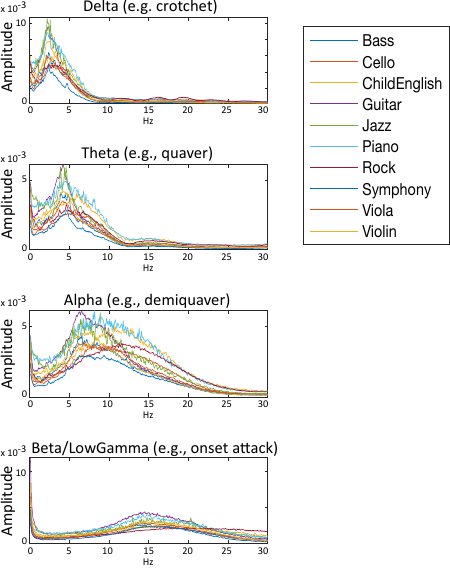
Figure **c** below shows the modulation spectra of FFT for each genre. The 10 genres are plotted in different colours. It may be observed that the 10 genres produced broadly consistent modulation spectra of FFT.

**Figure c.** Individual modulation spectra of FFT
