## Supplementary material for "The Temporal Statistics of Musical Rhythm across Western Genres: An Amplitude Modulation Phase Hierarchy Model": S4 Appendix (from another version of this preprint: version 2, 2021-04-01)

**Mutual Information Between different Bands in S-AMPH model**

Figure **a** shows the mutual information (MI) between different bands in S-AMPH model for each genre. The MI shown here are broadly similar across the 10 genres. Figure **b** shows the MI of non-music natural sounds such as rain and wind.

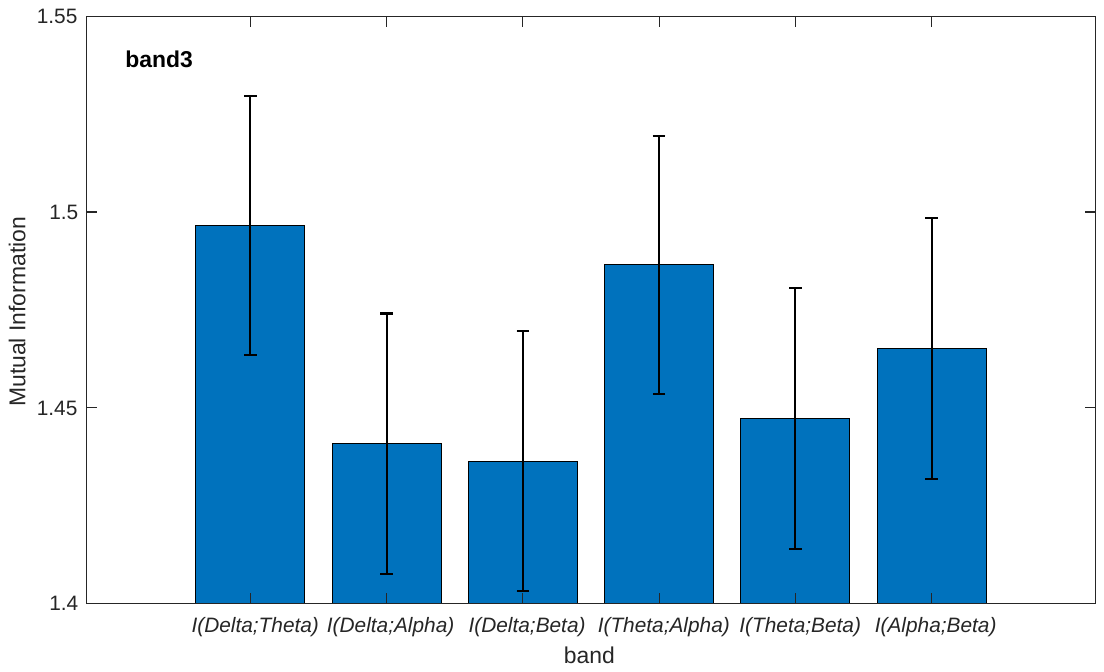

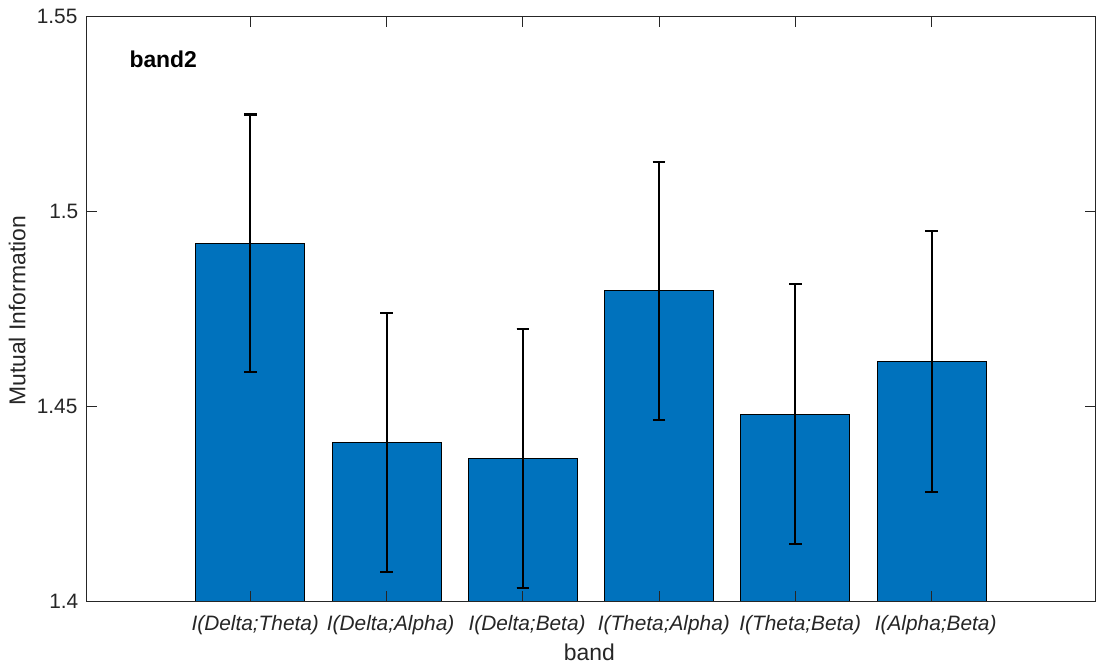

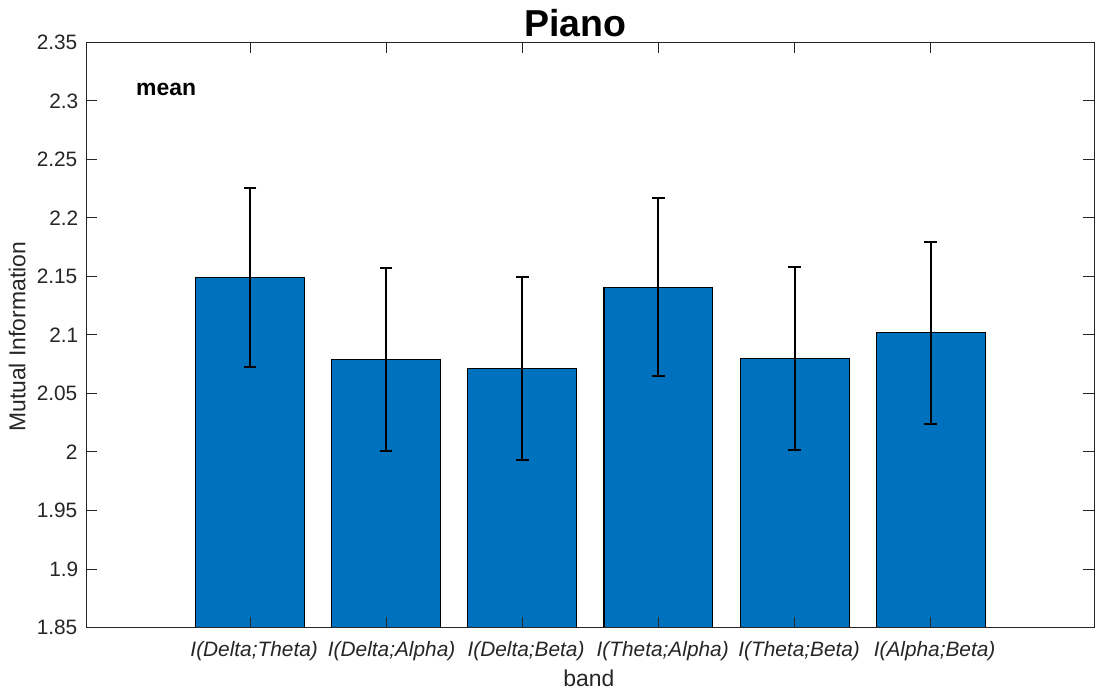

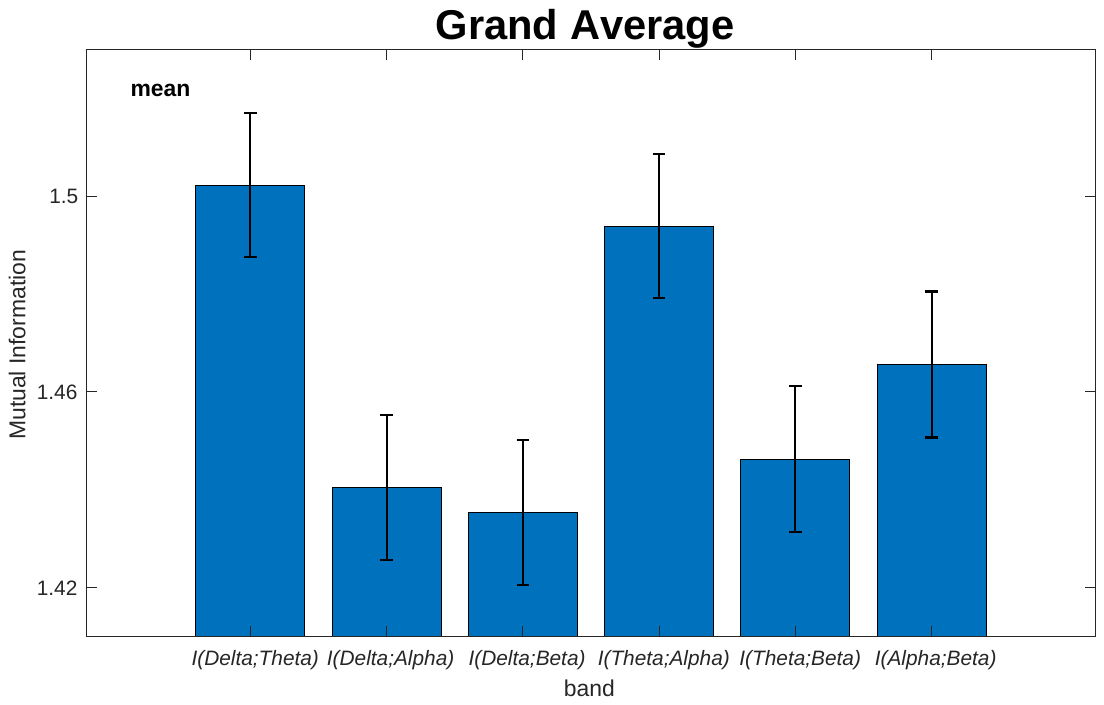

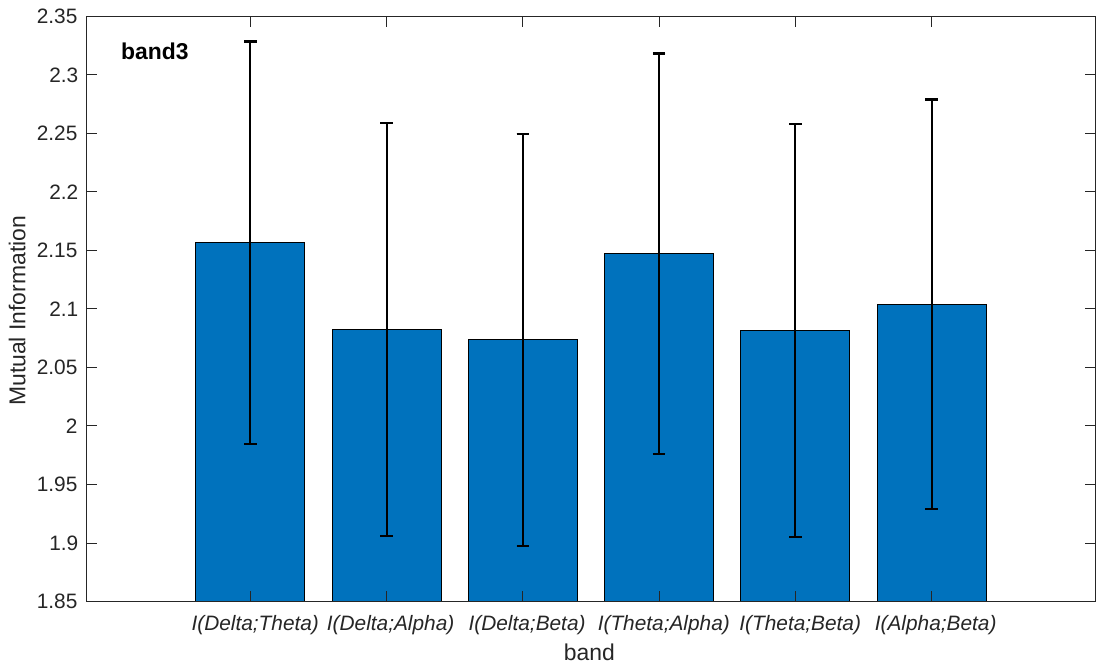

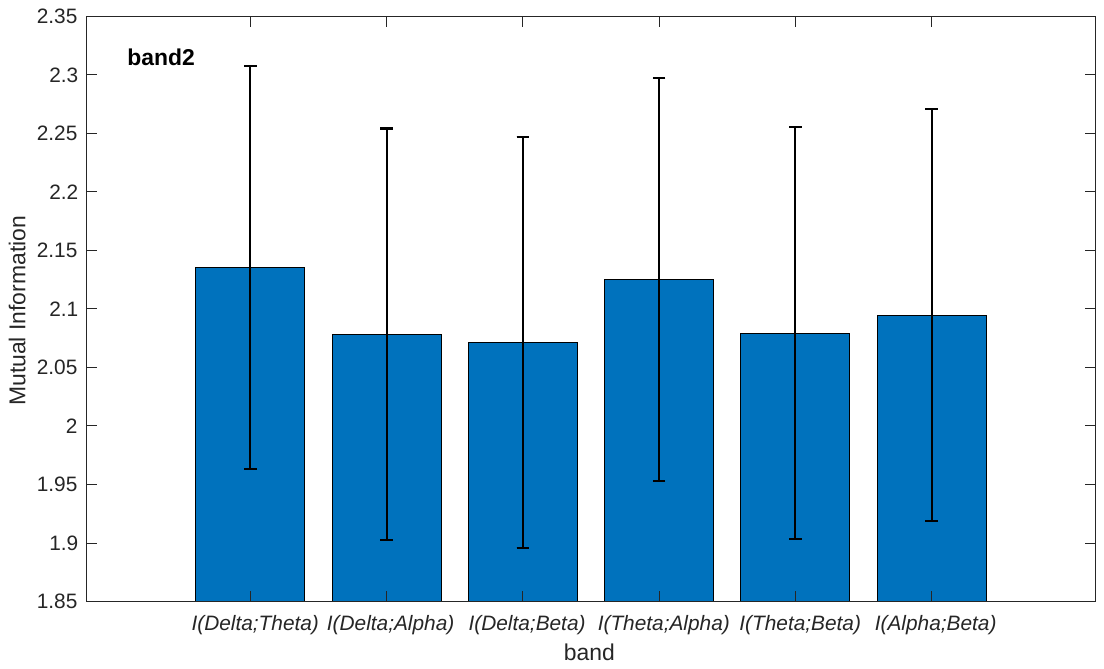

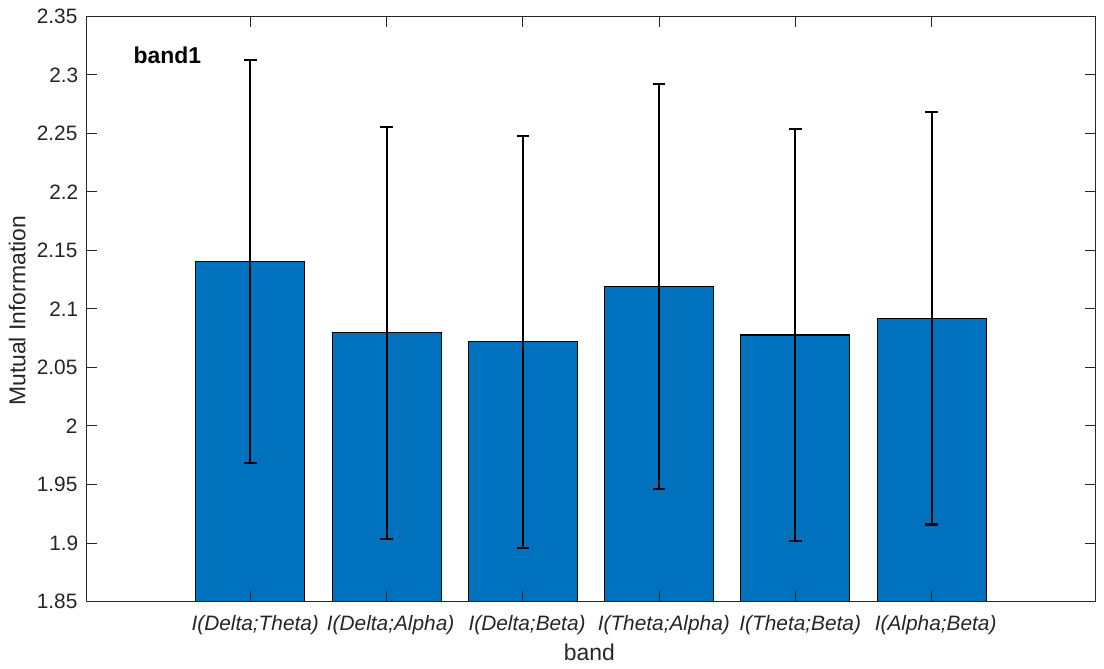

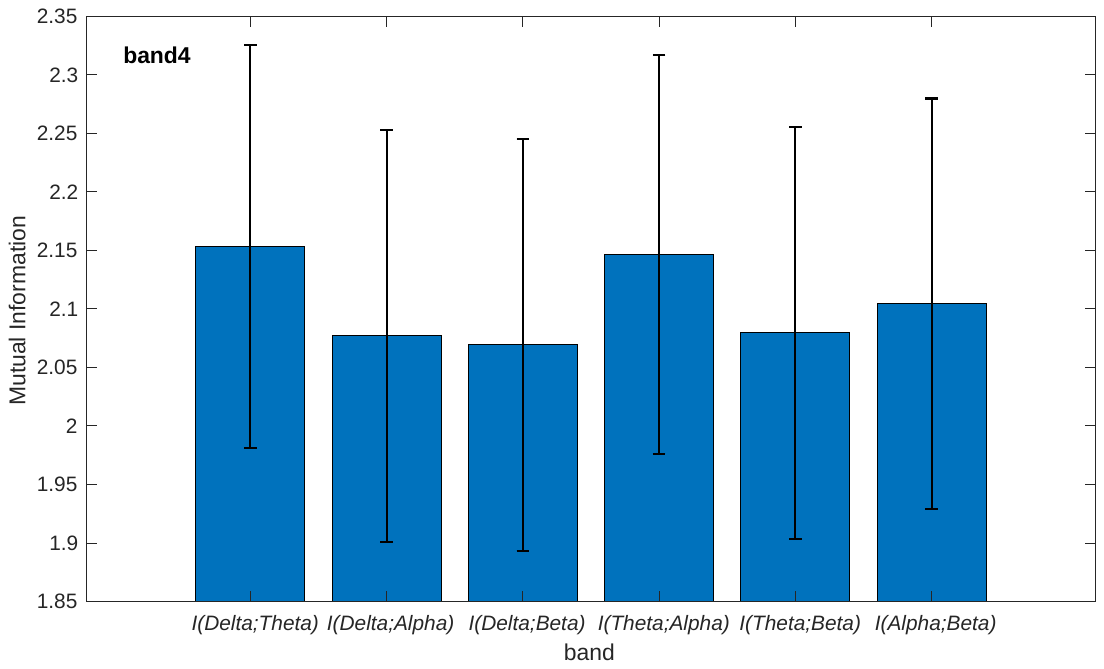

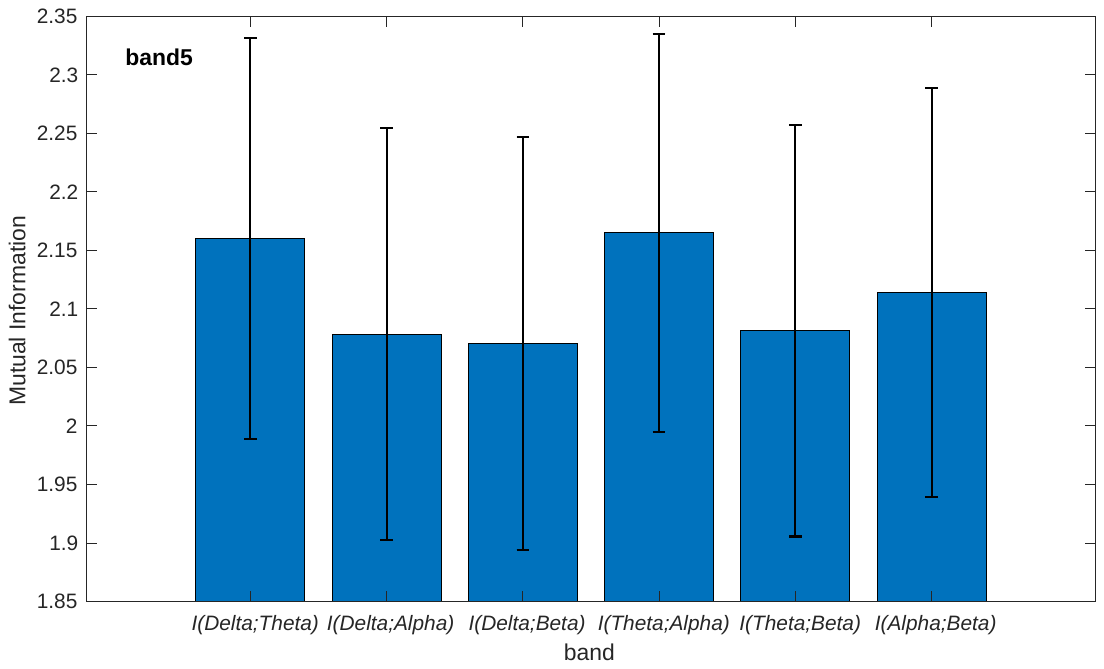

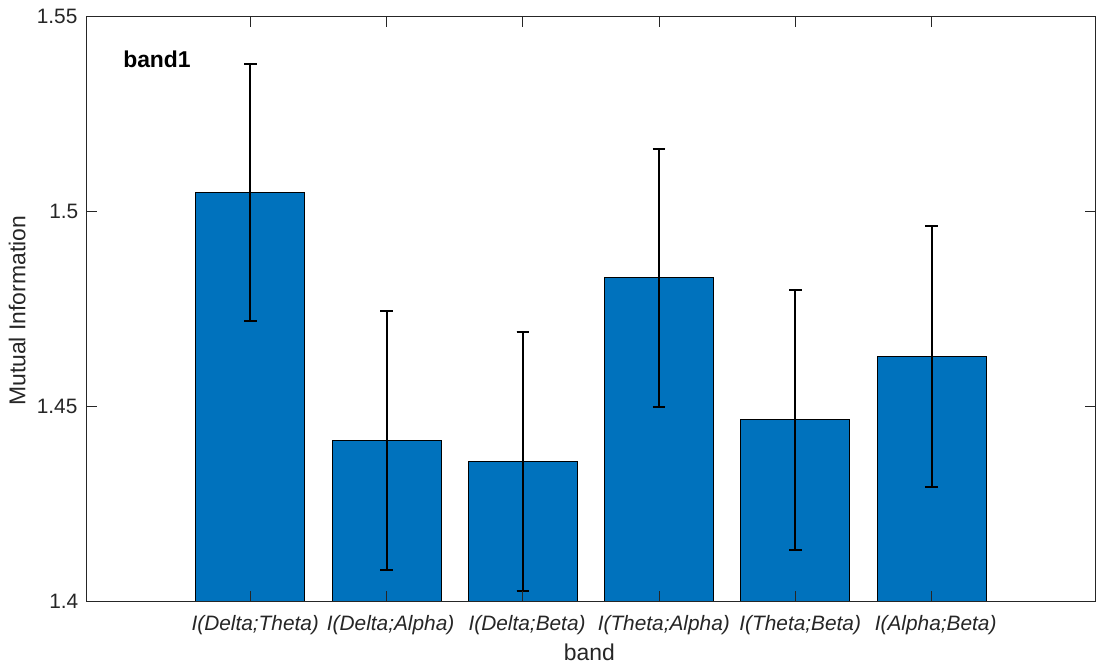

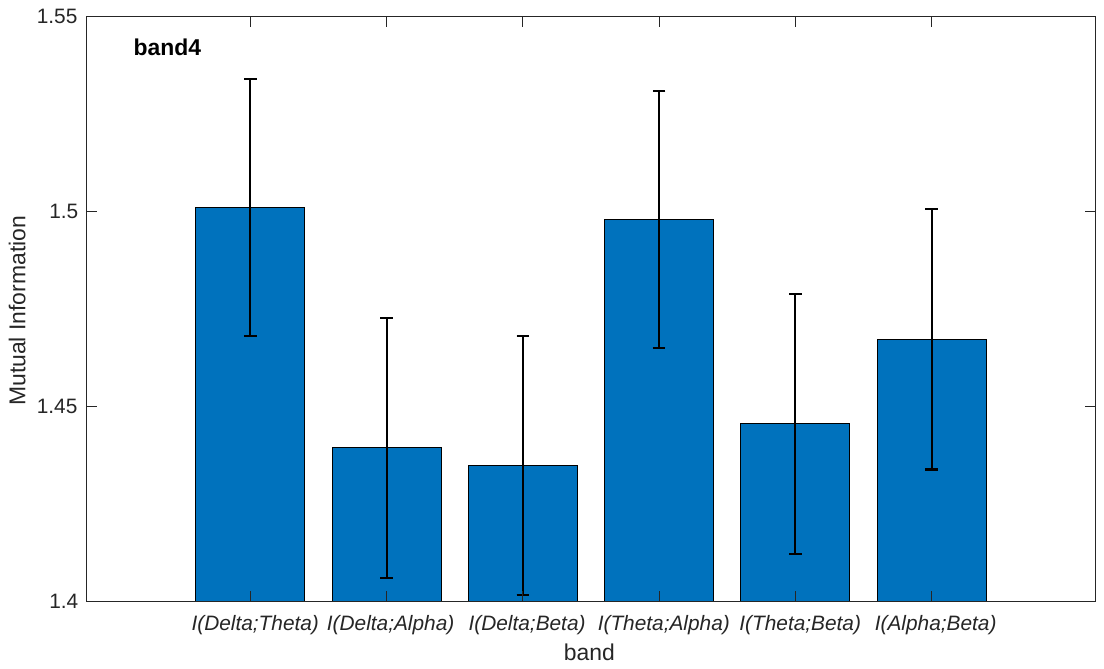

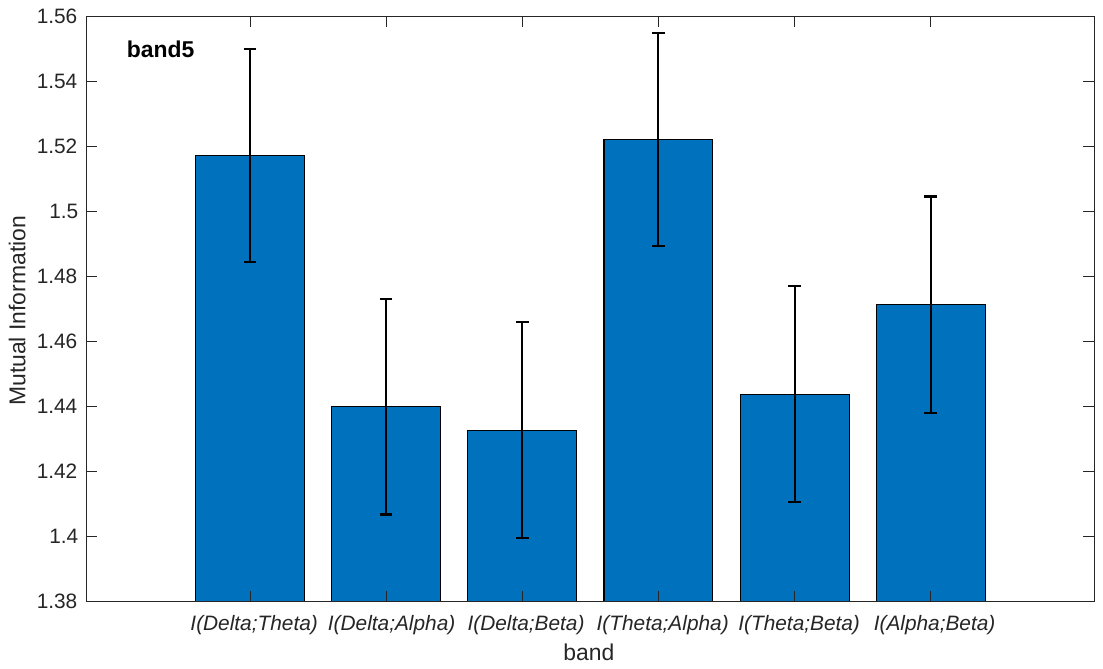

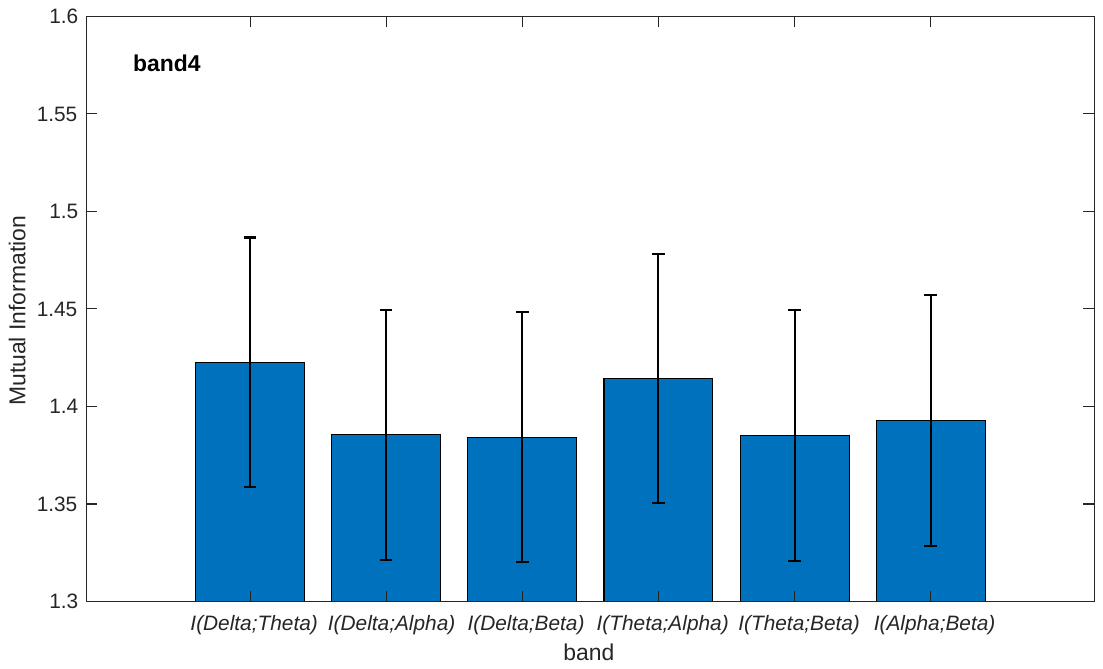

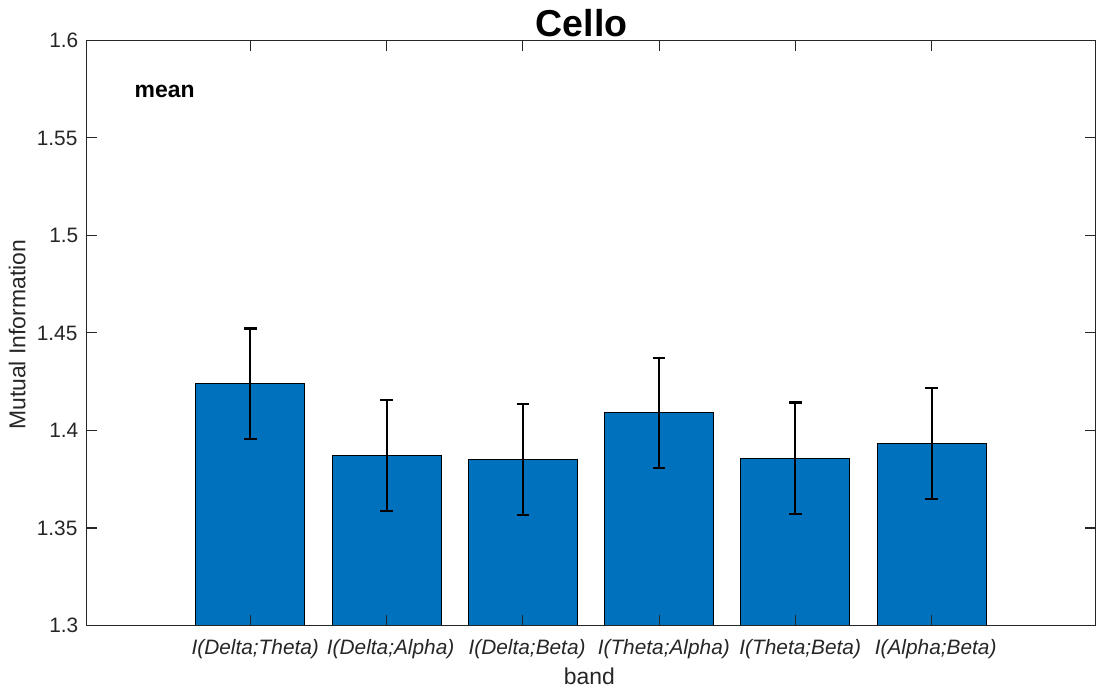

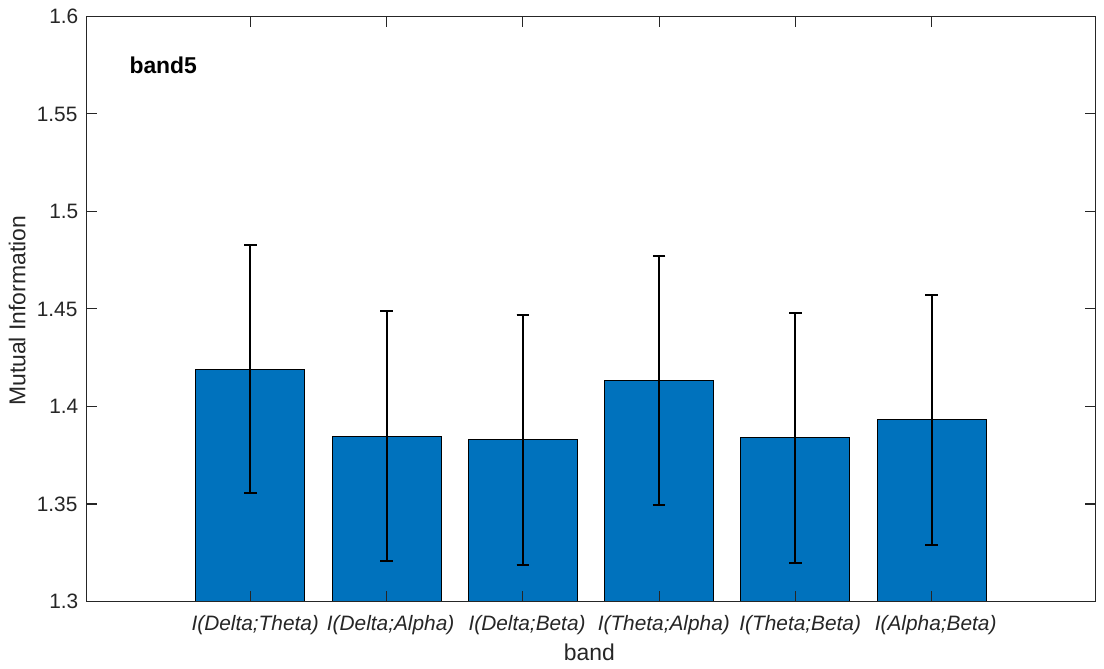

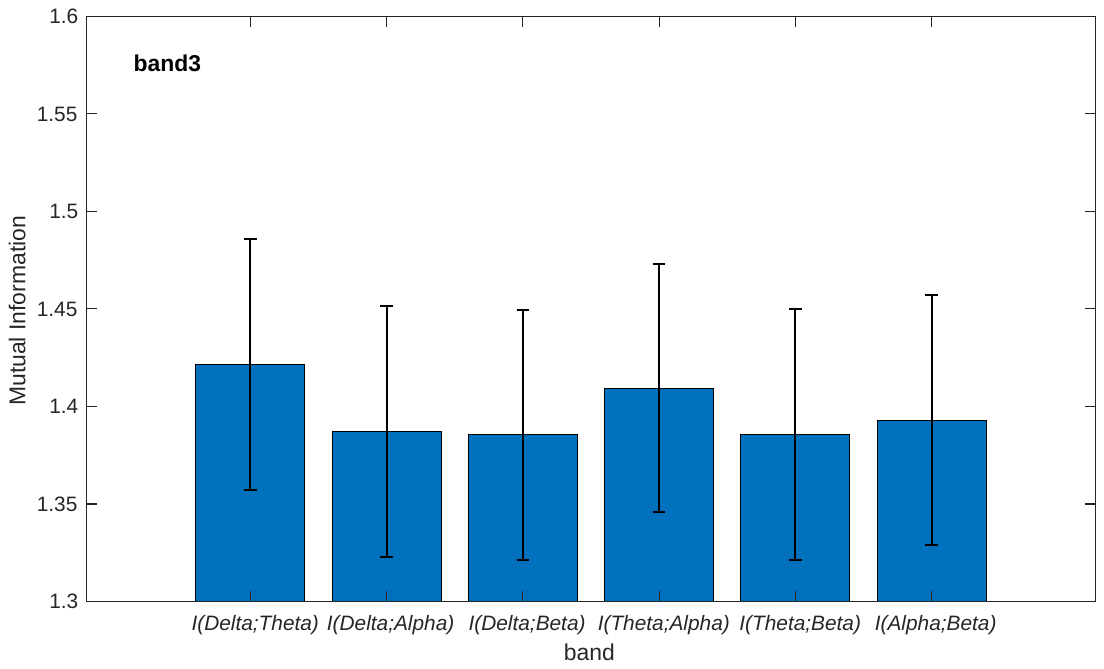

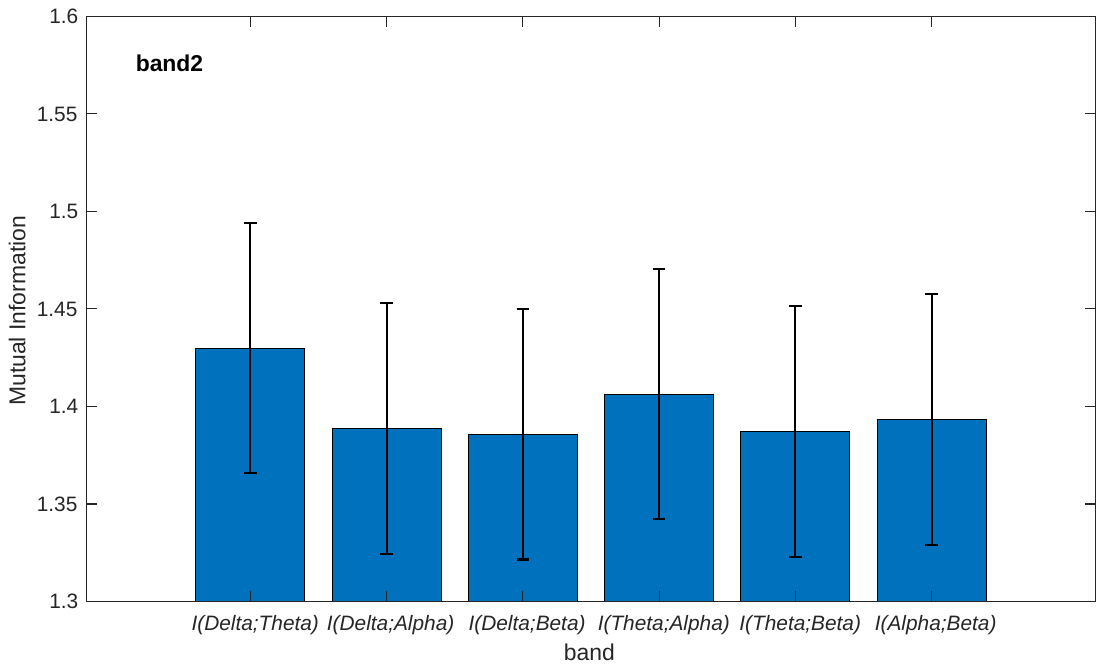

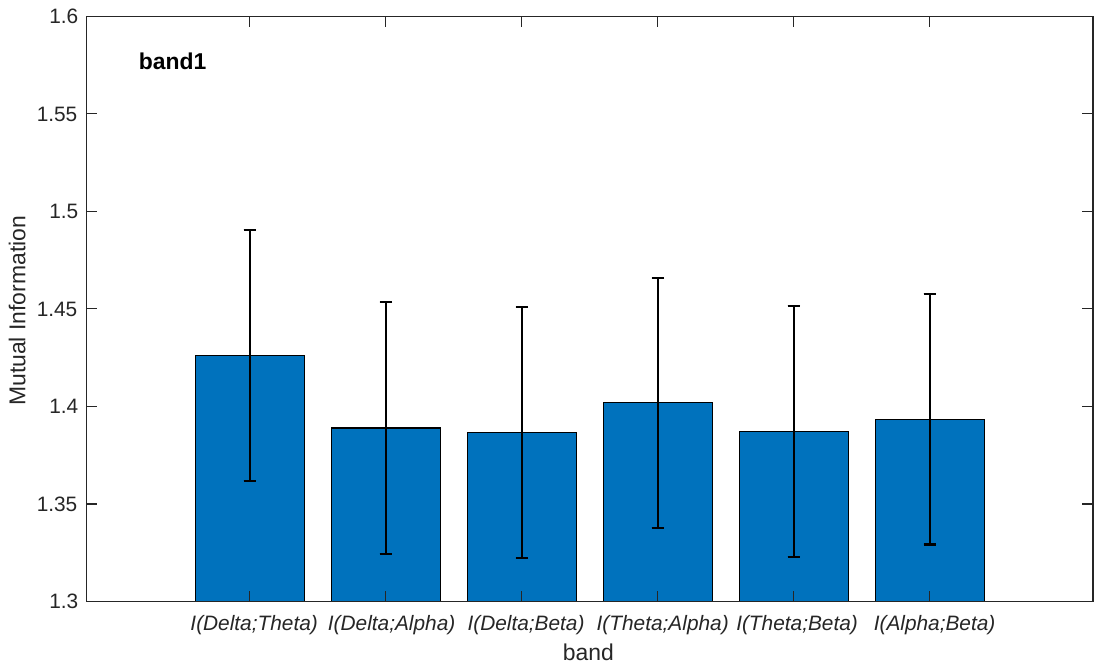

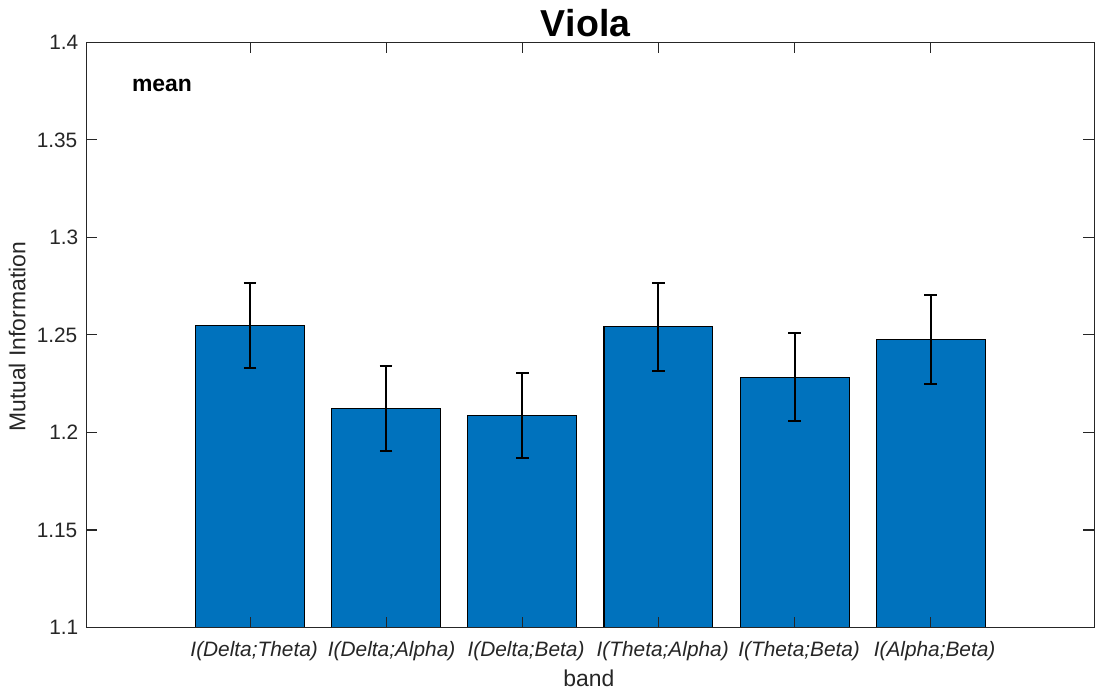

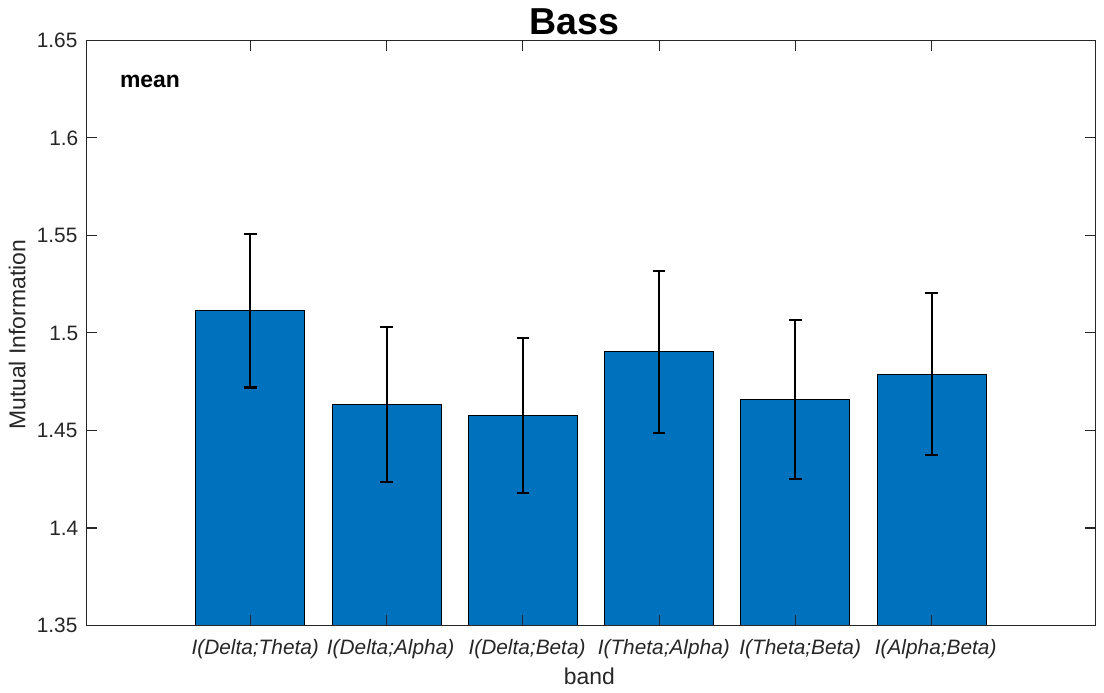

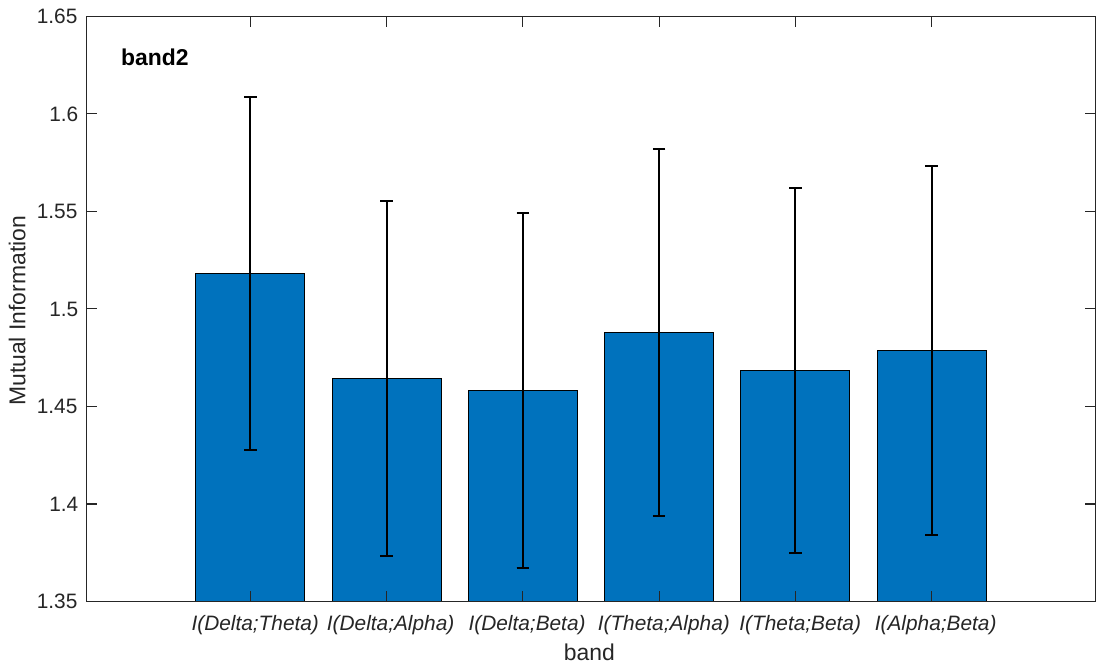

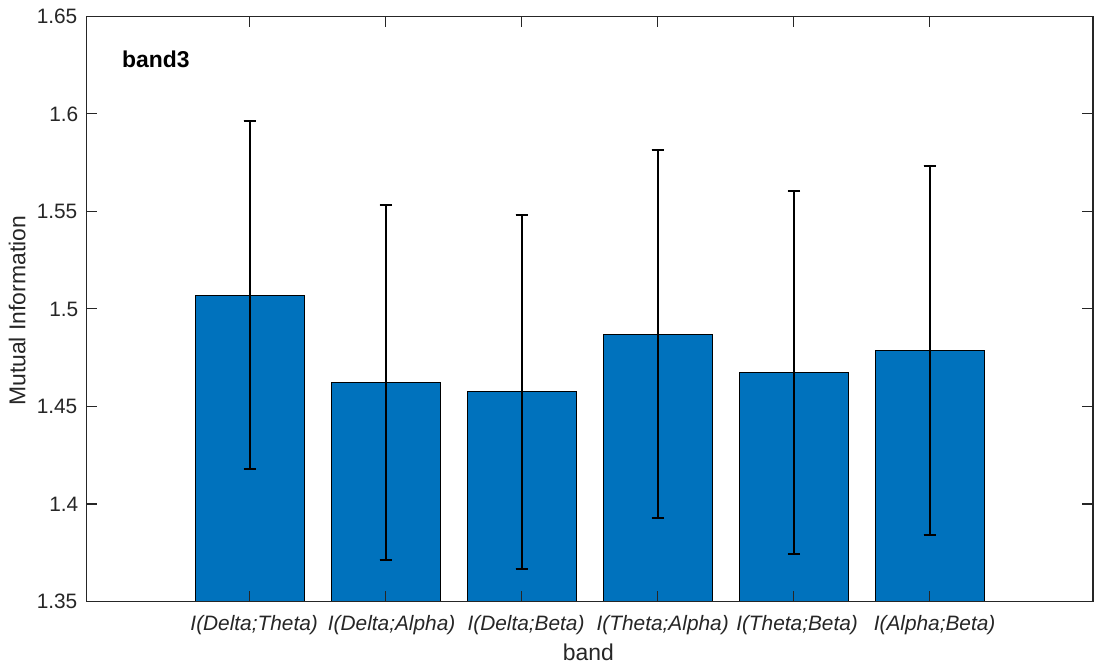

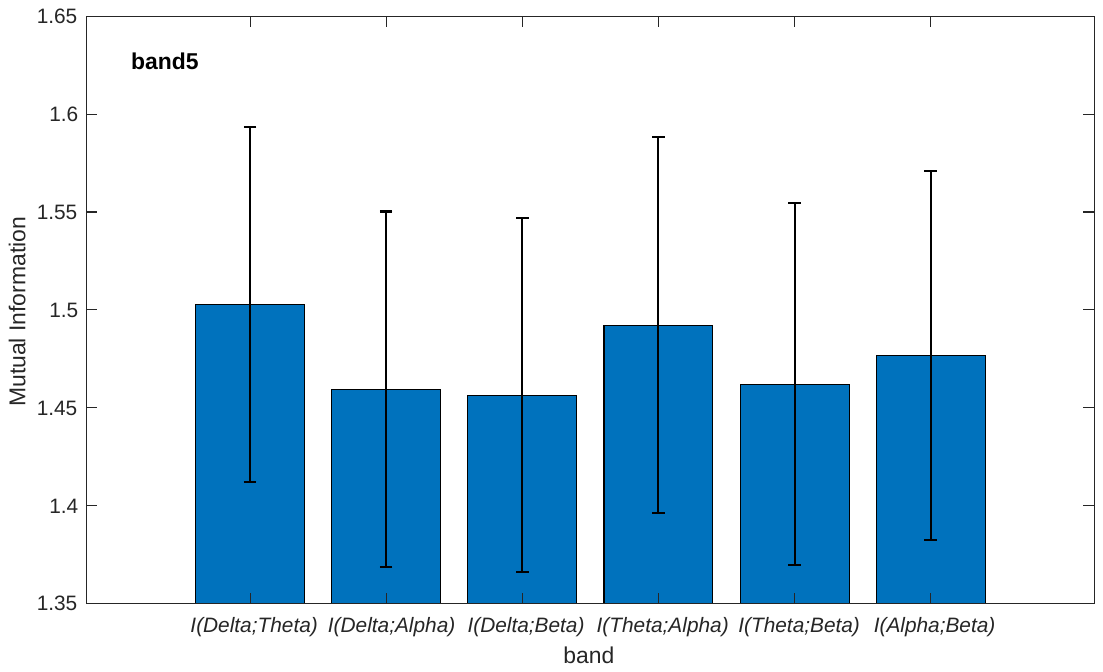

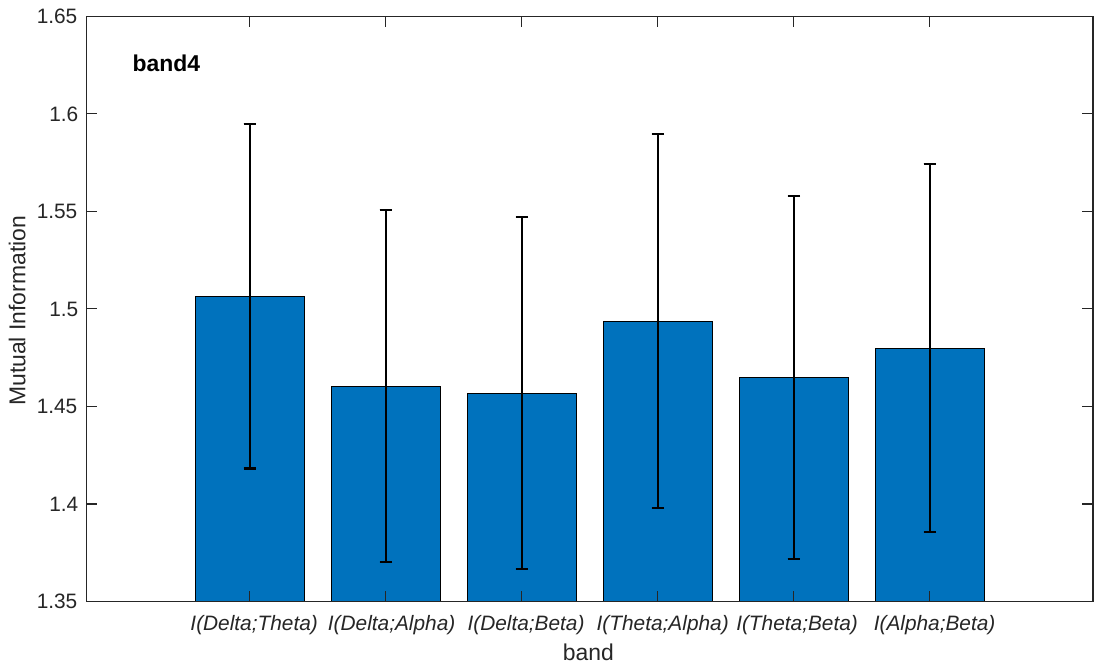

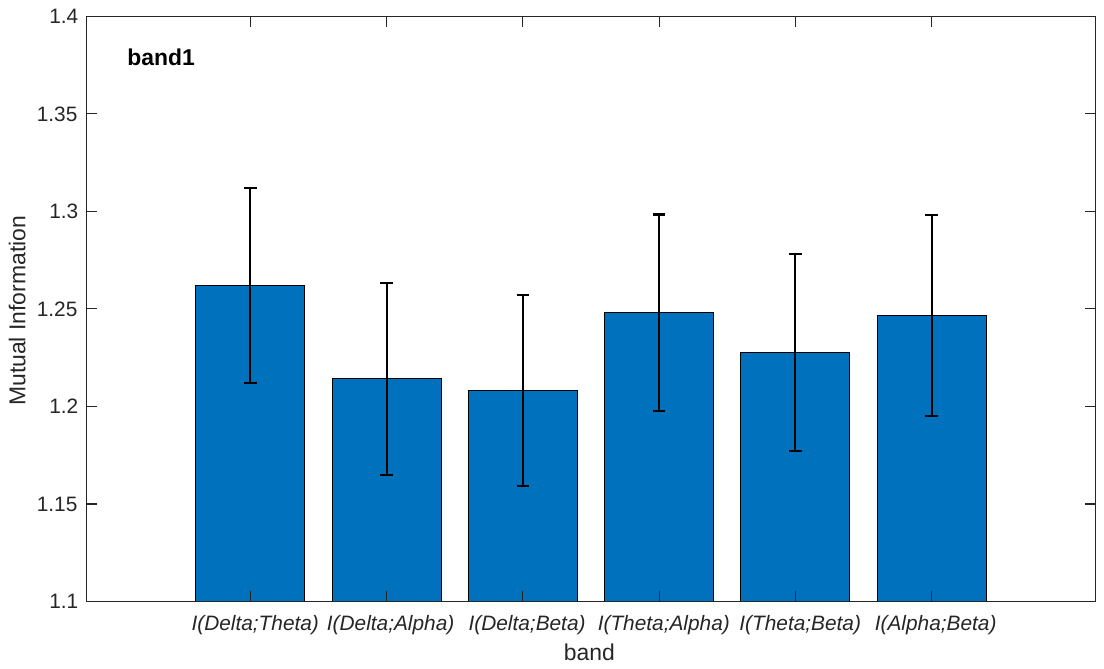

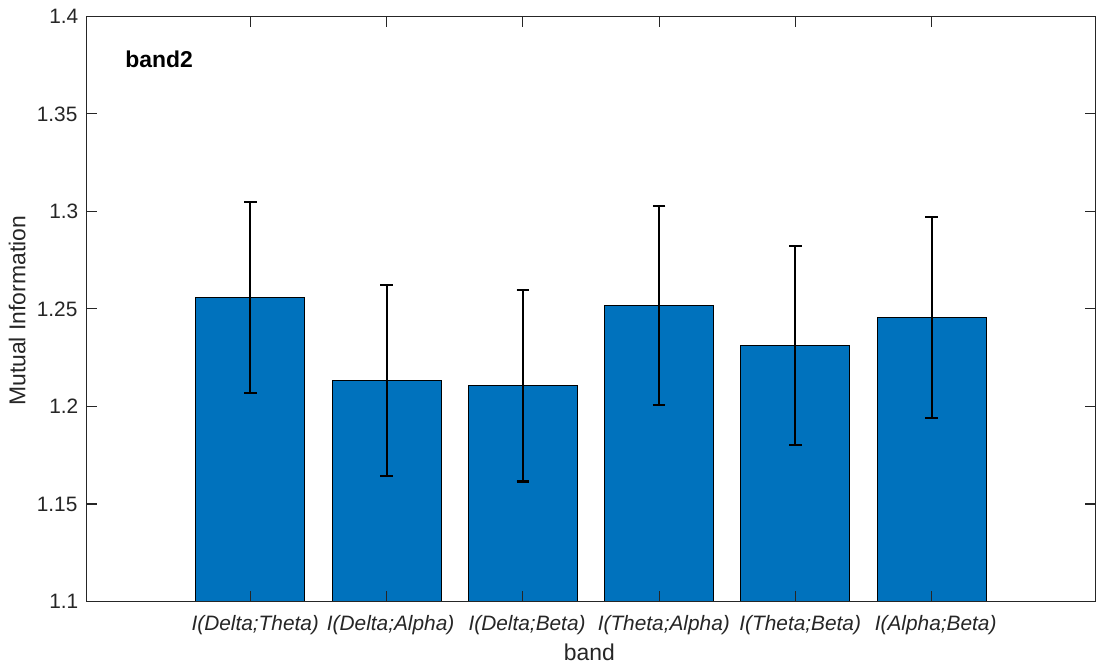

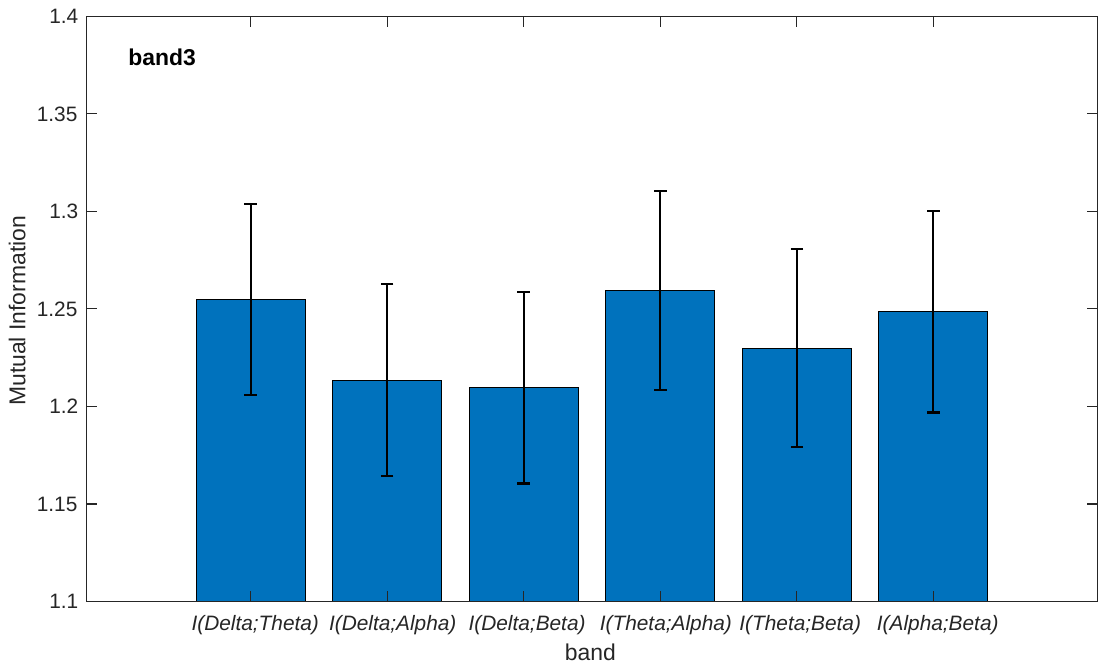

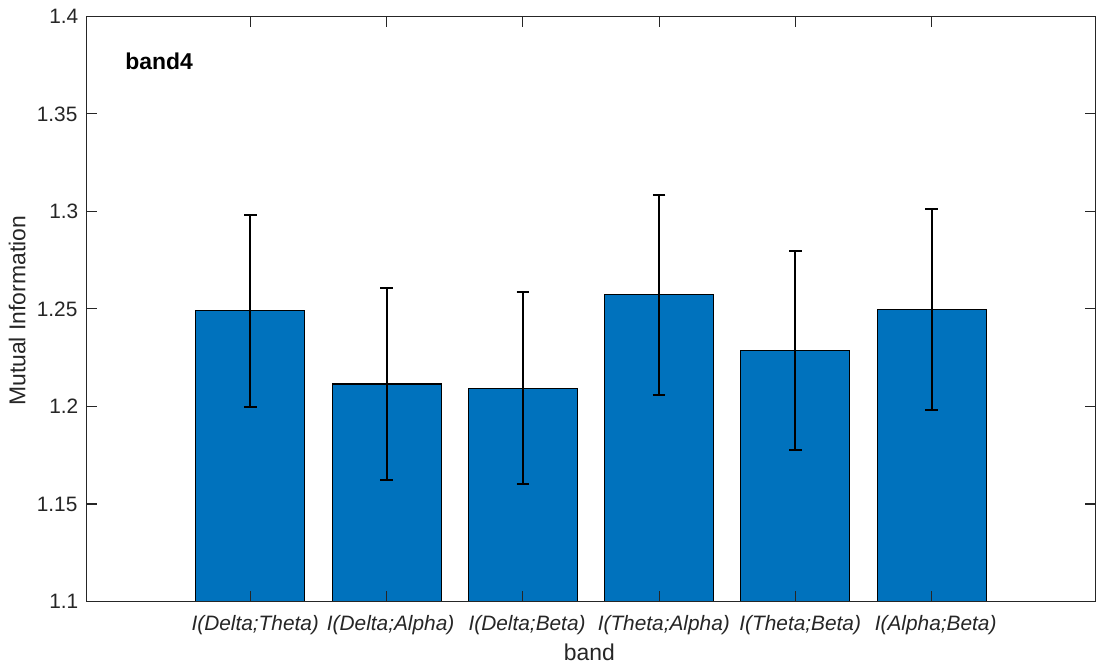

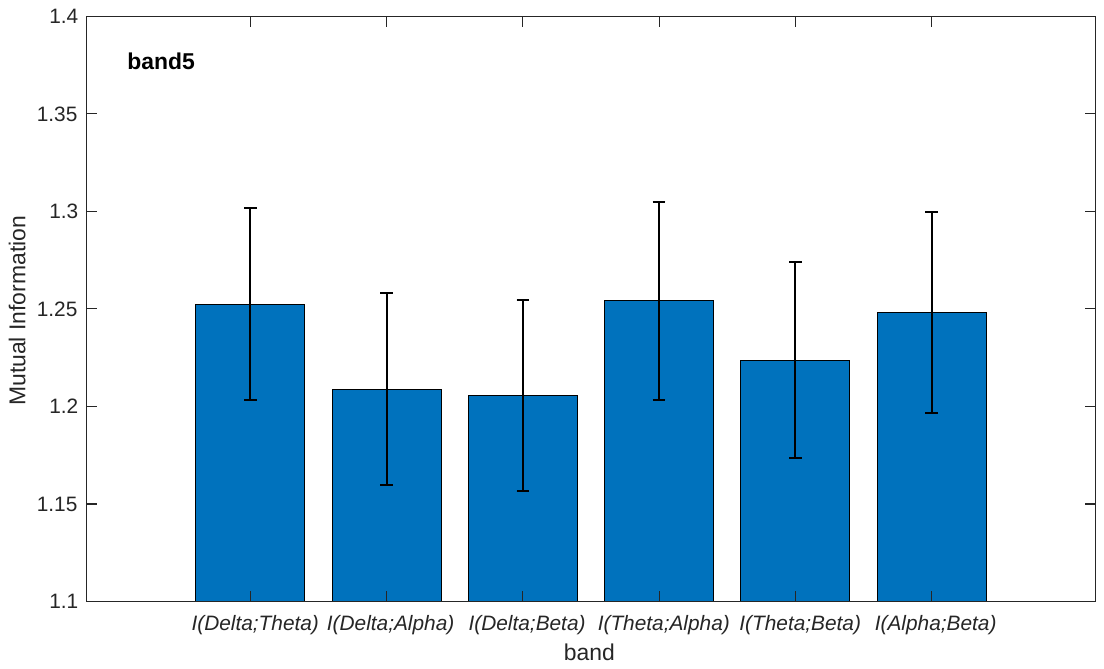

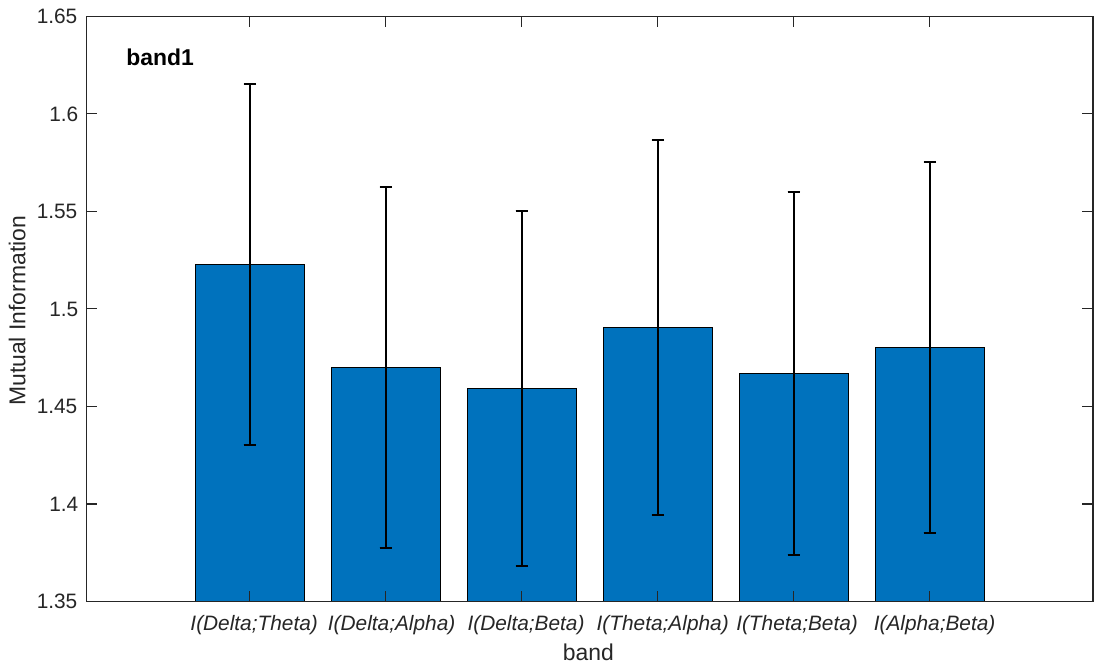

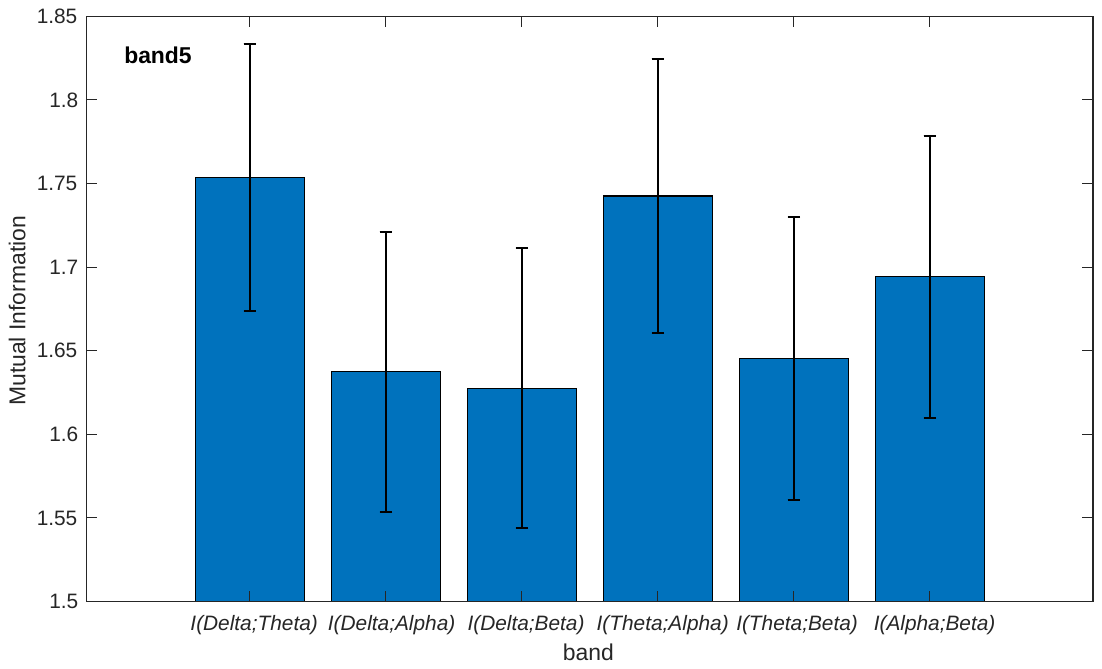

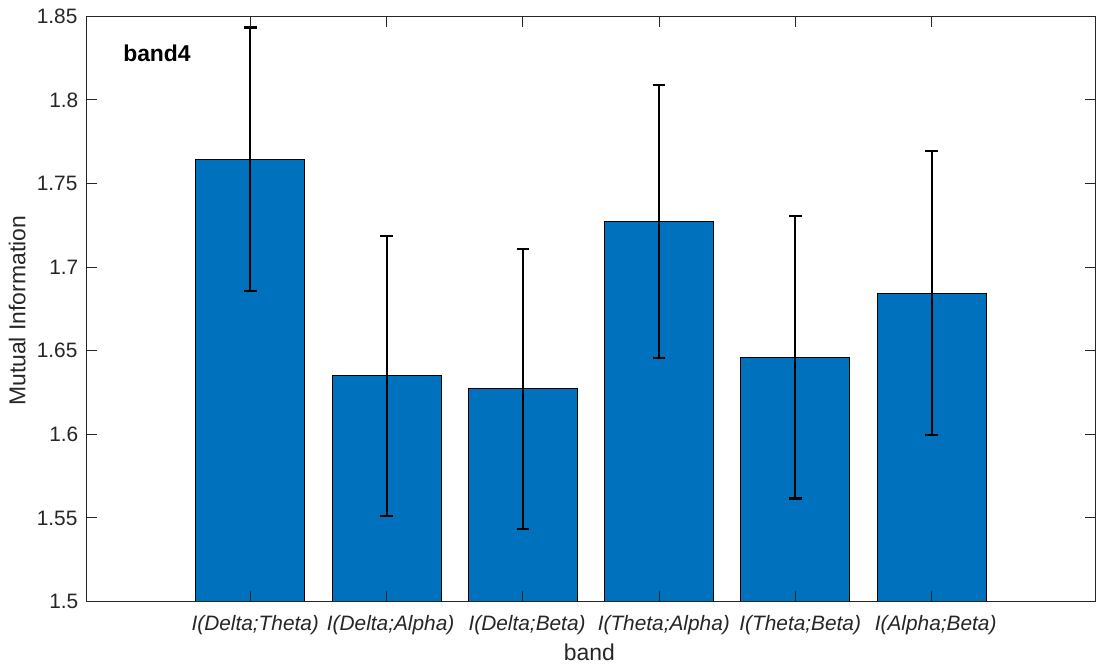

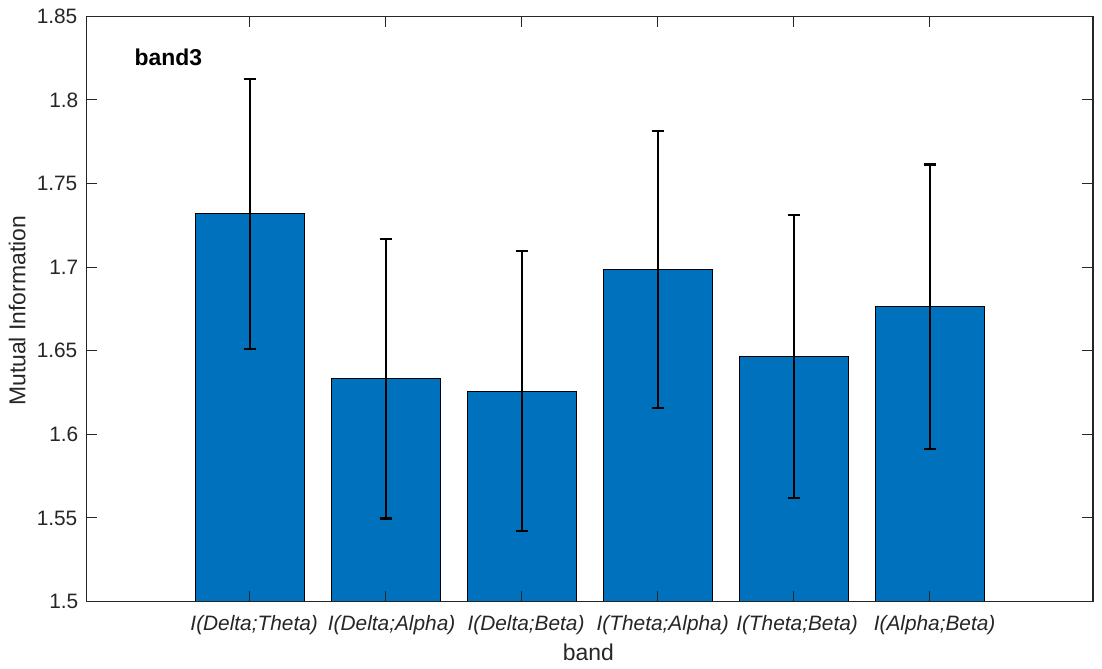

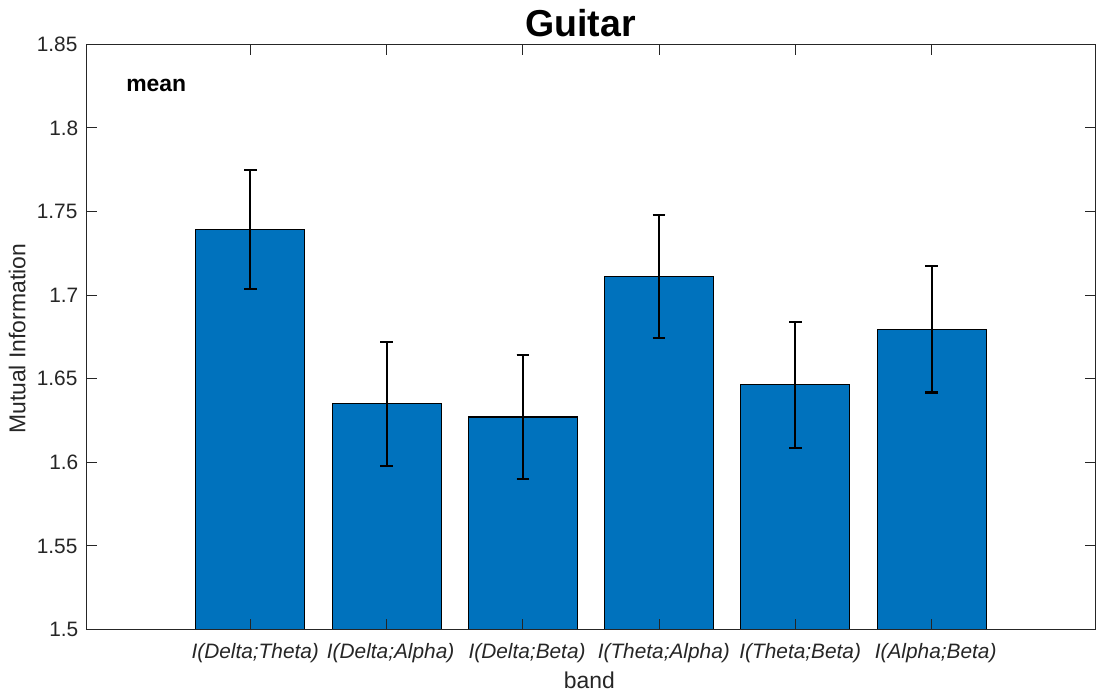

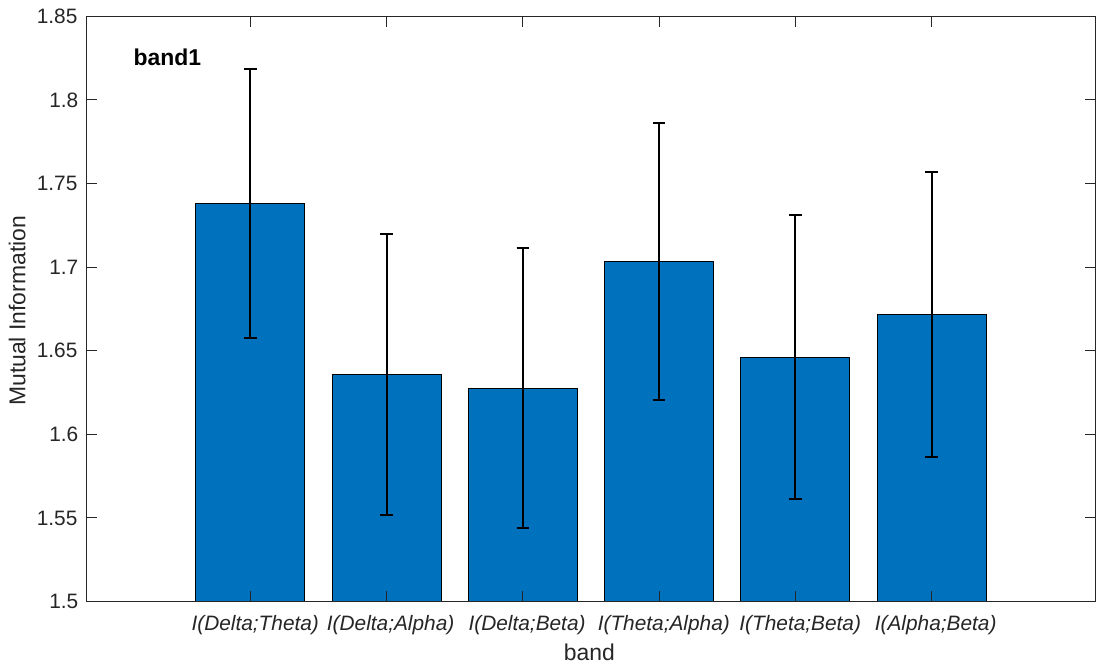

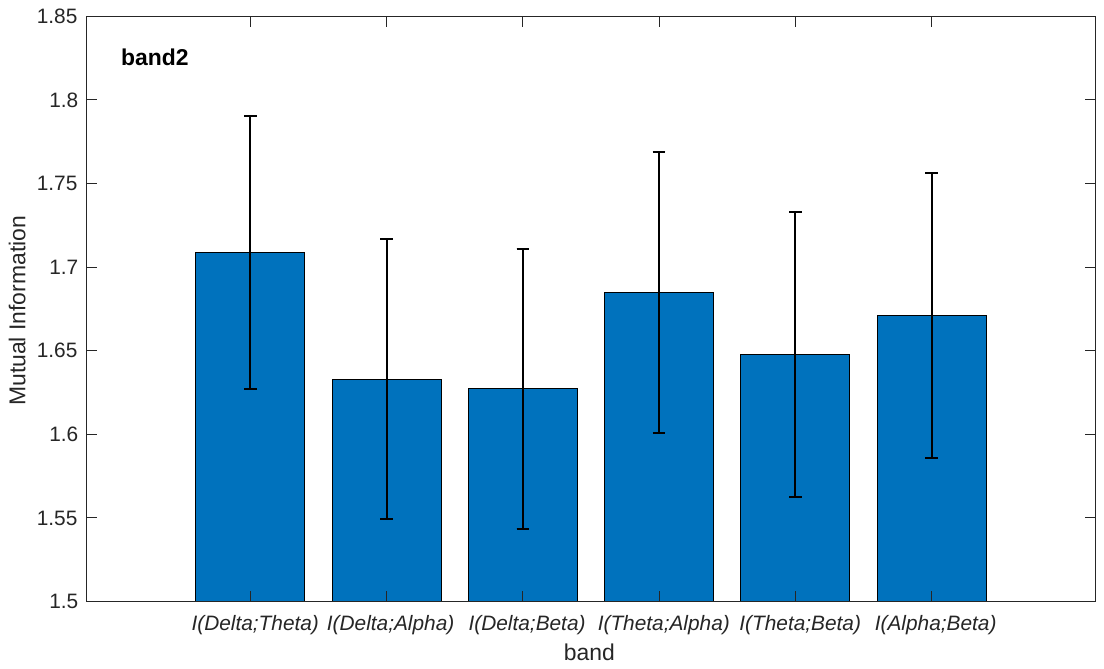

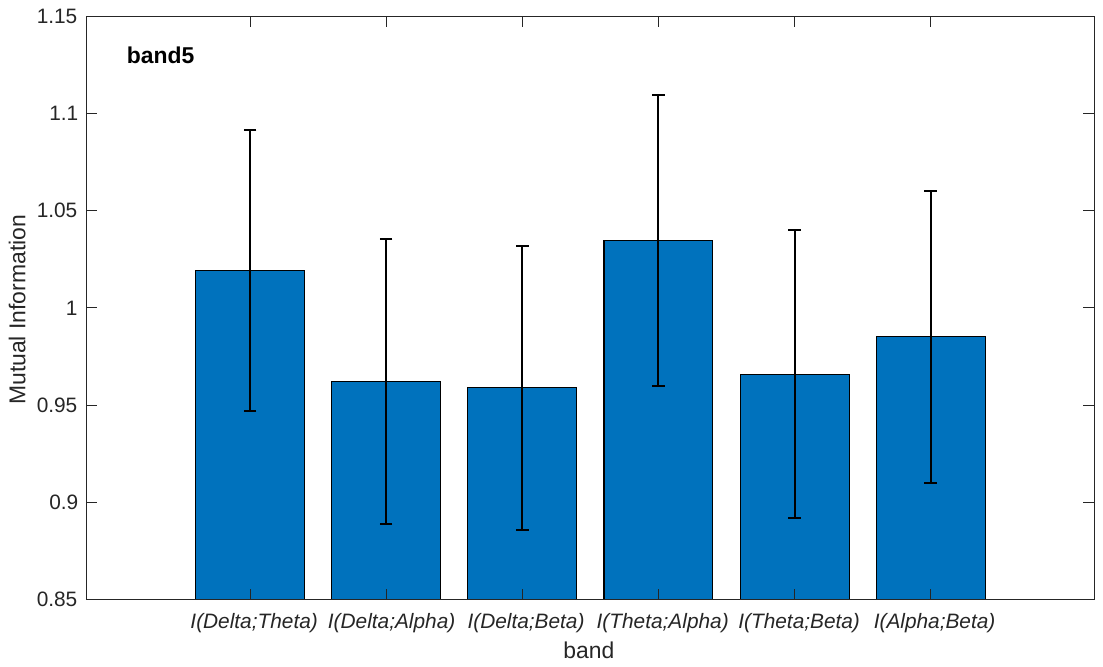

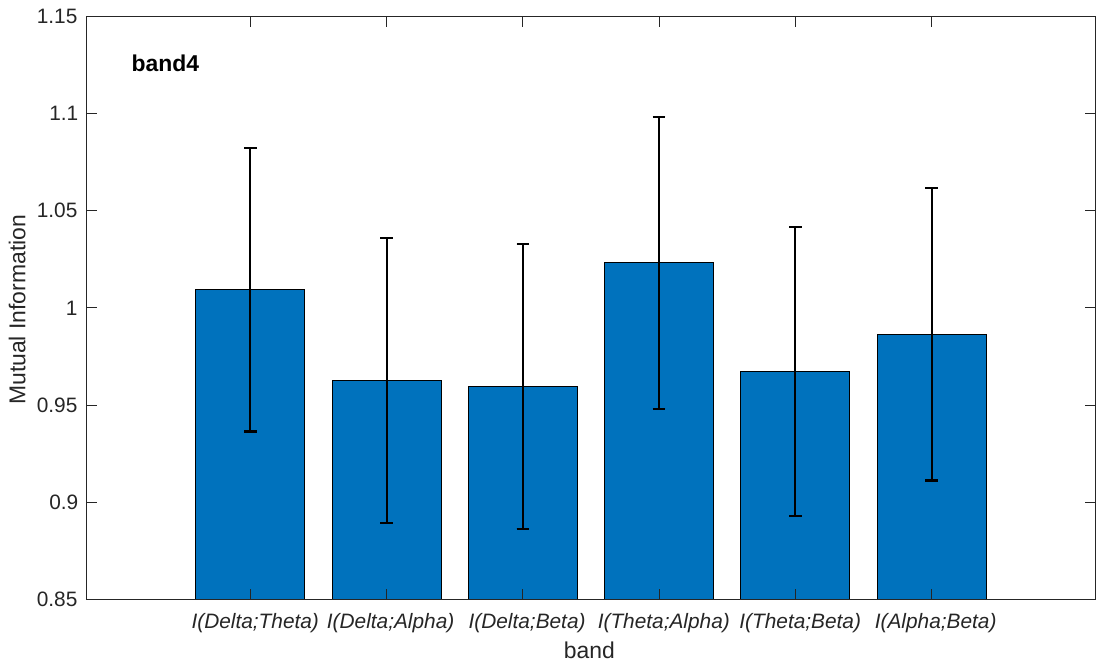

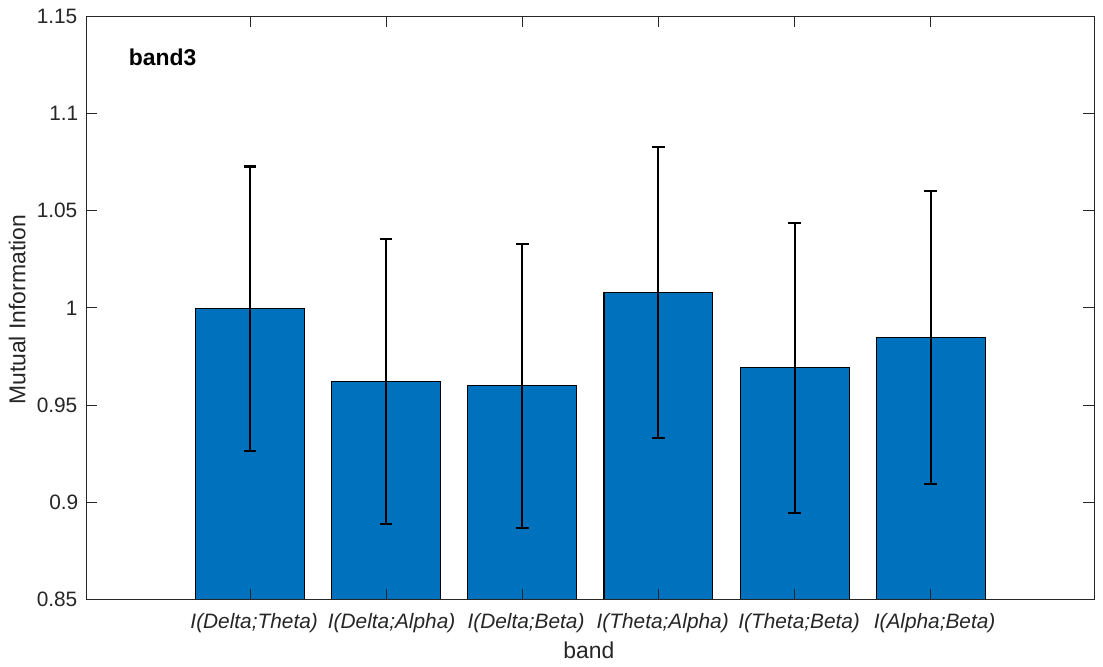

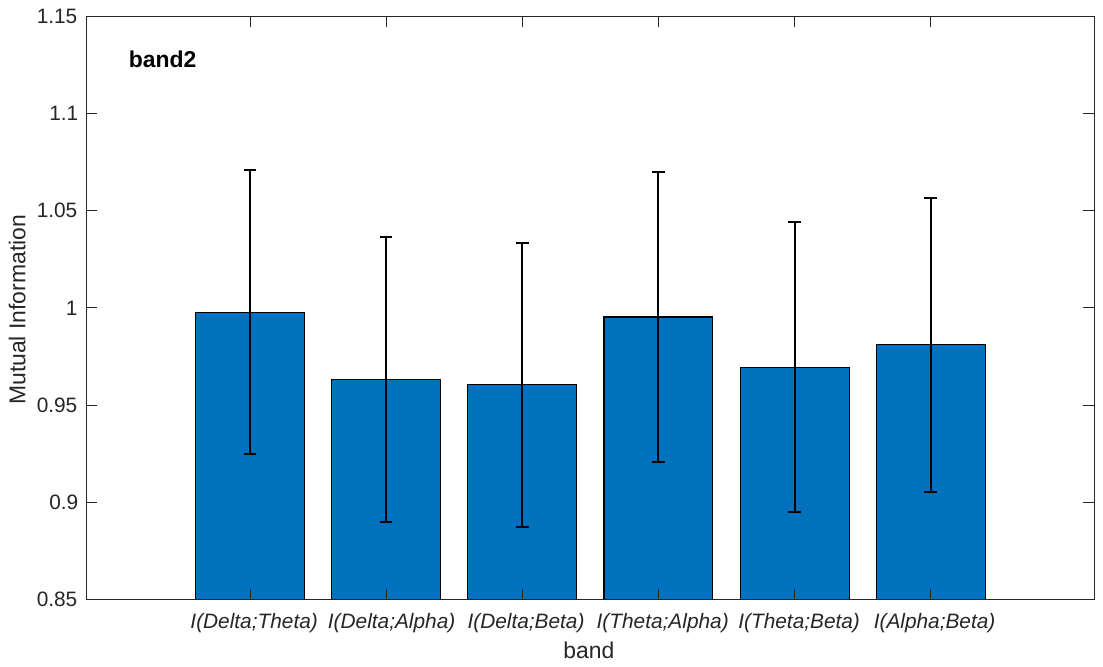

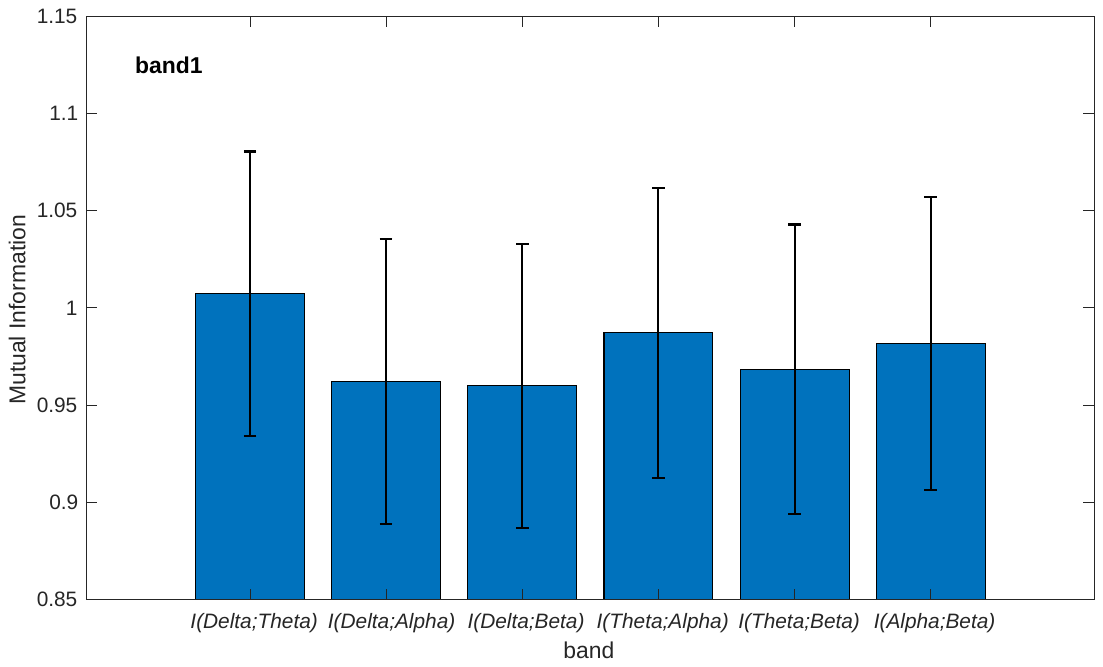

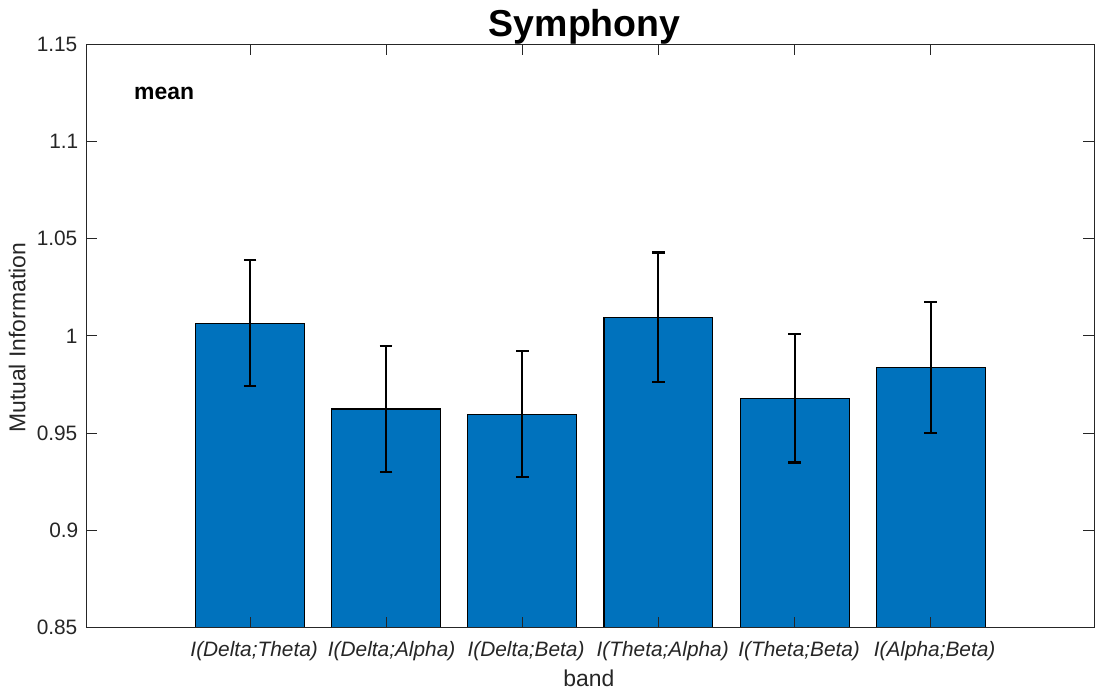

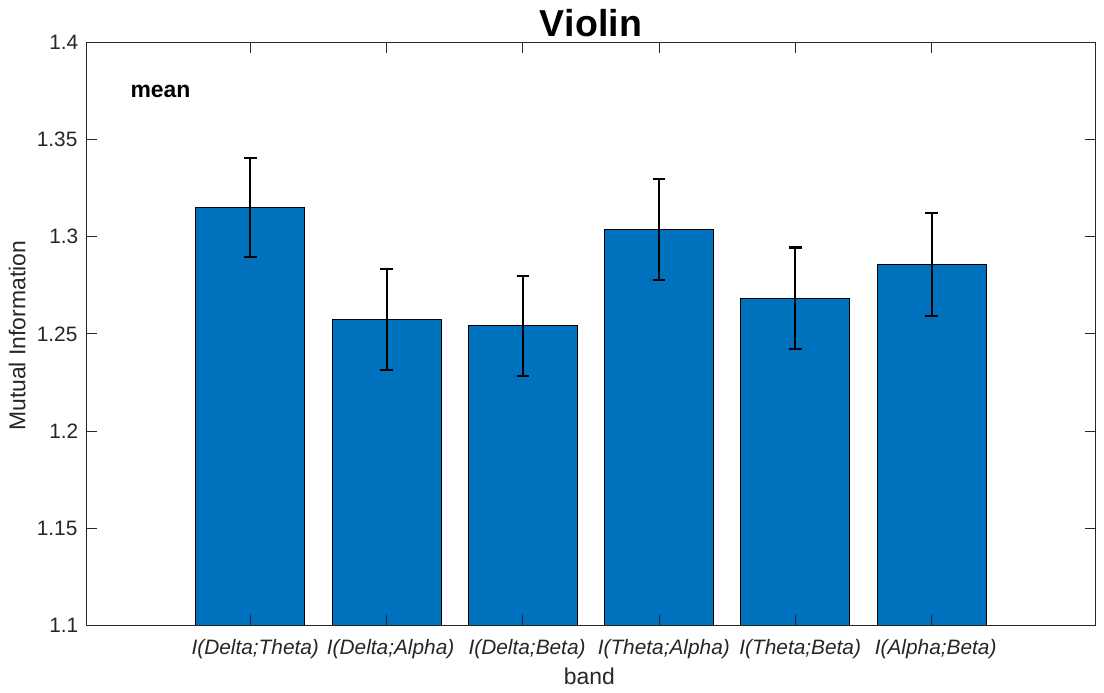

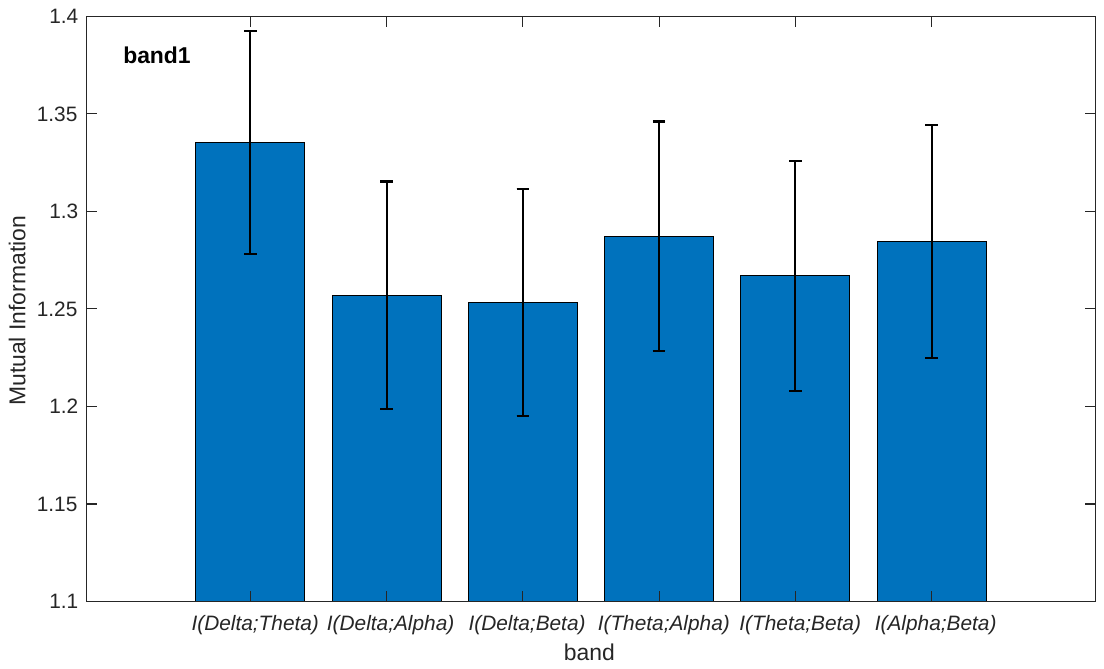

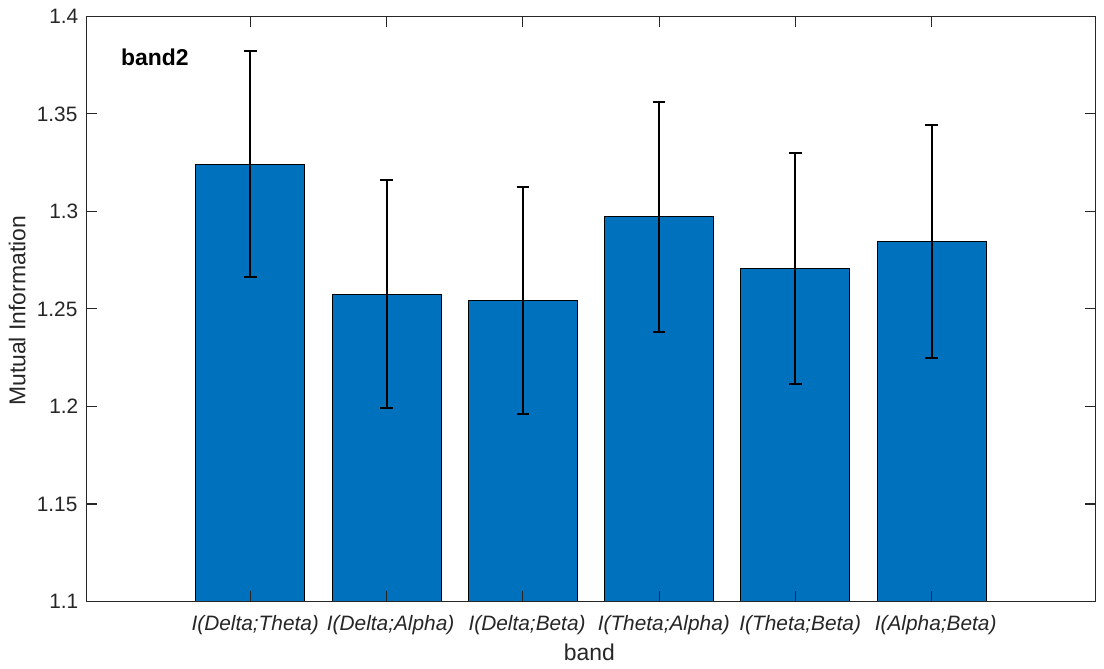

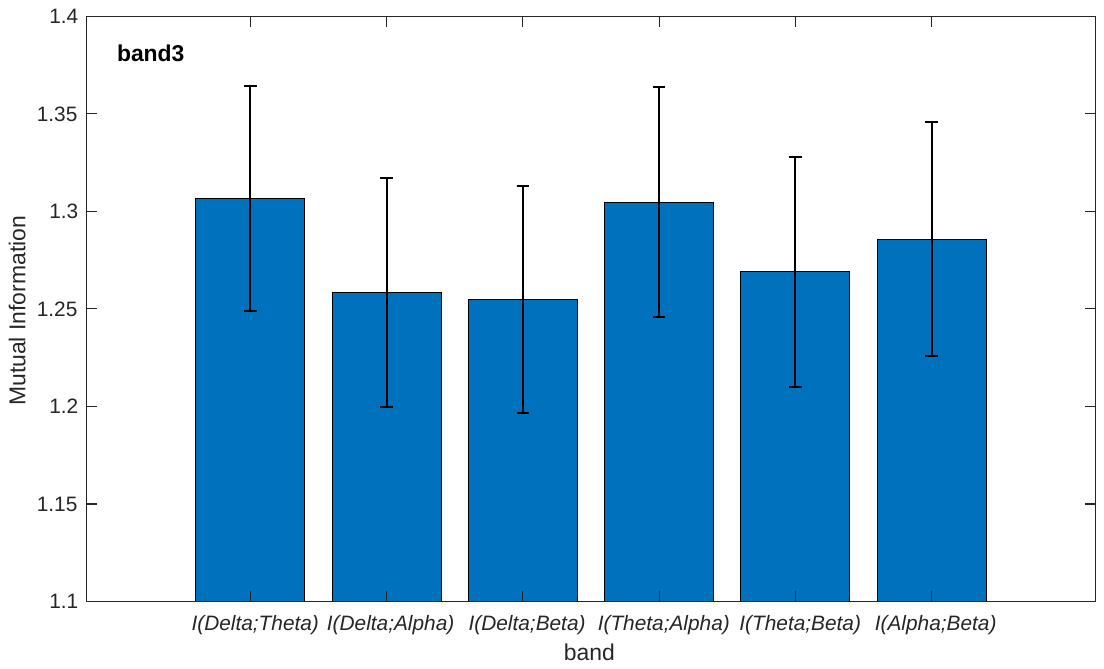

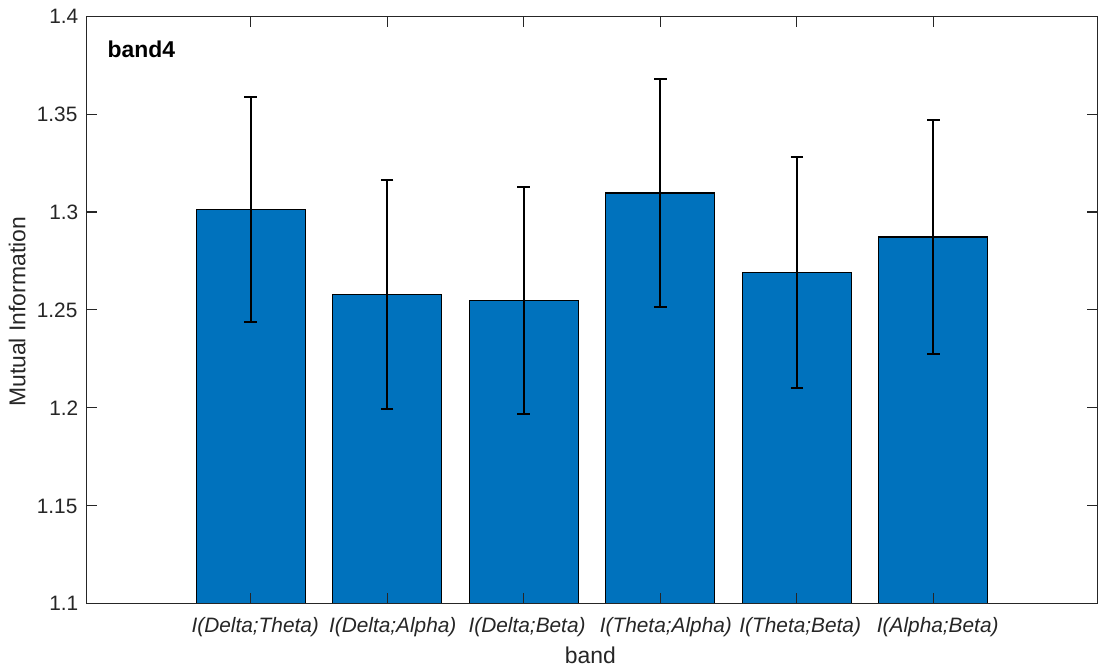

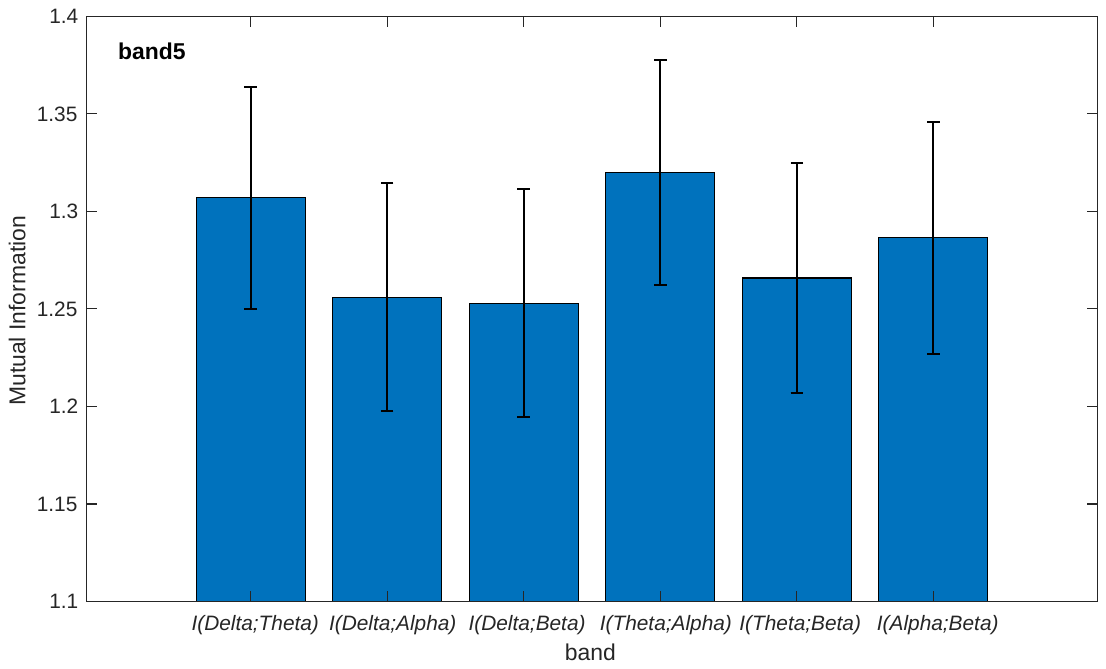

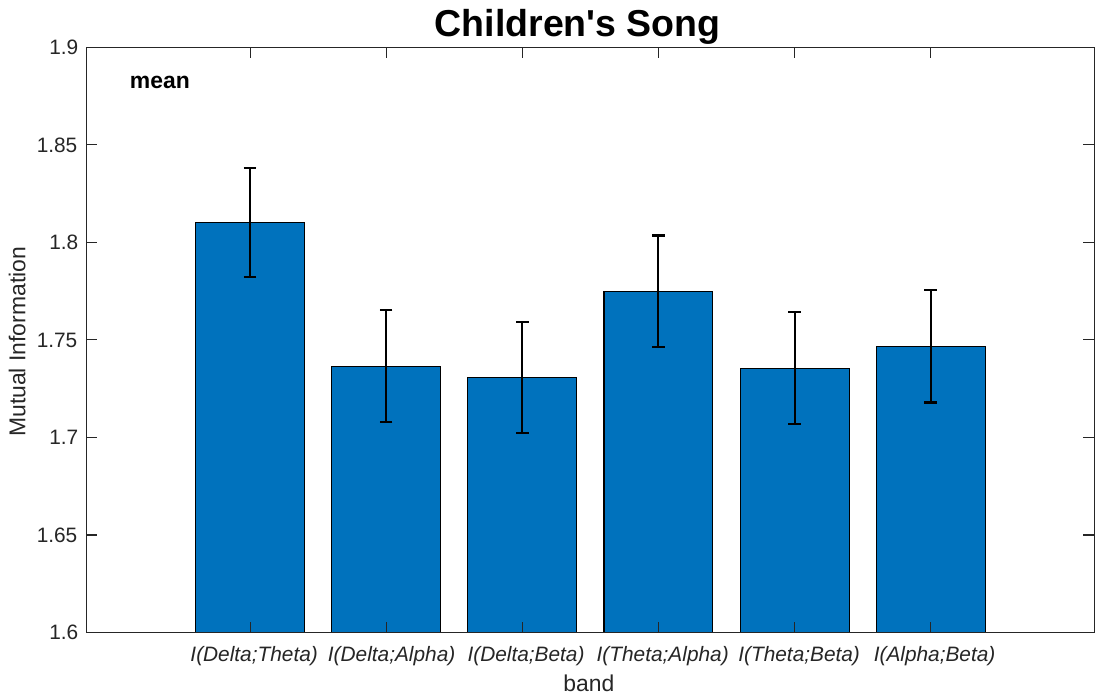

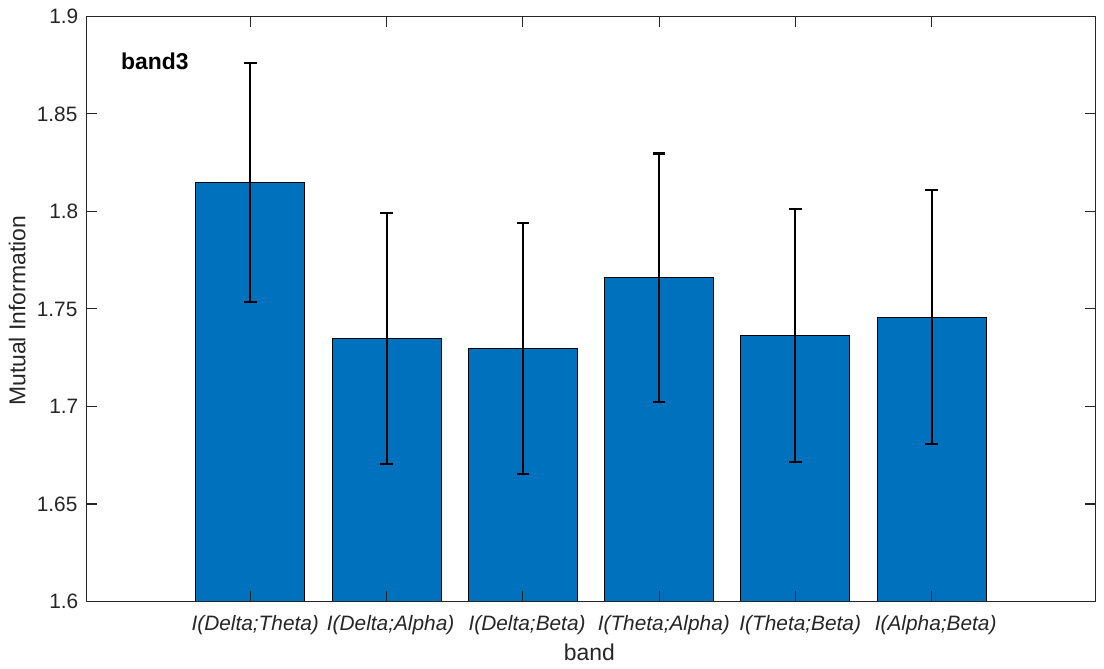

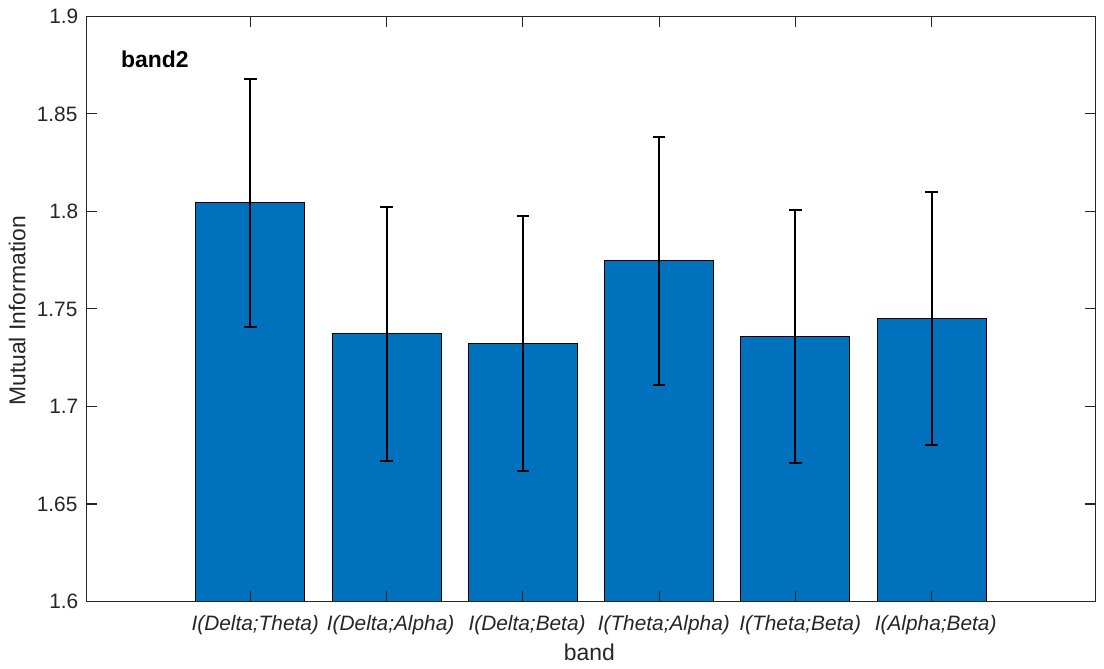

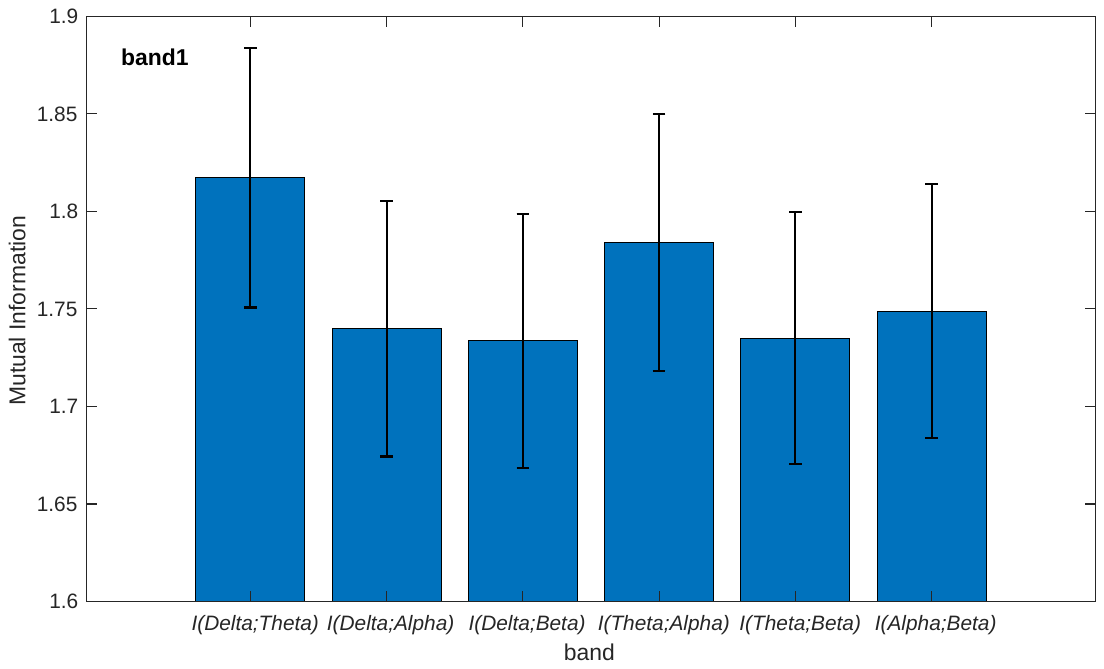

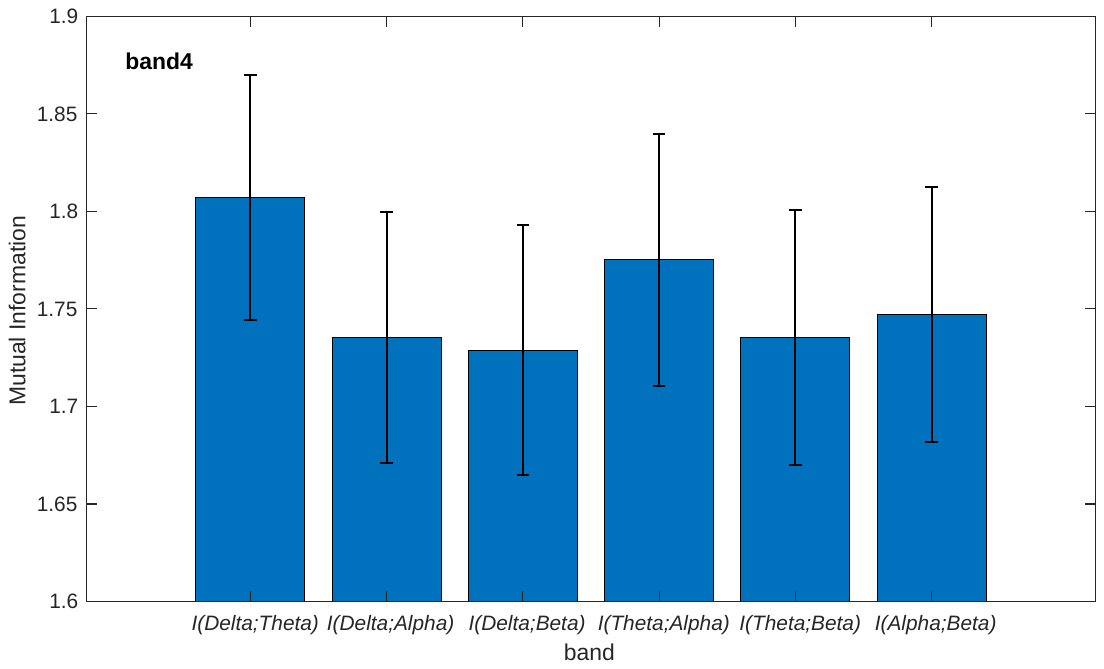

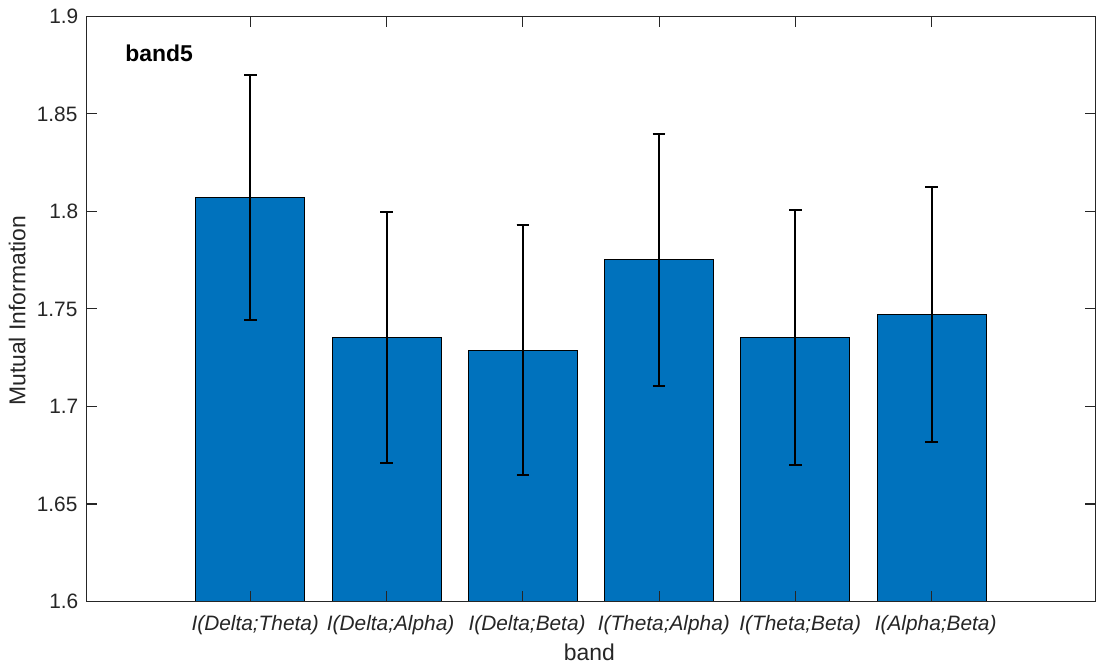

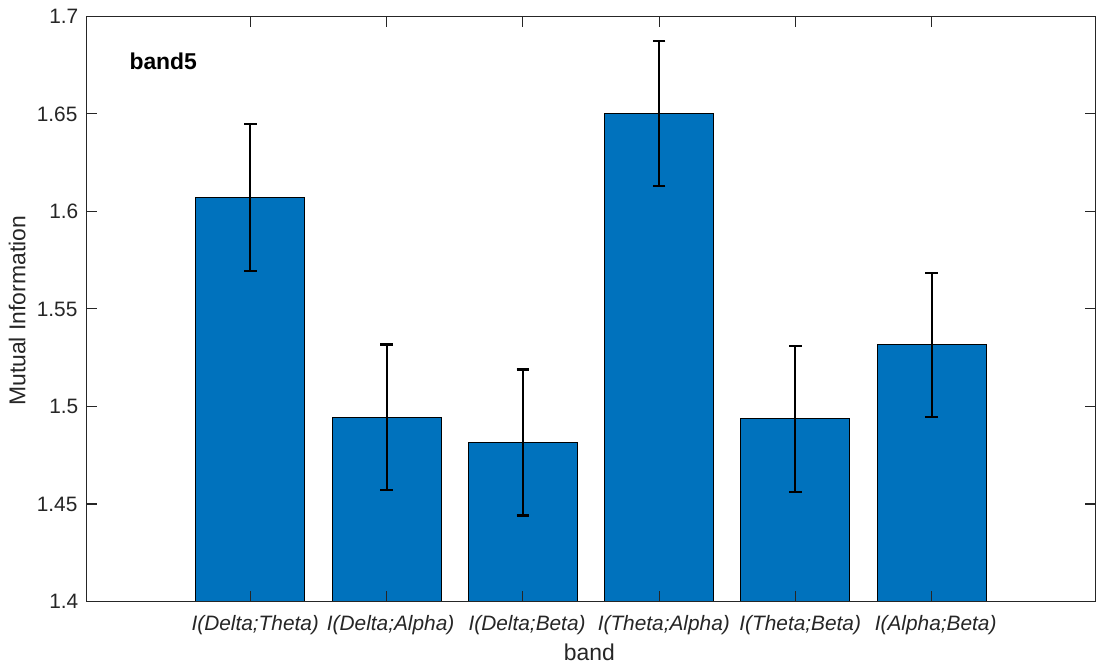

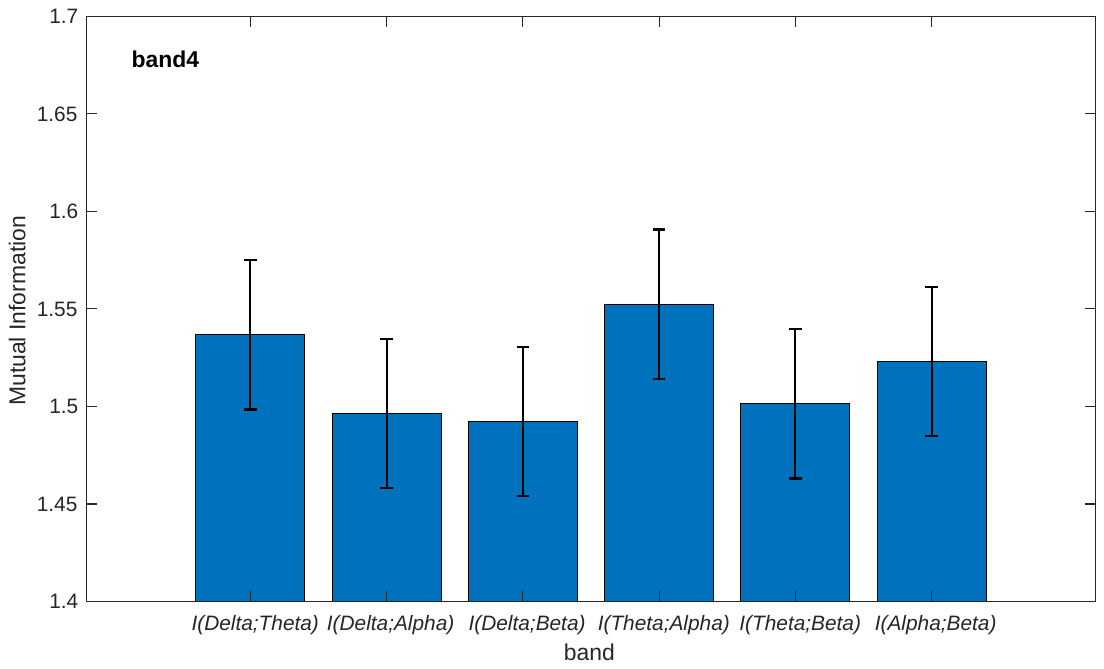

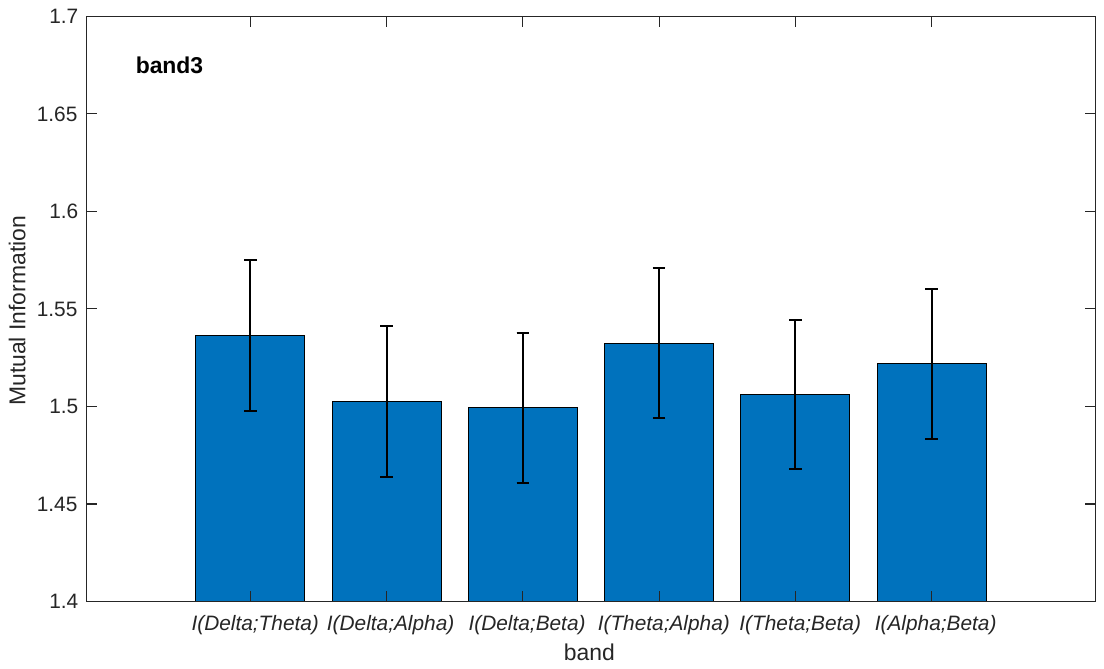

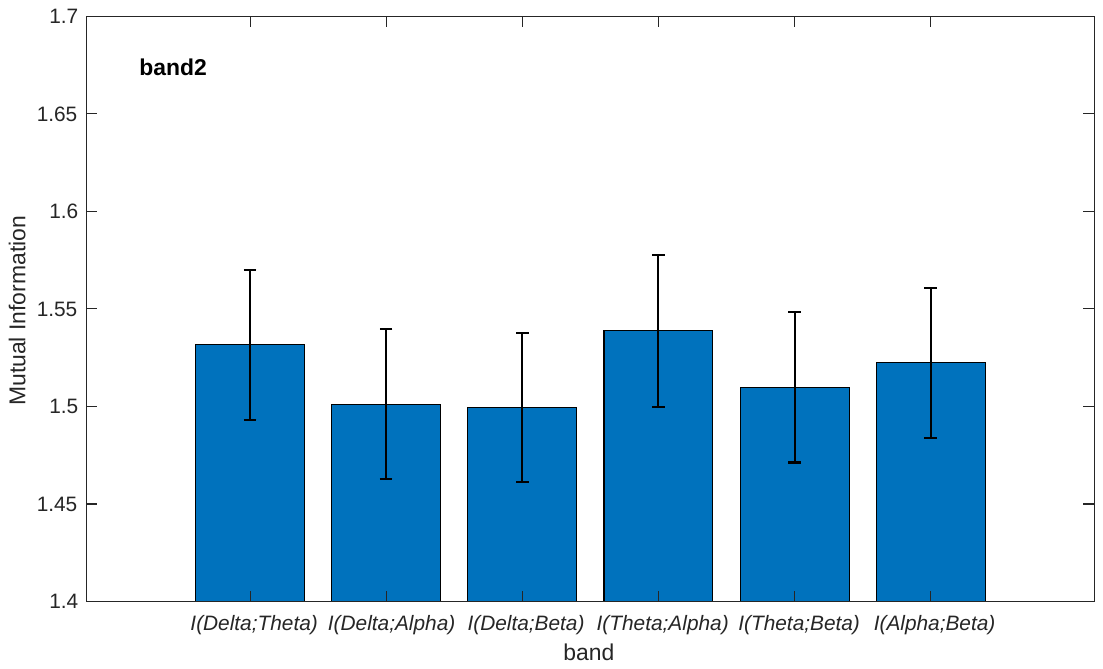

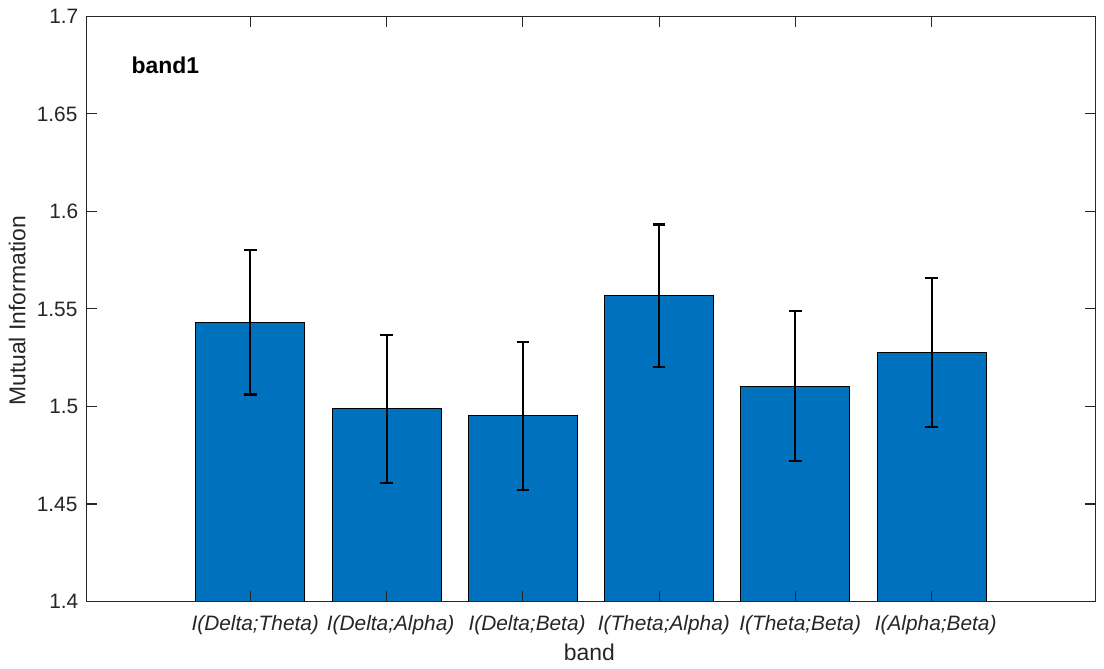

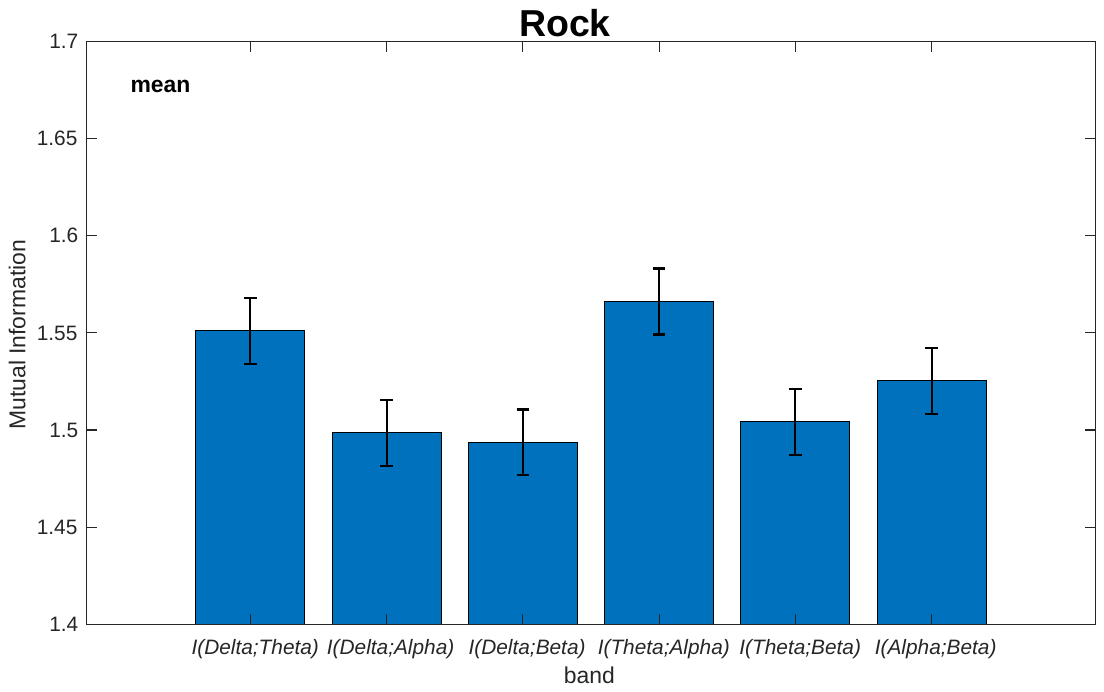

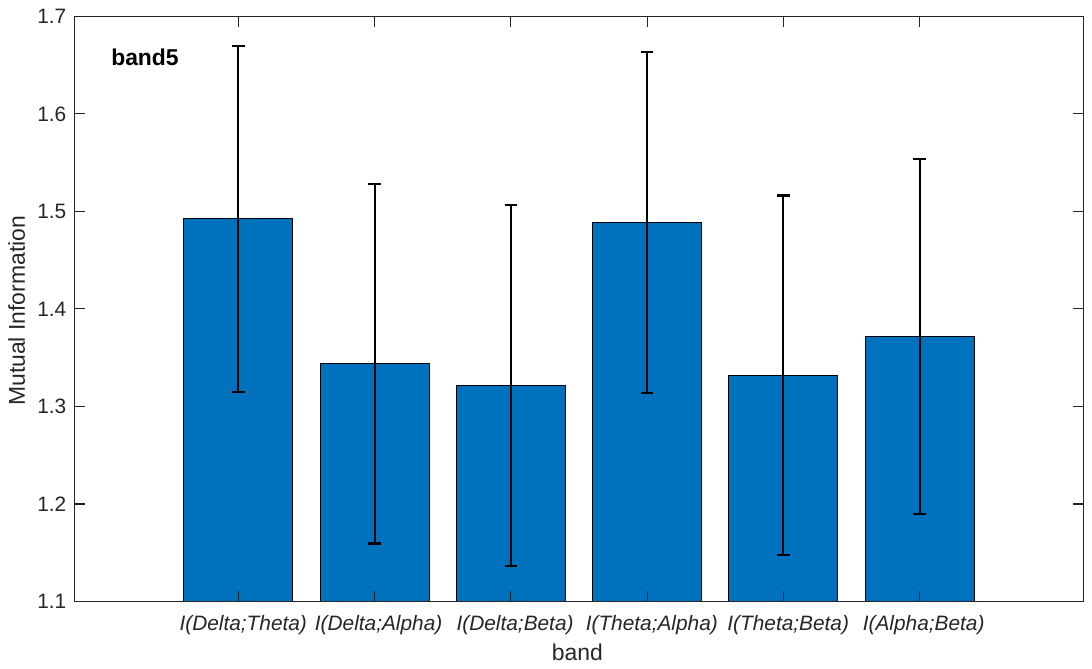

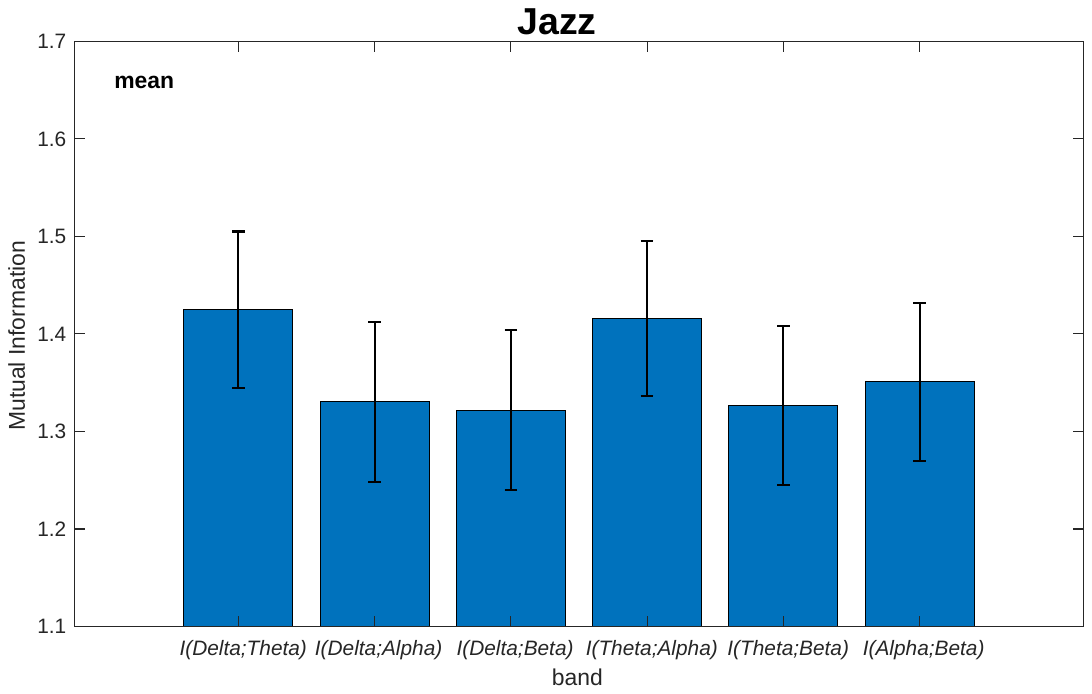

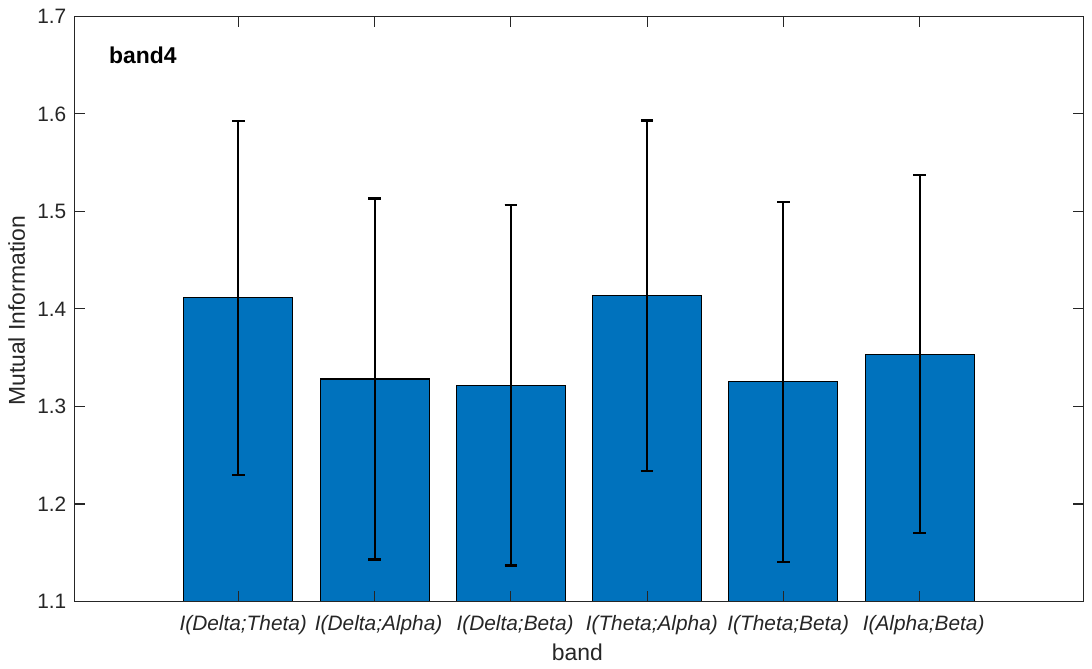

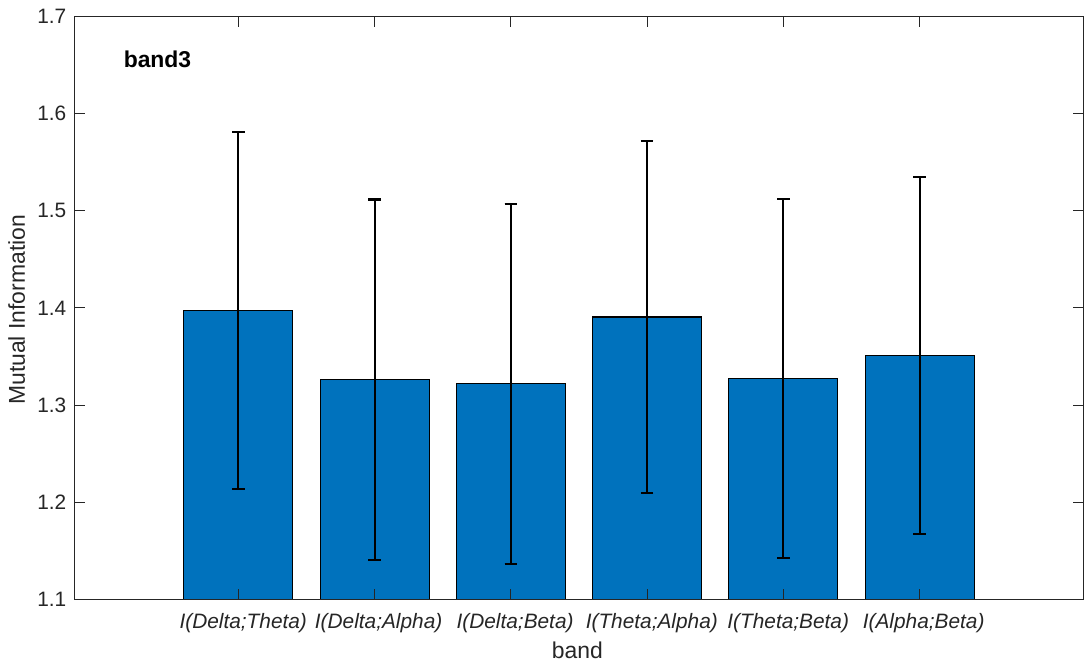

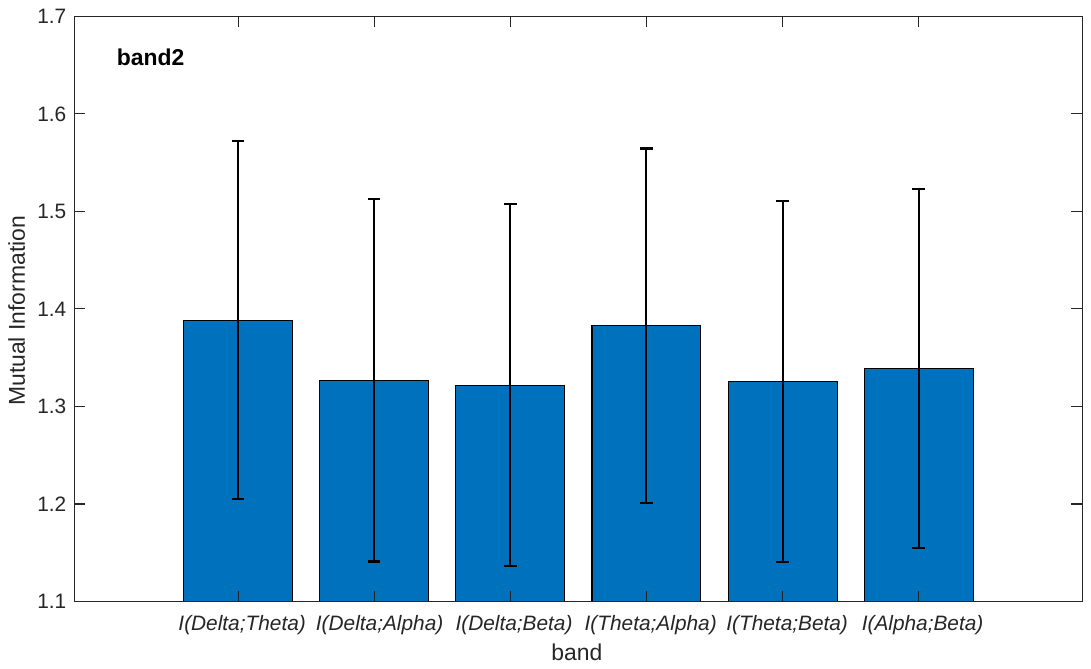

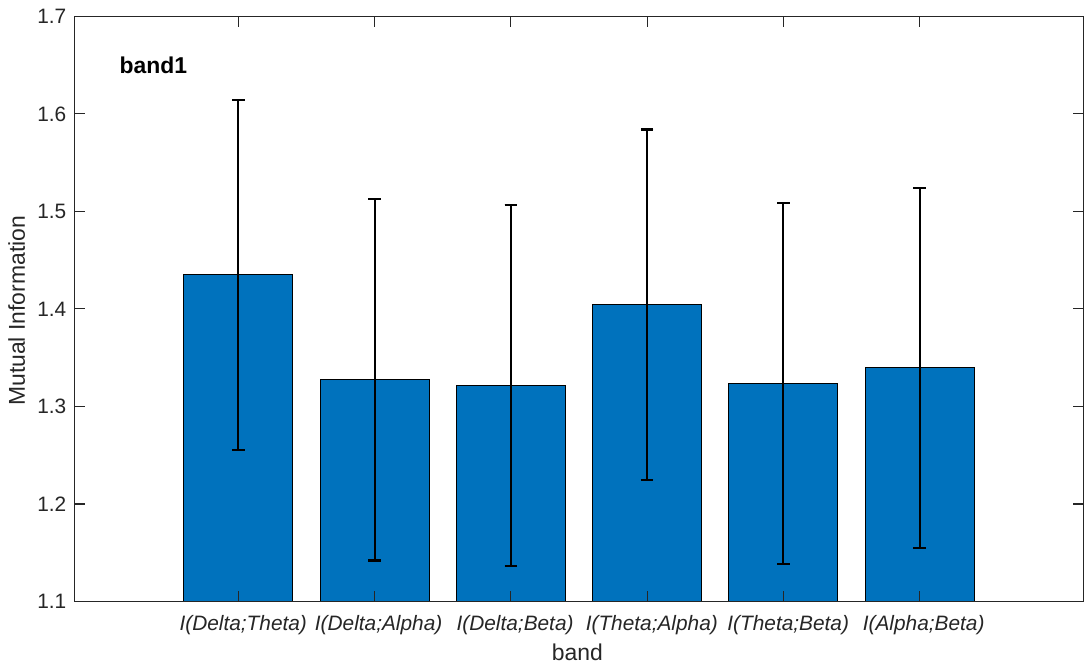

**Figure a**. Mutual information between different bands in S-AMPH model

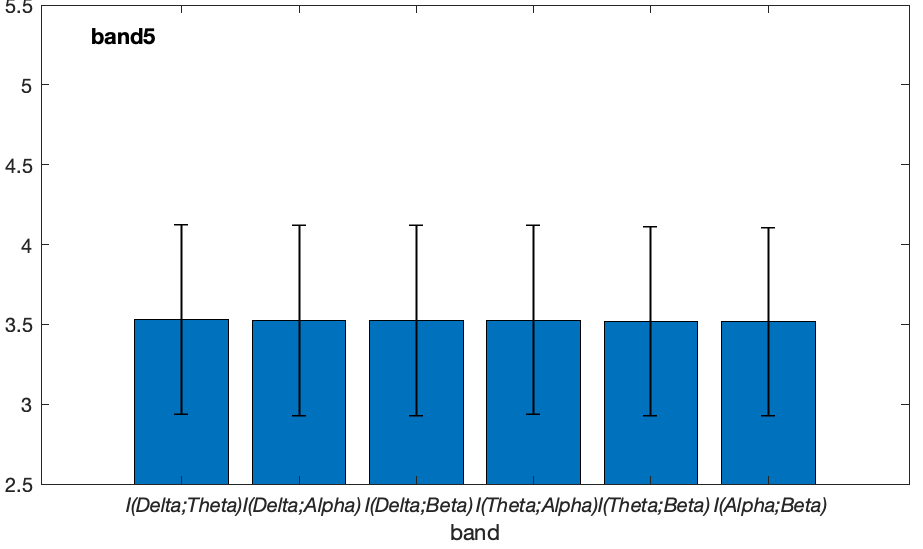

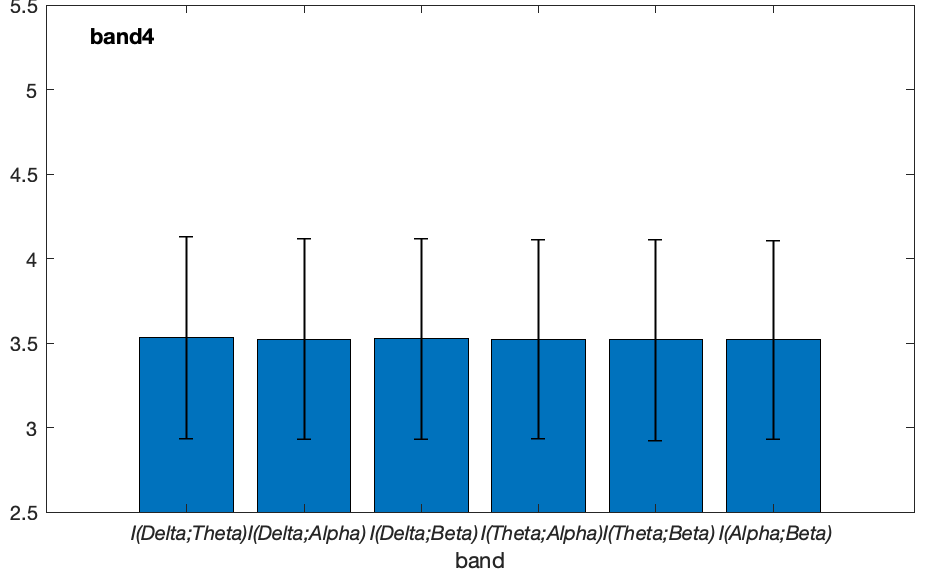

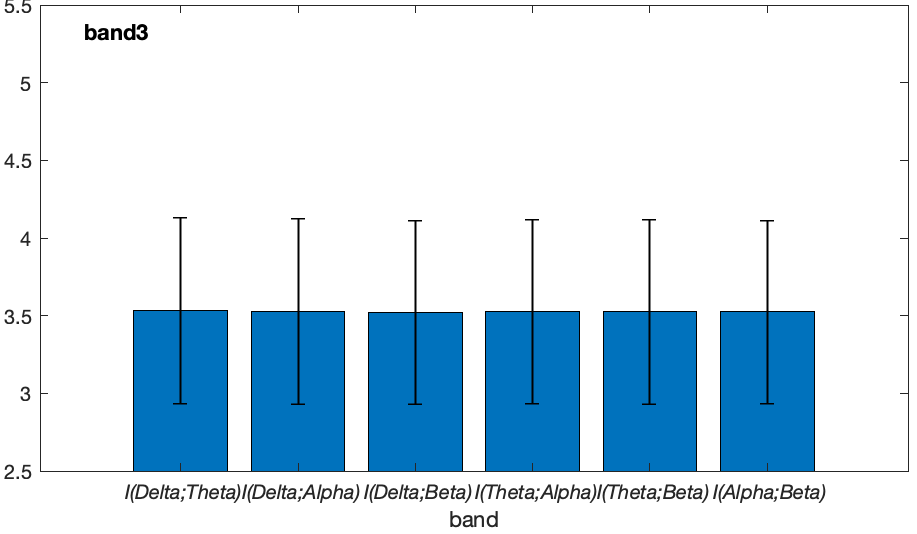

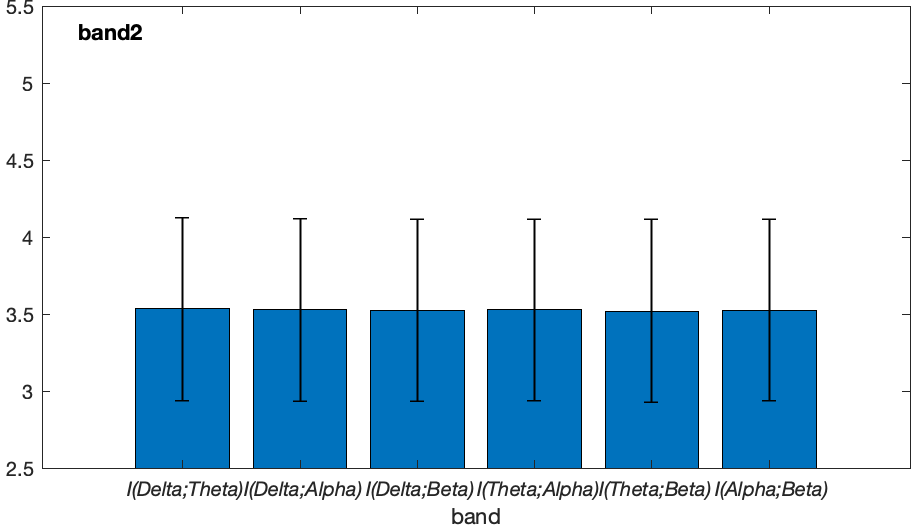

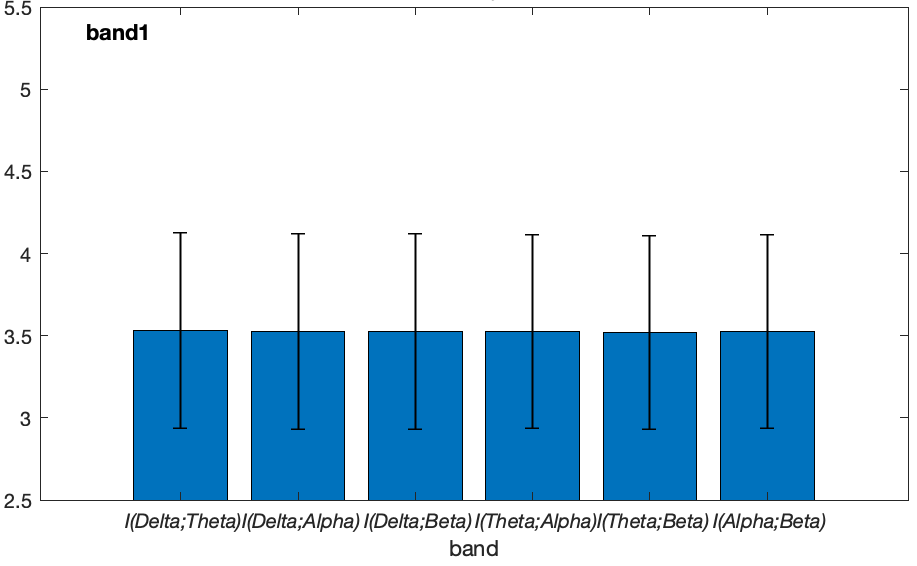

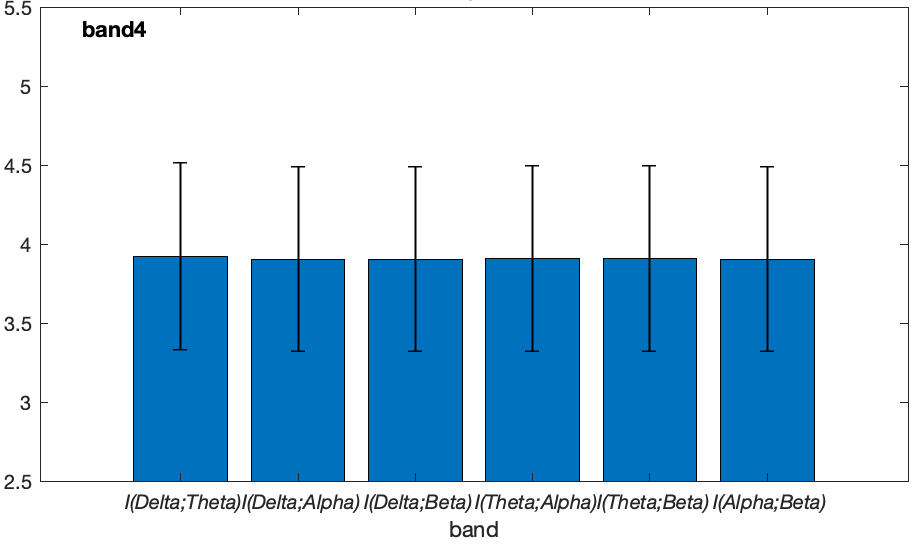

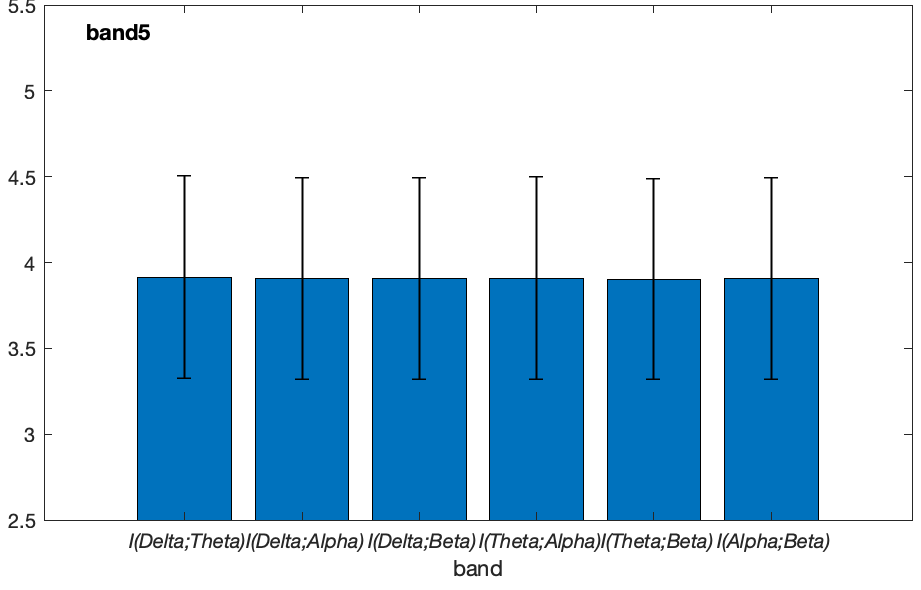

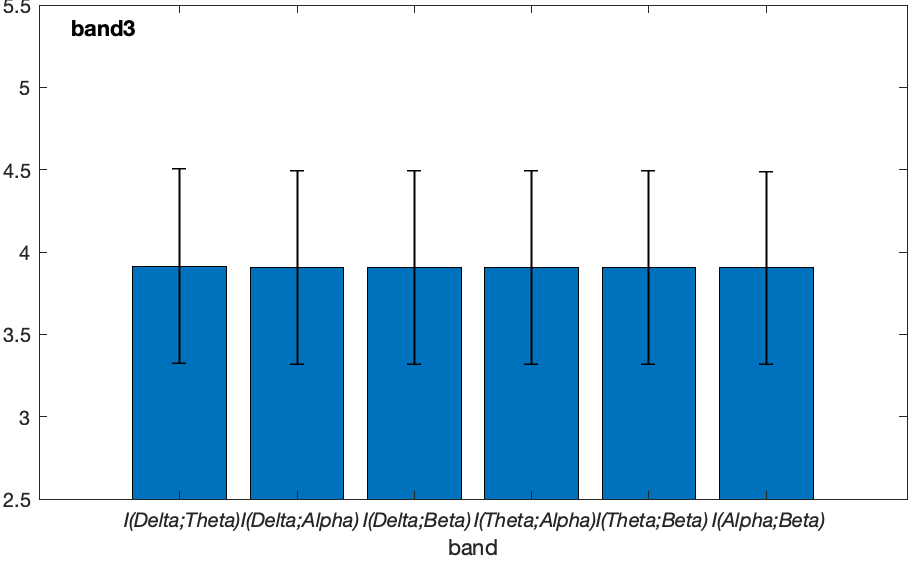

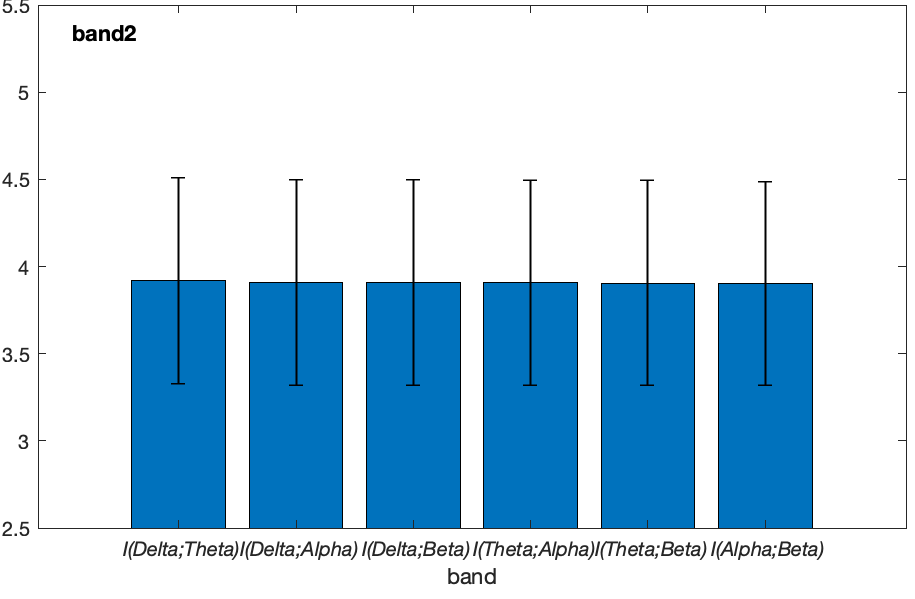

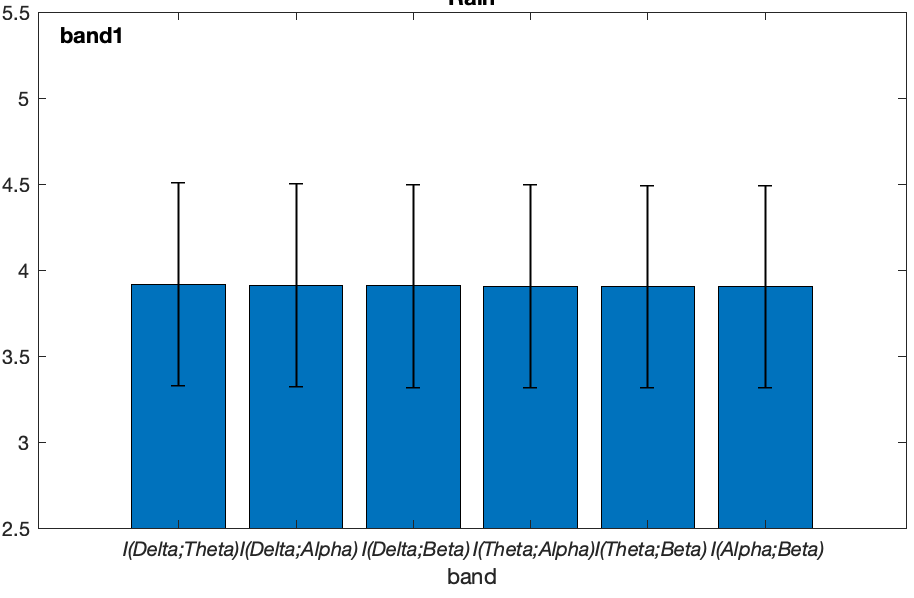

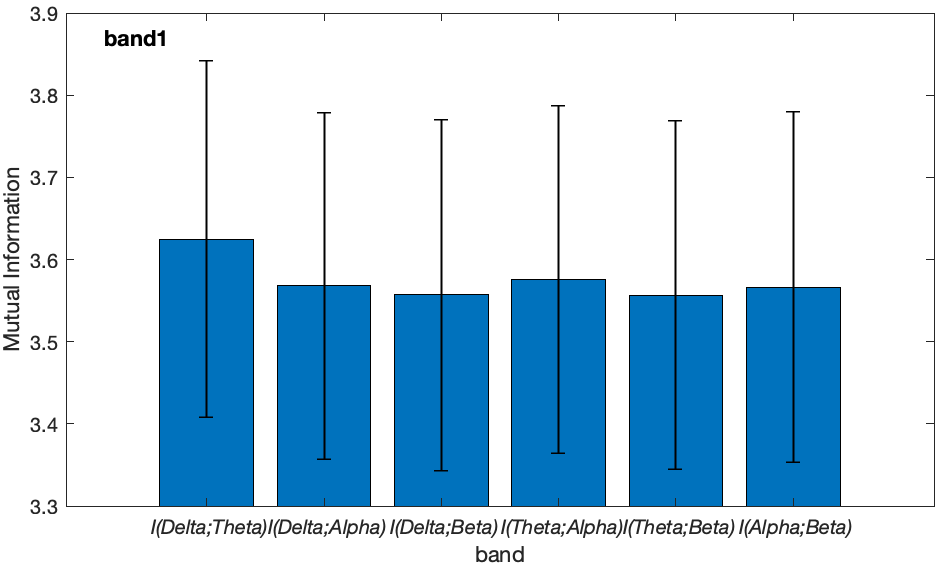

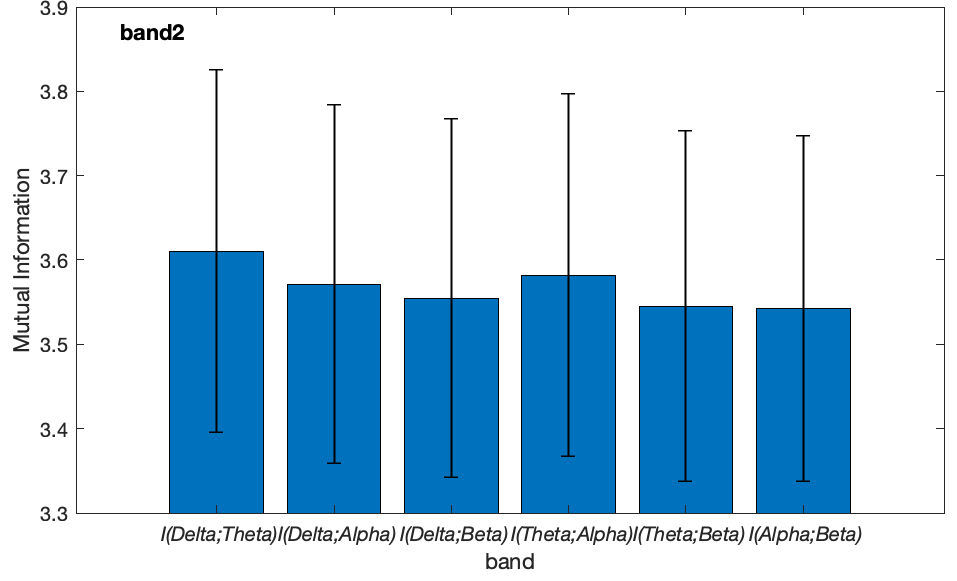

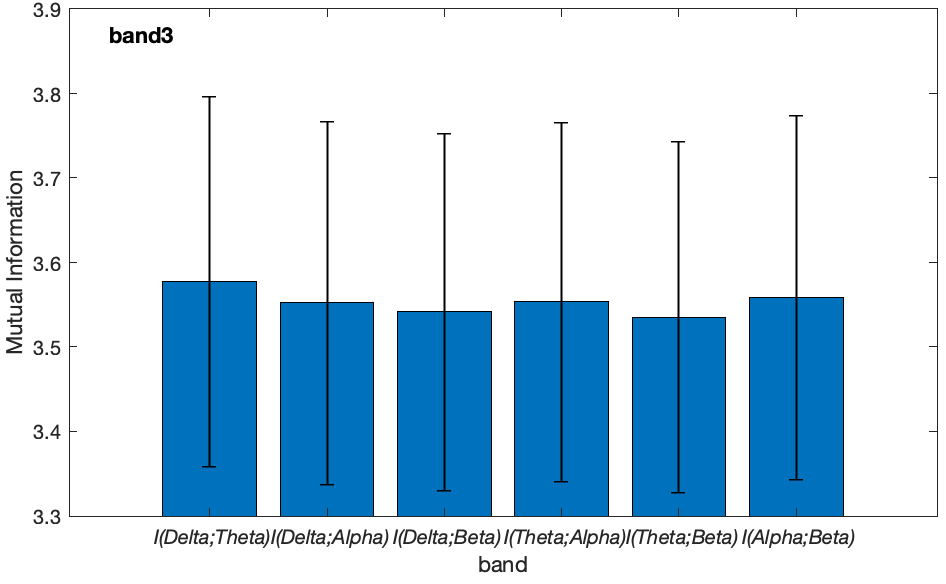

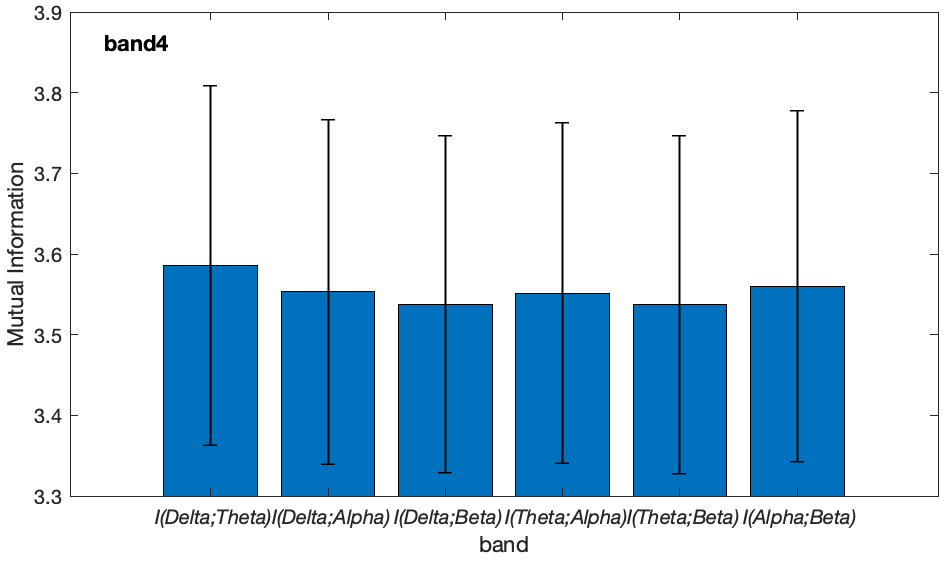

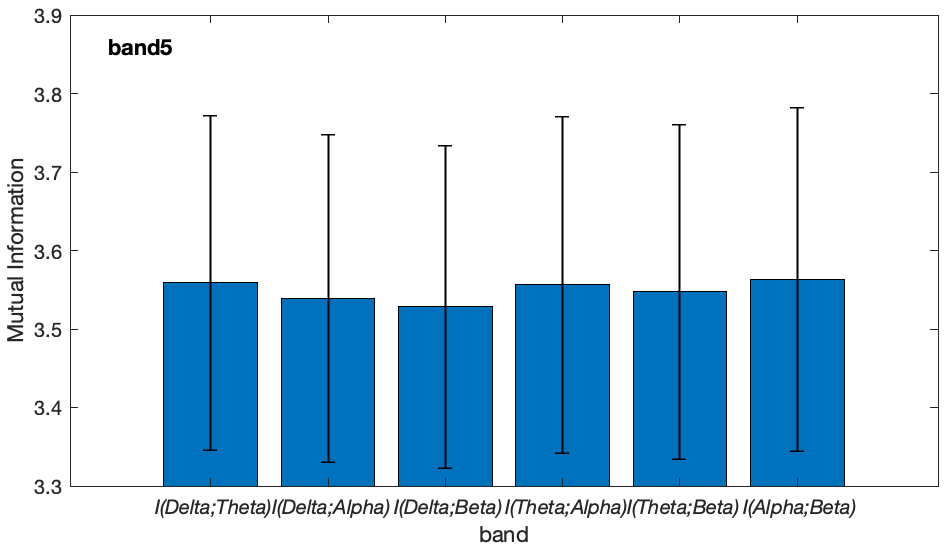

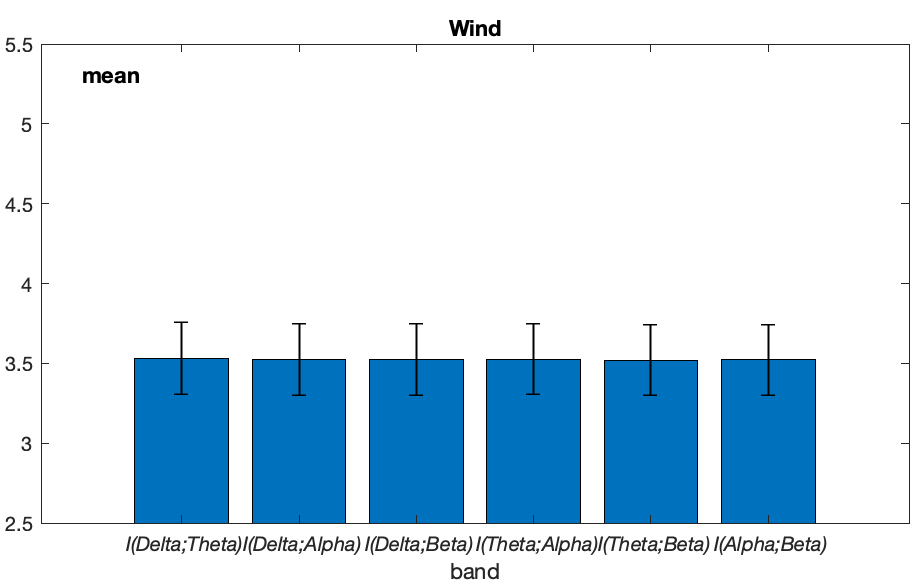

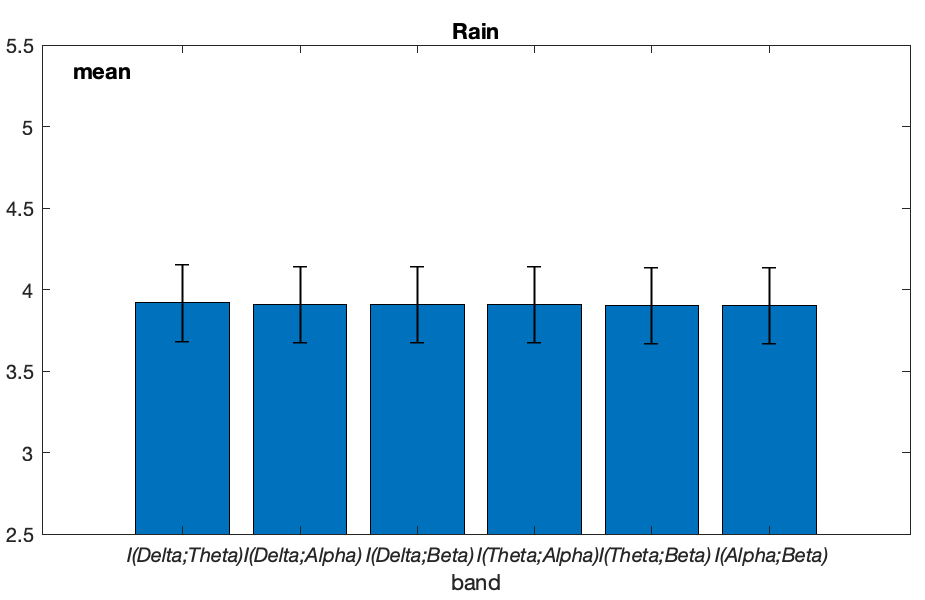

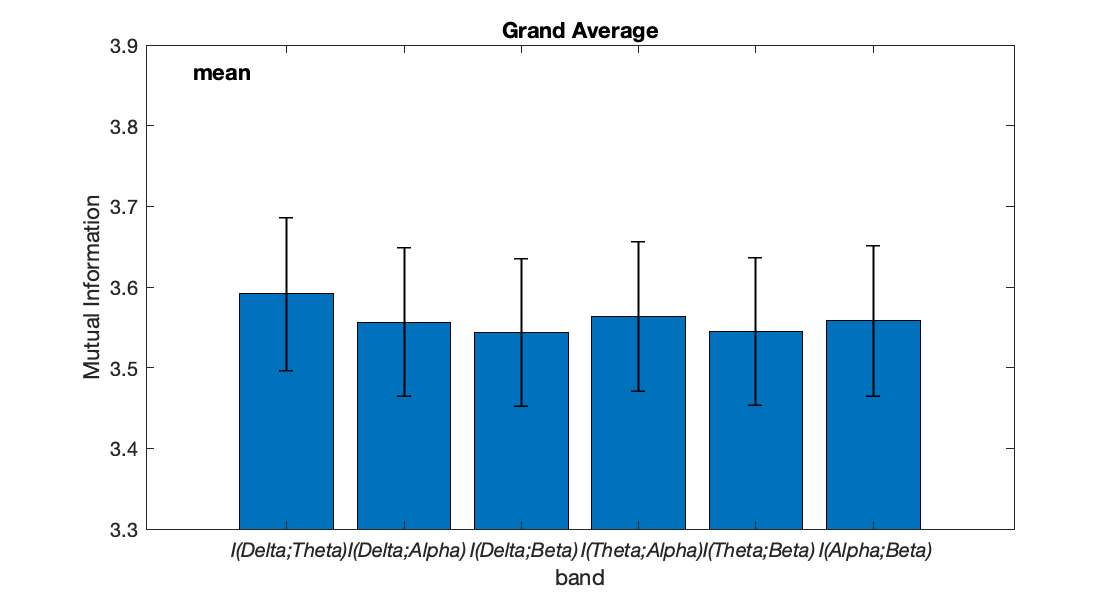

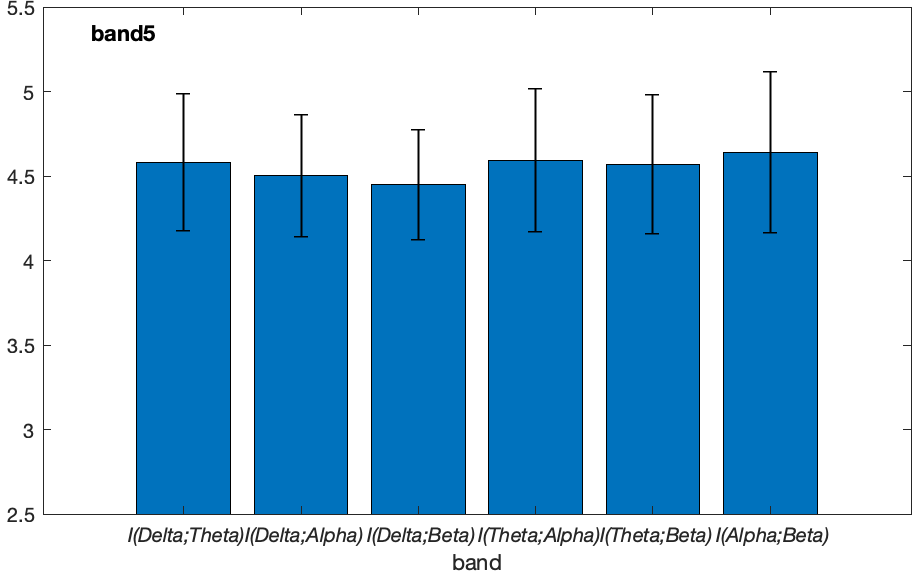

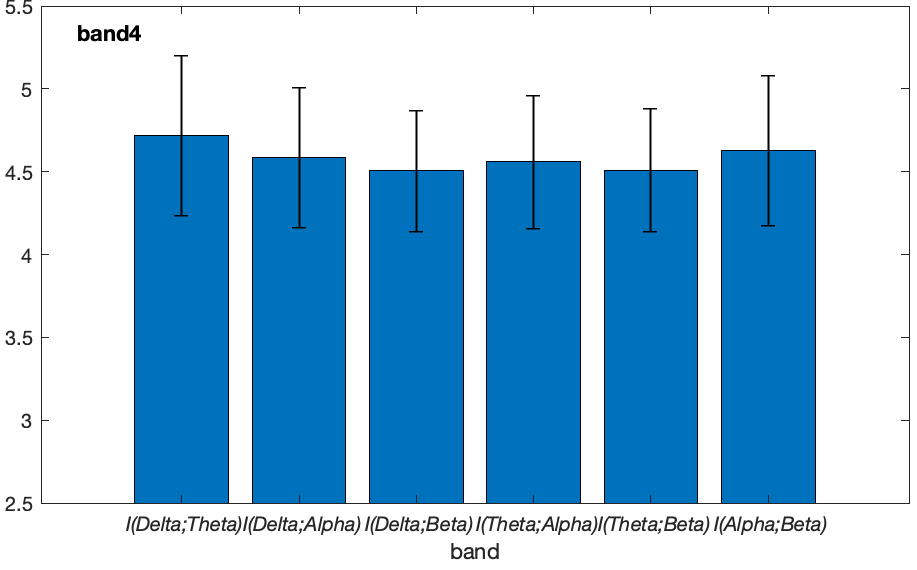

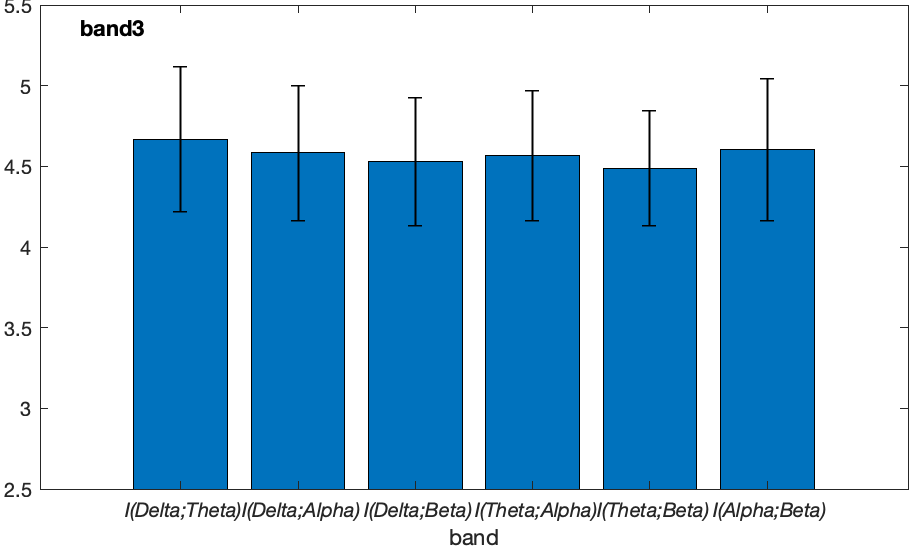

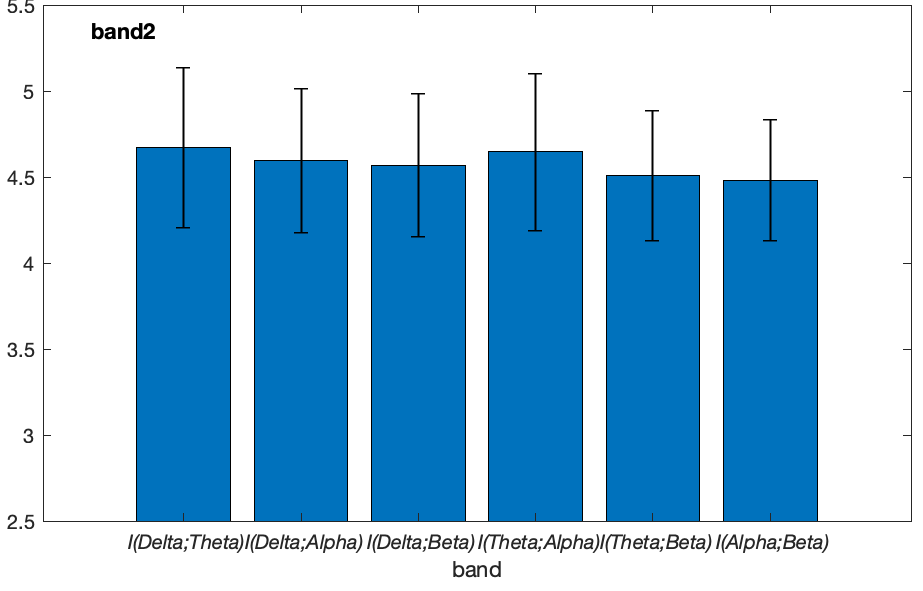

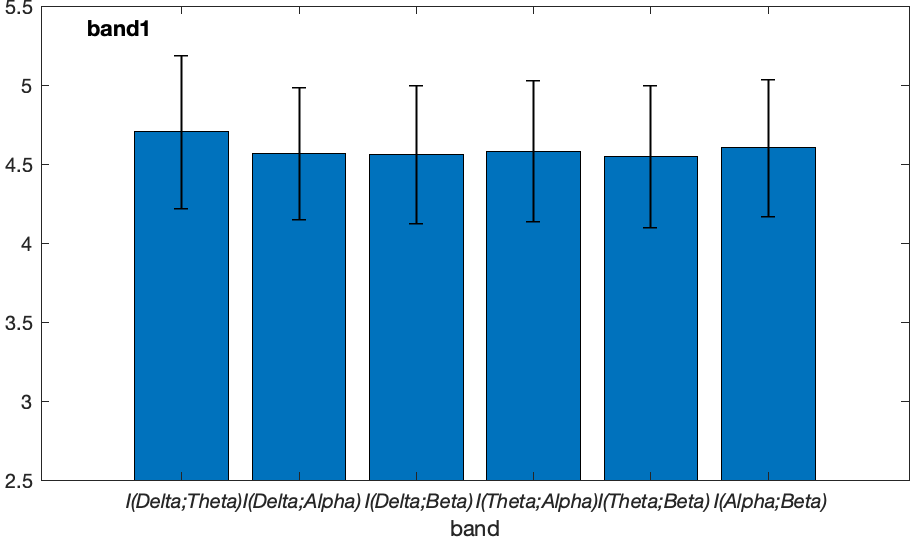

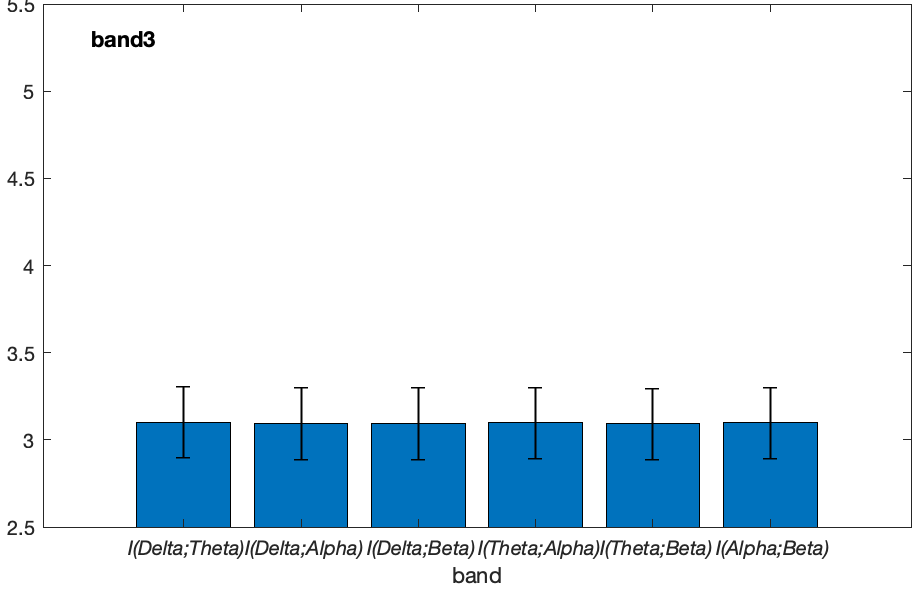

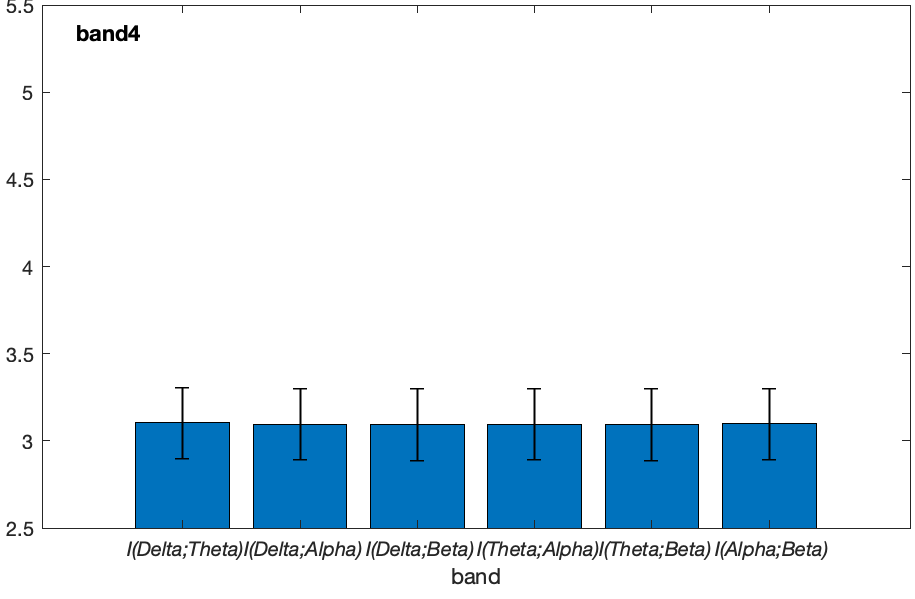

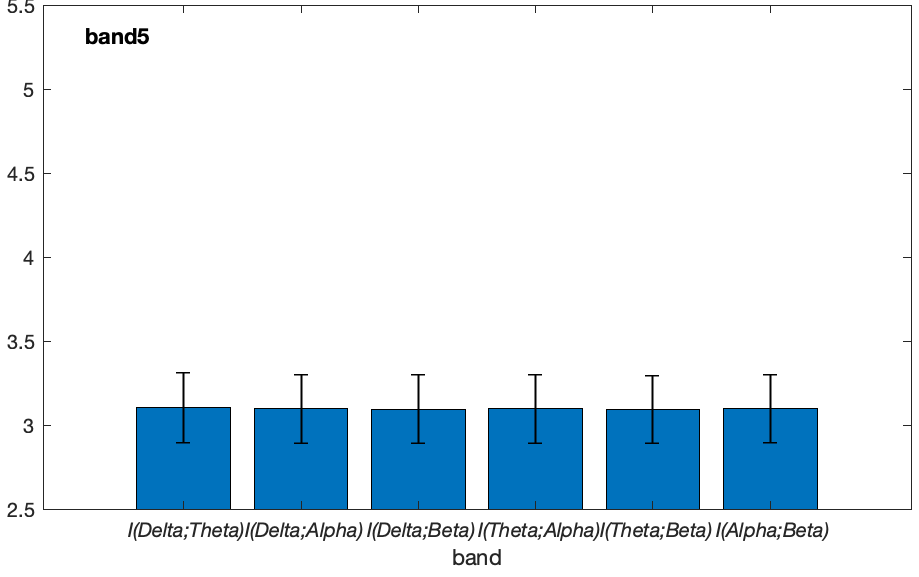

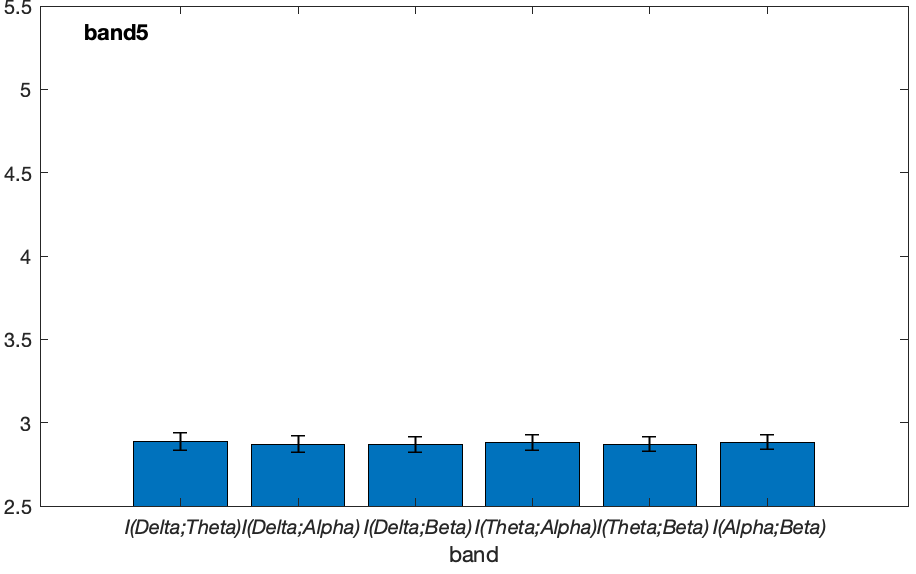

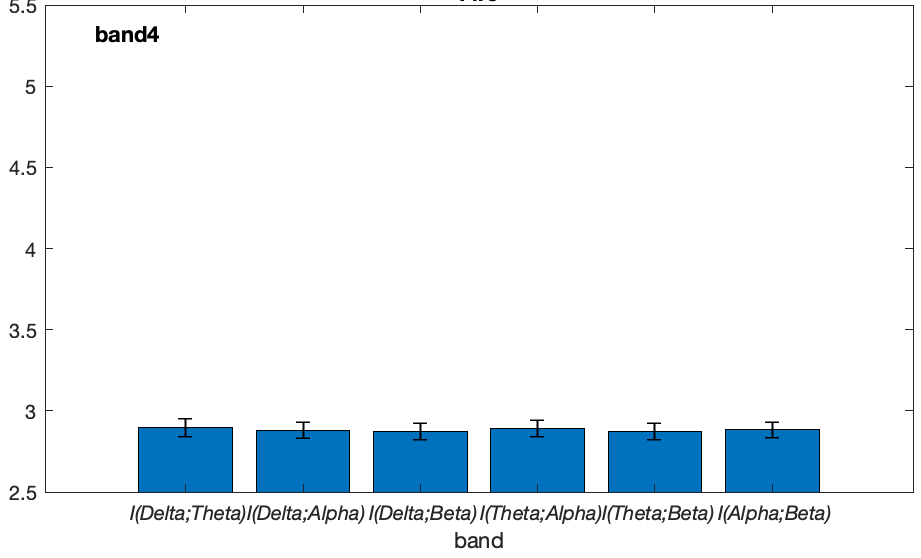

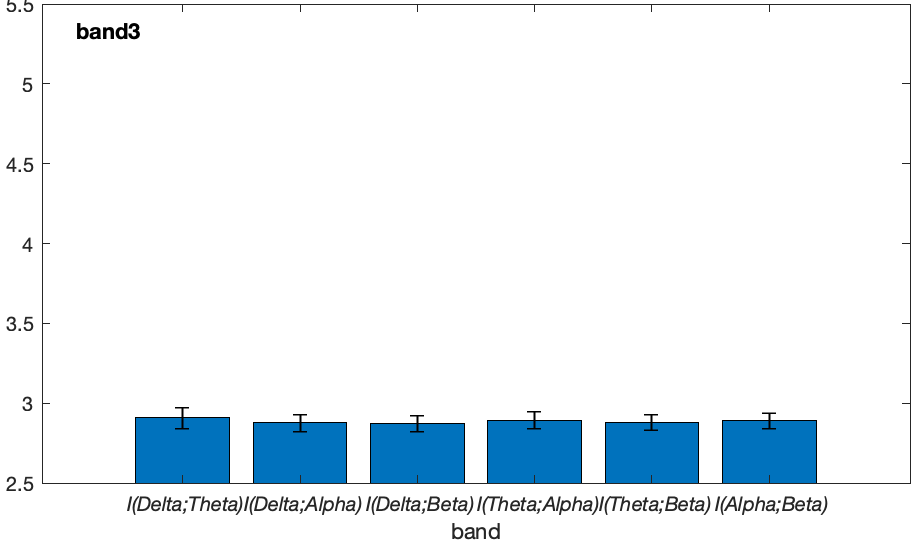

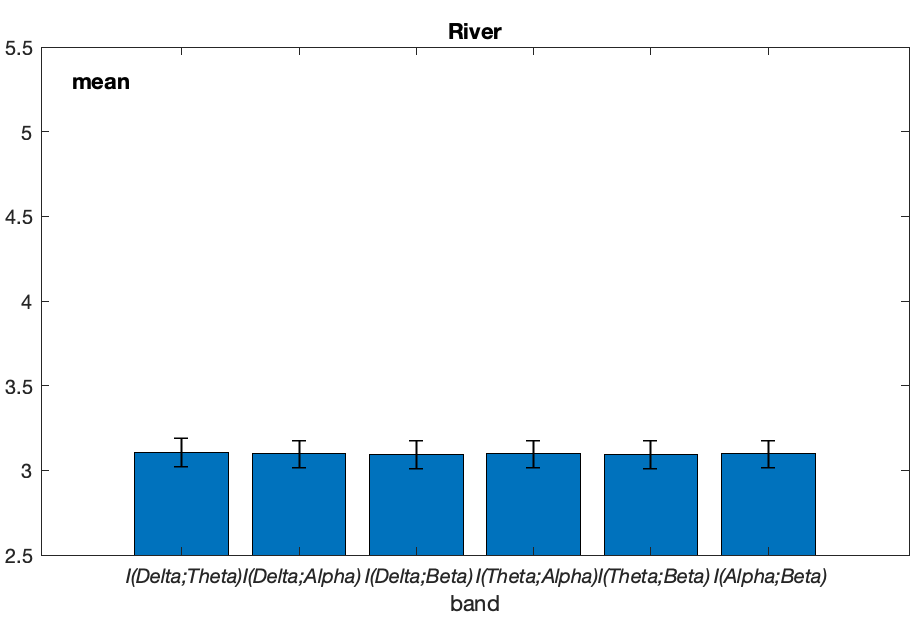

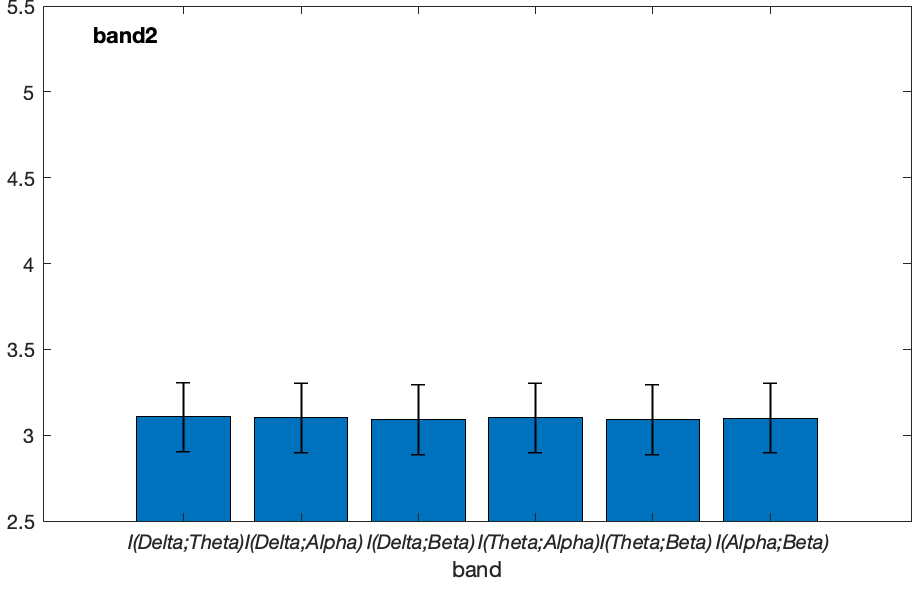

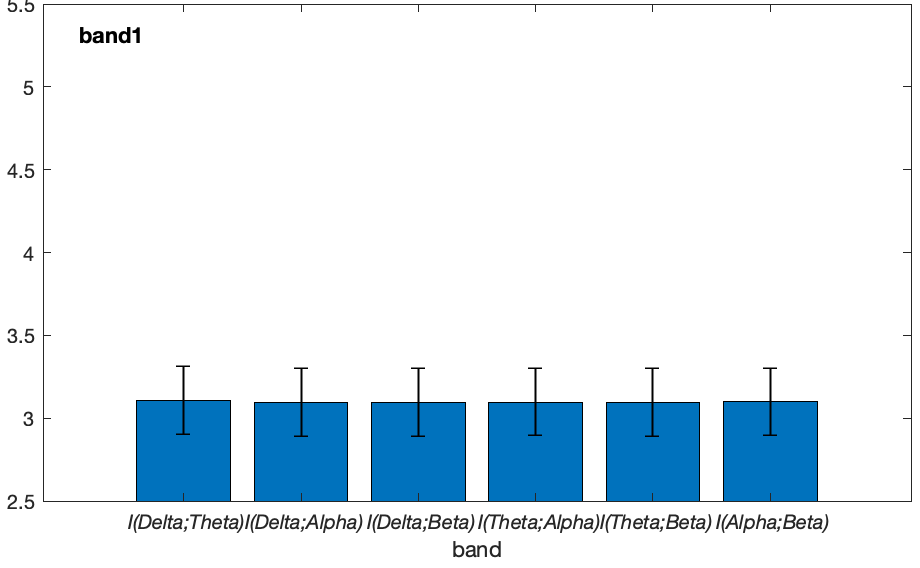

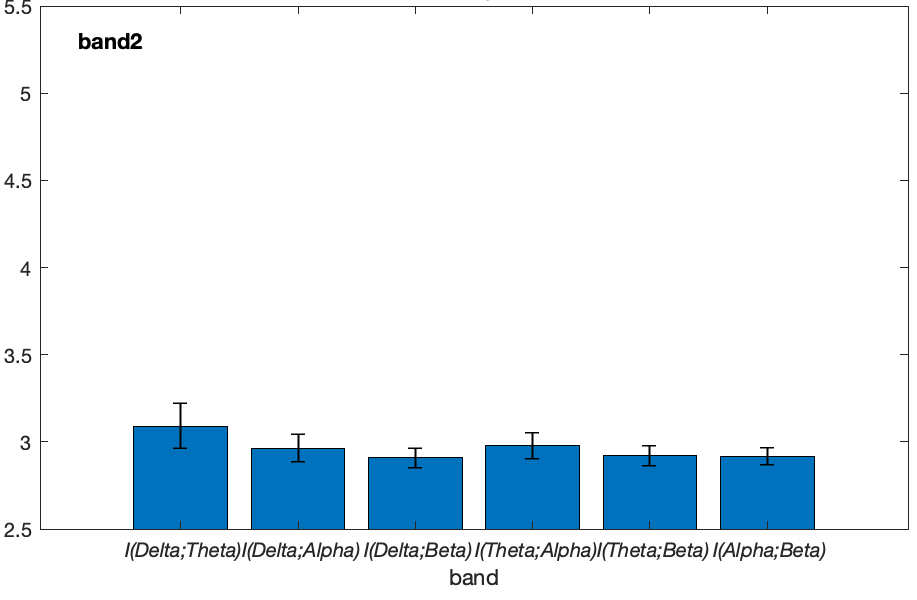

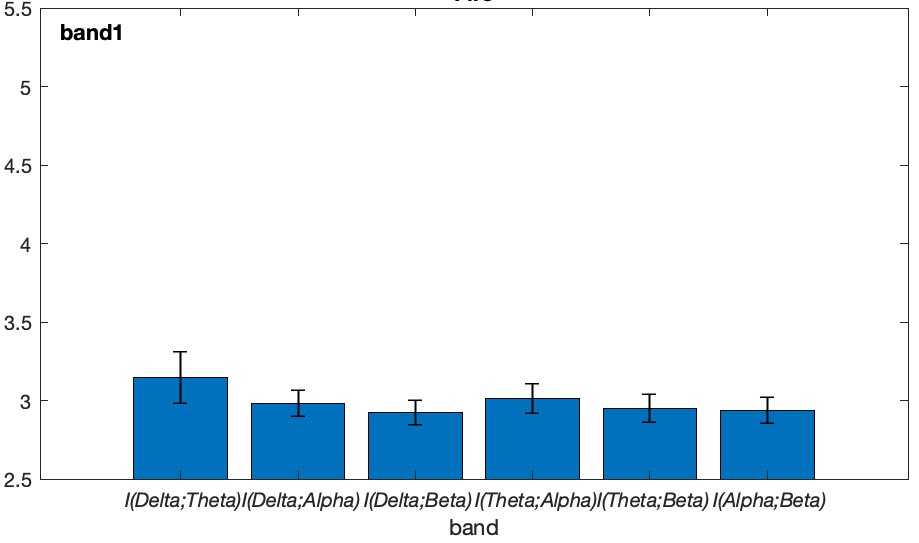

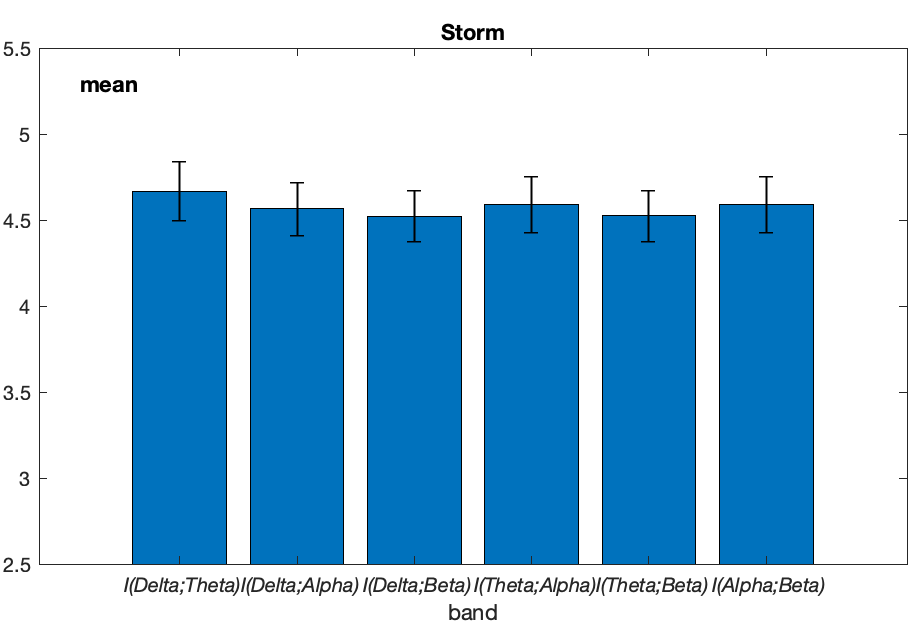

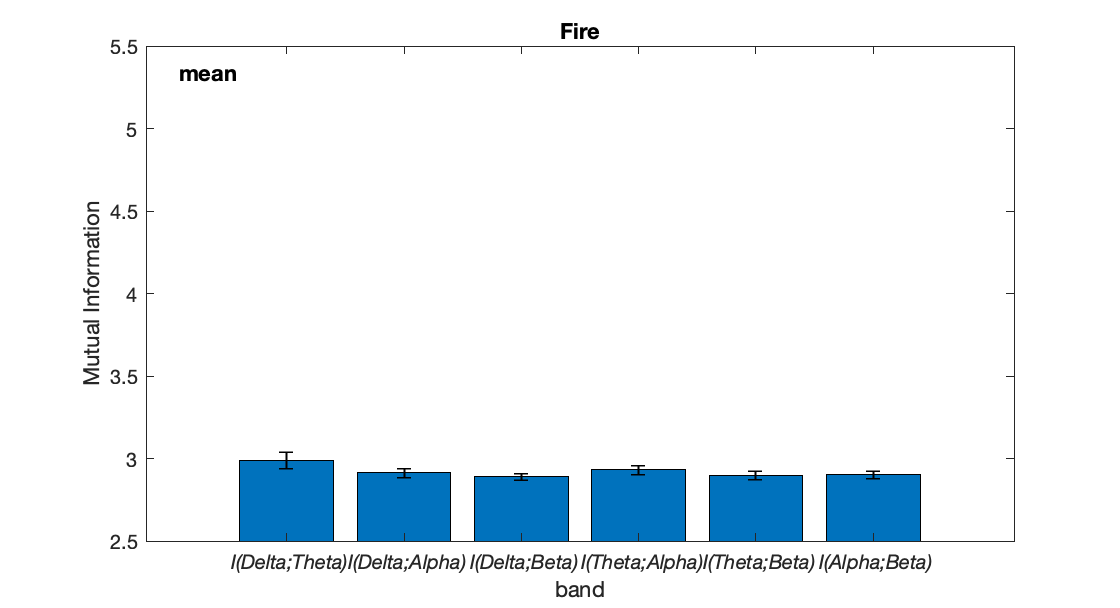

**Figure b**. Mutual information of nature sounds in S-AMPH model

**Mutual Information Between Adjacent Oscillatory Bands in PAD model**

Figure **c** shows the mutual information (MI) between different bands in PAD model for each genre. The MI shown here are broadly similar across the 10 genres: adjacent bands are predictable each other compared with non-adjacent band. Figure **d** shows the MI of non-music natural sounds such as rain and wind.

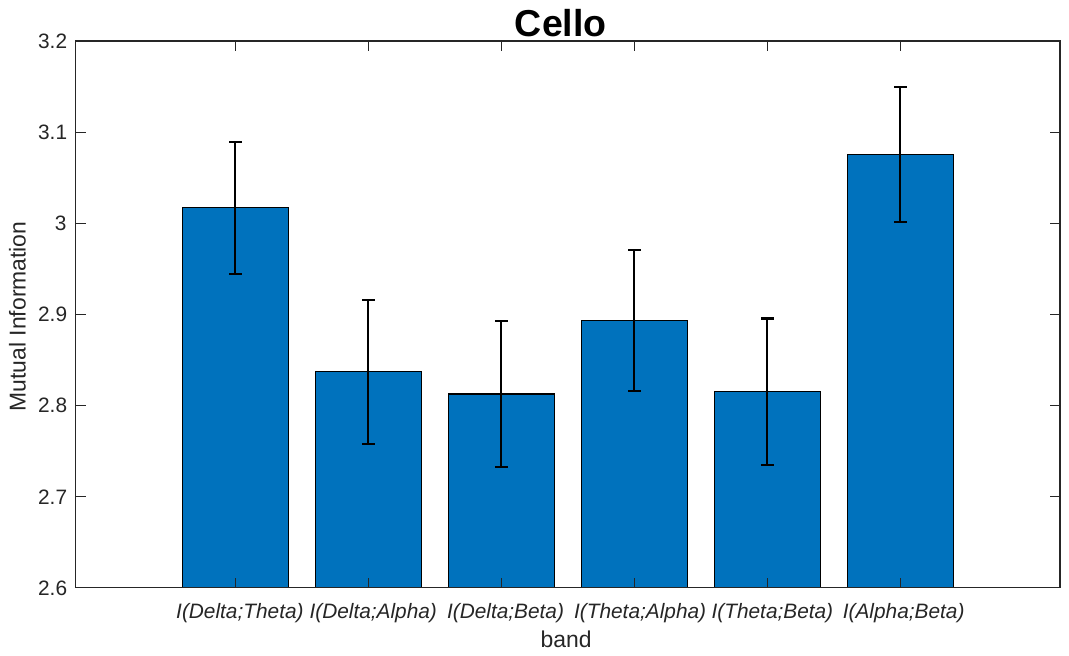

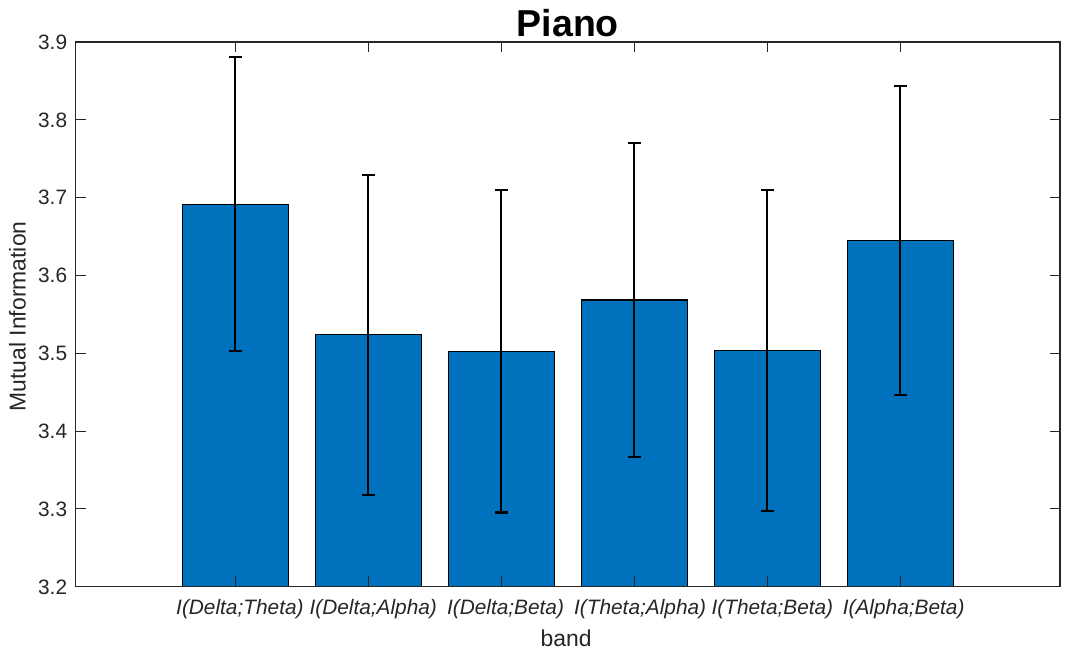

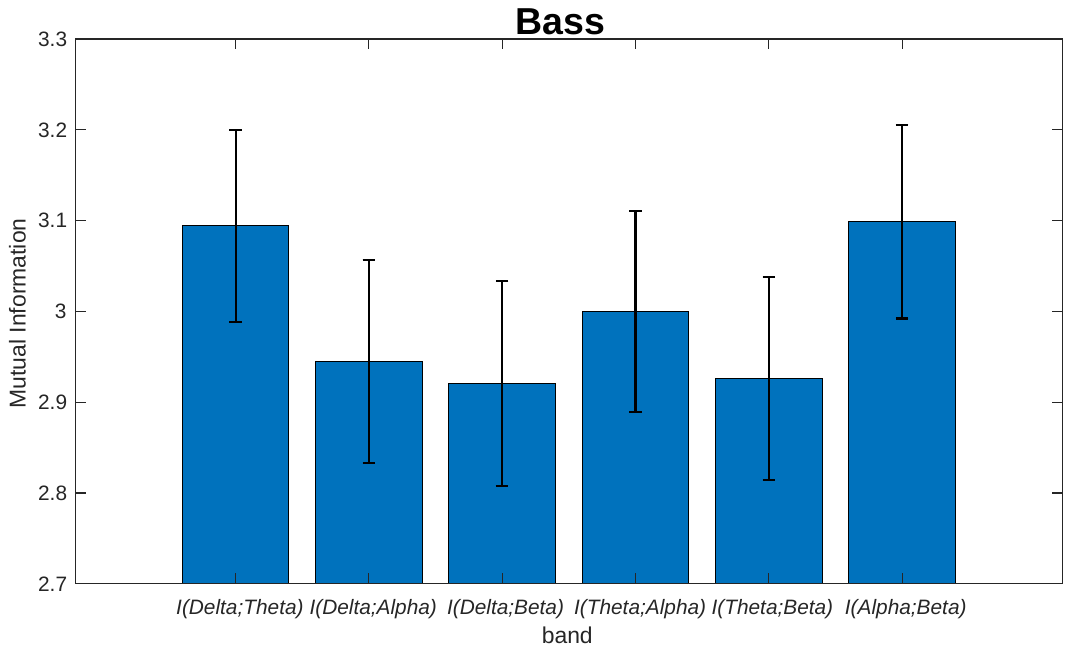

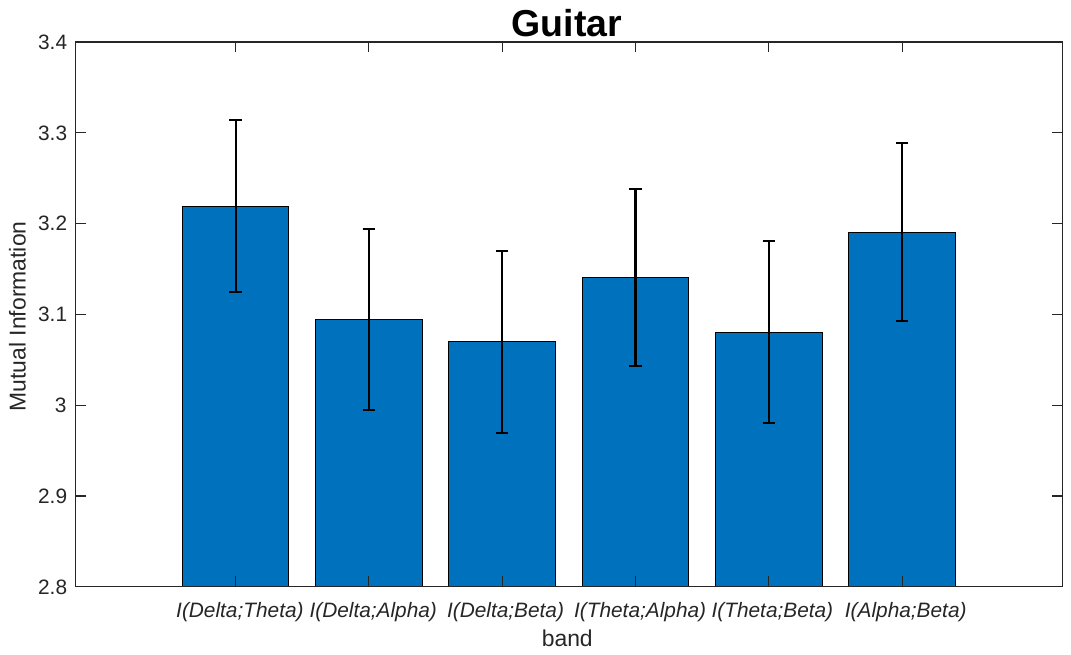

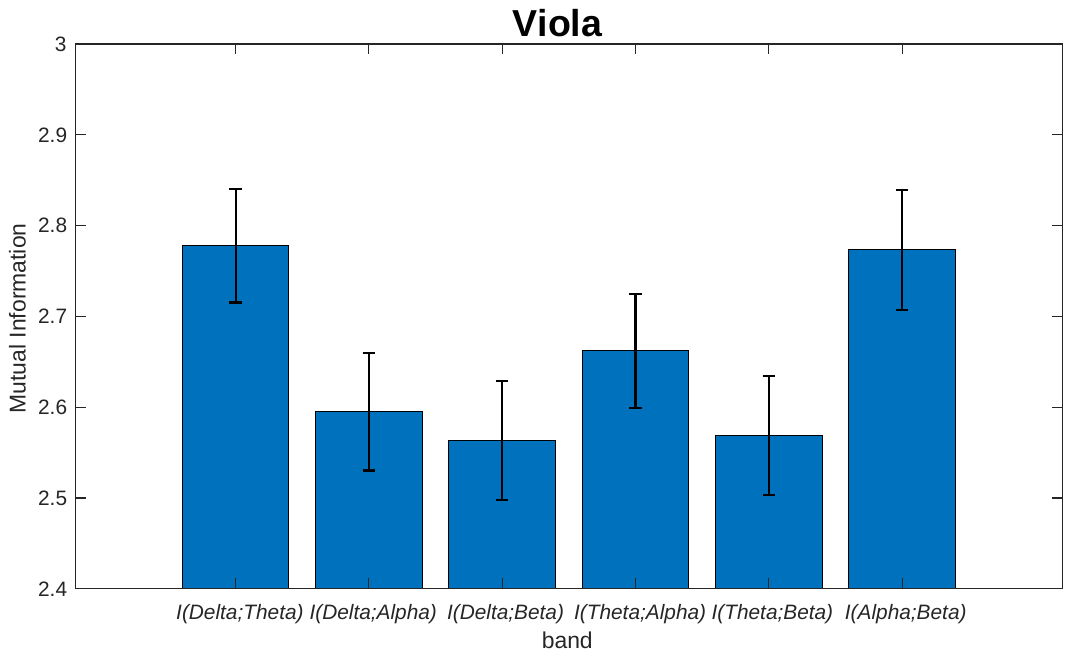

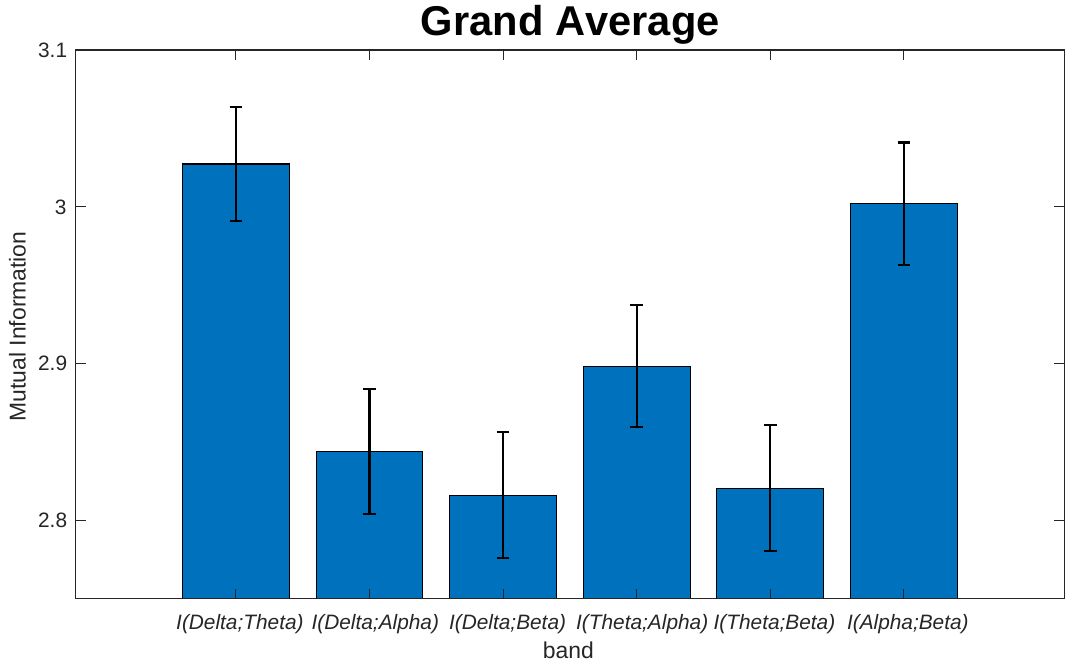

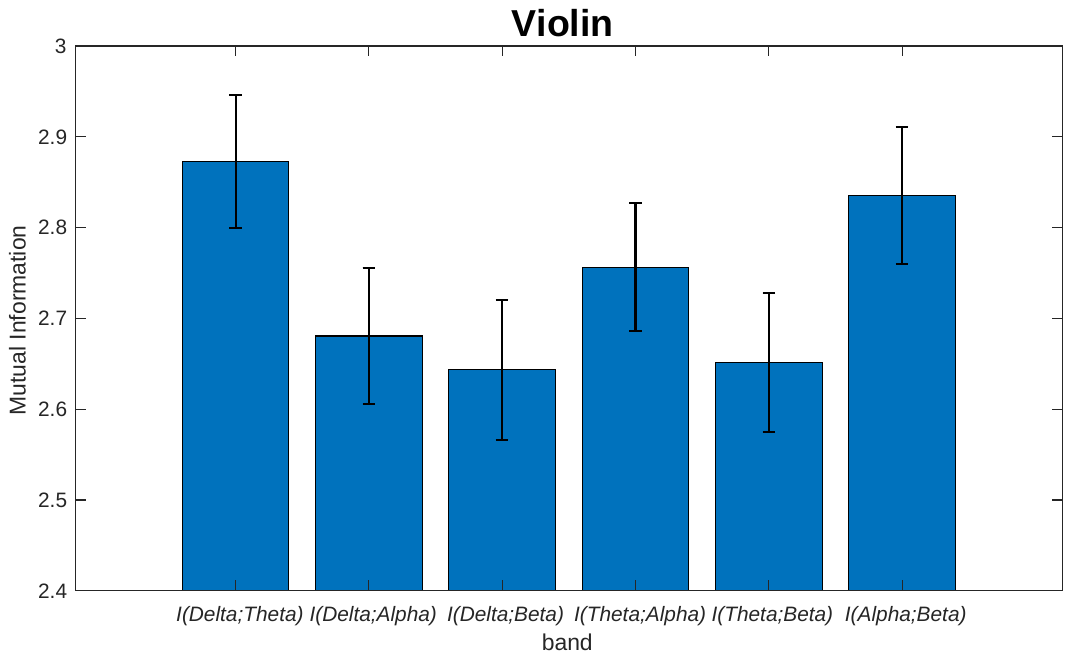

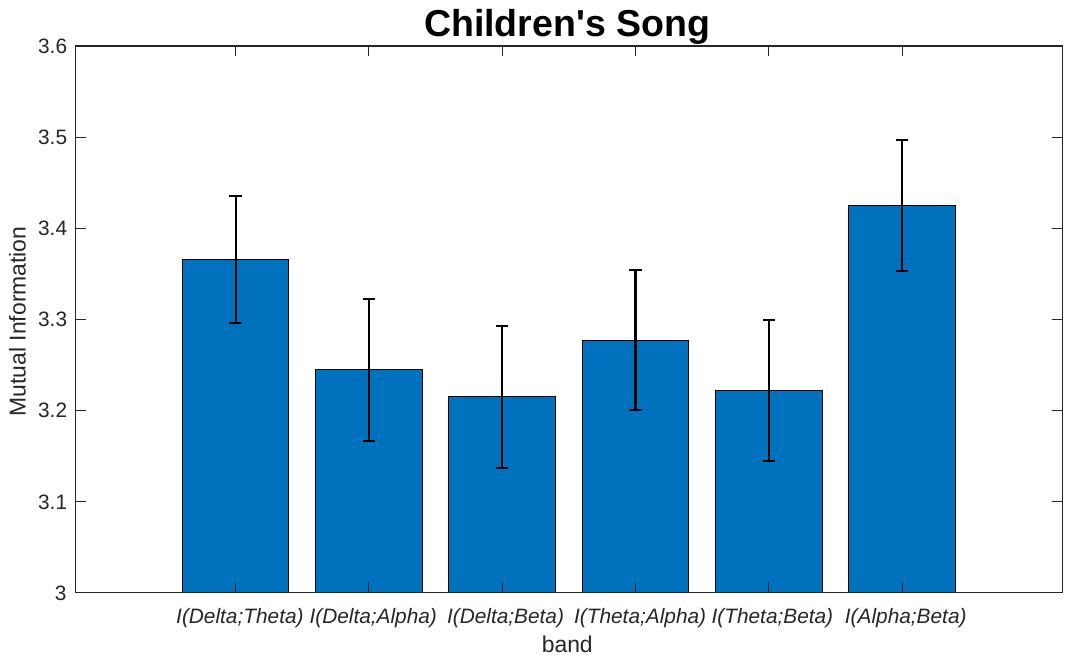

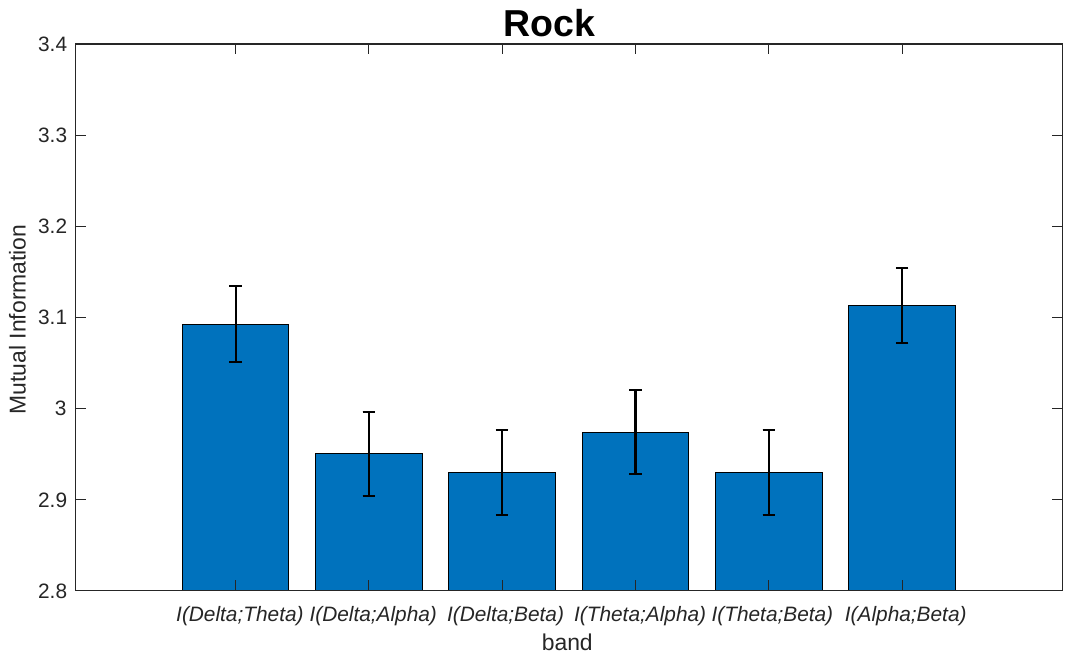

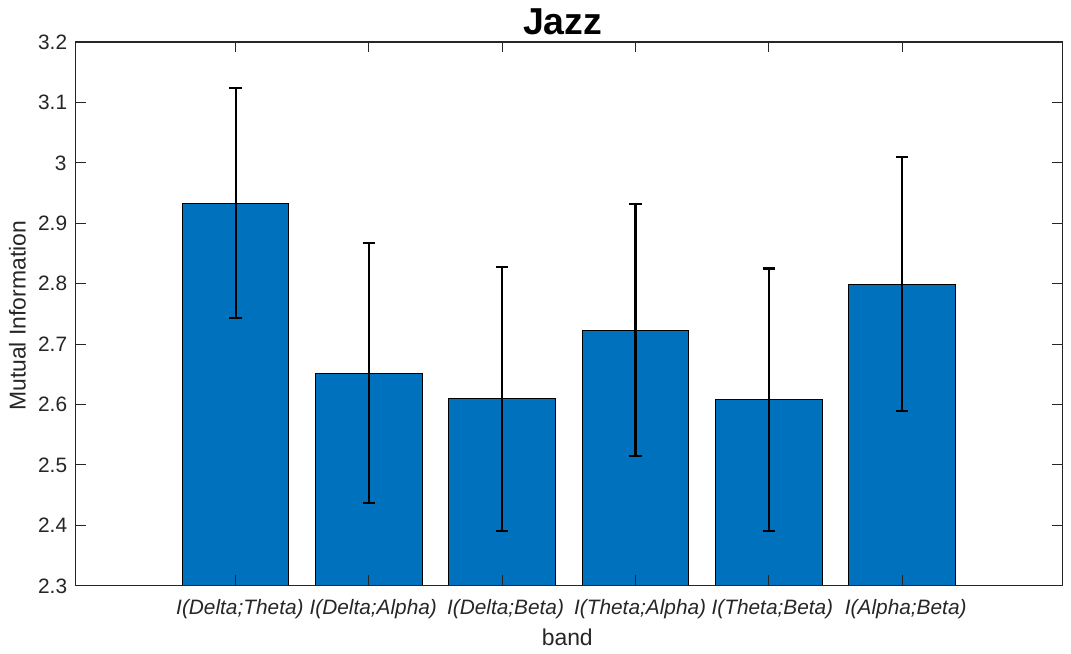

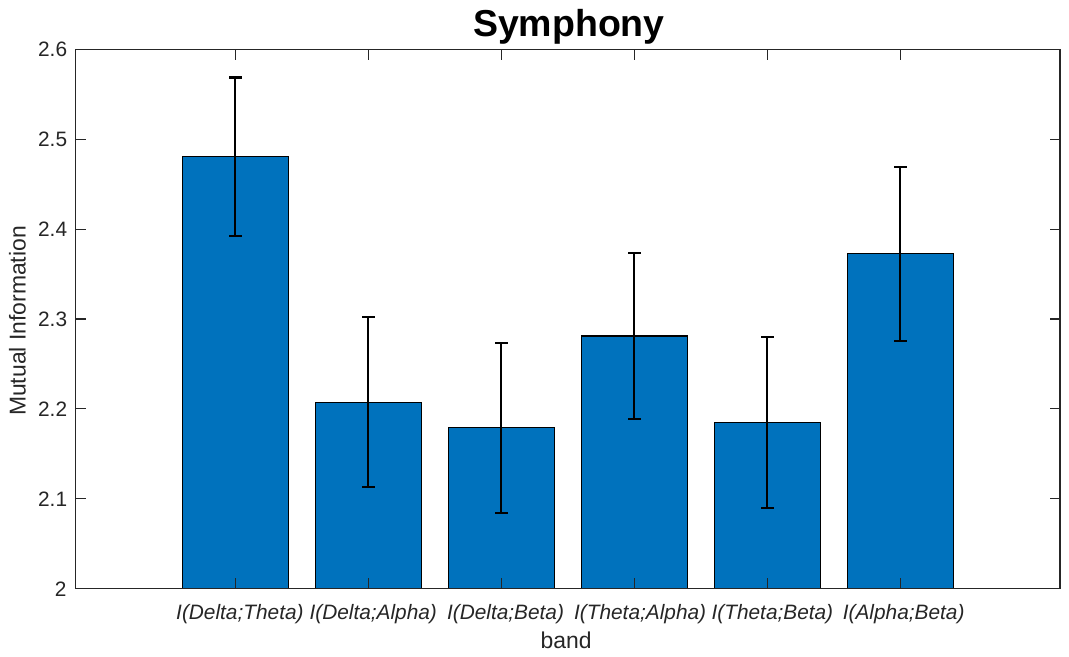

**Figure c**. Mutual information between different bands in PAD model

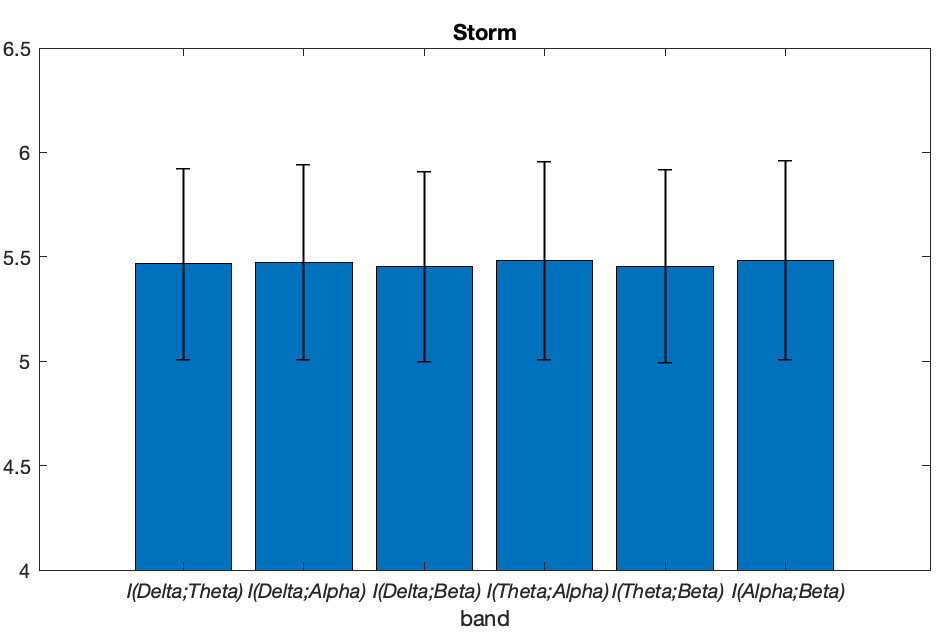

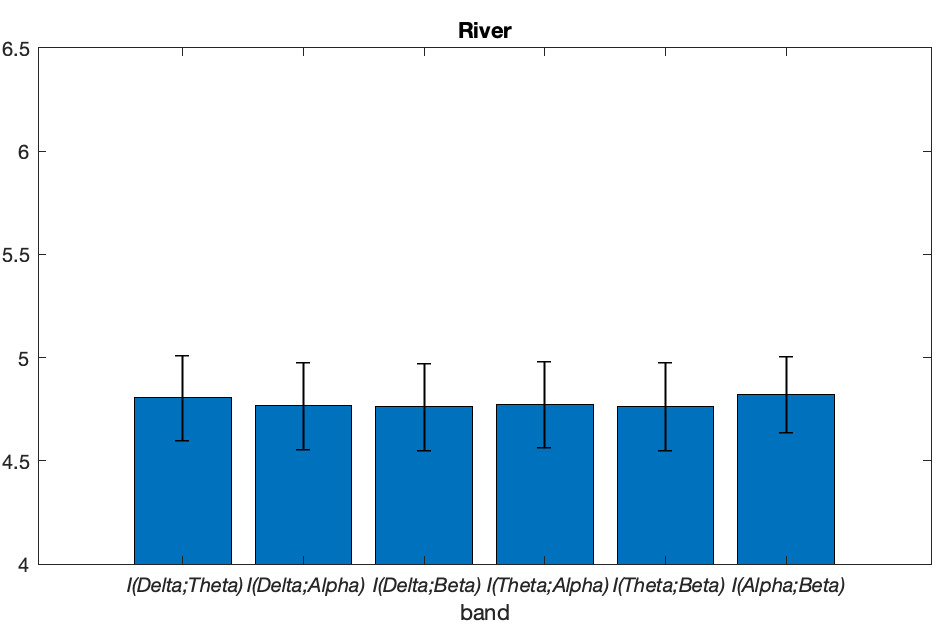

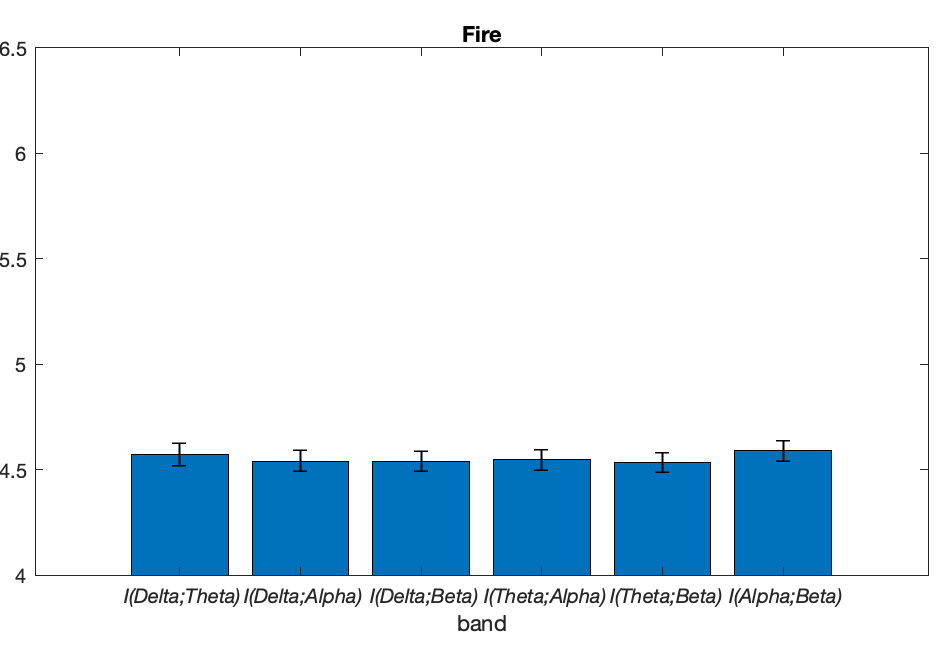

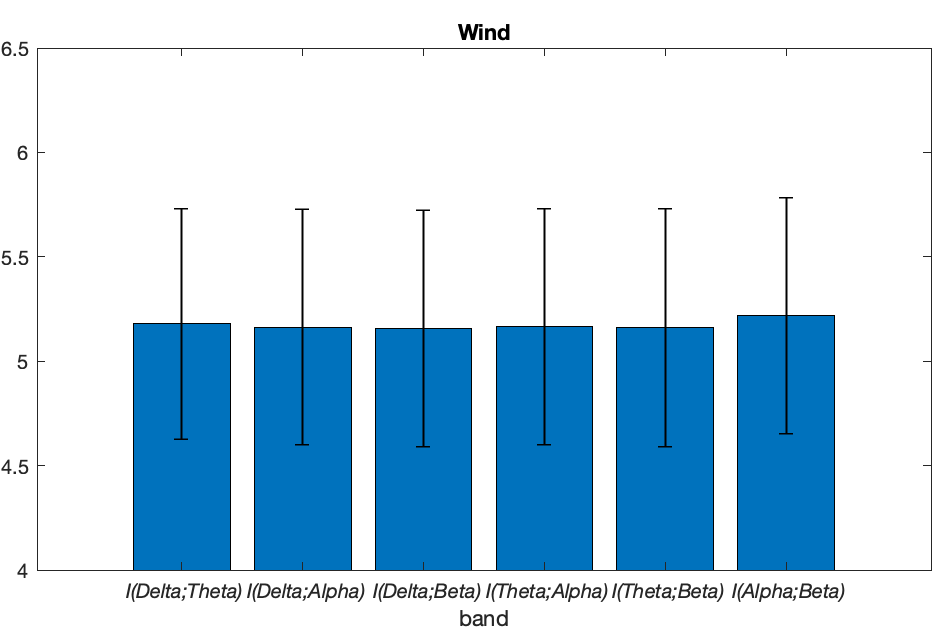

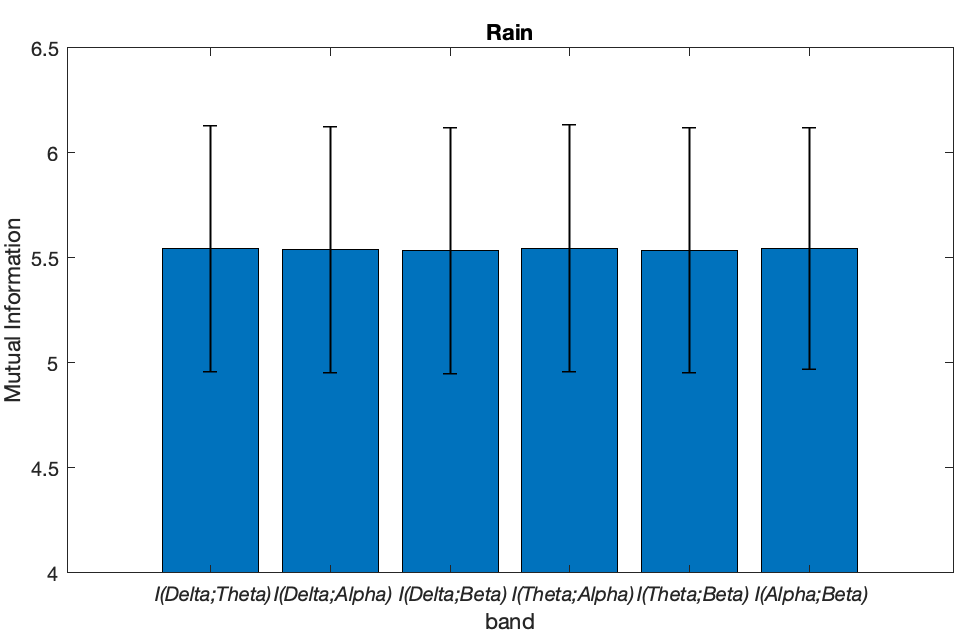

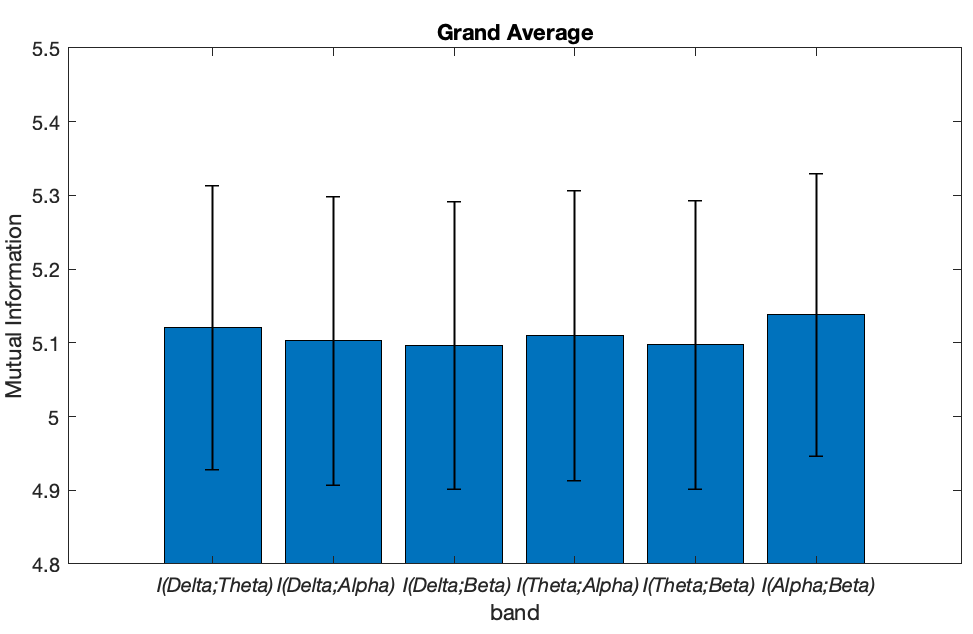

**Figure d**. Mutual information of nature sounds in PAD model
