## Supplementary material for "The Temporal Statistics of Musical Rhythm across Western Genres: An Amplitude Modulation Phase Hierarchy Model": S5 Appendix (from another version of this preprint: version 2, 2021-04-01)

**Averaged PSI in Each Integer Ratio by S-AMPH model**

Figure **a** shows the averaged PSI in each integer ratio by S-AMPH model for each genre. In each subplot, the lines of different thickness indicate different comparisons in oscillatory bands. The PSI shown here are broadly similar across the 10 genres and across different comparisons in oscillatory bands. It may be observed that the 10 genres produced broadly consistent PSIs.

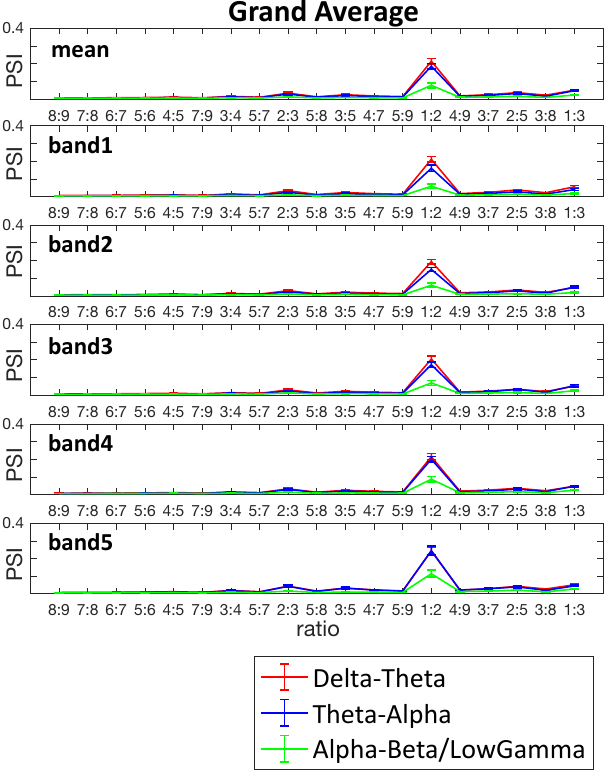

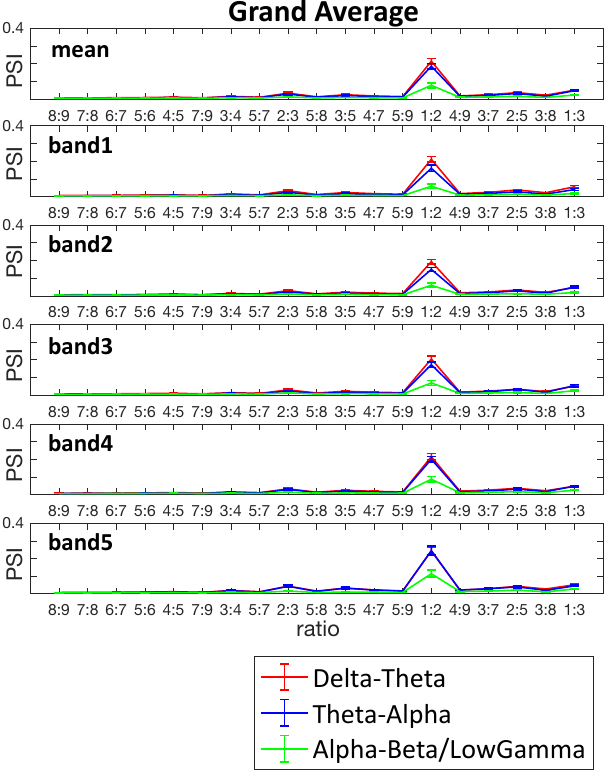

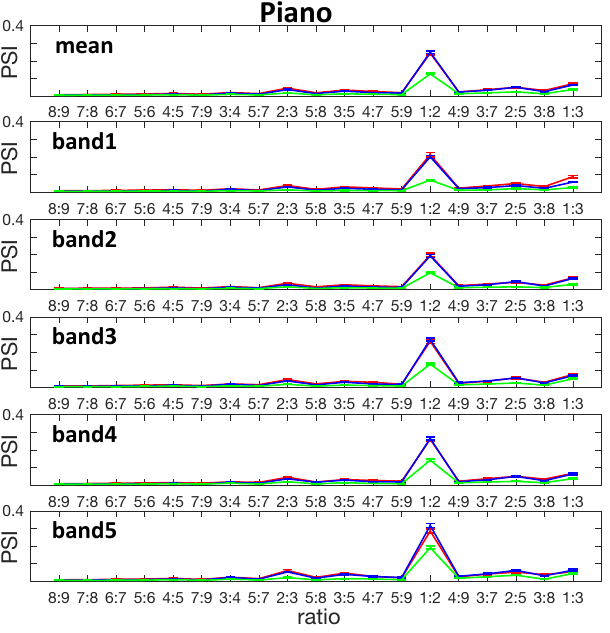

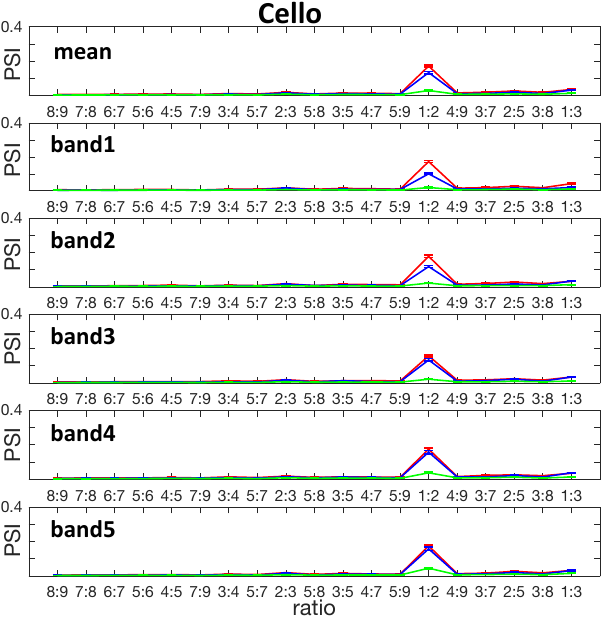

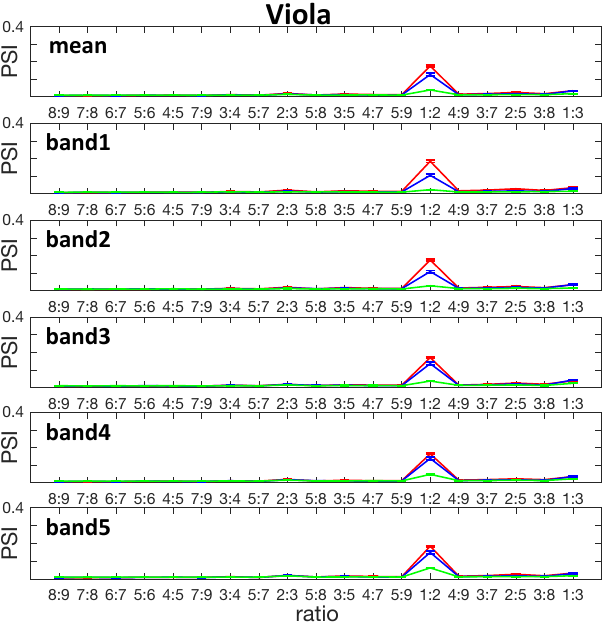

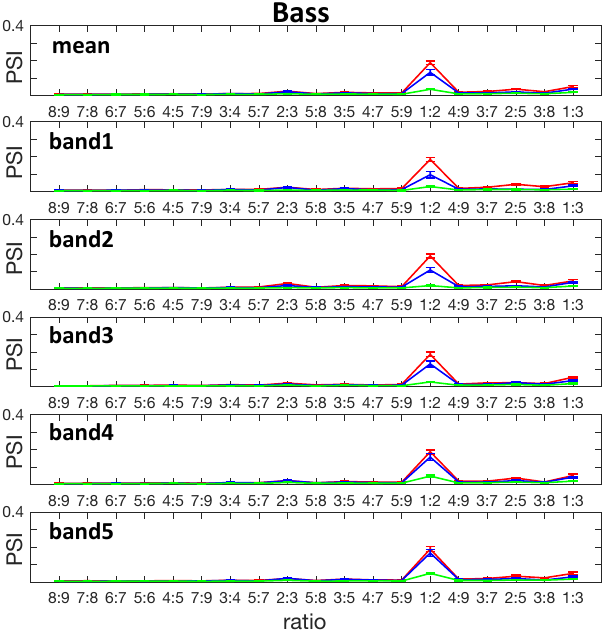

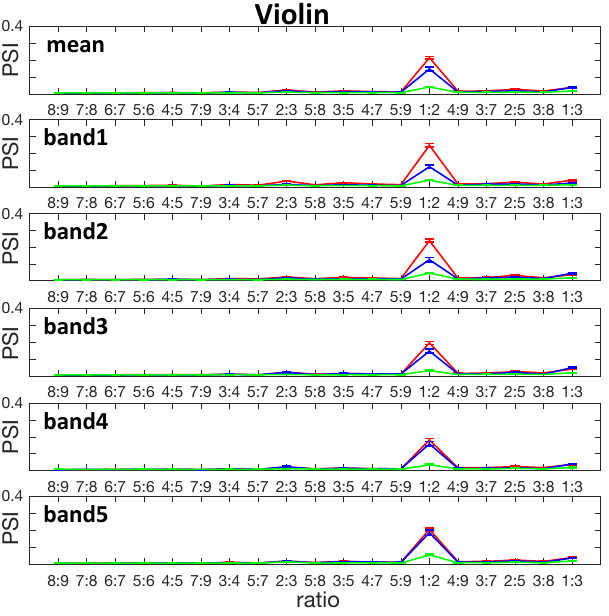

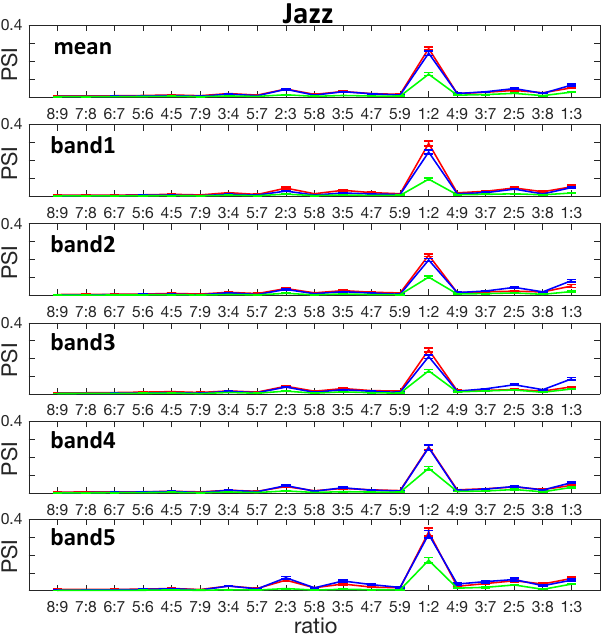

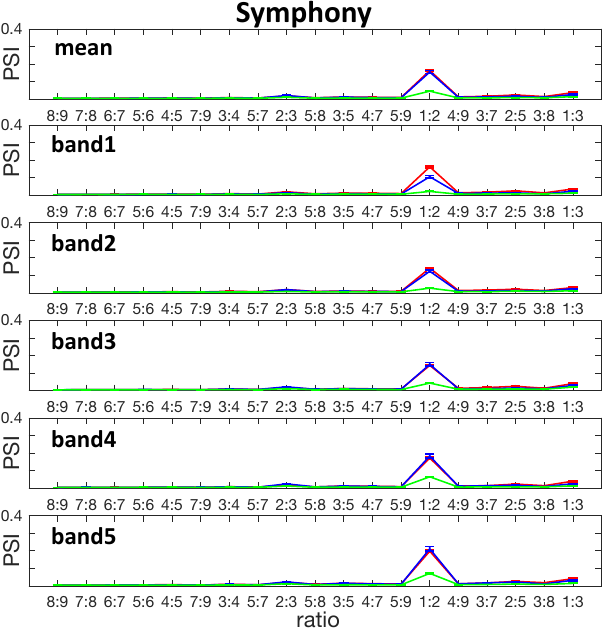

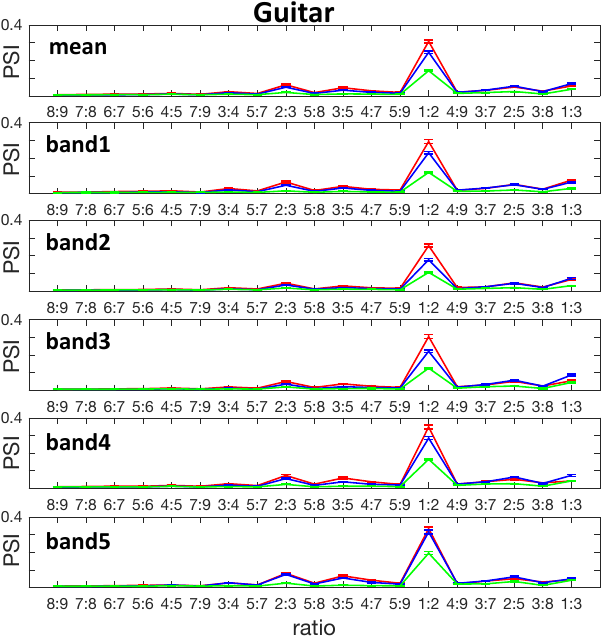

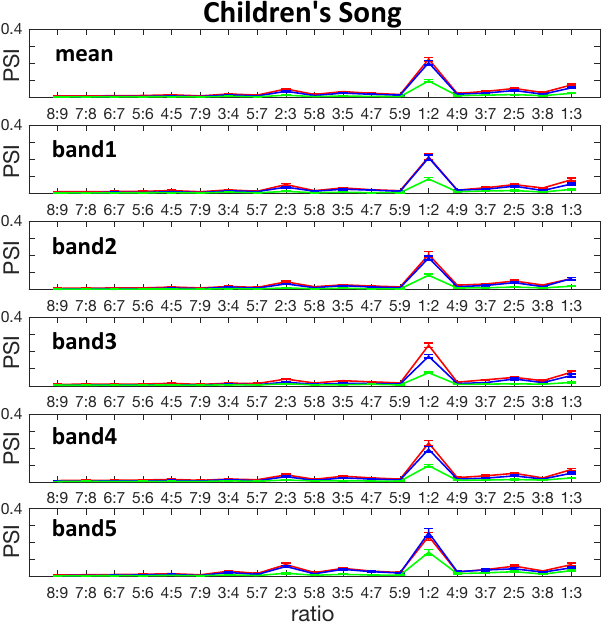

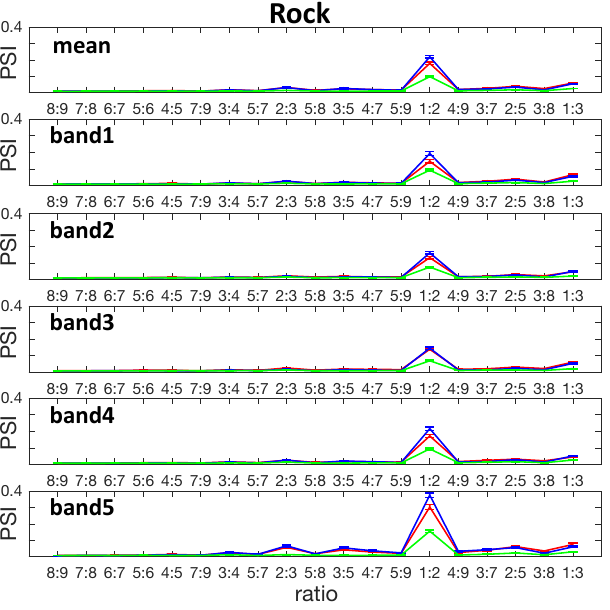

**Figure a**. Individual spectral PCA component loading patterns

Figure **b** shows the averaged PSI in each integer ratio by S-AMPH model for each genre of nature sounds. In each subplot, the lines of different thickness indicate different comparisons in oscillatory bands.

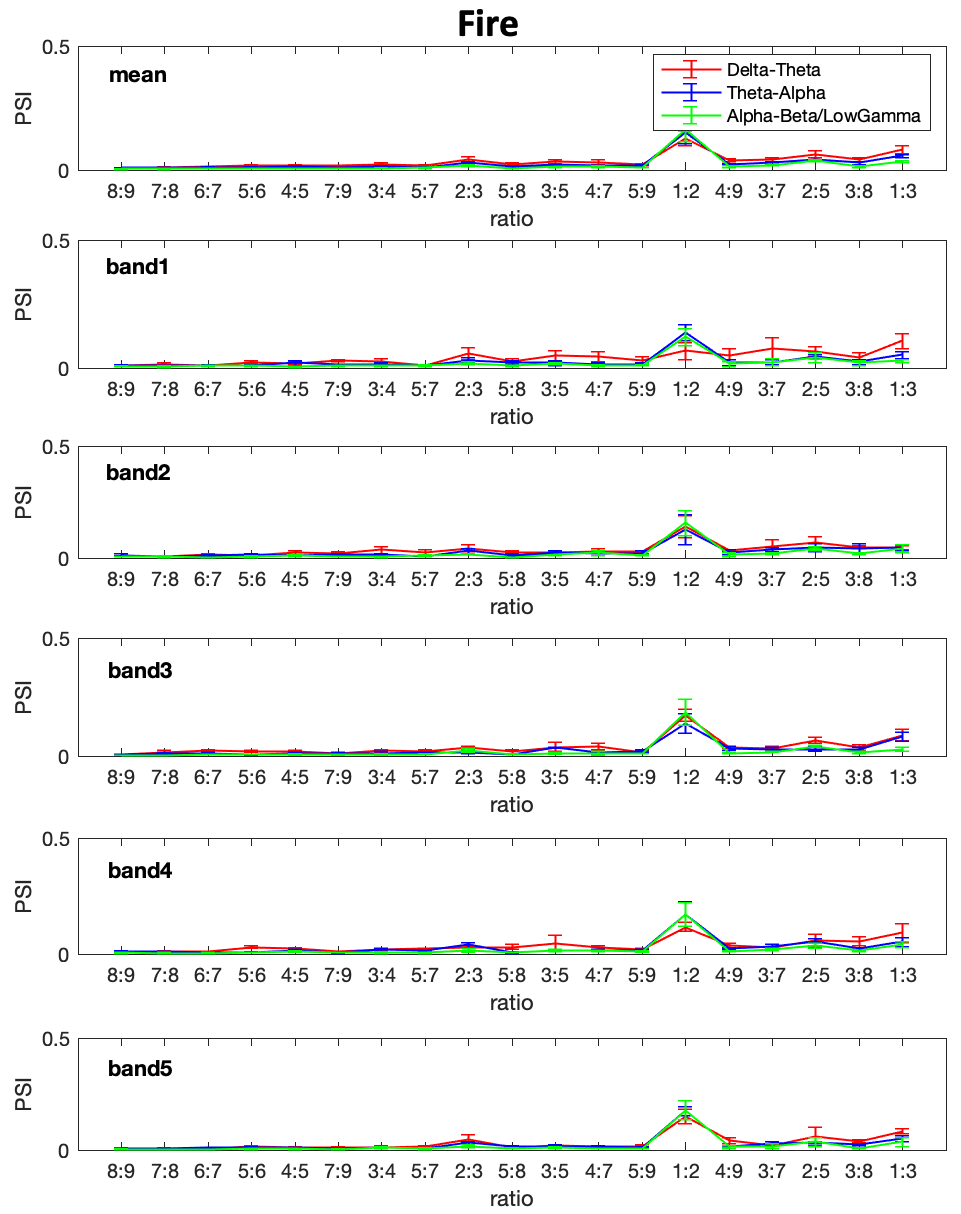

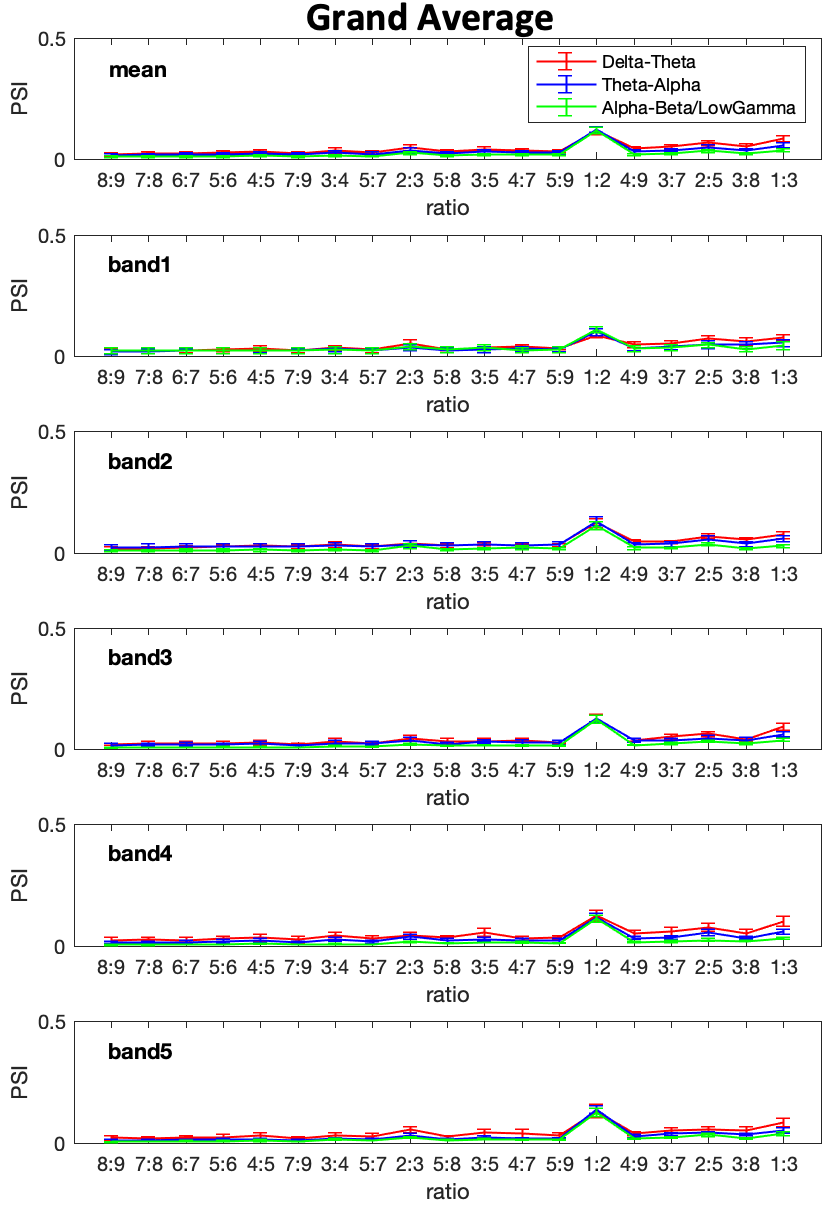

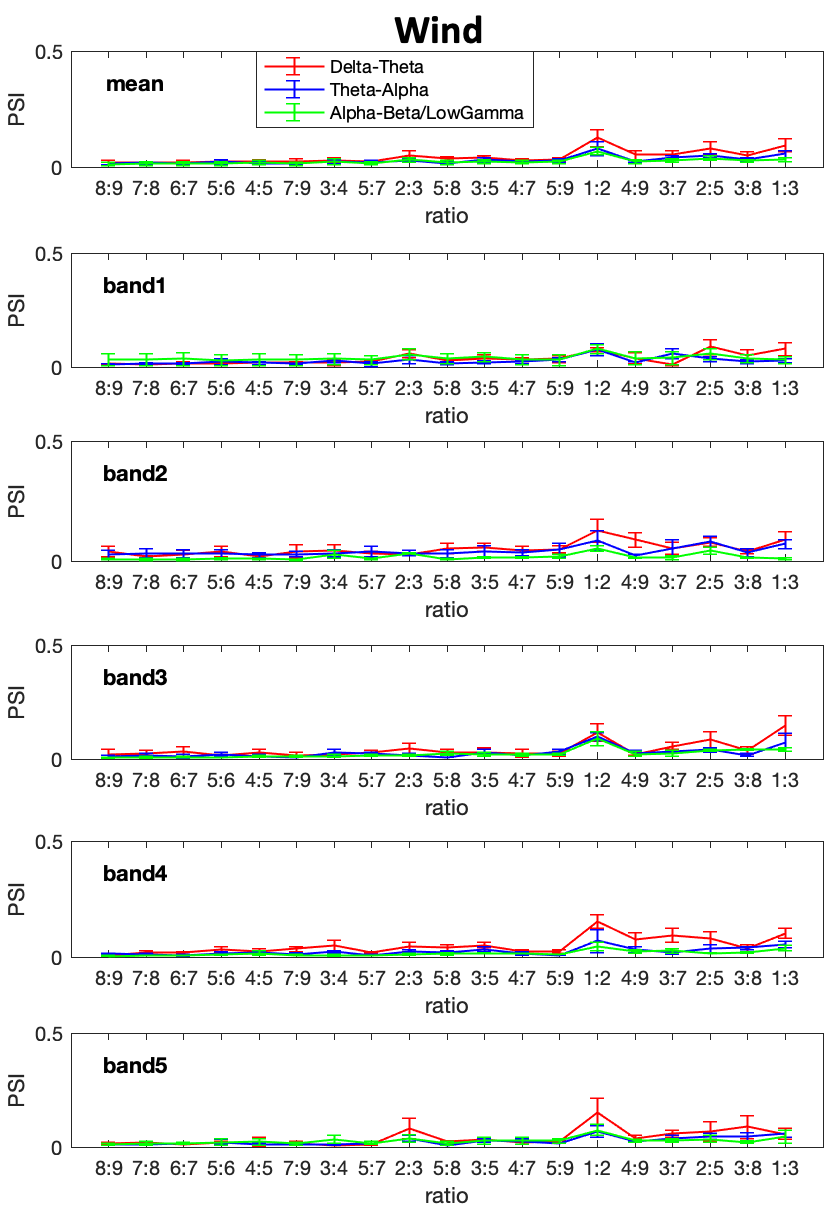

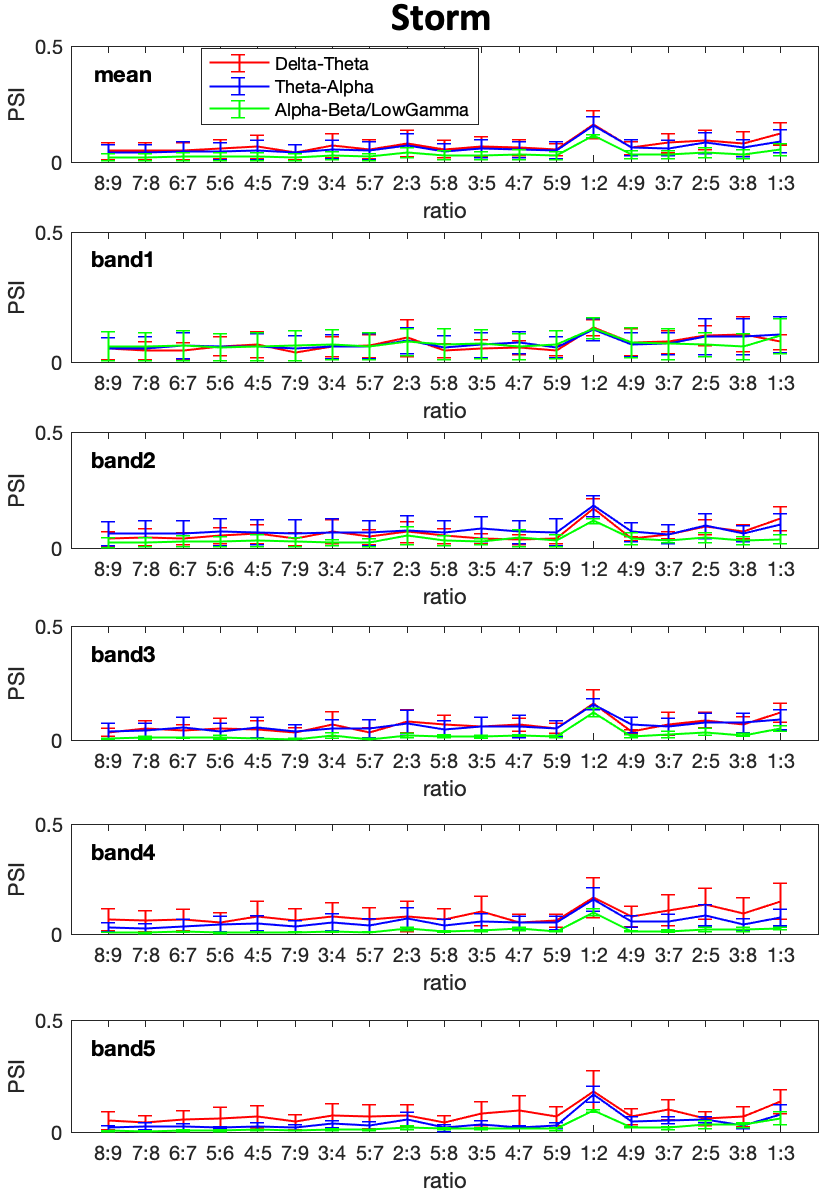

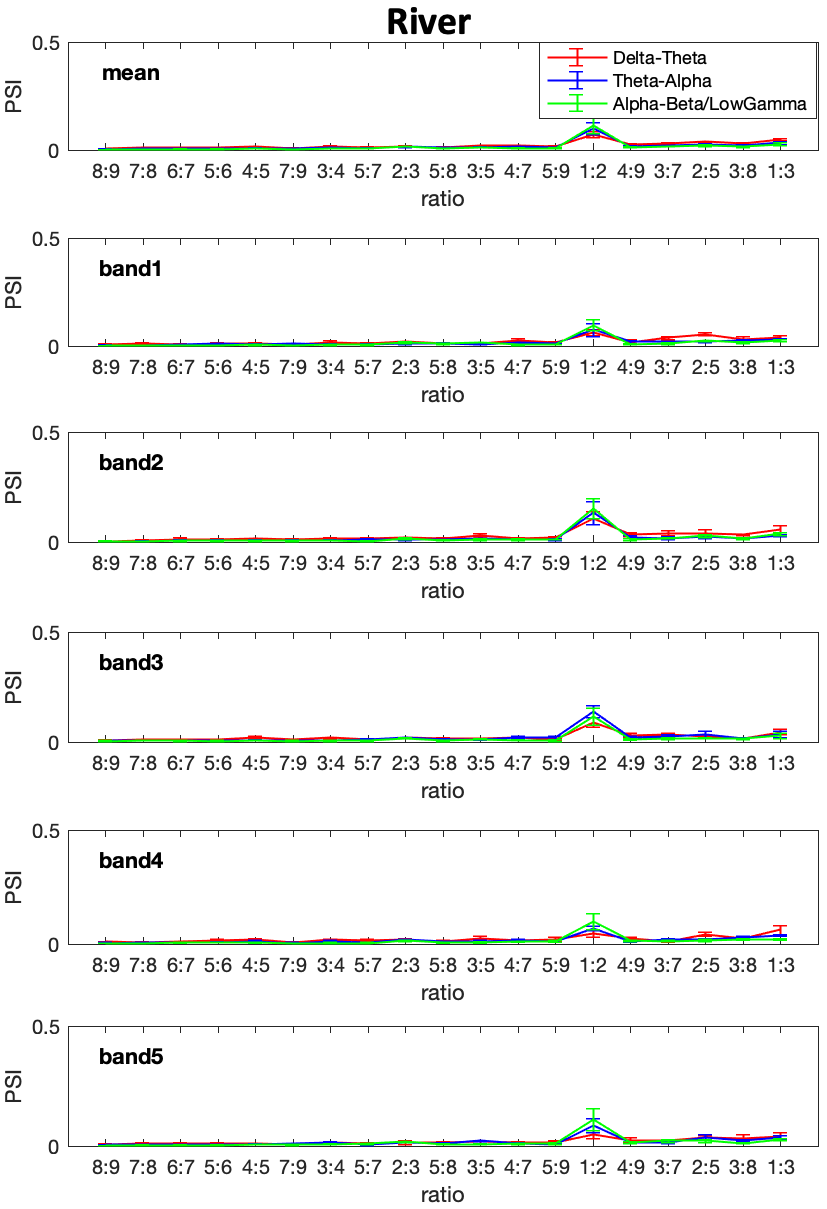

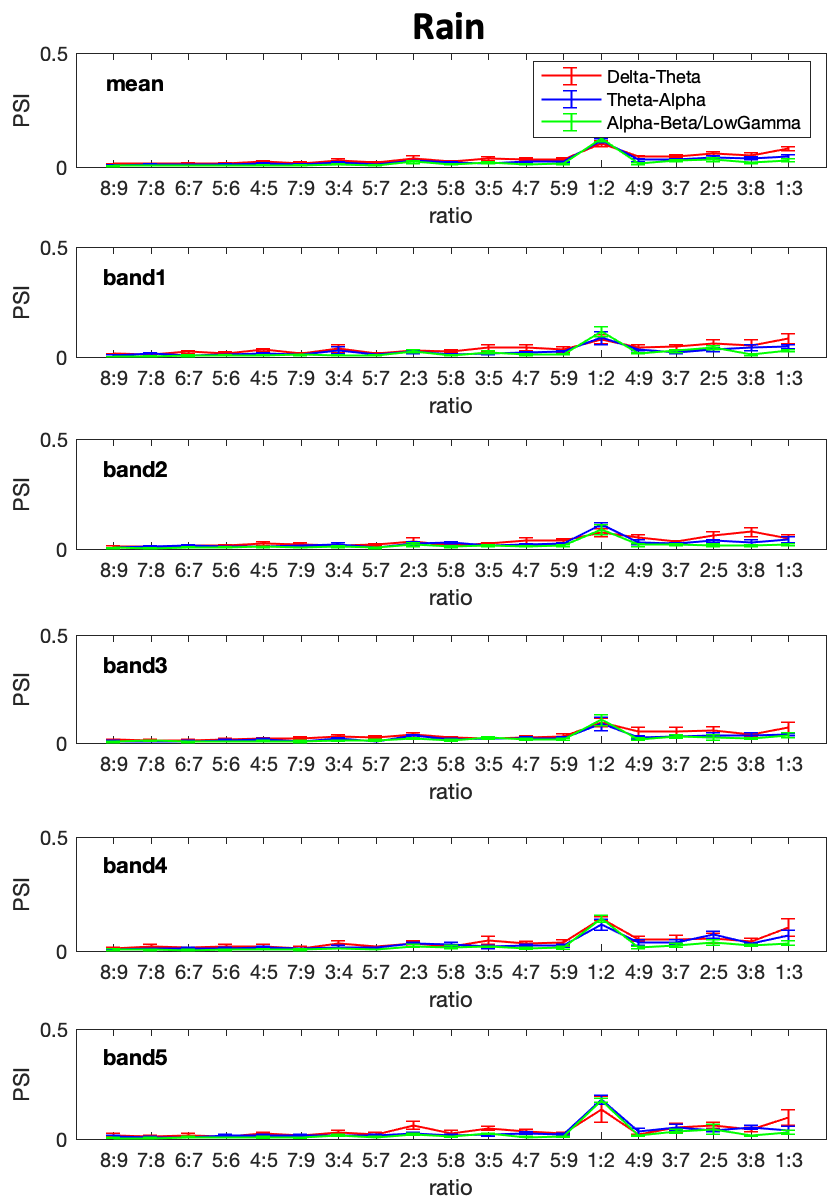

**Averaged PSI in Each Integer Ratio by PAD model**

Figure **c** shows the averaged PSI in each integer ratio by PAD model for each genre. In each subplot, the lines of different thickness indicate different comparisons in oscillatory bands. The PSI shown here are broadly similar across the 10 genres and across different comparisons in oscillatory bands. It may be observed that the 10 genres produced broadly consistent PSIs.

**Figure b.** Individual modulation spectra of FFT

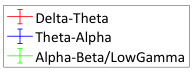

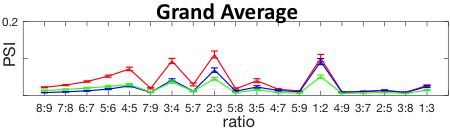

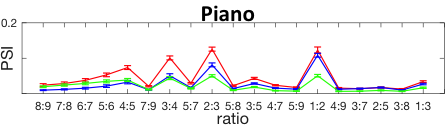

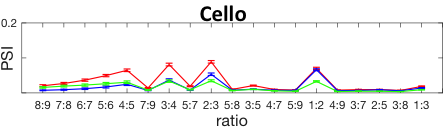

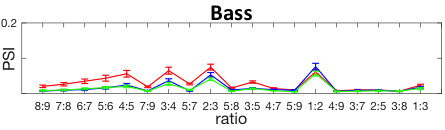

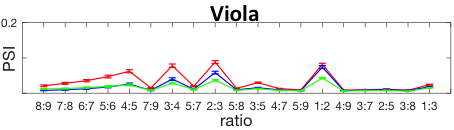

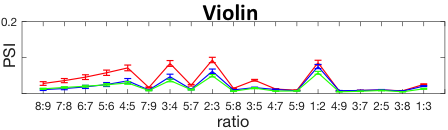

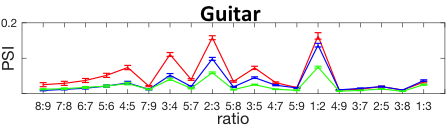

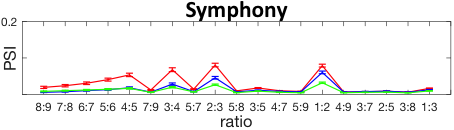

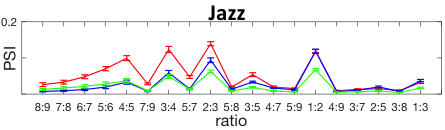

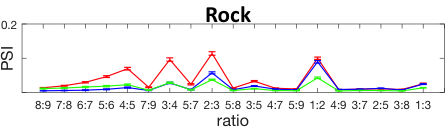

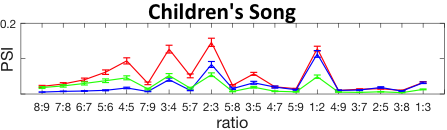

Figure **d** shows the averaged PSI in each integer ratio by PAD model for each genre of nature sounds. In each subplot, the lines of different thickness indicate different comparisons in oscillatory bands.

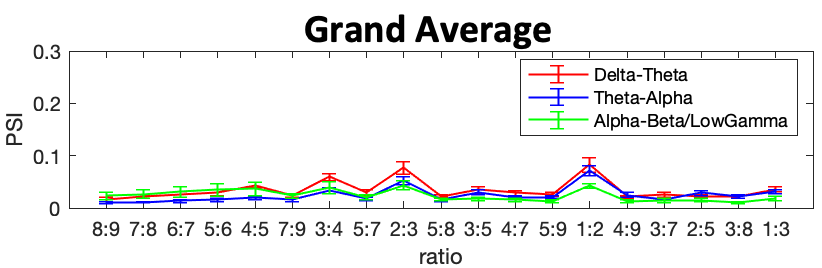

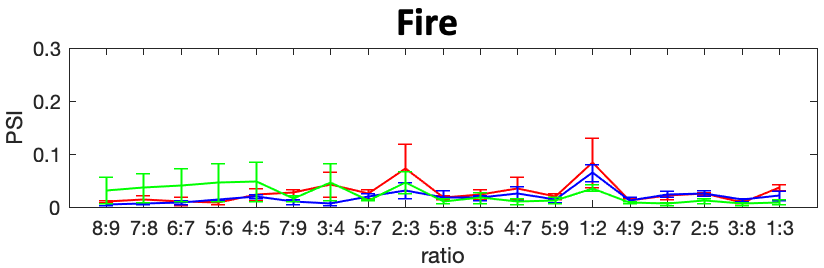

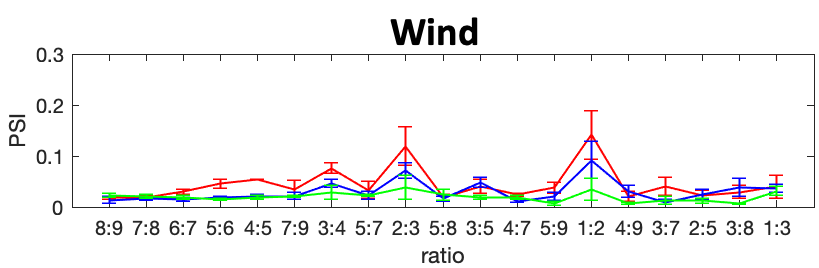

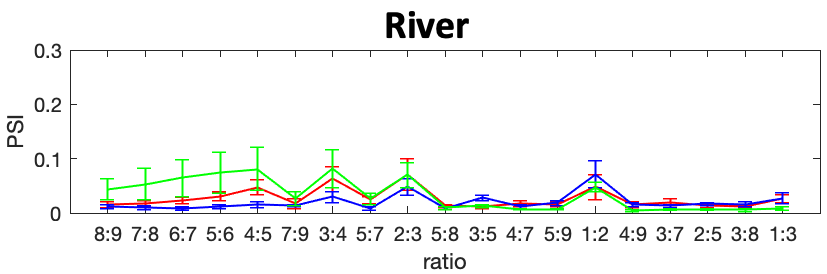

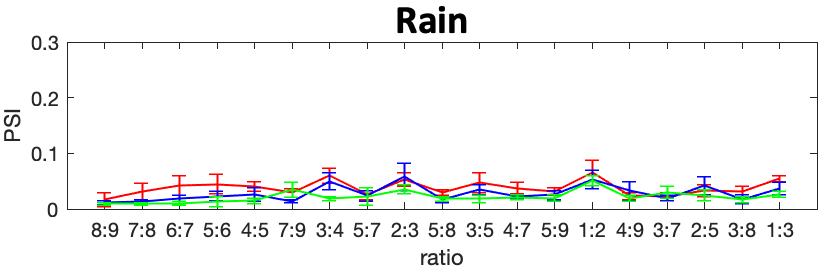

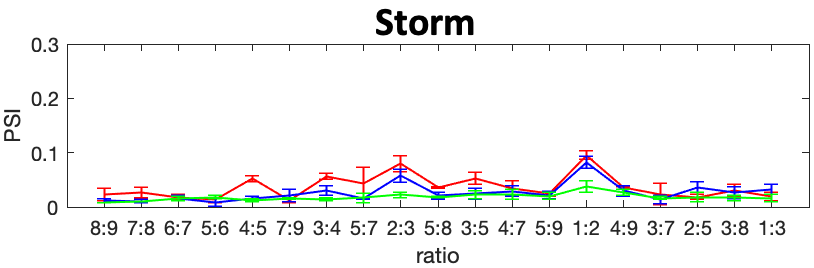

**Simulation results of PSI**

**Figure c**. FFT after the PAD.

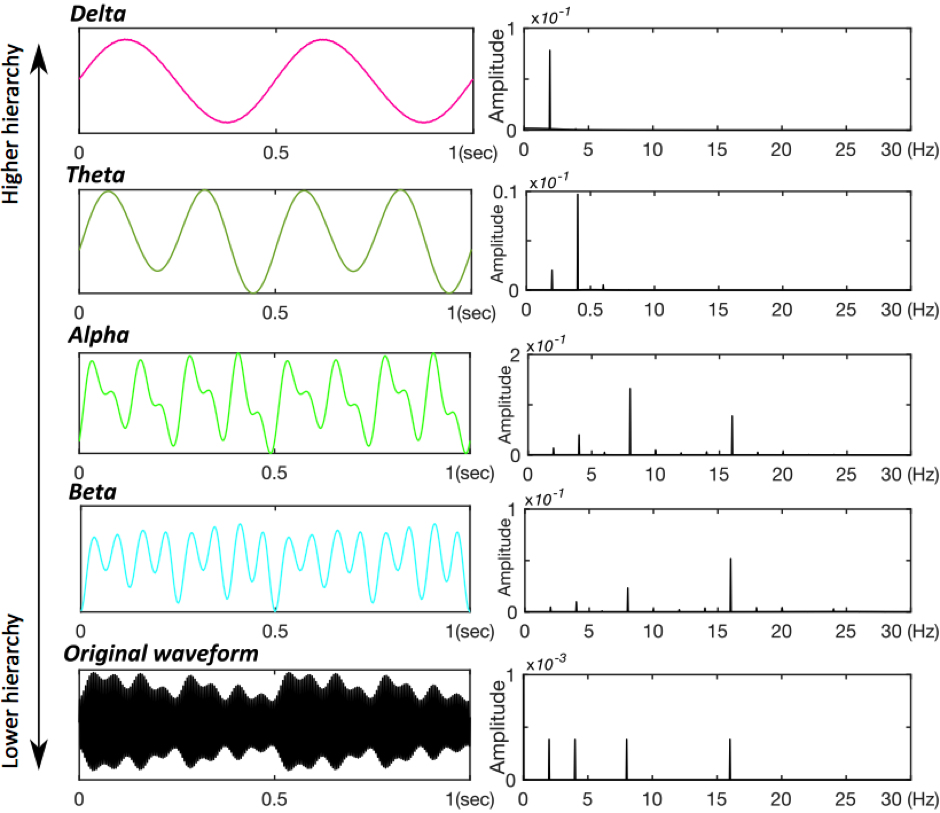

**Figure d**. PSI results

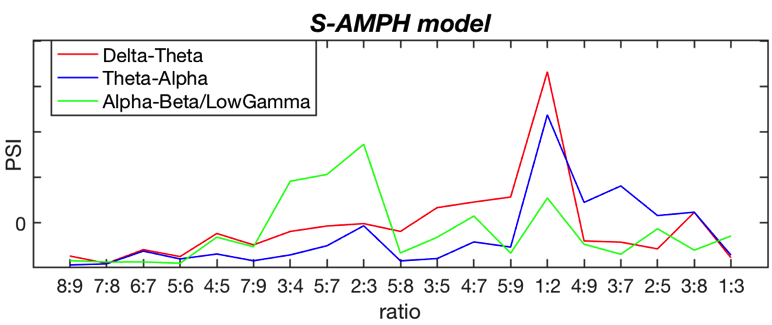

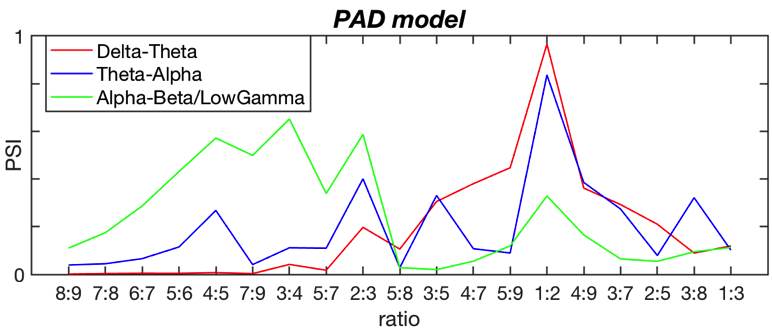
